# Biologically grounded cell profiling across microscopy modalities

**DOI:** 10.64898/2026.09.23.753678

**Authors:** Enze Ye, Xiaoxuan Wu, Rui Peng, Wenjia Hu, Xiangyou Li, Xuefei Zhang, Mengxiao Niu, Yaorong Guo, Xinlei Sheng, Jinzhuo Wang, Liangyi Chen, He Sun

## Abstract

Microscopy-based cell profiling has broad applications in biological discovery, disease characterization, and phenotypic drug screening. Modern microscopy continues to push the limits of resolution, speed, depth and throughput, but better imaging does not automatically lead to better biomedical discovery and translation. A key bottleneck is feature representation: existing features are either handcrafted or learned as black-box embeddings and often lack explicit biological meaning. Here we propose biological grounding as a first principle for cell profiling and implement it through MorphAgent, an AI agent framework in which each quantitative feature originates from a biological hypothesis and is anchored to a cellular structure or process. By integrating biological knowledge, multimodal reasoning and automated validation, MorphAgent makes biologically grounded feature design systematic and scalable. We show biological grounding fundamentally changes the properties of cell profiling features. Across three microscopy modalities, MorphAgent produces compact, expressive and transferable features for drug screening, cell state characterization and disease morphology analysis. In wide-field Cell Painting, a common assay for phenotypic drug screening, it improves perturbation-retrieval mean average precision by 48% over CellProfiler and 20% over DeepProfiler while using substantially fewer dimensions. In confocal mitochondrial imaging, biologically grounded features transfer across independently acquired datasets and imaging resolutions, supporting accurate aging-state classification. In structured illumination microscopy (SIM) imaging of Tau-labeled samples, biologically grounded super resolution features reveal nanoscale morphologies inaccessible at conventional resolution, enhancing the discrimination of disease-associated mutations. Moreover, biologically grounded features provide a hierarchical organization of cellular morphology, enabling multilevel alignment with transcriptomics data and facilitating mechanistic understanding. Biologically grounded cell profiling thus brings AI reasoning closer to biological mechanisms, advancing biomedical discovery and translation.

## Introduction

Microscopy-based cell profiling aims to decode cellular state by transforming cellular images into quantitative representations that reflect biological processes, perturbation responses and disease-associated phenotypes^1–4^. Over the past decade, standardized assays such as Cell Painting^5^ have enabled large-scale profiling of cellular responses to genetic and chemical perturbations, supporting mechanism-of-action (MoA) inference and phenotypic drug screening. Meanwhile, advances in microscopy continue to expand what can be observed: structures that were previously blurred become separable, puncta emerge from diffuse signal, bundles resolve from aggregates, and spatial organization becomes measurable across cellular compartments^6,7^. However, better images do not automatically yield better biomedical discovery or translation. Cell profiling must therefore evolve with microscopy through a generalizable feature design strategy.

Existing cell profiling approaches address feature design through two dominant paradigms. Handcrafted methods, such as CellProfiler^8–11^, describe cells using predefined features of intensity, shape, texture and spatial organization. Although effective for standardized high-content assays, these features are limited by what experts can specify in advance and are difficult to adapt to phenotypes that emerge in new biological systems or imaging modalities. Deep learning methods^12–15^, such as DeepProfiler^16^, can capture richer image variation through learned embeddings, but their features depend on the quality and diversity of training data and often lack explicit biological semantics. Both paradigms share a common limitation: the feature space is not biologically grounded by design. As a result, their features may not fully capture predictive information in the images, generalize poorly across imaging conditions and provide limited biological insight.

To address these limitations, we propose biologically grounded cell profiling. Rather than extracting or learning image features first and interpreting them afterwards, biological grounding builds meaning into each feature by construction. Each quantitative feature begins with a biological hypothesis and is anchored to a cellular structure or process and a defined spatial scale. This mirrors how cell biologists interpret microscopy images, making the features highly expressive, transferable across imaging modalities, and suitable for mechanistic understanding.

Systematically constructing biologically grounded features, however, cannot be achieved through human expert curation, because human knowledge may be incomplete or biased and many visual features are difficult to quantify. We develop MorphAgent, an AI agent framework that generates biologically grounded feature hypotheses through automated literature review, quantifies them using coding agents or vision-language models, and iteratively selects and validates features based on feature statistics and feedback from paired omics. MorphAgent thus provides a practical solution for scalable biologically grounded feature design.

Across wide-field, confocal and super resolution microscopy, we show that biological grounding fundamentally changes the properties of cell profiling features. First, biologically grounded features are compact yet expressive. In high-content wide-field chemical perturbation screening, 467 biologically grounded features improve perturbation retrieval by 48% over 6,256 CellProfiler features and by 20% over 672 DeepProfiler embeddings. The same advantage extends to MoA assignment and the matching of compounds sharing the same MoA, supporting phenotypic drug screening. Second, biologically grounded features are transferable. In confocal mitochondrial imaging, biologically grounded feature vocabulary designed from one HSC dataset transfers to an independently acquired dataset with higher image quality, supporting accurate cell state classification. Finally, biologically grounded features are resolution-aware and can therefore be hierarchically organized. In structured illumination microscopy (SIM) of Tau-labeled samples, these features capture fine-scale phenotypes resolved only by super resolution imaging, enabling accurate disease-associated mutation classification and bidirectional prediction between cellular morphology and single-cell transcriptomic programs. Their hierarchical organization, from individual features to subcellular compartments and morphology-defined biological processes, further aligns with multilevel gene ontology (GO) modules, enabling mechanistic understanding across scales. These results show how biologically grounded cell profiling can convert advances in microscopy into both biomedical discovery and translation.

## Results

### MorphAgent designs biologically grounded features for cell profiling across microscopy modalities

MorphAgent takes three inputs: an interactive dataset environment (microscopy images, documentation, channel and modality information and paired metadata), the biological question posed by the user, and an open-world knowledge base (expert knowledge, literature knowledge and automated deep research^17^). It uses these inputs in an iterative loop of planning, quantification and validation to design biologically grounded features (Fig. 1a, Supplementary Fig. 1).

**Figure 1.**
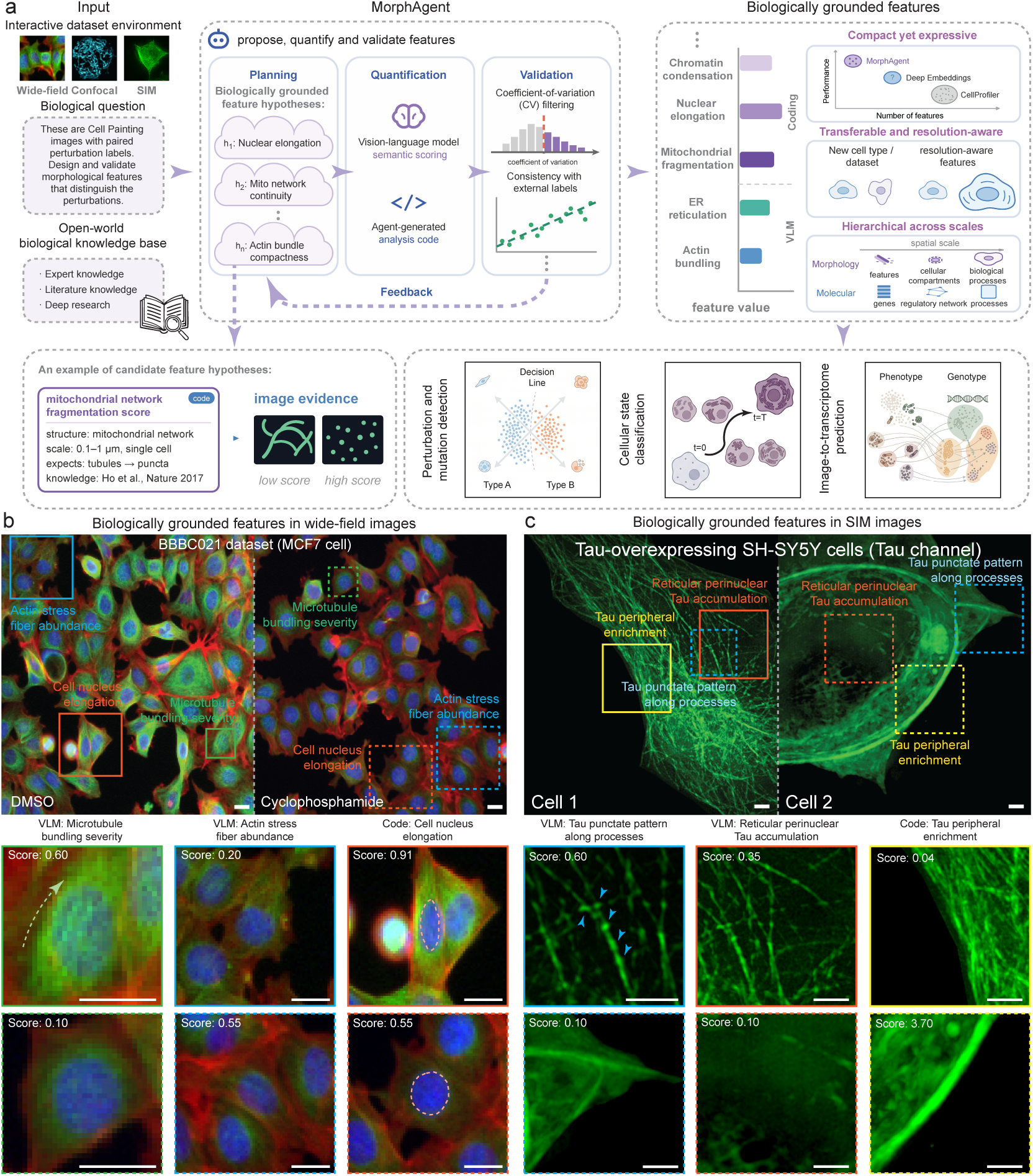
MorphAgent constructs biologically grounded image profiles across microscopy modalities. **(a)** Overview of the MorphAgent framework. MorphAgent takes an image dataset from any microscopy modality, a biological question, and an open-world biological knowledge base, and iteratively designs, quantifies and validates biologically grounded features. Each candidate feature is stated as a hypothesis (h₁–hₙ) naming a cellular or subcellular structure, its spatial scale and the expected visual evidence. Bottom left, one such hypothesis is shown, together with the retrieved literature^51^ and the image evidence expected at low and high scores. Explicitly quantifiable phenotypes are measured using agent-generated code with automated segmentation and bioimage analysis tools, whereas visually complex phenotypes are quantified through VLM semantic scoring. Candidates are screened by coefficient-of-variation filtering and consistency with external labels, and validated features guide subsequent rounds of discovery. Right, biological grounding confers three advantages, evaluated in the following sections: vocabularies are compact yet expressive, transferable and resolution-aware, and hierarchical across scales. Bottom right, the validated vocabularies support downstream analyses including perturbation and mutation detection, cellular state classification and image-to-transcriptome prediction. **(b)** Application to BBBC021 Cell Painting images of MCF7 cells. Representative MorphAgent features distinguishing cyclophosphamide-treated cells from DMSO controls include the VLM-derived features *microtubule bundling severity* and *actin stress fiber abundance*, together with the code-derived feature *cell nucleus elongation*. Zoomed-in views show representative image regions together with the corresponding feature scores. Scale bars, 5 μm. **(c)** Application to super resolution SIM images of SH-SY5Y cells across states defined by Tau expression. Representative MorphAgent features include the VLM-derived features *Tau punctate pattern along processes* and *reticular perinuclear Tau accumulation*, together with the code-derived feature *Tau peripheral enrichment*. Zoomed-in views show representative image regions together with the corresponding feature scores. Scale bars, 2 μm.

During planning, MorphAgent proposes candidate feature hypotheses, specifying the cellular structure or process they target, the spatial scale at which each feature becomes measurable, and the visual evidence expected from biological knowledge, making each feature biologically grounded. Each feature hypothesis is then converted into a score through one of two complementary routes (Methods). Explicitly quantifiable phenotypes, such as size, shape, intensity, texture, and topology of cells, nuclei and subcellular structures, are computed using automated segmentation with foundation models, open-source bioimage analysis toolkits, and agent-generated analysis code. Phenotypes that are biologically meaningful and visually recognizable but difficult to formalize analytically, such as organelle interactions, subcellular compartments, and protein-aggregate organization, are captured through VLM-based semantic scoring (Methods).

After quantification, MorphAgent validates features using statistical feedback, including coefficient of variation, redundancy and, when external labels are available, associations with external biological labels. Validated features are retained as accumulated knowledge to guide subsequent rounds of feature planning. Through this closed loop, MorphAgent screens, refines and expands its feature set (Methods), gradually converging to a compact, modality-specific feature vocabulary for downstream analysis, including perturbation and mutation detection, cellular state classification, and image-to-transcriptome prediction. Component-wise ablations showed that iterative validation, biological knowledge integration (Supplementary Fig. 2) and the two feature design pathways each contributed to the quality of the resulting feature vocabularies (Supplementary Fig. 3).

MorphAgent’s representative features in three microscopy modalities can be found in Supplementary feature lists 1–4. Figures 1b and 1c illustrate how biologically grounded features adapt to what each modality resolves. In wide-field Cell Painting, we highlighted three representative features that distinguish cyclophosphamide-treated MCF7 cells and DMSO-treated controls: two VLM-scored features, *microtubule bundling severity* and *actin stress fiber abundance*, and one code-derived feature, *cell nucleus elongation*. In each case the feature values agreed with expert assessments of the high- and low-scoring images. In super resolution SIM of Tau-overexpressing SH-SY5Y cells, MorphAgent additionally proposed features defined at the sub-diffraction scale, including *Tau punctate pattern along processes*, *reticular perinuclear Tau accumulation* and *Tau peripheral enrichment*, which resolved differences in Tau organization between individual cells. Because each feature is biologically grounded in a defined biological structure, the resulting vocabularies are compact yet expressive, transferable and resolution-aware, and hierarchical across biological scales (Fig. 1a, right). These properties are evaluated in the following sections.

### Biologically grounded features improve drug screening in wide-field Cell Painting

Microscopy-based perturbation profiling supports phenotypic drug screening by capturing cellular responses to genetic and chemical perturbations, inferring MoAs and identifying compounds with similar therapeutic effects. We evaluated biologically grounded features designed by MorphAgent on BBBC021, a canonical Cell Painting dataset for perturbation profiling in human MCF7 cells, comprising 3,552 images from 37 compounds across 26 MoA categories; the images were acquired in three fluorescence channels: DNA, tubulin and actin. Seven MoA groups contained more than one compound. MorphAgent designed 467 features, substantially more compact than CellProfiler, which contains 6,256 handcrafted features, or DeepProfiler, which contains 672 latent embeddings (Fig. 2a, Supplementary feature list 1). After removing redundant features with pairwise absolute Pearson correlation above 0.9, MorphAgent retained 291 features, compared with 440 for DeepProfiler and 1,284 for CellProfiler (Fig. 2b).

**Figure 2.**
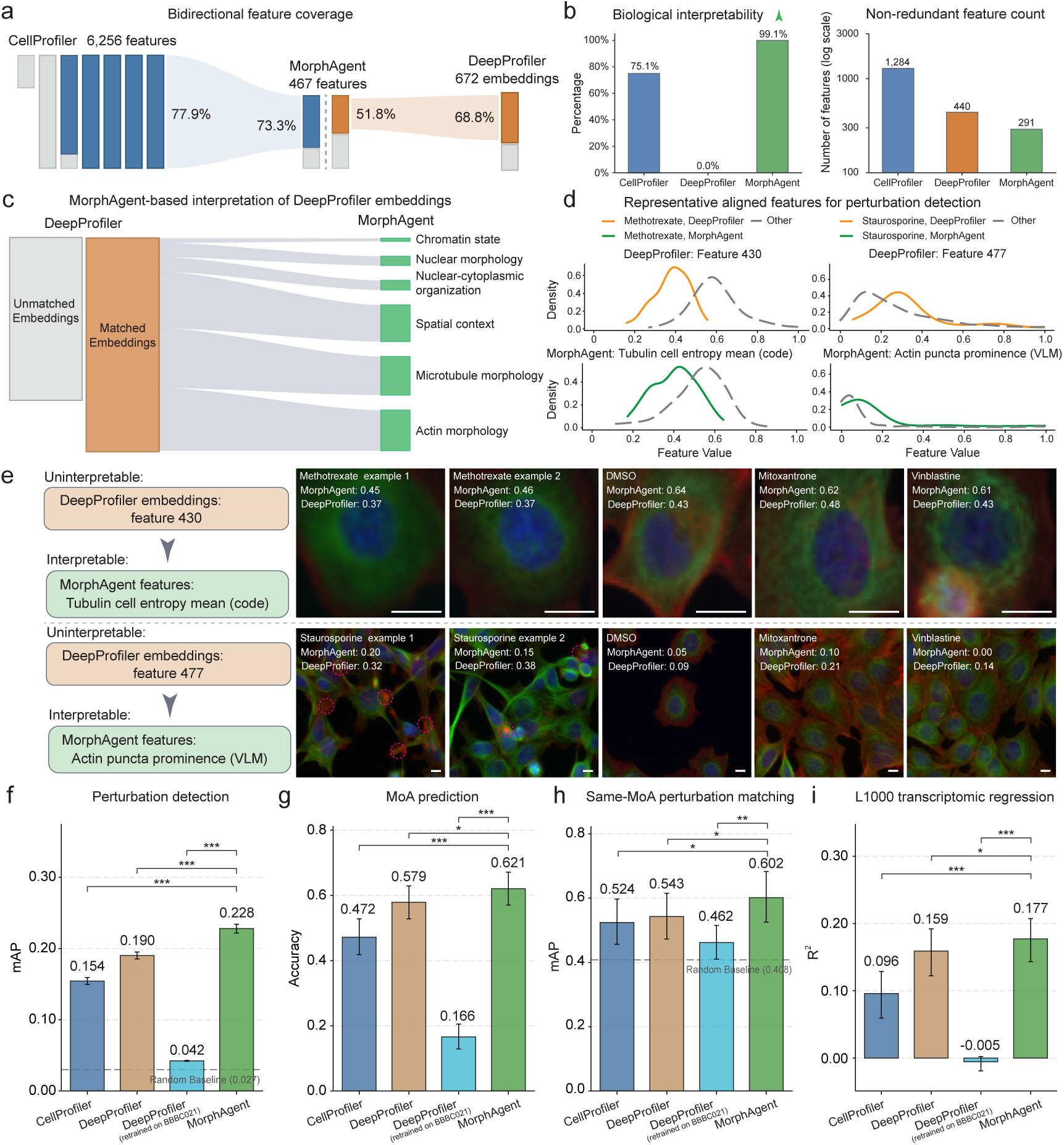
MorphAgent defines a compact, biologically grounded and predictive Cell Painting feature vocabulary. **(a)** Cross-representation correspondence among CellProfiler, MorphAgent and DeepProfiler on the BBBC021 MCF7 Cell Painting dataset. Bar lengths indicate the dimensionality of each representation (CellProfiler, 6,256 handcrafted features; MorphAgent, 467 biologically grounded features; DeepProfiler, 672 latent embeddings). A feature or embedding was considered represented by another feature space when its maximum absolute Spearman correlation with that space exceeded 0.50. MorphAgent represented 77.9% of CellProfiler features and 68.8% of DeepProfiler embeddings, whereas CellProfiler and DeepProfiler represented 73.3% and 51.8% of MorphAgent features, respectively. **(b)** Biological interpretability and redundancy of the three feature spaces. Expert review showed that 99.1% of MorphAgent features and 75.1% of CellProfiler features were biologically interpretable, whereas DeepProfiler embeddings lacked intrinsic biological semantics. After iterative redundancy pruning until no retained feature pair had |Pearson *r*| > 0.90, 291 MorphAgent, 440 DeepProfiler and 1,284 CellProfiler features remained (logarithmic scale). **(c)** Biological interpretation of DeepProfiler embeddings using matched MorphAgent features. Matched MorphAgent features provide interpretable annotations spanning chromatin state, nuclear morphology, nuclear–cytoplasmic organization, spatial context, microtubule morphology and actin morphology. **(d)** Representative matched feature pairs. Density distributions compare DeepProfiler feature 430 with the code-derived MorphAgent feature *tubulin cell entropy mean* for methotrexate-treated cells, and DeepProfiler feature 477 with the VLM-derived feature *actin puncta prominence* for staurosporine-treated cells. Features were independently min–max scaled to the range 0–1 before visualization. Curves compare the highlighted perturbation with all other conditions. **(e)** Representative single-cell images and corresponding feature values for the matched examples shown in (d) across the highlighted perturbation, DMSO controls and comparison compounds. Scale bars, 5 μm. **(f–i)** Benchmark comparison of CellProfiler, two DeepProfiler variants (pretrained on large-scale Cell Painting data and retrained solely on BBBC021) and MorphAgent. **(f)** Perturbation detection evaluated by image-level same-compound retrieval using mAP. **(g)** MoA prediction evaluated by image-level *k*-nearest-neighbour classification. **(h)** Same-MoA perturbation matching evaluated across the seven MoA groups containing at least two compounds. **(i)** L1000 gene-expression regression evaluated by predicting the 977 landmark genes from morphology profiles using a fixed training–validation–test split and a global *R*^2^ score. Across these benchmarks, MorphAgent showed the strongest overall performance, with the clearest gains on perturbation detection, same-MoA matching, and L1000 regression, while remaining comparable to DeepProfiler on MoA prediction. Error bars in f–i show 95% bootstrap percentile confidence intervals from 2,000 bootstrap resamples.

This compactness did not come at the expense of the information content of the features. At a Spearman correlation threshold of 0.5, MorphAgent covered 77.9% of CellProfiler features, with 73.3% of MorphAgent features reciprocally covered by CellProfiler. Of the DeepProfiler embeddings, 68.8% were covered by MorphAgent, whereas 51.8% of MorphAgent features were covered by DeepProfiler (Fig. 2a). These overlaps indicate that the biologically grounded vocabulary spans the variation captured by both existing paradigms with far fewer and less redundant dimensions. The vocabulary was biologically interpretable by construction. Expert review judged 463 of 467 MorphAgent features (99.1%) to be biologically interpretable. In comparison, 24.9% of CellProfiler features were not interpretable and all DeepProfiler latent embeddings lacked intrinsic biological semantics (Fig. 2b). Expert-assisted validation further confirmed that most features measured their associated biological concepts and that VLM-derived scores were grounded in the appropriate biological structures (Supplementary Fig. 4).

Leveraging cross-representation correlations, we annotated DeepProfiler embeddings using their most similar biologically grounded features (Methods). The corresponding biologically grounded features provided biological interpretations spanning chromatin state, nuclear morphology, nuclear–cytoplasmic relations, spatial context and cytoskeletal organization (Fig. 2c). Two representative examples linked DeepProfiler feature 430 to the code-derived feature *tubulin cell entropy mean* and DeepProfiler feature 477 to the VLM-derived feature *actin puncta prominence* (Fig. 2d). In both cases, the DeepProfiler embeddings and matched MorphAgent features showed similar distribution shifts between the selected perturbation and other conditions, which were also reflected in representative cell images (Fig. 2d, e).

We next benchmarked biologically grounded features on various Cell Painting tasks against CellProfiler and two DeepProfiler variants: the original model pretrained on large-scale Cell Painting data and a model retrained solely on the 3,552 BBBC021 images (Methods). We evaluated four tasks: perturbation detection, measured by image-level same-compound retrieval using mean average precision (mAP) (Fig. 2f); MoA prediction, measured by k-nearest-neighbour classification (Fig. 2g); same-MoA perturbation matching, measured by compound-level retrieval among perturbations sharing a MoA (Fig. 2h); and transcriptomic regression, measured by prediction of paired L1000^18^ signatures using a shared multilayer perceptron (Fig. 2i). Across the four tasks, MorphAgent achieved the best overall performance: perturbation detection mAP = 0.228, MoA prediction accuracy = 0.621, same-MoA matching mAP = 0.602 and L1000 regression R² = 0.177. These results exceeded large-scale pretrained DeepProfiler (0.190, 0.579, 0.543, and 0.159) and CellProfiler (0.154, 0.472, 0.524 and 0.096) in the same task order. Notably, DeepProfiler retrained solely on BBBC021 performed poorly across all four tasks (0.042, 0.166, 0.462 and −0.005). The advantage of biological grounding remained robust when feature design used as few as one reference image per compound. MorphAgent outperformed additional label-free representation learning methods^12,15^ and generalized to the other Cell Painting benchmarks, such as BBBC022 (Supplementary Figs. 5–7).

This performance gain arises from biological grounding. It is particularly evident for DNA-targeting compounds, where MorphAgent’s stronger performance is driven largely by biologically grounded nuclear features that directly capture cellular responses associated with DNA damage (Supplementary Fig. 8 and Supplementary feature list 5). For example, one such feature, *nuclear boundary irregularity*, quantifies the loss of nuclear shape integrity associated with nuclear blebbing, a phenotype previously reported after treatment with DNA-targeting compounds such as cisplatin^19^. MorphAgent converts this biological phenotype into an explicit quantitative feature through image segmentation and computational measurement and retains it through statistical validation. By contrast, CellProfiler represents nuclear morphology largely through generic geometric and texture descriptors, such as Zernike features (Supplementary feature list 6), which capture shape variation but are not explicitly defined around the underlying biological process. Biological grounding therefore produces more expressive features that directly capture biologically relevant phenotypes.

### Biologically grounded features transfer across confocal imaging resolutions for cell state classification

Spinning-disk confocal microscopy resolves mitochondrial architecture in substantially finer detail than wide-field imaging. We applied MorphAgent to a published omics-paired HSC dataset comprising 110 single cells from mice (48 young and 62 aged), with paired mitochondrial images (approximately 200 nm resolution) and single-cell RNA-seq labels^20^. Guided by a 107-article knowledge library, a description of acquisition protocols, and a deep research report on mitochondrial biology, MorphAgent generated and iteratively refined candidate features, ultimately selecting a compact feature vocabulary of 25 transcriptome-anchored mitochondrial morphology features for characterizing age-associated HSC states (Methods; Supplementary feature list 2). This feature vocabulary was predefined before transfer to an independently acquired HSC validation dataset with higher image quality (approximately 120 nm resolution), allowing us to test whether the discovered mitochondrial features generalized across datasets and imaging experiments.

In the feature discovery dataset comprising 110 cells, the 25 biologically grounded features distinguished young from aged HSCs more effectively than manually curated mitochondrial features. Using images alone, MorphAgent features achieved an AUC of 0.739, compared with 0.575 for expert-designed features^20^, including the short-to-long and dispersed-to-polarized mitochondrial patterns previously used for HSC state analysis (Fig. 3a). For reference, the HVG500 transcriptomic baseline achieved an AUC of 0.920. The transcriptome-anchored feature vocabulary aligned with transcriptional programs related to aging, proliferation and stress, including genes such as Stmn1, Cdk1 and Gdf15 (Fig. 3b), consistent with their well-established roles in a previous publication^20^. A STRING^21^-based protein–protein interaction analysis further showed that genes most strongly associated with MorphAgent features were more likely to lie near the highest-degree node of the network (Trp53) than weakly associated genes (Fig. 3c, Supplementary Fig. 9 and Methods).

**Figure 3.**
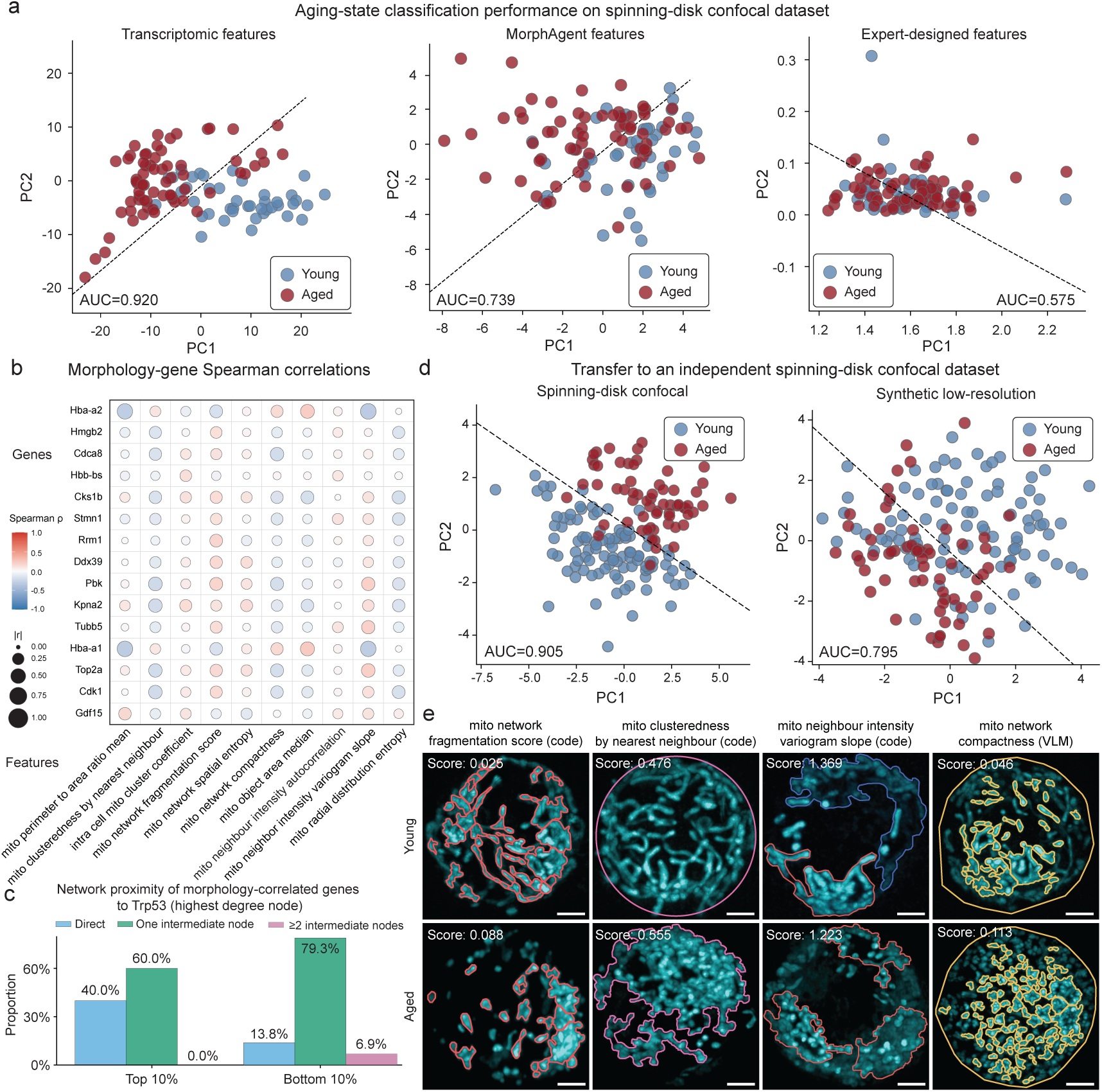
MorphAgent discovers transferable mitochondrial features across independent mouse HSC datasets. **(a)** Age-state discrimination in the discovery dataset of 110 single HSCs. PCA projections are shown for transcriptomic profiles, the 25 MorphAgent mitochondrial features and expert-designed mitochondrial features. Classification AUCs were 0.920, 0.739 and 0.575, respectively. Dashed lines indicate decision boundaries. **(b)** Spearman correlations between representative MorphAgent mitochondrial features and age-associated genes. Circle color indicates correlation direction and magnitude, and circle size indicates absolute correlation magnitude. **(c)** Distribution of genes in the top and bottom 10% of morphology–gene association strength according to their shortest-path distance from Trp53 in a STRING protein–protein interaction network. Trp53 was selected post hoc as the highest-degree node in the analyzed network. Compared with the bottom decile, the top decile contained a larger proportion of genes directly connected to Trp53. **(d)** Transfer of the fixed 25-feature vocabulary, without feature reselection, to an independent validation dataset of 162 HSCs. PCA projections and decision boundaries are shown for the original spinning-disk confocal images and matched synthetic low-resolution representations generated by two-dimensional Gaussian blurring. Classification AUCs were 0.905 and 0.795, respectively. **(e)** Representative young and aged HSCs showing four transferred features: *mitochondrial network fragmentation score*, *mitochondrial clusteredness by nearest neighbour*, *mitochondrial neighbour intensity variogram slope* and *mitochondrial network compactness*. Overlays indicate the structures or regions used for quantification, with feature scores shown. Scale bars, 2 μm.

We next applied the 25 biologically grounded features, without feature reselection, to an independent in-house HSC dataset of 162 cells (99 young and 63 aged) acquired by spinning-disk confocal microscopy (Methods). The validation images showed higher image quality and finer visible mitochondrial detail than the discovery images (Supplementary Fig. 10). On the validation dataset, the feature vocabulary achieved an AUC of 0.905 for young-versus-aged classification. Conversely, when we applied two-dimensional Gaussian blurring to the validation images to mimic the image quality of the discovery dataset, the classification AUC decreased to 0.795, similar to the results of the discovery dataset (Fig. 3d and Methods). Representative features, including *mitochondrial network fragmentation*, *clusteredness*, *neighbour-intensity variogram slope* and *network compactness*, consistently captured the transition from integrated mitochondrial networks in young HSCs to more fragmented and dispersed mitochondrial organization in aged HSCs (Fig. 3e). These findings demonstrate that biologically grounded features can transfer across imaging experiments, connecting mitochondrial architecture with molecular cell state variation.

### Biologically grounded super resolution features enable accurate disease-associated mutation classification and bidirectional morphology–transcriptome prediction

We next applied MorphAgent to super resolution SIM of Tau-labeled samples, which resolves subcellular architecture beyond the diffraction limit at approximately 100 nm resolution. This resolution reveals distinct Tau organizations (puncta, fibrils, bundles) that are clearly distinguishable. While traditional methods struggle to capture these patterns due to reliance on limited expert knowledge for feature engineering, MorphAgent enables comprehensive characterization of disease-associated mutations. In a discovery dataset of 58 Tau-overexpressing SH-SY5Y cells with paired SIM images and single-cell transcriptomes, we focused on Tau protein organization, a structural biomarker of tauopathy whose hallmark phenotypes become accessible only with super resolution imaging. Guided by 31 articles on Tau imaging, expert knowledge on Tau-associated pathogenic phenotypes, and an automated deep research report, MorphAgent generated and iteratively refined candidate features using cross-cell variability, yielding a 301-feature biologically grounded Tau morphology vocabulary (Supplementary feature list 3).

We compared this biologically grounded feature vocabulary with a conventional expert-designed feature vocabulary comprising 16 features based on biological processes central to Tau biology (Supplementary Table 2). MorphAgent recovered all 16 expert-designed features, with each feature matched by at least one MorphAgent feature at high Spearman correlation (ρ > 0.7), while preserving a similar global organization of cells in PCA space (Fig. 4a, b). Notably, 48.6% of MorphAgent features were largely orthogonal to the expert-designed feature vocabulary (maximum absolute Spearman ρ < 0.5), indicating substantial additional information beyond the expert-designed features. Expert review judged 295 of 301 features (98.0%) to be biologically meaningful, and semantic analysis grouped these features into coherent structural themes, including Tau-positive puncta, fibril orientation and bundle formation (Fig. 4c, Methods).

**Figure 4.**
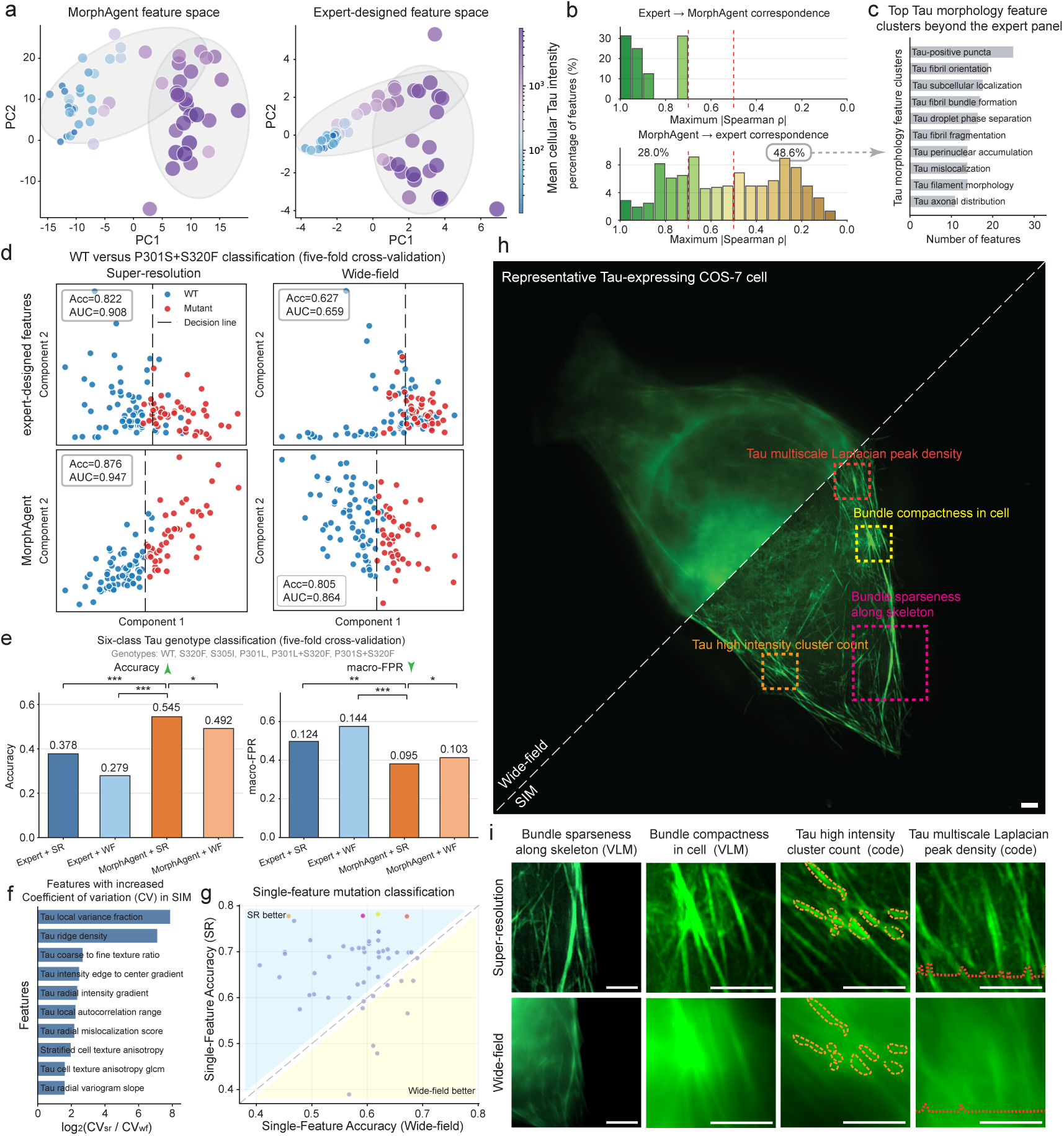
MorphAgent discovers informative Tau features revealed by super resolution microscopy. **(a)** Principal-component projections of the 301 MorphAgent features and the 16 expert-designed Tau features in the wild-type (WT) Tau discovery dataset. Cells are colored by mean cellular Tau intensity. MorphAgent recovers the broad organizational structure of the expert-designed features while revealing greater phenotypic diversity. **(b)** Cross-space feature correspondence in the WT dataset. The upper distribution shows, for each expert feature, its maximum absolute Spearman correlation with any MorphAgent feature; all 16 expert features had a MorphAgent match with |ρ| > 0.70. The lower distribution shows the reverse mapping; 48.6% of MorphAgent features had maximum |ρ| < 0.50 with the expert feature vocabulary. **(c)** LLM-based biological-process clustering of MorphAgent features weakly represented by the expert feature vocabulary, revealing recurrent Tau structural phenotypes beyond expert-designed descriptors. **(d)** Binary classification of WT versus the P301S+S320F double mutant (five-fold cross-validation, L2-regularized logistic regression), shown as decision-axis projections for both modalities and feature sets. MorphAgent provided the strongest separation between WT and mutant cells, with the largest improvement observed for super resolution imaging (super resolution: MorphAgent (Acc = 0.876, AUC = 0.947) versus expert-designed features (Acc = 0.822, AUC = 0.908); wide-field: MorphAgent (Acc = 0.805, AUC = 0.864) versus expert-designed features (Acc = 0.627, AUC = 0.659)). **(e)** Six-class genotype classification across WT and the five Tau mutants (five-fold cross-validation). MorphAgent on super resolution images achieved the best genotype discrimination, demonstrating that resolution-dependent phenotypes improve multiclass mutation profiling. **(f)** Features exhibiting the largest increase in cross-cell variability under super resolution imaging, indicating phenotypic heterogeneity revealed only at higher spatial resolution. **(g)** Single-feature mutation-classification accuracy on super resolution versus wide-field images for the top 50 mutation-discriminative features. The dashed diagonal indicates equal accuracy; more than 80% of the mutation-discriminative features became more informative under super resolution imaging. Colored points indicate the representative features shown in (h). **(h)** Matched wide-field and super resolution views of the same representative cell, split along the diagonal. Boxes mark regions corresponding to four informative MorphAgent features: *bundle sparseness along the skeleton*, *bundle compactness in cell*, *Tau high-intensity cluster count* and *Tau multiscale Laplacian peak density*. Scale bar, 2 μm. **(i)** Local zoom-ins for the features in (h), showing how super resolution resolves bundle architecture, punctate structure and textural detail that are blurred in wide-field. Scale bar, 2 μm.

We then examined whether the expanded Tau morphology feature vocabulary could distinguish disease-associated mutations and capture resolution dependent phenotypes. We transferred the vocabulary unchanged to a different cell type and species: 495 COS-7 cells expressing wild-type Tau or one of five Tau mutants, imaged with both SIM and wide-field microscopy. For wild-type versus P301S+S320F classification, MorphAgent features achieved an accuracy of 0.876 and an AUC of 0.947 on SIM images, outperforming the expert-designed features (accuracy = 0.822, AUC = 0.908; Fig. 4d). When applied to paired wide-field images, MorphAgent’s performance decreased to an accuracy of 0.805 and an AUC of 0.864, but remained better than the expert-designed features on the same images (accuracy = 0.627, AUC = 0.659; Fig. 4d). Biologically grounded features also achieved the best overall performance in the more challenging six-class genotype classification task, with an accuracy of 0.545 and a macro-averaged false-positive rate (macro-FPR) of 0.095 (Fig. 4e; Supplementary Fig. 11).

The performance gain on SIM images reflected the contribution of features sensitive to fine structural information resolved only at higher optical resolution. Morphology features showed greater cross-cell variation in SIM than in paired wide-field images, expanding the measurable dynamic range of specific Tau phenotypes (Fig. 4f). Accordingly, when individual features were used for binary classification, more than 80% of the top 50 mutation-discriminative features classified mutation states more accurately from SIM images than from wide-field images (Fig. 4g, Supplementary Fig. 12). Representative features captured both higher-order bundle architecture and fine-scale Tau organization, including *bundle sparseness along the skeleton*, *bundle compactness*, *high-intensity cluster count* and *multiscale Laplacian peak density* (Fig. 4h, i). Progressive image blurring further reduced classification performance as fine structural information was lost, confirming that MorphAgent’s advantage on SIM images derives from resolution-dependent morphology features (Supplementary Fig. 13).

We further examined whether the super resolution Tau morphology captured molecular variation beyond its ability to distinguish genotypes. Using the transcriptomics-paired SIM discovery dataset, we redesigned the feature vocabulary using transcriptomic profiles as validation signals, yielding 400 morphological features (Supplementary feature list 4). These features covered 79% of the original feature set at |ρ| > 0.7 and 96% at |ρ| > 0.5, while identifying additional features that were predictive of transcriptomic variation. These results indicate that transcriptomic profiles provide richer feedback for iterative feature design (Supplementary Fig. 14).

We trained support-vector regression (SVR) models to predict each gene’s expression from biologically grounded features, measured by the mean Pearson correlation between predicted and measured expression in the test dataset (Methods). Across 27,476 genes, MorphAgent achieved a mean prediction correlation of 0.100 ± 0.004 (3σ). By comparison, expert-designed Tau features and Tau intensity alone showed little predictive capacity (0.005 ± 0.003 and −0.002 ± 0.003; Fig. 5a). Predictability was concentrated in a subset of genes: when genes were ranked by each method’s own prediction performance, mean correlation for the top 5,000 genes reached 0.340 for MorphAgent, 0.175 for the expert-designed feature vocabulary, and 0.156 for Tau intensity, respectively. For the top 1,000 genes, the corresponding values were 0.553, 0.317 and 0.296. We quantified shared information between Tau morphology and transcriptomics through bidirectional prediction (Fig. 5b). MorphAgent features predicted a subset of genes (mean r = 0.198; 3.0% with r > 0.50), whereas transcriptomic profiles predicted MorphAgent features more strongly (mean r = 0.423; 25.5% with r > 0.50), indicating overlapping but complementary information between the two modalities. This asymmetry is expected because the available microscopy images capture Tau architecture alone, while the transcriptome reflects a broader range of cellular processes.

**Figure 5.**
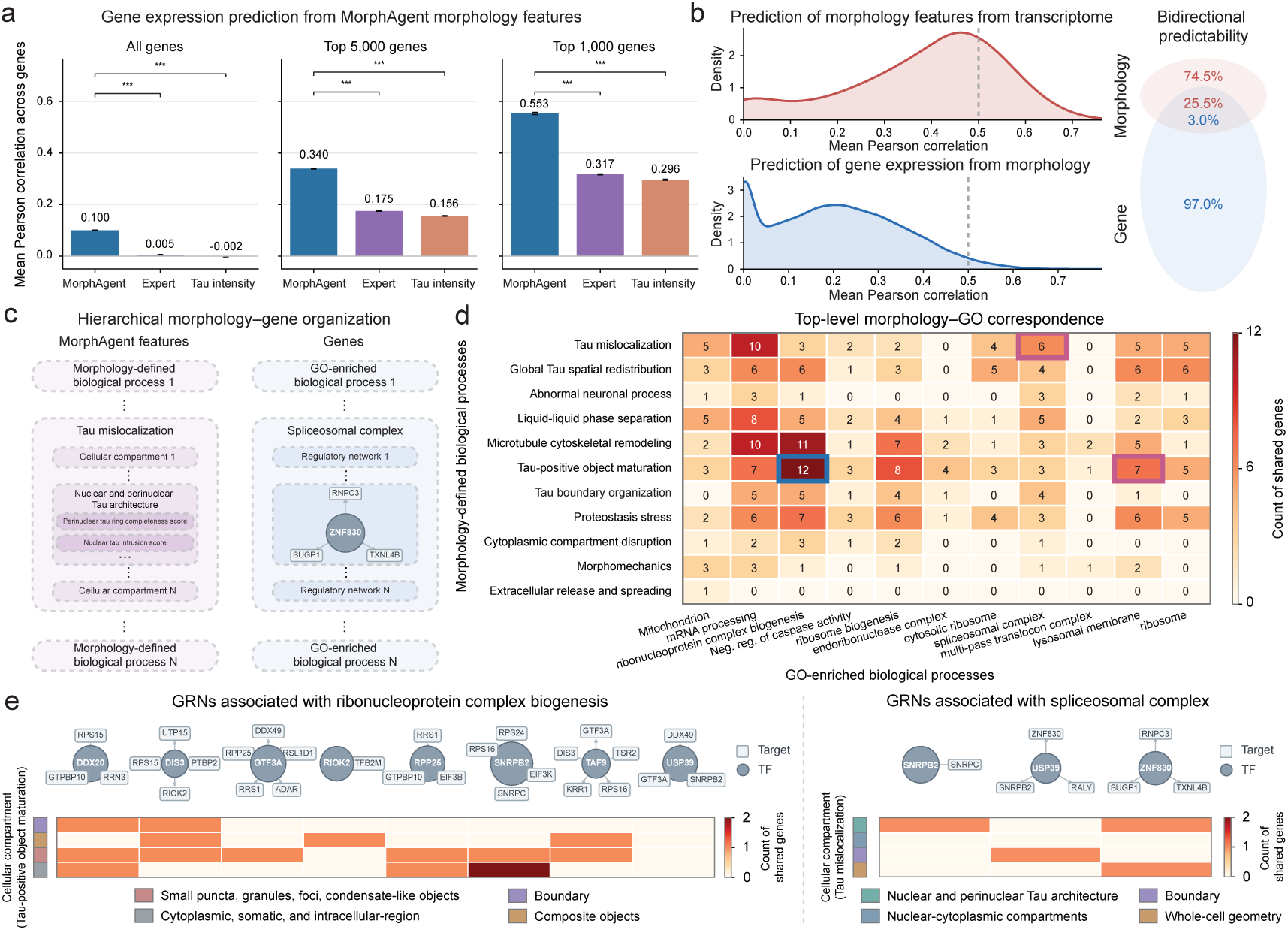
MorphAgent links super resolution Tau morphology to single-cell transcriptomic programs through a hierarchical biological representation. **(a)** Prediction of single-cell gene expression from Tau morphology. Bars show the mean held-out Pearson correlation between predicted and measured expression using the expanded 400-feature MorphAgent vocabulary, the expert-designed Tau features or whole-cell Tau intensity. Results are shown for all 27,476 measured genes and for the top 5,000 and top 1,000 genes ranked separately by each method’s prediction performance. Error bars show the standard error of the mean across genes. **(b)** Bidirectional prediction between super resolution Tau morphology features and transcriptomic profiles. The upper density plot shows prediction performance for MorphAgent features from transcriptomic profiles, and the lower plot shows prediction performance for genes from MorphAgent morphology. Dashed lines mark Pearson r = 0.50. The schematic summarizes the proportions of MorphAgent features and genes with prediction performance exceeding this threshold: 25.5% of MorphAgent features and 3.0% of genes. **(c)** Three-level morphology–gene hierarchy. MorphAgent features are organized from local features to cellular compartments and morphology-defined biological processes, whereas genes are organized from individual genes to GRN modules and Gene Ontology (GO)-enriched biological processes. **(d)** Top-level morphology–GO correspondence map based on the top 1,000 genes most predictable from MorphAgent morphology. Rows show 11 Tau-pathology morphology classes, and columns show 11 GO terms. Each cell reports the number of shared genes. The strongest correspondence was between Tau-positive object maturation and ribonucleoprotein complex biogenesis, with 12 shared genes. Two additional literature-supported correspondences linked Tau mislocalization to the spliceosomal complex (six genes) and Tau-positive object maturation to the lysosomal membrane (seven genes). **(e)** Intermediate-level correspondence between cellular compartments and GRN modules for ribonucleoprotein complex biogenesis (left) and the spliceosomal complex (right). Circles denote transcription factors, boxes denote target genes, and heatmap cells report shared-gene counts. Ribonucleoprotein complex biogenesis GRNs were linked mainly to punctate, granule-like and condensate-like structures and broader cytoplasmic architectures. Spliceosomal complex GRNs were linked primarily to nuclear and perinuclear Tau architecture.

### Hierarchical organization of biologically grounded features supports mechanistic understanding

Because every biologically grounded feature identifies a specific structure at a stated spatial scale, the features can be grouped into cellular compartments and then into morphology-defined biological processes (Fig. 5c, left). Genes were grouped in parallel into regulatory networks and then into GO-enriched biological processes (Fig. 5c, right; Methods). The two sides were linked through the bidirectional prediction: among the 1,000 most morphology-predictable genes, each feature was assigned the genes it predicted, and every cellular compartment and biological process inherited the genes of the features it contained. At the top level, we therefore quantified gene overlap between each morphology-defined biological process and each GO-enriched process (Fig. 5d and Methods). In the morphology–GO correspondence map, the strongest association was between *Tau-positive object maturation* and *ribonucleoprotein complex biogenesis*, with 12 shared genes (Fig. 5d). Two additional associations were supported by prior studies. *Tau mislocalization* shared six genes with the *spliceosomal complex*. Pathological Tau can interact with spliceosomal proteins and mislocalize nuclear-speckle components, leading to pre-mRNA splicing defects^22,23^. *Tau-positive object maturation* shared seven genes with the *lysosomal membrane*. This association is biologically plausible because pathological Tau accumulates in and damages lysosomes, while lysosomal trafficking and exocytosis modulate its clearance and propagation^24–26^. A permutation test was used to support the significance of the morphology–GO associations (Supplementary Fig. 15; Methods).

At the intermediate level, we linked cellular compartments to gene regulatory networks (GRNs) using the same gene overlap calculation (Methods). GRNs associated with *ribonucleoprotein complex biogenesis* were linked mainly to *small puncta, granule-like and condensate-like structures*, as well as *cytoplasmic and broader intracellular-region architectures* (Fig. 5e). GRNs associated with the *spliceosomal complex* were linked primarily to *nuclear and perinuclear Tau architecture* (Fig. 5e, Supplementary Fig. 16). The intermediate-level analysis of the *lysosomal membrane* association is shown separately in Supplementary Fig. 17. These results show that the morphology–gene hierarchy connects nanoscale Tau architecture with molecular functions and regulatory programs, positioning super resolution morphology as a biologically interpretable complement to transcriptomic profiling for mechanistic understanding.

## Discussion

Microscopy-based cell profiling provides a quantitative bridge between image data and cellular phenotypes for biological discovery, disease characterization and phenotypic drug screening. Existing cell profiling approaches, however, rely largely on handcrafted geometric features or black-box deep learning embeddings whose biological meaning is unclear, limiting their predictive power, interpretability, transferability and ability to support biological understanding or translational research. Here we introduce biologically grounded cell profiling, in which each feature originates from a biological hypothesis and is anchored to a cellular structure or process. Because feature identity is defined by biology rather than by a particular staining protocol, sample preparation or imaging modality, biologically grounded features are compact yet expressive, transferable across imaging conditions, and naturally organized hierarchically across spatial scales. These properties enable broad applications across distinct microscopy modalities and biomedical settings: biologically grounded features enable accurate profiling of chemical perturbations in Cell Painting for phenotypic drug screening and MoA analysis, quantify aging-associated states in hematopoietic stem cells across independently acquired datasets and resolutions, and resolve disease-associated Tau mutation phenotypes that become accessible only with super resolution imaging. Beyond prediction, their explicit biological organization enables bidirectional prediction between multiscale morphology and single-cell transcriptomic programs and reveals hierarchical associations linking morphology to genes, regulatory networks and biological processes, extending cell profiling toward broader mechanistic understanding and translational discovery.

MorphAgent provides an agentic AI framework for making biologically grounded feature design systematic and scalable. Recent scientific agents such as Biomni^27^ and Robin^28^ have demonstrated the potential of agentic systems for reasoning over structured molecular, tabular and sequence data^29–42^. Bioimaging poses a different challenge: biologically meaningful information is encoded implicitly in high-dimensional visual patterns and must first be recognized and converted into quantitative features before downstream reasoning is possible. MorphAgent automates this process through an iterative loop of literature-grounded hypothesis generation, multimodal image reasoning, quantitative feature implementation and validation, using feature variation and external omics data, to refine the resulting vocabulary. The central advance is therefore not automation alone, but the scalable construction and testing of image features that retain explicit biological meaning. This strategy could extend beyond fluorescence microscopy to other image-based biomedical analyses, including histopathology, multiplexed tissue imaging and clinical imaging.

Nonetheless, several limitations remain. First, although MorphAgent expands the measurable morphological space, its feature vocabulary is not yet exhaustive. The observation that combining MorphAgent features with redundancy-filtered CellProfiler features further improved profiling performance (Supplementary Fig. 18) indicates that conventional low-level features capture complementary information not represented by the current biologically grounded feature vocabulary. Second, the quality of agent-designed features depends on the biological knowledge and validation signals available to the system. In MorphAgent, biological knowledge guided feature proposal, whereas iterative validation provided feedback for feature refinement (Supplementary Fig. 2). In biological systems with limited prior knowledge or unavailable validation signals, feature discovery may therefore remain challenging. Third, VLM-based semantic scoring expands the accessible feature space, but its ability to encode complex biological phenotypes remains limited. Expert evaluation showed that semantic plausibility does not always correspond to accurate phenotype ranking (Supplementary Fig. 19), and performance varies across model backbones (Supplementary Fig. 20). Moreover, large-scale VLM scoring remains computationally expensive (Supplementary Fig. 21).

These limitations suggest several directions for future development. First, future agentic profiling systems could incorporate existing profiling approaches, such as CellProfiler and DeepProfiler, as computational tools within the agent framework. Beyond improving predictive performance through complementary feature vocabularies, such systems could use agentic reasoning to uncover the biological meanings underlying previously unexplained morphological descriptors. Second, building modality-adaptive biological knowledge bases and large-scale validated feature libraries could reduce dependence on experts’ manual input and enable more robust feature discovery across imaging platforms. Third, fine-tuning vision-language models on large-scale microscopy datasets may enable more faithful biological reasoning and more efficient semantic feature discovery. Finally, extending agentic feature design to live-cell video profiling may enable discovery of dynamic phenotypes that complement static molecular measurements. Because many biological processes are encoded in temporal trajectories rather than captured by static snapshots, agent-designed temporal features could provide new representations for modeling cellular dynamics and predicting future cell states.

Looking forward, MorphAgent represents an initial step toward agentic microscopy, accelerating imageomics-driven biological discovery. More broadly, biological grounding may offer a general principle for trustworthy biomedical AI, linking increasingly powerful imaging and AI more directly to biological mechanisms, cellular state modeling, and biomedical translation.

## Methods

### Data collection and preprocessing

MorphAgent was evaluated on six imaging datasets spanning wide-field microscopy, spinning-disk confocal microscopy and SIM. Supplementary Table 1 summarizes the imaging modalities and acquisition details, sample sizes and biological labels, and paired transcriptomic profiles or other paired biological metadata for all six datasets, where available.

### BBBC021 Cell Painting dataset using wide-field microscopy

The wide-field Cell Painting dataset used in our analysis was constructed as a subset of the public BBBC021 dataset of MCF7 cells by retaining compounds with curated MoA annotations. This yielded 37 annotated compounds from the original 113 compounds. Data were downloaded from the BBBC021 public repository^43,44^ and included 49 imaging plates from Weeks 1–10, comprising 3,552 images across 26 MoA categories. Seven of these categories contained multiple compounds, enabling evaluation of same-MoA matching. Each image contained three fluorescence channels corresponding to DNA, β-tubulin and F-actin, acquired at 20× magnification on the original BBBC021 imaging platform. Per-image information included plate identifier, well position, imaging site, compound identity, compound concentration, and MoA annotation. We matched BBBC021 compound conditions to L1000 transcriptional profiles^18^, an open collection of compound-induced gene-expression signatures, using compound identity and concentration when matched profiles were available. These matched conditions were used to construct paired image–transcriptomic profiles for comparing MorphAgent-derived features with molecular perturbation responses.

### BBBC022 Cell Painting dataset using wide-field microscopy

We used a single-plate subset of the public BBBC022 release (plate 20585), comprising 1,026 wide-field Cell Painting fields from U2OS cells: 450 compound-treated images (50 bioactive compounds, nine sites per well) and 576 mock controls (64 wells, nine sites each). Images contained five fluorescence channels (ER, DNA, mitochondria, AGP/actin, RNA). Metadata included plate, well, site, compound name, BROAD ID and condition (compound or mock), and were used for perturbation-detection benchmarks.

### Mitochondrial imaging datasets using spinning-disk confocal microscopy

We analyzed two mouse HSC mitochondrial imaging datasets acquired by spinning-disk confocal microscopy. The first dataset was a publicly available image–transcriptomics-paired dataset consisting of 110 single HSCs with paired single-cell Smart-seq2 transcriptomic profiles^20,45^. This dataset was used to construct MorphAgent mitochondrial morphology features and to associate these features with transcriptomic cell states.

The second dataset was an independently acquired validation dataset comprising 162 single HSCs. Before imaging, mitochondria were labeled in live cells with MitoTracker Green (0.67 μM; Cell Signaling Technology, 9074S) under 488-nm excitation in the presence of verapamil (83 μM) to reduce dye efflux. Imaging was performed using a CSU-X1 Yokogawa spinning-disk confocal head mounted on an inverted Olympus IX81 microscope equipped with a 100× oil-immersion objective lens. Images were sampled at 38.2 nm laterally and 200 nm axially, and the estimated lateral optical resolution was approximately 120 nm. 3D image stacks were converted into 2D maximum-intensity projections for downstream morphological profiling. The predefined MorphAgent feature vocabulary was evaluated on the original spinning-disk confocal images and on paired synthetic low-resolution representations generated from the same cells by two-dimensional Gaussian blurring, as described below.

In both datasets, long-term HSCs were classified according to the chronological age of donor mice (young: 6–8 weeks old; aged: 12–22 months old). Mitochondrial morphology reflects a continuous and heterogeneous cellular state rather than an absolute age-specific feature.

### Tau imaging datasets using SIM

We analyzed two Tau-overexpression datasets acquired on the same SIM platform (HIS-SIM, Guangzhou Computational Super resolution Biotech) with Tau-Enhanced Green Fluorescent Protein (EGFP) imaged under 488-nm excitation. SIM image stacks were acquired with a lateral pixel size of 32.5 nm and an axial step size of 300 nm, yielding a nominal reconstructed lateral resolution of approximately 100 nm.

The first dataset was a paired imaging and transcriptomic dataset used for MorphAgent feature design, consisting of 58 SH-SY5Y cells overexpressing wild-type human 0N4R Tau-EGFP. Cells were imaged as three-dimensional stacks, with paired single-cell Smart-seq2 transcriptomic profiles covering 27,476 genes. Morphological profiling was performed on 2D maximum-intensity projections of the corresponding 3D stacks. The second dataset was a mutant validation dataset consisting of 495 COS-7 cells spanning six genotypes: wild type (WT), S320F, S305I, P301L, P301L+S320F and P301S+S320F. For each cell, raw SIM data were used to generate two matched image types: super resolution SIM images reconstructed using Wiener filtering and wide-field images generated by averaging the corresponding raw SIM frames. This dataset was used to evaluate whether MorphAgent-derived Tau morphology features generalized across Tau variants and imaging resolutions.

### MorphAgent framework

MorphAgent is an agentic framework for biologically grounded feature design in cell profiling. It iteratively designs, quantifies and validates morphology features to build compact, modality-adaptive feature vocabularies (Algorithm 1). Explicitly quantifiable phenotypes are implemented through agent-generated code, whereas visually recognizable but hard-to-formalize phenotypes are quantified by VLM-based semantic scoring. Perturbation labels, phenotypic annotations and omics readouts, when available, provide feedback signals for iterative feature validation.

### Dataset preparation

MorphAgent first constructs an interactive environment for each microscopy dataset. The required inputs include the raw microscopy dataset and a user query prompt defining the biological question. Optional inputs include a dataset description file and paired metadata. The dataset description file specifies image shapes, channel definitions, sample identifiers and file organization, enabling the coding agent and VLM modules to correctly load, inspect and interpret the dataset. Paired metadata provide paired biological information for each sample, including perturbation labels, genotype annotations, or transcriptomic programs, which are used as validation signals during the validation stage.

Before entering the agentic feature-design loop, MorphAgent performs dataset-specific image preprocessing and segmentation. Based on the dataset description and channel information, modality-appropriate segmentation models are selected to generate masks for relevant cellular or subcellular structures. In the wide-field Cell Painting dataset, whole-cell and nuclear masks were generated using Cellpose-SAM^46^, and cytoplasmic masks were obtained by subtracting nuclear masks from whole-cell regions. For spinning-disk confocal and SIM datasets, detailed mitochondrial structures and Tau architectures were segmented using the AICS segmentation framework^47^. For VLM-based semantic scoring, images were resized to 512×512, normalized to 0–255 using clipping at the 1st and 99th percentiles, and converted into 2D PNG inputs, with 3D or multi-channel images exported as channel-specific slices or projection images when appropriate.

### Biological knowledge integration

MorphAgent integrates task-specific biological knowledge bases to guide biologically grounded feature discovery. Each knowledge base consists of three complementary sources: expert knowledge, research articles and LLM-assisted deep research summaries. Expert knowledge provides task-specific assumptions, sample and imaging-platform information, candidate morphological concepts and other experiment-specific knowledge. Relevant research articles were automatically retrieved from PubMed Central Open Access when their titles or abstracts matched at least one term from the sample types, biomarkers, perturbation terms and other biological keywords. Articles were converted into machine-readable text with PaddleX. A deep research report was prepared offline by Gemini deep research, or during runtime by Tongyi-DeepResearch-30B-A3B^17^.

### Feature design and dual-path quantification

MorphAgent builds a growing feature vocabulary through iterative feature design, quantification, and validation. In each round, MorphAgent uses a planner to combine the interactive environment, the biological knowledge base and feedback from previous rounds to propose candidate features. Each candidate feature is specified by a structured feature card containing its name, biological interpretation, expected visual signature, required channels or masks, candidate operators and summary statistics. Candidate features are then assigned to one of two execution pathways. Features with explicit quantitative definitions, such as area, intensity, texture, colocalization, spatial distribution or shape measurements, are measured by agent-generated code. Features with biologically meaningful but difficult-to-formalize phenotypes, such as subtle organelle organization, Tau architecture or higher-order biomarker patterns, are routed to the VLM-based semantic scoring pathway.

For the code-based pathway, MorphAgent generates an executable Python script conditioned on the feature card and dataset description file. Each script follows a fixed input-output contract: it receives the image and available masks, and returns one numerical feature value. Code generation is embedded in a bounded ReAct-style self-correction loop^48^. When execution fails, MorphAgent records the traceback and invokes the coding agent to revise the extractor. A maximum of five execution–correction cycles is allowed for each candidate feature, and candidates that still fail after five cycles are discarded. Each executable script is then reviewed by a VLM-based plausibility critic, which inspects representative images together with the quantified feature values to assess whether the numerical outputs are visually consistent with the intended feature definition. When inconsistencies are detected, MorphAgent enters a plausibility-correction loop of up to five cycles; candidates that remain unresolved are retained in the audit record with a warning flag rather than being automatically promoted to the validated feature set.

For the VLM-based pathway, MorphAgent prompts the VLM to assign each sample a score from 0 to 100 for the candidate feature, where 0 indicates absence of the visual phenotype and 100 indicates maximal presence within the visual range of the dataset. Scores are then normalized to 0–1 for downstream tasks. Each VLM response returns both a numerical feature score and a brief evidence-grounded rationale, allowing semantically rich phenotypes to be incorporated into the feature table while preserving interpretability and auditability.

### Feedback-driven feature validation and selection

After each round of feature design and quantification, MorphAgent removes duplicate candidates and evaluates the remaining features using predefined validation criteria. Features are first screened for sufficient across-sample variability, with a coefficient of variation (CV) greater than 0.05 used as the default retention threshold. Furthermore, any features sharing identical names or exhibiting a Pearson correlation coefficient exceeding 0.98 with existing features are classified as redundant and excluded.

When paired metadata are available, MorphAgent uses predefined dataset-specific validation metrics. These metadata may include perturbation identity, treatment condition, genotype or mutation label, age group, paired molecular measurements or other biological annotations. For categorical annotations, MorphAgent quantifies whether these features distinguish the annotation categories. For continuous or high-dimensional paired metadata, MorphAgent computes association scores between each morphology feature and metadata readout across samples. For example, given paired transcriptomic metadata, MorphAgent computes the maximum absolute Spearman correlation between each MorphAgent feature and transcriptomic variables, and features showing no correlation with any transcriptomic variable are removed. The exact validation metric and the code for executing the validation process for each dataset are automatically specified during the dataset preparation stage by MorphAgent. Features passing the validation criteria can further be ranked according to the validation metric, enabling MorphAgent to prioritize biologically informative features. The resulting validation report is fed back to the planner in the next iteration.

### Agent settings

In this study, the planning, code-generation and validation modules were implemented using Gemini 2.5 Pro. The planner used temperature = 0.3 to encourage hypothesis diversity, whereas code generation, validation and final scoring used deterministic decoding (temperature = 0.0). The VLM scorer and plausibility critic were implemented using Qwen3-VL-8B-Instruct with deterministic inference. Unless otherwise stated, each round proposed k = 20 candidate features. Detailed system prompts used in MorphAgent are available in Prompt 1. We further investigated the performance of MorphAgent across nine mainstream LLMs and five VLMs. In the Tau-genotype classification task, Gemini 2.5 Pro in the code-based pathway achieved the best overall performance, while Claude 4.8 Opus emerged as the top-performing VLM (Supplementary Fig. 20).

### Computational cost

We quantified the computational cost of MorphAgent separately for the code-based and VLM-scoring pathways (Supplementary Fig. 21). For the code-based pathway, we analyzed a representative five-round BBBC021 discovery run using stored prompt–response logs, from which the number of priced text-model API calls, input tokens and output tokens were recorded. Remote API cost was estimated by multiplying the total input and output token counts by the corresponding public model pricing schedule. In this representative run, the code-based pathway involved 106 priced text-model API calls and 718,748 total tokens, corresponding to an estimated remote API cost of approximately US$5.55. For the VLM-scoring pathway, local inference cost was quantified by recording the number of input images, number of feature scores, model configuration, hardware configuration and total wall-clock runtime. In a representative BBBC021 VLM-scoring run, a locally deployed Qwen3-VL-8B-Instruct model running on two NVIDIA GeForce RTX 4090 GPUs generated 177,600 feature scores for 3,552 images in 101.8 h.

### BBBC021 Cell Painting analysis

#### Feature-space coverage, compactness and interpretability

The coverage, compactness and interpretability of MorphAgent features relative to CellProfiler^11^ handcrafted features and DeepProfiler^16^ latent embeddings were assessed using the following criteria. For two representation spaces A and B, a feature or latent dimension in A was considered covered by B if its maximum absolute Spearman correlation with any feature in B exceeded 0.50 across matched image-level profiles. Coverage from A to B was reported as the fraction of features in A satisfying this criterion. Feature-space compactness after redundancy filtering was quantified within each representation space by iteratively pruning features until no retained pair had an absolute Pearson correlation greater than 0.90. The resulting non-redundant feature counts were 291 for MorphAgent, 440 for DeepProfiler and 1,284 for CellProfiler (Fig. 2a, b).

Feature interpretability was assessed by human expert review. For MorphAgent and CellProfiler features, the name and description of each feature were reviewed by three biologists using a five-point scoring scale, where 1 indicated no clear biological meaning and 5 indicated a fully biologically interpretable cellular or subcellular phenotype. Features with an average expert score greater than 3 were classified as biologically interpretable. To assess the fidelity of MorphAgent-derived measurements to their intended definitions, code-based features were evaluated using an LLM-as-a-judge implementation audit^49^ that assessed whether each generated script matched the corresponding feature name and description. For VLM-derived features, experts additionally assessed whether the semantic scores were grounded in visible image content by inspecting model attention maps highlighting image regions associated with the responses.

#### DeepProfiler feature interpretation

Each DeepProfiler dimension was assigned to a MorphAgent feature only when two criteria were met: their absolute Spearman correlation across matched image-level profiles exceeded 0.50, and the pair showed concordant perturbation discrimination, defined as a same-compound absolute Cohen’s d of at least 0.50. For a feature z and focal compound c, Cohen’s d was calculated as

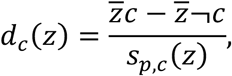

where 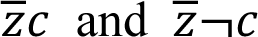 are the mean feature values for images treated with compound c and all other compounds, respectively, and *s_p_*_,*c*_(*z*) is their pooled standard deviation. Cohen’s d was calculated separately for the DeepProfiler dimension and the MorphAgent feature. Because feature matching was based on absolute correlation, the direction of the compound response was considered concordant when

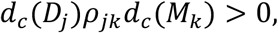

where *ρ*_j*k*_ is the Spearman correlation between DeepProfiler dimension *D*_j_ and MorphAgent feature *M_k_*. Among all qualifying pairs, the MorphAgent feature with the largest absolute Spearman correlation was used for annotation (Fig. 2c–e).

### Benchmark tasks

We evaluated CellProfiler, two DeepProfiler variants (the original model pretrained on large-scale Cell Painting data and a variant retrained solely on the 3,552 BBBC021 images; see Supplementary Methods) and MorphAgent on four Cell Painting tasks: perturbation detection, image-level MoA prediction, same-MoA perturbation matching and L1000 transcriptomic regression (Fig. 2f–i). For perturbation detection, each compound was treated as a one-versus-rest retrieval task, using same-compound image-level profiles as positives and all other profiles as negatives. Profiles were ranked by cosine distance, and performance was measured by mAP, computed as:

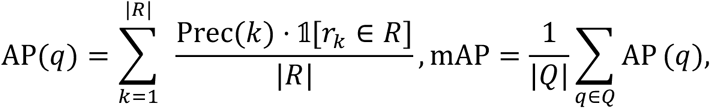

where Prec(k) is the precision at rank k, R is the set of profiles sharing the compound identity of q, and Q is the image dataset. The analytic random baseline for 37-way matching is D[AP] ≈ 0.027. Uncertainty was estimated using 2,000 bootstrap resamples and reported as 95% bootstrap percentile confidence intervals.

For MoA prediction, image-level profiles were classified into 26 MoA categories using k-nearest-neighbour classification ( *k* is selected automatically by cross-validation) with cosine distance under five-fold cross-validation. For same-MoA perturbation matching, we considered the seven MoA groups containing at least two compounds. For each valid query, one compound served as the query, another compound with the same MoA served as the positive, and five compounds with different MoAs formed the negative candidate set, yielding a six-way ranking task. Performance was reported as mAP averaged across all valid same-MoA query pairs, and the analytic random baseline for six-way matching is D[AP] ≈ 0.408. For L1000 regression, morphology profiles were paired with L1000 gene-expression profiles from the same compound and concentration conditions. The resulting image-level morphology–transcriptome pairs were divided into training, validation and test sets at a ratio of 80:10:10. A multilayer perceptron with two hidden layers of 512 and 256 ReLU units and a dropout rate of 0.3 was trained under mean squared error loss to predict the 977 L1000 landmark genes. Training was performed for 300 epochs, and the checkpoint with the lowest validation loss was evaluated on the held-out test set. Performance was reported as a single global *R*^2^ across all test samples and genes. Additional implementation details are provided in the Supplementary Methods.

### HSC mitochondrial imaging analysis

#### Aged-cell discrimination task on confocal HSC dataset

We used MorphAgent-derived mitochondrial features to classify young versus aged HSCs. For the HSC discovery dataset, paired single-cell Smart-seq2 transcriptomic profiles were available and were used to construct a transcriptomic reference baseline. Genes were ranked by expression variance across the 110 discovery cells, and the top 500 most variable genes were retained as the HVG500 transcriptomic feature set. This transcriptomic baseline was evaluated using the same 2D PCA projection and L2-regularized logistic regression framework as the image-derived features. For MorphAgent, features were first restricted to valid numerical values with sufficient variance. The remaining features were pre-ranked using one-way ANOVA F statistics for the young-versus-aged labels in the 110-cell discovery dataset. Greedy forward selection was then performed. At each step, each remaining candidate was added in turn to the selected feature set. The resulting feature set was projected into two dimensions by PCA and evaluated using L2-regularized logistic regression. Balanced accuracy was the primary selection criterion, and AUC was used only to break ties. The first 25 features in the resulting selection order were retained as the fixed mitochondrial feature vocabulary and locked before analysis of the independent validation dataset (Fig. 3a, d).

The synthetic low-resolution representations were generated from the confocal maximum-intensity projections by convolution with a normalized two-dimensional Gaussian kernel with a full width at half maximum (FWHM) of 200 nm, corresponding to a standard deviation of 84.9 nm (approximately 2.22 pixels at 38.2 nm per pixel). Assuming Gaussian lateral point spread functions and an original nominal lateral resolution of approximately 120 nm, this convolution yielded a nominal effective lateral resolution of approximately 233 nm, calculated as 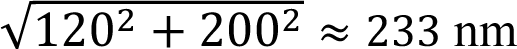.

#### STRING-based network proximity analysis

To compare the molecular network context of morphology-associated genes, each gene was assigned an association score defined as the maximum absolute Spearman correlation between its expression and any MorphAgent mitochondrial feature across the image–transcriptome-paired HSC discovery dataset. Genes were ranked by this score, and those in the top and bottom 10% were mapped to the STRING protein–protein interaction network^21^. Within the retained network, Trp53 had the highest node degree and was therefore used as the reference node for proximity analysis. For each mapped gene connected to Trp53, the shortest-path distance to Trp53 was calculated and grouped as direct interaction (distance = 1), one intermediate node (distance = 2), or at least two intermediate nodes (distance ≥ 3). Genes lacking a STRING mapping or lying outside the Trp53-connected component were excluded from the shortest-path distribution analysis. The proportions of genes in these distance categories were compared between the top and bottom 10% groups (Fig. 3c).

### Tau super resolution imaging analysis

#### Tau expert feature comparison

The MorphAgent Tau features were first compared with the 16 expert-designed Tau phenotype features in the 58-cell discovery dataset. For each expert feature, recovery by MorphAgent was quantified as the maximum absolute Spearman correlation with any MorphAgent feature across cells; an expert descriptor was considered recovered when this value exceeded 0.70. Conversely, a MorphAgent feature was considered weakly represented by the expert feature vocabulary when its maximum absolute Spearman correlation with all 16 expert features was below 0.50. For Fig. 4c, these weakly represented MorphAgent features were grouped into structural themes using GPT-5.5 Pro, and the resulting assignments were reviewed by human experts.

#### Mutation state classification

Mutation state classification was evaluated on the 495-cell COS-7 dataset using paired SIM and wide-field images of the same cells. The same MorphAgent features were applied independently to the two image modalities. For binary WT-versus-P301S+S320F classification, features were standardized using statistics calculated from the training fold, and an L2-regularized logistic regression classifier was trained and evaluated under five-fold cell-level cross-validation. Identical cell folds and classifier settings were used for SIM and wide-field comparisons and for the MorphAgent and expert-designed feature vocabulary. Performance was quantified by accuracy and area under the receiver operating characteristic curve (AUC). Six-genotype classification across WT, S320F, S305I, P301L, P301L+S320F and P301S+S320F was evaluated under the same five-fold cell-level split structure, and performance was summarized by overall accuracy and macro-FPR.

#### Resolution dependency

Resolution-dependent feature behavior was analyzed in three ways. First, for each feature, the cross-cell coefficient of variation was computed separately from paired SIM and wide-field measurements, and the SIM-to-wide-field change was summarized as log2(CVSIM/CVWF) (Fig. 4f). Second, individual features were evaluated as single-feature classifiers using the same five-fold cross-validation framework. Within each training fold, features were ranked by their mutation discrimination accuracy, and SIM and wide-field accuracies were compared for the 50 highest-ranked features on the corresponding held-out cells (Fig. 4g). Third, reconstructed SIM images, with an estimated lateral resolution of approximately 100 nm, were progressively smoothed using normalized two-dimensional Gaussian kernels with FWHM values of 100, 200, 400, 800, 1,600 and 3,200 nm (Supplementary Fig. 13). Assuming Gaussian point spread functions, these blur-kernel scales corresponded to estimated effective lateral resolutions of approximately 141, 224, 412, 806, 1,603 and 3,202 nm, respectively. MorphAgent features were re-extracted at each blur level, and WT-versus-P301S+S320F classification was repeated using the same five-fold cross-validation protocol. Matched wide-field images were generated by averaging the 15 raw SIM frames used for SIM reconstruction (Fig. 4h, i).

#### SIM morphology-based transcriptomic prediction

MorphAgent redesigned the feature vocabulary using transcriptomic profiles as paired metadata for validation, yielding an expanded set of 400 morphology features. Each target gene was predicted separately using 10 repeated train–test splits. In each repeat, 80% of the cells were assigned to the training set and the remaining 20% to the test set. Candidate MorphAgent features were evaluated individually by five-fold cross-validation within the training set. For each cross-validation fold, feature standardization and a univariate SVR model with an RBF kernel (C = 1.0, gamma = scale, and epsilon = 0.1) were fitted using four folds and evaluated on the remaining fold. Features were ranked by their mean validation Pearson correlation across the five folds, and the 16 highest-ranked features were selected for each gene. A final multivariate SVR model was then fitted using these 16 features on the complete training set and evaluated on the corresponding test set. Gene-level prediction performance was calculated as the mean test set Pearson correlation across the 10 repeated splits.

Two reference feature sets were evaluated using the same train–test splits and model fitting procedure. The expert feature model used the 16 manually defined Tau morphology features (Supplementary Table 2), whereas the Tau intensity baseline used whole-cell Tau intensity as a single input feature. Prediction performance was first averaged across the 10 test splits for each gene. Bar heights represent the mean of these gene-level Pearson correlations across all genes in each target-gene set, and error bars represent the standard error of the mean across genes. Pairwise comparisons between methods were performed using Welch’s t-test on the gene-level mean Pearson correlations.

#### Bidirectional morphology–transcriptome prediction

Transcriptome-to-morphology prediction was evaluated analogously, with each MorphAgent feature treated as a prediction target and transcriptomic measurements used as candidate predictors. Predictor ranking, scaling, model selection and evaluation were nested within the same repeated split structure used for morphology-to-gene prediction. For both directions, the distribution of target-level Pearson correlations was summarized, and targets with mean Pearson correlation greater than 0.50 were counted as strongly predictable (Fig. 5b).

#### Biologically grounded three-level morphology–gene hierarchy

We built parallel three-level hierarchies for both morphology features and genes. On the morphology side, MorphAgent Tau features were organized from local features (bottom) to cellular compartments (middle) and to morphology-defined biological processes (top). The hierarchical grouping of features was assisted by a large language model (GPT-5.5 Pro) and verified by three human experts. For the top level, MorphAgent features were grouped into morphology-defined biological processes according to the biological process represented by their feature description. For the middle level, MorphAgent features in each morphology-defined biological process were grouped according to the cellular compartment or spatial scale of the measured structure. On the gene side, genes were organized from individual genes (bottom) to regulatory modules (middle) and to GO-enriched biological processes (top). For the top level, GO terms were obtained by over-representation testing of the top 1,000 genes (*clusterProfiler::enrichGO*; biological process and cellular component ontologies; Supplementary Methods). For the middle level, we inferred a GRN from the Smart-seq2 expression profiles of the 58 WT cells using GRNBoost2^50^. 109 transcription factors within the top 1,000 genes acted as candidate regulators, whereas all genes were candidate targets in our analysis. To reduce stochastic variation in network inference, we ran GRNBoost2 50 times using distinct random seeds. In each run, GRN edges were ranked by their importance scores, and only those ranked top 10% in at least 40 of the 50 runs were retained in the final GRN.

#### Correspondence map

Correspondence maps at the top level (Fig. 5d) and middle level (Fig. 5e) were constructed using the same gene-overlap procedure. As described in the morphology–transcriptome prediction analysis, the 16 most predictive MorphAgent features were identified for each gene. We then inverted this relationship to associate each feature with the genes for which it was selected as a top predictor. Because individual features were further organized hierarchically into cellular compartments and biological processes, gene sets could be aggregated at each level of the hierarchy. Correspondence between morphology and gene modules was quantified by counting the number of genes shared between the corresponding gene sets. The P value for each cell in the correspondence maps was further calculated using the one-sided permutation test (Supplementary Methods and Supplementary Figs. 15 and 16).

## Code availability

The code for implementing MorphAgent and reproducing the analyses is openly available at https://github.com/ai4imaging/MorphAgent. An interactive UI demo and usage instructions are also provided in this repository.

## Supporting information

Algorithm

Prompt

Supplementary feature list 1

Supplementary feature list 2

Supplementary feature list 3

Supplementary feature list 4

Supplementary feature list 5

Supplementary feature list 6

