## Supplementary Information for "Biologically grounded cell profiling across microscopy modalities"

### Supplementary Methods

#### Wide-field Cell Painting analyses

**Perturbation detection.** Perturbation detection was formulated as image-level compound retrieval. Each image served as a query; positives were all other images sharing the same compound identity; all remaining images were negatives. The query image was excluded from its own ranking. Profiles were ranked by cosine distance. Mean average precision (mAP) was computed as:

$$\mathrm{AP}(q) = \sum_{k=1}^{|R(q)|} \frac{\mathrm{Prec}(k) \cdot \mathbb{1}[r_k \in R(q)]}{|R(q)|}, \mathrm{mAP} = \frac{1}{|Q|} \sum_{q \in Q} \mathrm{AP}\,(q),$$

where $Q$ is the set of query images, $R(q)$ is the set of same-compound positives for query $q$, $r_k$ is the profile retrieved at rank $k$, and $Prec(k)$ is precision at rank $k$. The analytic random baseline for this 37-compound matching task was $\mathbb{E}[AP] \approx 0.027$. Task performance was summarized as the mean of per-query AP values; 95% CIs were obtained from 2,000 bootstrap resamples over queries.

**Mechanism-of-action (MoA) prediction.** MoA prediction was performed at the image level. Each image was assigned one of 26 MoA labels. Classification used $k$-nearest-neighbour (kNN) classification with cosine distance under five-fold profile-level cross-validation, where $k$ is selected automatically by cross-validation. Features were standardized using training-fold statistics only. Accuracy was calculated from the pooled out-of-fold predictions across the five folds, and 95% percentile confidence intervals were obtained from 2,000 bootstrap resamples of the image-level predictions.

**Same-MoA perturbation matching.** Same-MoA matching was restricted to the seven MoA categories containing at least two compounds. For each ordered pair of distinct compounds $(d_q, d_+)$ sharing a MoA, one compound served as the query source $(d_q)$ and the other as the positive candidate $(d_+)$. For each ordered query–positive pair, five negative compounds were sampled randomly, with each negative drawn from a different MoA category. Each compound $d$ was represented by its mean image-level profile:

$$\overline{x}_d = \frac{1}{|\mathcal{I}_d|} \sum_{i \in \mathcal{I}_d} x_i,$$

where $\mathcal{I}_d$ is the set of images for compound $d$. For each query image from $d_q$, cosine distance was computed to the six candidate centroids. With exactly one positive among six candidates, average precision reduces to:

$$AP_i = \frac{1}{\mathrm{rank}(d_+)}.$$

AP values were averaged over query images within each valid $(d_q, d_+)$ pair, then over all valid pairs. The analytic random baseline for six-way ranking with one positive is:

$$\mathbb{E}[AP_{\text{6-way,1-pos}}] = \frac{1}{6}\sum_{j=1}^{6} \frac{1}{j} \approx 0.408.$$

Results were aggregated over all valid query–positive pairs; 95% CIs were obtained using 2,000 bootstrap resamples at the pair level.

**L1000 regression.** Morphology profiles were paired with L1000 gene-expression profiles from the same compound and concentration conditions for the shared BBBC021 compounds. When multiple image profiles mapped to the same compound–concentration signature, each image retained the corresponding condition-level expression profile, yielding image-level morphology–transcriptome pairs.

The prediction target comprised the 977 L1000 landmark genes. The paired samples were randomly partitioned into 80% training, 10% validation and 10% test sets using a fixed random state of 42. For each morphology representation, a multilayer perceptron with two hidden layers of 512 and 256 units, respectively, ReLU activation and a dropout rate of 0.3 was trained to predict the 977-dimensional L1000 expression vector. The model was optimized under mean squared error loss using Adam with a learning rate of $10^{-3}$, weight decay of $10^{-5}$ and a batch size of 128. Training was performed for the full 300 epochs without early stopping. Model performance on the validation set was recorded after each epoch, and the checkpoint with the lowest validation MSE was retained for evaluation on the held-out test set.

The reported $R^2$ was calculated as a single global score across all test samples and the 977 landmark genes:

$$R^2 = 1 - \frac{\mathrm{MSE}\,(\hat{y}, y)}{\mathrm{MSE}\,(\bar{y}_{\mathrm{test}}, y)},$$

where $y$ and $\hat{y}$ denote the measured and predicted expression profiles, respectively, and $\bar{y}_{\mathrm{test}}$ denotes the vector of per-gene mean expression values in the test set. Ninety-five percent confidence intervals were obtained from 2,000 bootstrap resamples of the held-out test samples, with the global $R^2$ recalculated for each resample. Because the random image-level split could place profiles associated with the same compound–concentration condition in different data partitions, this analysis was not interpreted as out-of-compound or out-of-condition generalization.

**DeepProfiler baseline.** Two DeepProfiler variants were benchmarked, each producing a 672-dimensional embedding. Because BBBC021 contained three fluorescence channels whereas the DeepProfiler pipeline expected five-channel Cell Painting input, the channels were mapped as follows: the DNA channel was assigned to the DNA input, the tubulin channel to the endoplasmic-reticulum input and the actin channel to the AGP input; the RNA and mitochondrial inputs were set to zero. The same channel mapping was used for both DeepProfiler variants. **(i) DeepProfiler (pretrained).** The publicly released Cell_Painting_CNN_v1 checkpoint with an EfficientNet backbone, pretrained on a large external Cell Painting dataset, was applied to BBBC021 without further training. **(ii) DeepProfiler (trained on BBBC021).** The same EfficientNet architecture was trained de novo on the 3,552 BBBC021 images. Training used treatment-identity classification for 30 epochs with a sampled-crop generator and on-the-fly augmentation. The learning rate was 0.005, the batch size was 32, label smoothing was set to 0 and no additional convolutional blocks were used. Training was monitored using accuracy, top-five accuracy and average-class precision. All runs followed the standard DeepProfiler framework configuration.

**Unsupervised representation-learning baselines.** We compared MorphAgent with two label-free deep representation-learning methods: CellPaintSSL and MorphoGenie on BBBC021. Because the officially released weights could not be retrieved in our computational environment, both models were trained directly on the 3,552 BBBC021 images. Their results therefore represent in-domain, label-free representations learned from a single mid-sized dataset rather than large-scale pretrained foundation models.

CellPaintSSL[1] is a self-supervised Cell Painting framework that learns image embeddings using the DINO objective with a Vision Transformer backbone. We used a ViT-Small/16 encoder, which generated a 384-dimensional embedding. Because BBBC021 contained three fluorescence channels whereas the framework expected five-channel Cell Painting input, the DNA, tubulin and actin channels were assigned to the DNA, endoplasmic-reticulum and AGP inputs, respectively, and the RNA and mitochondrial inputs were set to zero. The DINO model was trained from scratch on the BBBC021 images for 80 epochs. Optimization used AdamW with a batch size of 32 images per GPU. The reference learning rate of 0.004, defined for a batch size of 256, was linearly scaled to an effective peak learning rate of approximately $5 \times 10^{-4}$ for a batch size of 32. Because CellPaintSSL used the DINO objective, no KL-divergence or VAE regularization term was applied. At inference, tile-level embeddings were extracted using the teacher encoder and averaged to obtain one field-level profile per image. The resulting profiles were post-processed using per-plate ZCA spherizing followed by median-absolute-deviation robust normalization with pycytominer.

MorphoGenie[2] learns a disentangled representation of cellular morphology using a FactorVAE-style generative model. We used the FactorVAE256 fallback implementation, which generated a ten-dimensional latent representation. Single cells were extracted from connected components of the nuclear mask, retaining up to the ten largest cells in each field by nuclear area. The three BBBC021 channels were arranged as an RGB image in DNA–tubulin–actin order. Each selected cell was cropped and resized to $256 \times 256$ pixels. The model was trained from scratch for 30 epochs using Adam with a batch size of 64 and a learning rate of $10^{-4}$. The training loss was the sum of the mean squared reconstruction loss and the KL-divergence term:

$$\mathcal{L} = \mathcal{L}_{\text{reconstruction}} + \beta \mathcal{L}_{\text{KL}},$$

with $\beta = 1$, because no additional multiplier was applied to the KL term. Features were extracted as the encoder posterior mean $\mu$ for each single-cell crop and averaged across the retained cells to produce one ten-dimensional profile per image.

**STRING-based Trp53 proximity analysis.**

To compare the molecular network context of morphology-associated genes, each gene was assigned an association score

$$s_g = \max_{f \in F_{25}} | \rho_S(g,f) |$$

where $F_{25}$ denotes the fixed top-25 MorphAgent feature vocabulary and $\rho_S(g,f)$ is the Spearman correlation between gene $g$ and feature $f$ across the paired discovery cells. Genes were ranked by $s_g$, and only the top 10% and bottom 10% groups were compared.

The 500 HVGs were mapped to the STRING v12.0 network[3]. Gene identifiers were matched to STRING protein identifiers, and only interactions with a combined score ≥ 0.4 were retained, yielding 354 mapped nodes and 3,936 edges. Trp53 had the highest node degree in this retained input-restricted subgraph and was therefore used as the reference node; it was not pre-specified independently of the network. Disconnected nodes were excluded from shortest-path analysis, leaving 329 genes in the Trp53-connected component. For each extreme group, the denominator was the number of group genes mapped to this connected component. Shortest-path distances to Trp53 were binned as direct interaction (distance = 1), one intermediate node (distance = 2), or at least two intermediate nodes (distance ≥ 3).

**Gene Ontology enrichment.**

Gene names were normalized by case folding, removal of punctuation-dependent variation and whitespace harmonization, mapped to official human gene symbols using the paired transcriptomic annotations, and converted to Entrez identifiers using org.Hs.eg.db. Among all 27,476 measured genes, 19,810 were protein-coding genes detected by sequencing and were used as the sampling universe for permutation testing. All query genes were matched to both an official gene symbol and an Entrez identifier.

GO enrichment[4] was performed with *clusterProfiler::enrichGO*[5] separately for biological-process (BP) and cellular-component (CC) ontologies. For each ontology, enrichment was formulated as a one-sided over-representation test. For a GO term containing $K$ genes among the background of $N = 19{,}810$ Entrez-mapped genes and for a query set containing all query genes, the observed overlap $x$ was evaluated using the upper tail of the hypergeometric distribution, $P(X \geq x)$. GO terms with gene-set sizes between 5 and 500 genes were tested. Raw $P$ values were adjusted within each ontology using the Benjamini–Hochberg procedure. Terms were considered significant when the BH-adjusted P-value was less than 0.05. GeneRatio was reported as $\mathrm{x/n}$, with genes in each GO-term converted back to readable gene symbols. This analysis identified 19 significant GO terms, including 6 BP terms and 13 CC terms. For visualization, 9 mitochondrion-related terms were consolidated into a single category, resulting in 11 displayed GO categories.

**Permutation test for correspondence maps.**

Significance was evaluated with a one-sided permutation test using the 19,810 detected protein-coding genes as the sampling universe. Before testing, both the morphology-defined gene set and the fixed GO-enriched gene set were restricted to this universe. For a morphology-defined gene set of size m, 10,000 random sets of size m were sampled without replacement, and their overlap with the fixed GO set was calculated. The empirical P value was

$$P_{\mathrm{emp}} = \frac{1 + \sum_{b=1}^{B} 1(O_b \geq O_{\mathrm{obs}})}{B + 1}, B = 10{,}000.$$

Nominal empirical P values were reported without multiplicity correction.

### Supplementary Figures

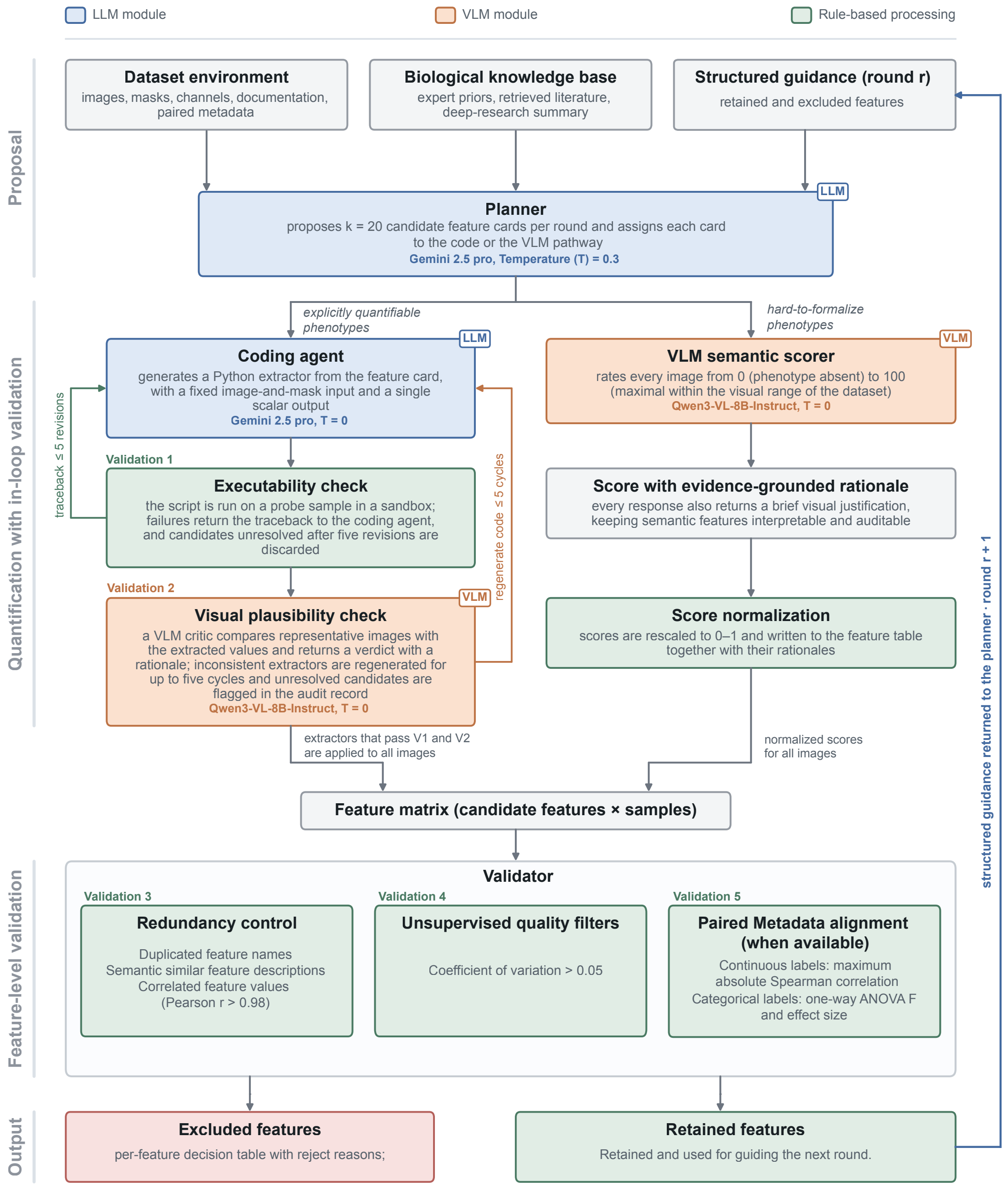


**Supplementary Fig. 1.** Schematic overview of the MorphAgent workflow, in which an LLM planner proposes candidate feature cards that are quantified through the code or the VLM pathway, filtered by five validation checkpoints (V1–V5: executability, VLM-based visual plausibility, redundancy, unsupervised quality and metadata alignment), and returned to the planner as structured guidance for the next round.

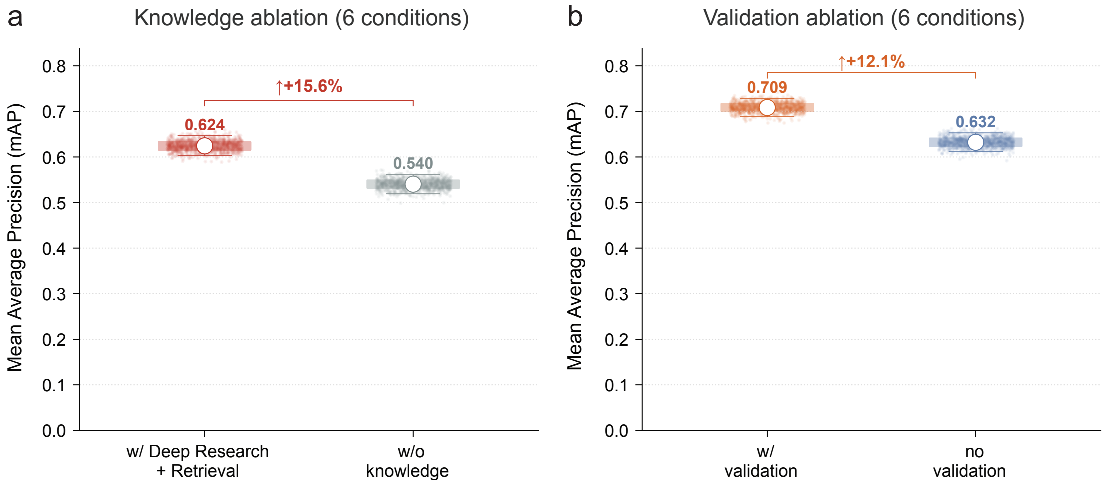


**Supplementary Fig. 2.** Knowledge and validation ablations on a BBBC021 subset (n = 568 images across six conditions: five compound treatments and DMSO controls). **(a)** Contribution of external biological knowledge to MorphAgent feature discovery. Knowledge-guided feature discovery (deep research and literature retrieval) improved perturbation-detection performance from 0.540 to 0.624 mAP, corresponding to a 15.6% relative increase. **(b)** Validation ablation on the same 6-condition subset shows that enabling compound-based validation improved mAP from 0.632 to 0.709 (+12.1%).

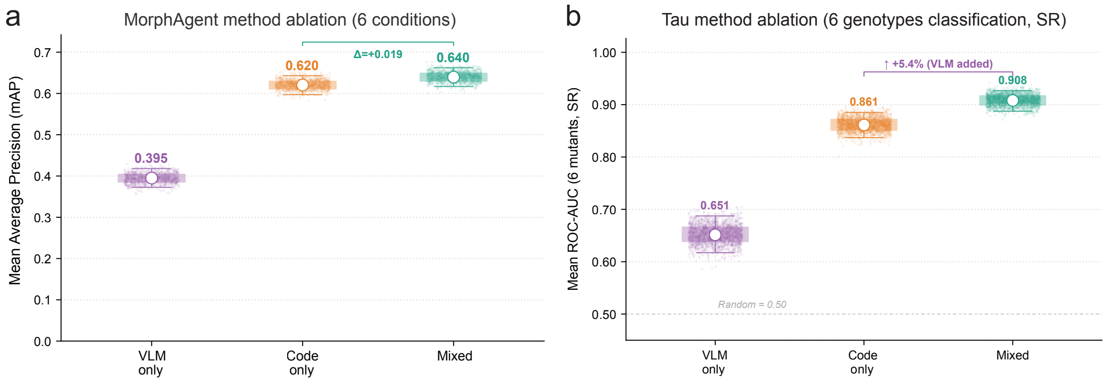


**Supplementary Fig. 3.** MorphAgent method ablation across BBBC021 and Tau six-genotype classification. **(a)** Perturbation-detection performance on a BBBC021 subset (n = 568 images across six conditions) for VLM-only features, code-only features and mixed features, reported as mean average precision (mAP). Although the VLM-only features exhibit limited standalone performance, integrating them with code features yields a measurable improvement over the code-only baseline, increasing the mAP from 0.620 to 0.640 ($\Delta = +0.019$). **(b)** Analogous method ablation for Tau morphology–based genotype classification on super resolution images, evaluated as mean ROC-AUC across six WT-versus-mutant tasks. VLM-only features again underperform in isolation (AUC = 0.651), whereas combining VLM and code features improves performance relative to the code-only baseline, raising mean AUC from 0.861 to 0.908 (+5.4%). Points indicate resampling estimates, open circles denote means, and the dashed line in (b) marks the random baseline (AUC = 0.50).

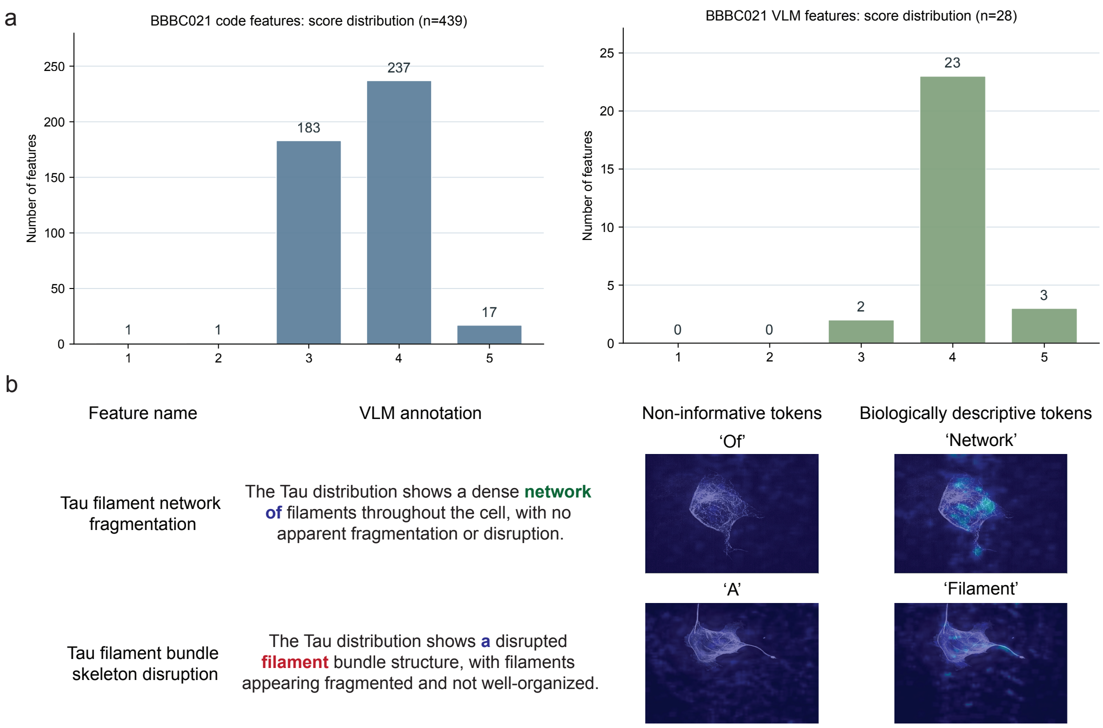


**Supplementary Fig. 4.** Technical review and visual grounding of MorphAgent features. **(a)** Left, BBBC021 code-based feature extractors ($n$ = 439) were assessed using an LLM-as-a-judge framework that reviewed consistency with the intended feature definition, executability on the target image inputs and production of interpretable numerical measurements; 437/439 received a score of ≥3 (1-5 integer scale). Right, BBBC021 VLM-derived semantic features ($n$ = 28) were assessed under the same framework; all 28/28 received a score of ≥3. **(b)** Token-activation maps provide a qualitative visual-grounding analysis for VLM-derived phenotypes, showing that high-activation regions in representative examples overlap with relevant Tau-positive structures such as networks and filament bundle structures.

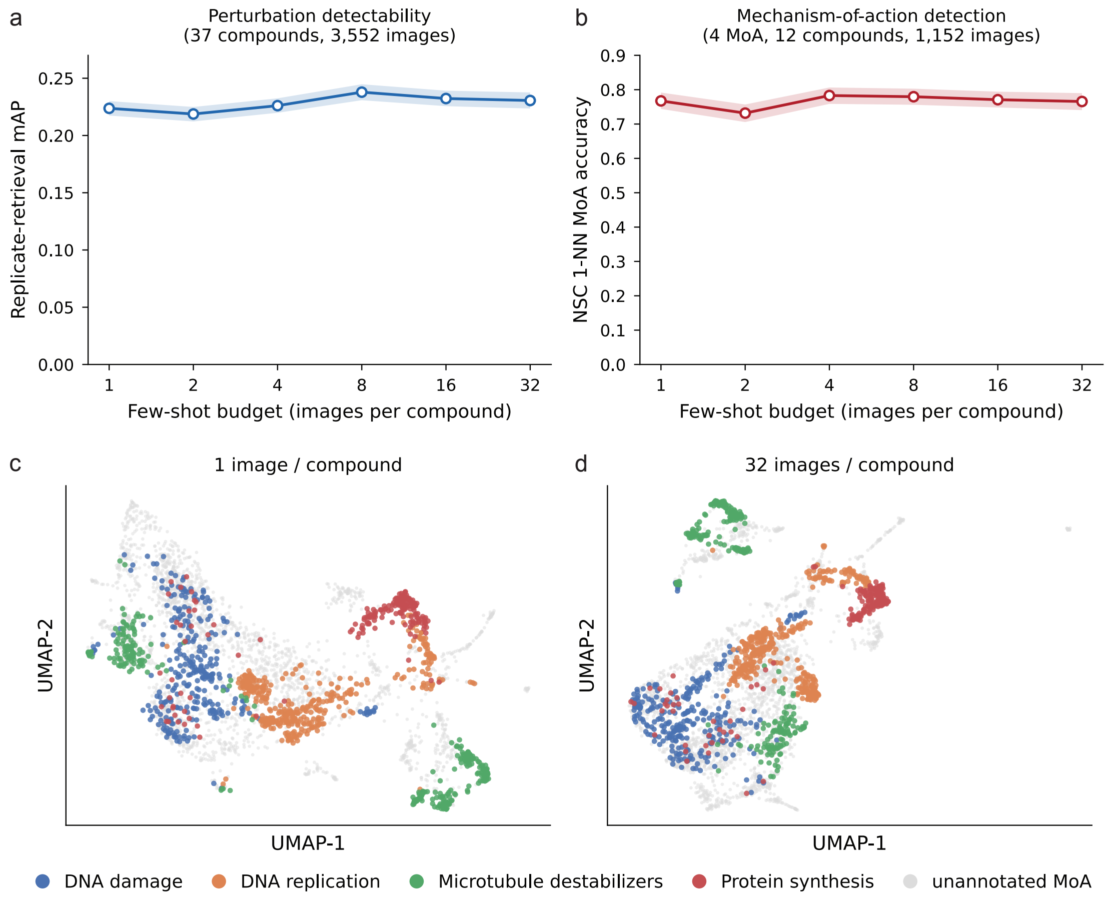


**Supplementary Fig. 5.** Few-shot-designed morphological feature sets detect perturbations and mechanisms with high data efficiency on BBBC021. Feature sets were designed using 1–32 reference images per compound and then applied without redesign to all 3,552 BBBC021 MCF7 fluorescence images. **(a)** Perturbation detectability, quantified by same-compound replicate-retrieval mAP across all 37 compounds and 3,552 images. **(b)** Mechanism-of-action evaluation using not-same-compound (NSC) 1-nearest-neighbour classification, restricted to four MoA classes represented by at least two compounds: DNA damage, DNA replication, microtubule destabilization and protein synthesis (12 compounds; 1,152 images). This evaluation is distinct from the 26-class, five-nearest-neighbour MoA benchmark in Fig. 2g. In (a) and (b), points indicate means, and shaded bands indicate 95% bootstrap confidence intervals based on 2,000 resamples of the per-image scores. **(c,d)** UMAP embeddings of the full dataset for the smallest (one image per compound, c) and largest (32 images per compound, d) budgets. Points represent individual images and are coloured by MoA; grey indicates images without an annotated MoA. Performance was similar across the tested budgets, indicating that in the evaluated runs, a feature vocabulary designed from one reference image per compound approached the performance obtained with larger reference-image budgets.

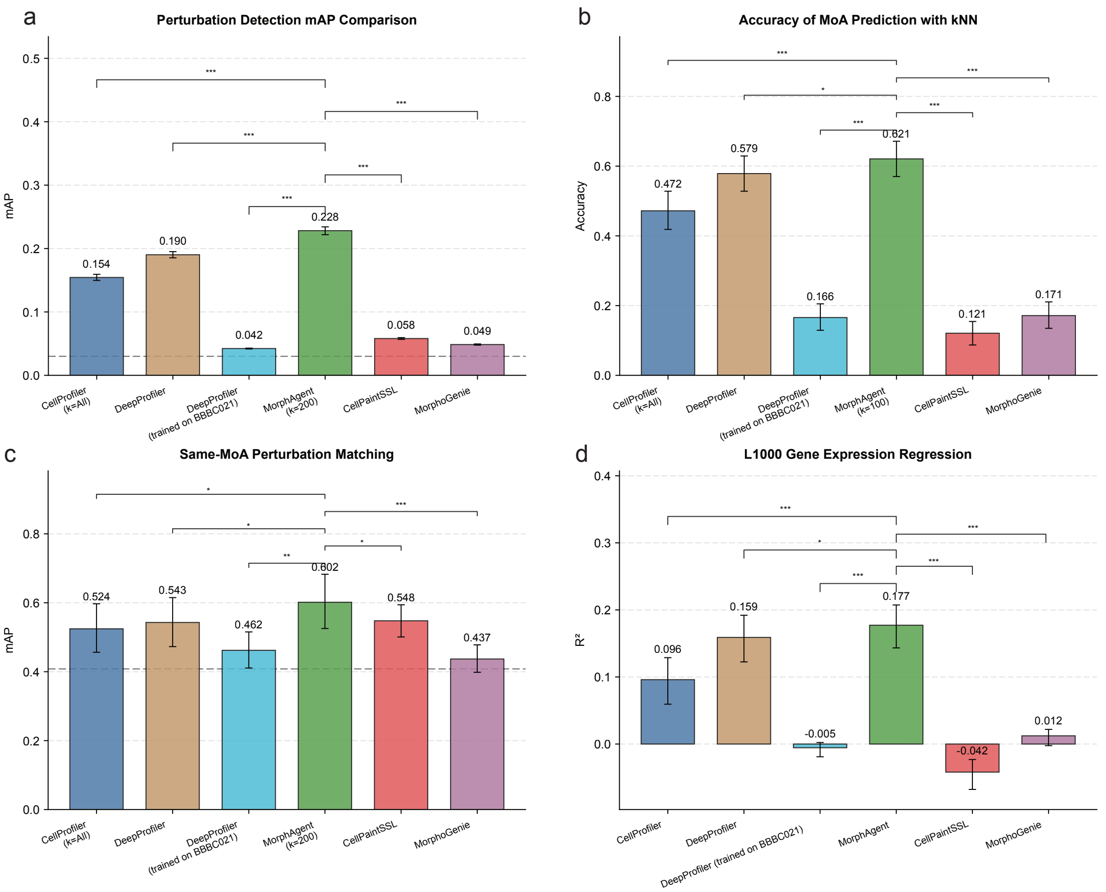


**Supplementary Fig. 6.** Benchmarking on BBBC021 including unsupervised representation-learning baselines. Performance of six morphological feature methods on BBBC021 (3,552 image-level profiles, 37 compounds): CellProfiler[6], pretrained DeepProfiler[7], DeepProfiler retrained on BBBC021, MorphAgent, and two label-free deep baselines: CellPaintSSL[1] and MorphoGenie[2]. **(a)** Perturbation detection (image-level same-compound retrieval, mAP). **(b)** Mechanism-of-action prediction (kNN classification, accuracy). **(c)** Same-MoA perturbation matching (compound-level retrieval among perturbations sharing a MoA, mAP). **(d)** L1000 transcriptomic regression (prediction of paired L1000 signatures by a shared MLP, $R^2$). Bars show point estimates and error bars show 95% bootstrap confidence intervals (2,000 resamples). The two unsupervised baselines were trained directly on BBBC021 and show generally weak performance across all four tasks.

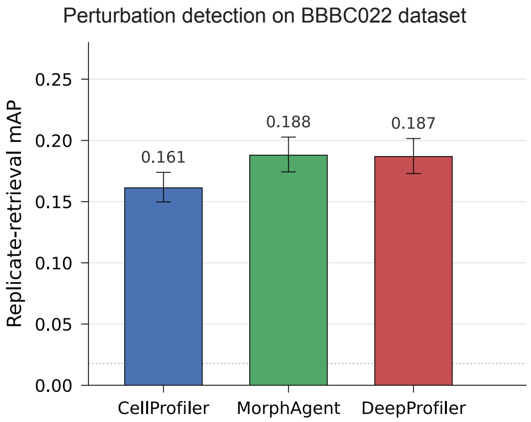


**Supplementary Fig. 7.** Perturbation retrieval on the BBBC022 compound-treated subset. Replicate-retrieval mAP was evaluated for CellProfiler, MorphAgent and DeepProfiler using 450 compound-treated image fields from 50 compounds on plate 20585. The 576 mock-control fields were used for normalization but excluded from retrieval. Each query had eight positive fields from the same compound.

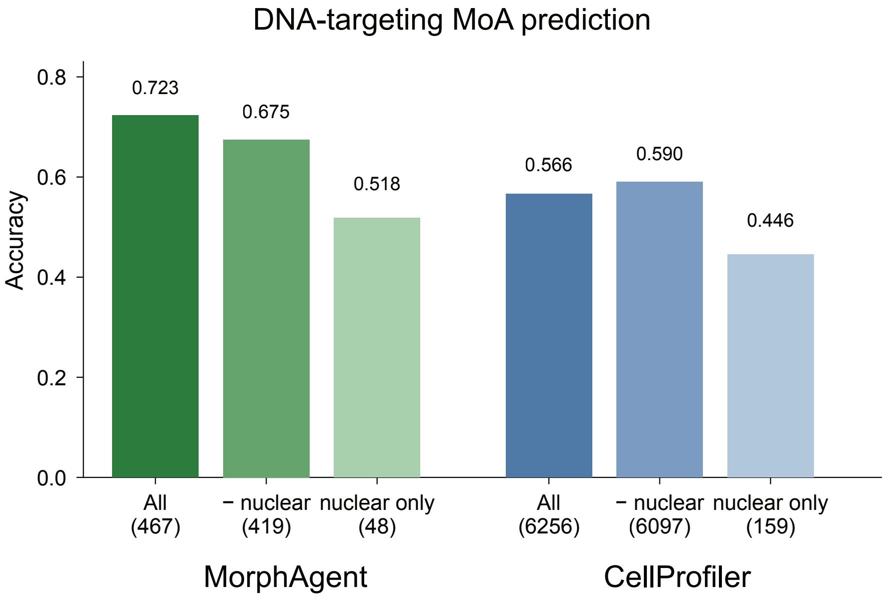


**Supplementary Fig. 8.** K-nearest-neighbour MoA prediction on DNA-targeting test images, with and without nuclear-associated features. Bars show all features, nuclear features removed, and nuclear features only, for MorphAgent (467 / 419 / 48 features) and CellProfiler (6,256 / 6,097 / 159). The drop in MorphAgent after nuclear feature removal indicates that this feature family is enriched for DNA-targeting signal, whereas CellProfiler does not drop; MorphAgent nuclear-only also far exceeds CellProfiler nuclear-only (0.518 vs 0.446).

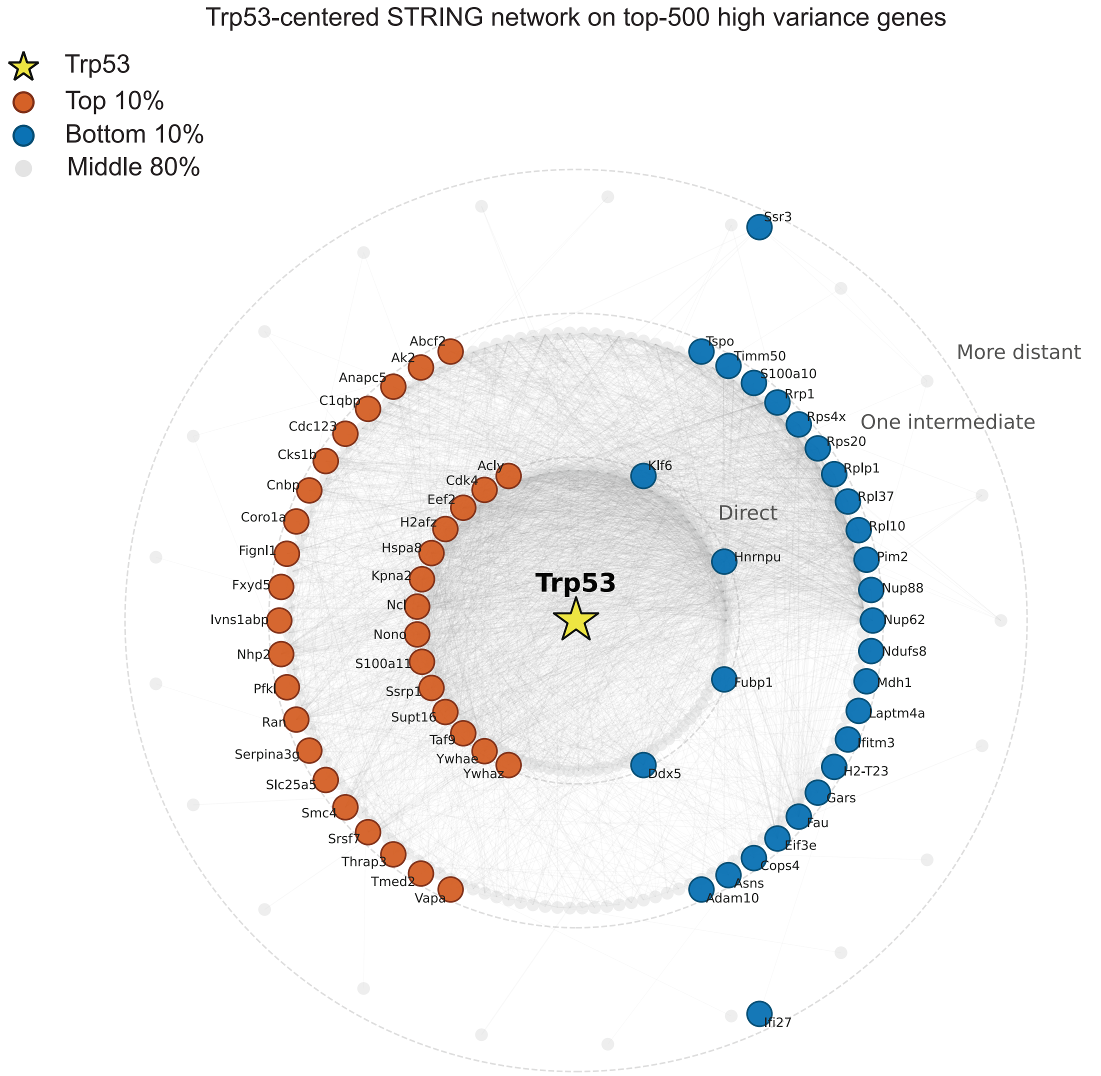


**Supplementary Fig. 9.** Trp53-centered STRING interaction network[3] constructed from the top 500 highly variable genes in the HSC discovery dataset. Genes were ranked by their maximum absolute Spearman correlation with the top MorphAgent-derived mitochondrial features. Nodes corresponding to the top 10% and bottom 10% morphology-associated genes are highlighted, while the remaining genes are shown in grey. Network distance from Trp53 is indicated by concentric layout levels, distinguishing direct, intermediate and more distant network neighborhoods. Trp53 was selected post hoc as the highest-degree node in the retained input-restricted network.

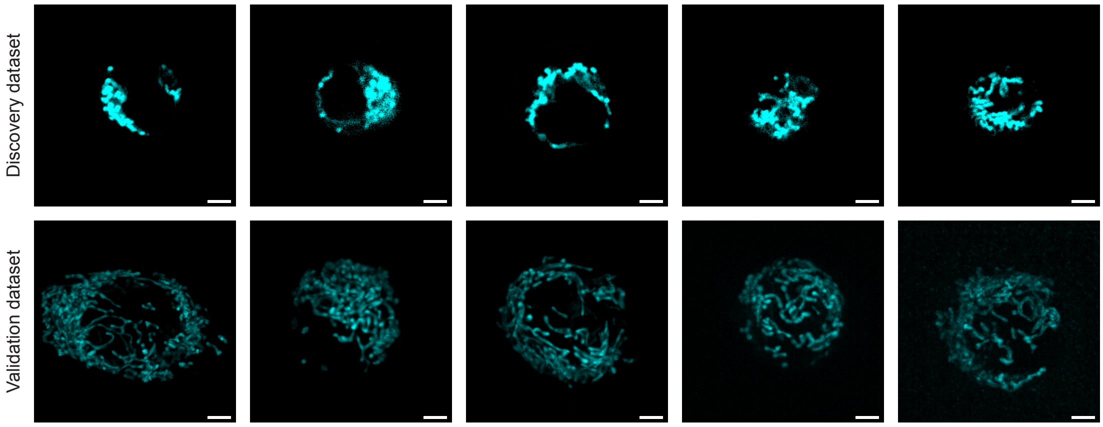


**Supplementary Fig. 10.** Representative mitochondrial images of single HSCs from the discovery dataset (top) and validation dataset (bottom). Both datasets were acquired by spinning-disk confocal microscopy. The validation images show higher image quality and finer visible mitochondrial-network detail than the discovery images. Scale bars, 5 μm.

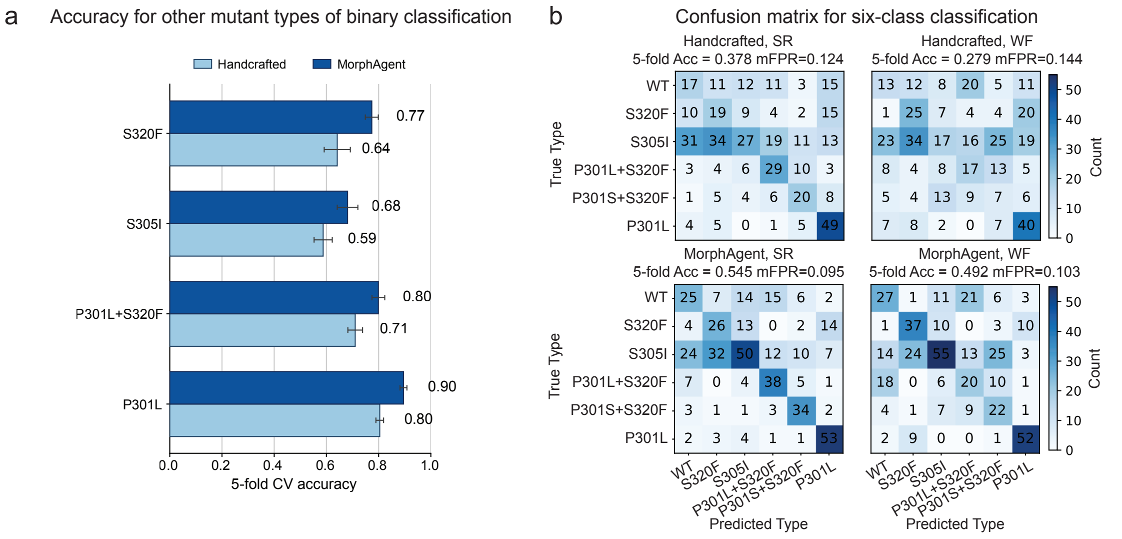


**Supplementary Fig. 11.** MorphAgent improves Tau-mutant genotype classification across binary and multiclass settings. **(a)** Five-fold cross-validation accuracy for one-versus-rest binary classification of four Tau-mutant classes not shown in the main figure (S320F, S305I, P301L+S320F and P301L). For each mutant type, MorphAgent features consistently outperform the handcrafted feature panel under super resolution imaging. Bars denote mean accuracy across folds, with error bars indicating cross-validation variability. **(b)** Confusion matrices for six-class Tau-mutant classification using handcrafted and MorphAgent feature spaces under super resolution (SR) and wide-field (WF) imaging. Rows indicate true genotypes and columns indicate predicted genotypes. MorphAgent improves overall six-class classification in both imaging modalities, with the strongest performance observed in SR images (5-fold accuracy = 0.545, macro-FPR = 0.095), compared with handcrafted SR features (accuracy = 0.378, macro-FPR = 0.124). Under WF imaging, MorphAgent also outperforms handcrafted features (accuracy = 0.492 versus 0.279). These results indicate improved genotype discrimination with MorphAgent features, with an additional performance gain under super resolution imaging.

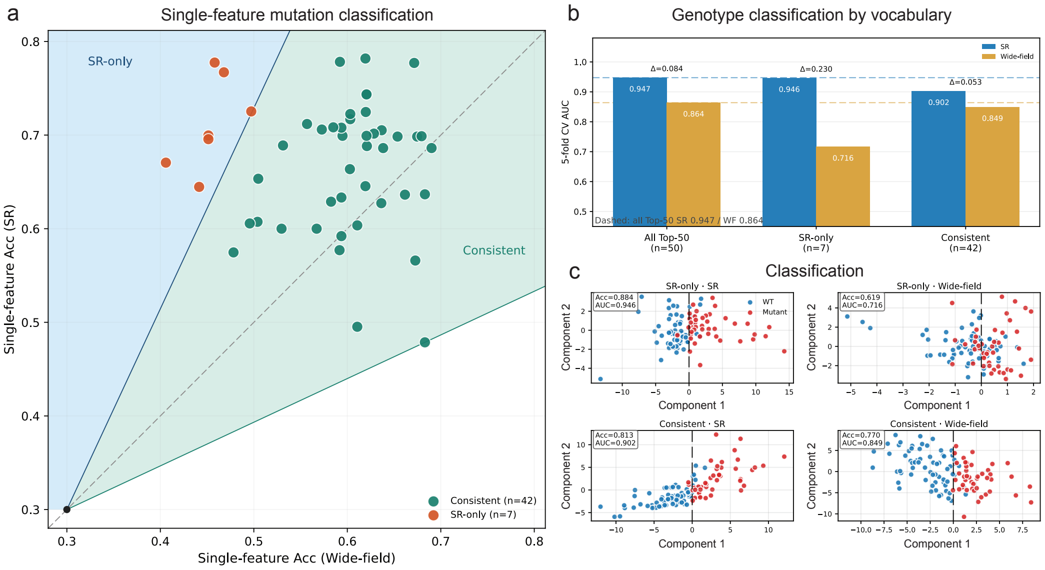


**Supplementary Fig. 12.** Super resolution specific features account for the gain in genotype discrimination. **(a)** Single feature 5-fold logistic accuracy for the Top-50 MorphAgent descriptors in paired super resolution (SR) versus wide-field images (WT versus P301S+S320F; same feature table as Fig. 4g). Two rays originating at (0.30, 0.30) and symmetric about the diagonal (y = x, ±20°) split the vocabulary into an SR-only set (upper-left sector, n = 7) and a consistent set (wedge between the rays, n = 42); features below the lower ray were excluded. **(b)** Multivariate 5-fold CV AUC using the same class-balanced logistic protocol as Fig. 4d. The full Top-50 panel recovered the original gap (AUC 0.947 versus 0.864; Δ = 0.084). The SR-only set retained nearly the same SR performance (AUC 0.946) but collapsed in wide-field images (AUC 0.716; Δ = 0.230). The consistent set performed similarly at both resolutions (AUC 0.902 versus 0.849; Δ = 0.053). **(c)** Decision-plane projections of the two vocabularies. SR-only features separate WT and mutant cells in super resolution but mix them in wide-field, whereas consistent features separate both genotypes to a similar degree in both modalities. SR-only features: *nonbundled Tau compact aggregate fraction*, *hotspot size heterogeneity index*, *Tau mislocalization pattern score*, *nonbundled tau fraction above local otsu*, *nonbundled tau aggregation area fraction*, *Tau high intensity area fraction*, *high intensity cluster count*.

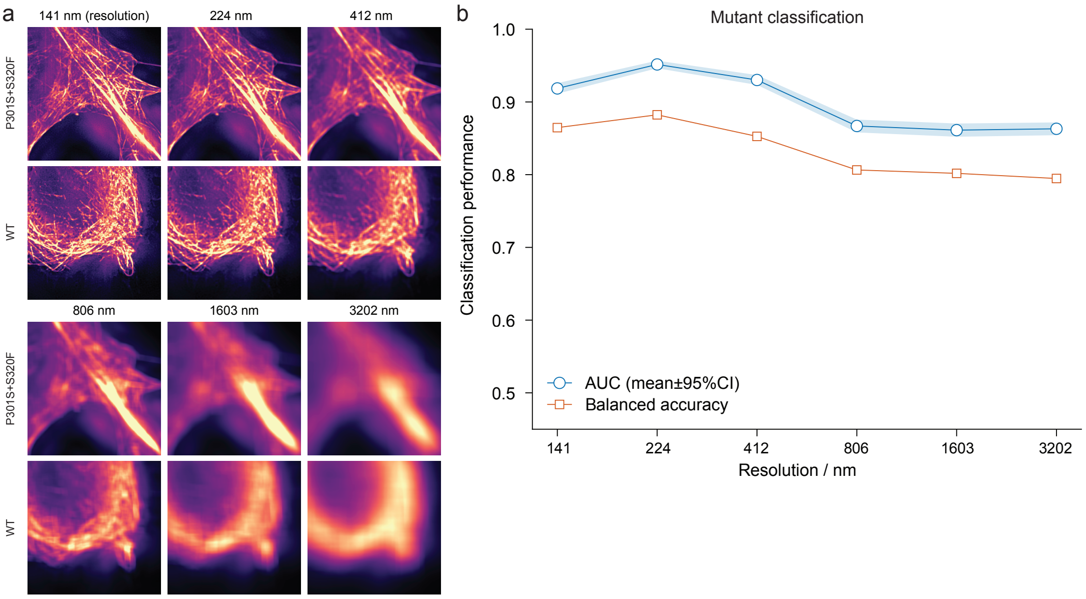


**Supplementary Fig. 13.** MorphAgent Tau morphological features progressively lose mutation-discriminative power with decreasing spatial resolution. **(a)** Representative P301S+S320F mutant (top) and wild type (bottom) cells at each resolution (141, 224, 412, 806, 1,603, 3,202 nm). **(b)** Cross-validated AUC (orange) and balanced accuracy (blue) as functions of estimated effective lateral resolution.

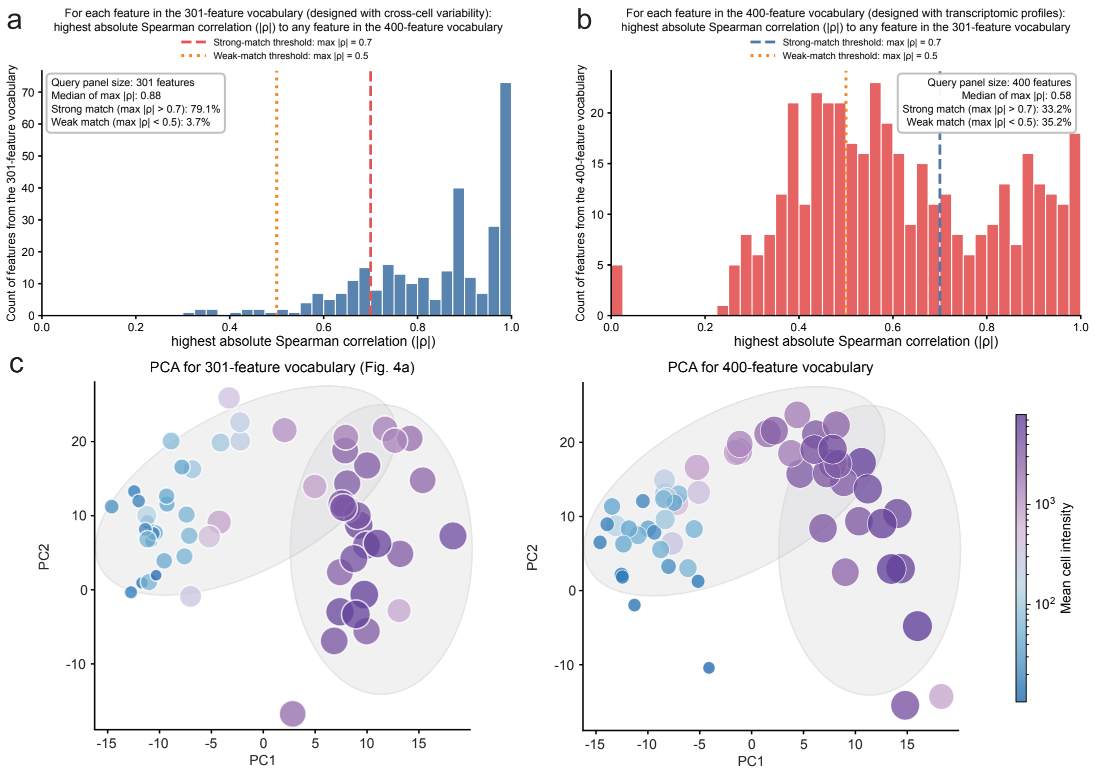


**Supplementary Fig. 14.** Cross-vocabulary correspondence and PCA comparison of the 301- and 400-feature vocabularies. Feature variation for the 301-feature vocabulary was assessed by coefficient of variation (CV) alone, whereas the 400-feature vocabulary was validated against transcriptome profiles. **(a)** Distribution of the maximum absolute Spearman correlation between each feature in the 301-feature vocabulary and its most highly correlated feature in the 400-feature vocabulary. Vertical lines mark the strong-match (max $|\rho| > 0.7$) and weak-match (max $|\rho| < 0.5$) thresholds; 79.1% of features show a strong match and 3.7% a weak match. **(b)** Reciprocal analysis from the 400-feature vocabulary to the 301-feature vocabulary; 33.2% of features show a strong match and 35.2% a weak match. **(c)** PCA of WT cells (n = 58) in the 301- and 400-feature vocabularies, colored by mean cell intensity. The two embeddings are largely consistent and both recover the same L-shaped intensity-associated trajectory.

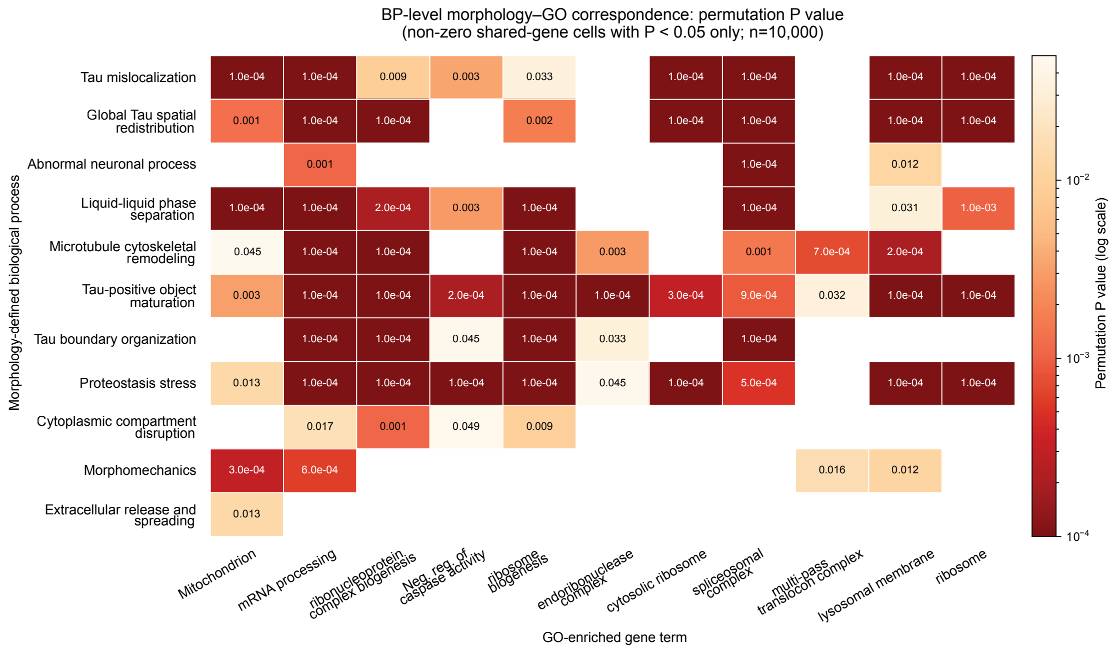


**Supplementary Fig. 15.** Permutation-test significance of the morphology–GO correspondence map (**Fig. 5d**). For each morphology-defined biological process and GO-enriched biological process, the significance of the count of shared genes was assessed with a one-sided permutation test (10,000 permutations per pair). Only pairs with a non-zero shared-gene count are shown, and blank cells indicate zero shared genes.

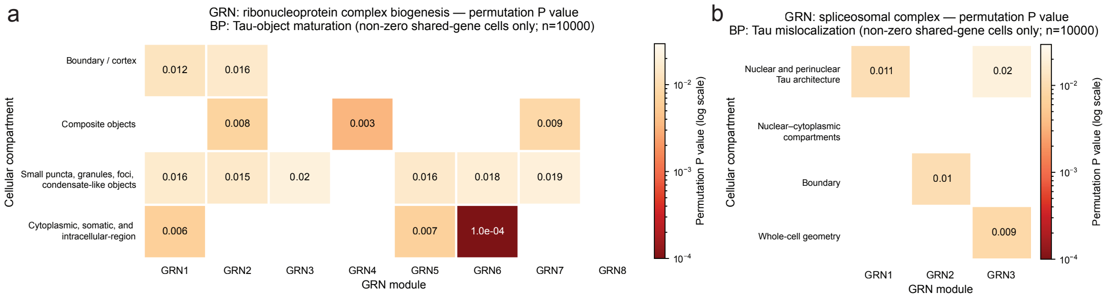


**Supplementary Fig. 16.** Permutation-test significance underlying the cellular-compartment–GRN correspondence maps (**Fig. 5e**). For each cellular compartment and GRN module pair within the ribonucleoprotein-complex-biogenesis (left) and the spliceosomal complex (right) GO terms, the significance of the count of shared genes was assessed with the same one-sided permutation test (10,000 permutations per pair). GRN1–GRN8 (left) and GRN1–GRN3 (right) follow the same left-to-right module order shown in **Fig. 5e**. Only pairs with a non-zero shared-gene count are shown (all satisfy nominal $P < 0.05$), and blank cells indicate zero shared genes.

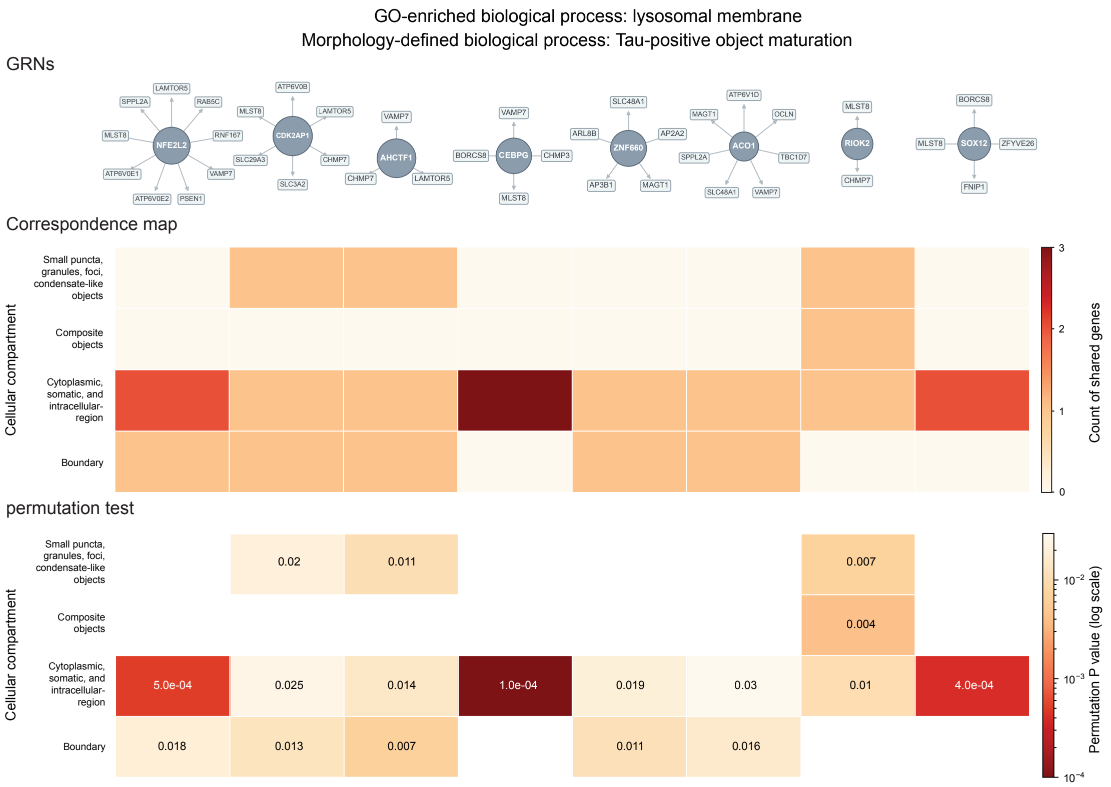


**Supplementary Fig. 17.** Top: GRN modules for lysosomal membrane. Because no lysosomal-membrane gene is a TF in the network, modules were defined from upstream TFs: among TFs whose top-decile targets include at least two of the seven genes shared by Tau-positive object maturation and lysosomal membrane, the top eight by shared-target count are shown (circle, TF; box, its lysosomal-membrane targets). Middle: the count of shared genes between each module and Tau-positive object maturation cellular compartments. Bottom: permutation *P* values for non-zero cells (10,000 size-matched draws).

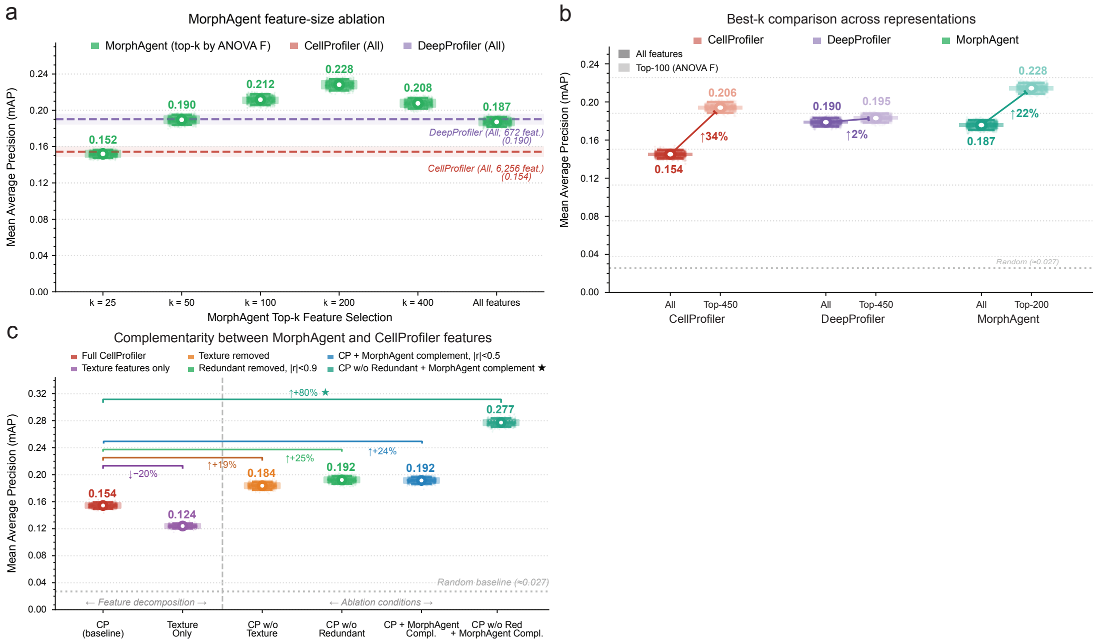


**Supplementary Fig. 18.** Feature-set ablations and cross-representation comparisons on BBBC021. **(a)** MorphAgent feature-size ablation for perturbation detection. Features were ranked by ANOVA F statistic, and performance was evaluated using the top-k selected features. MorphAgent achieved its best mAP at k = 200, outperforming both the full CellProfiler and full DeepProfiler feature spaces while using a substantially smaller feature set. **(b)** Best-k comparison across representations. Feature selection improved all three methods, but the gain was strongest for MorphAgent using only the top-200 subset (mAP = 0.228), outperforming the best-performing subsets of both CellProfiler (top-450, mAP = 0.206) and DeepProfiler (top-450, mAP = 0.195). **(c)** Complementarity between MorphAgent and CellProfiler features. Decomposing the CellProfiler space showed that texture-only features underperformed the full CellProfiler baseline, whereas removing texture or redundant features modestly improved performance. Adding MorphAgent features that were weakly correlated with CellProfiler ($|r| < 0.5$) further increased mAP, and the strongest performance was achieved when MorphAgent complements were added after redundant CellProfiler features were removed (mAP = 0.277). Together, these results indicate that MorphAgent provides a compact feature space with strong standalone performance and additional information not captured by classical handcrafted descriptors. Points indicate resampling estimates, open circles denote mean mAP, and dashed lines indicate random baseline performance where shown.

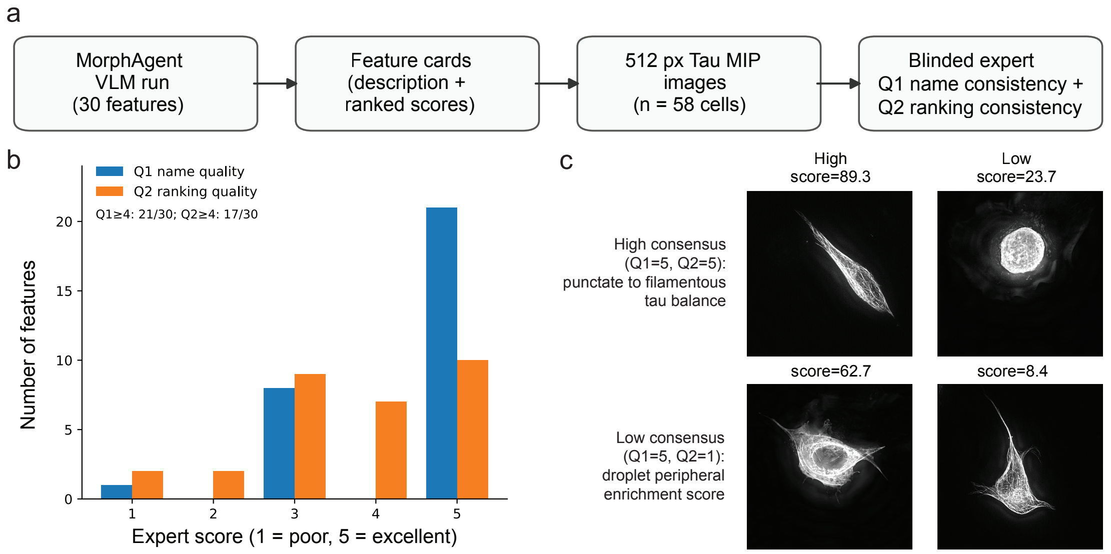


**Supplementary Fig. 19.** Blinded expert proofreading of MorphAgent VLM feature cards. Expert review workflow for automated validation of VLM-derived morphological features on Tau super resolution microscopy (n = 58 SH-SY5Y cells). **(a)** For each of the 30 retained VLM features, experts received a feature card containing the registry description, per-cell scores sorted high-to-low, and 512-pixel Tau maximum-intensity projections. Two independent criteria were scored on a 1–5 scale, with 5 being the highest score: Q1, biological specificity and interpretability of the feature name; Q2, whether the score ranking matched visually apparent phenotype extremes. **(b)** Distribution of expert scores across the 30 features (Q1≥4: 21/30; Q2≥4: 17/30). **(c)** Top row, punctate-to-filamentous Tau balance (Q1 = Q2 = 5): high- and low-scoring cells match the expected punctate-versus-filamentous phenotype. Bottom row, droplet peripheral enrichment score (Q1 = 5, Q2 = 1): despite a descriptive name and high automated validation, the expert judged that score extremes did not correspond to peripheral droplet enrichment in WT cells.

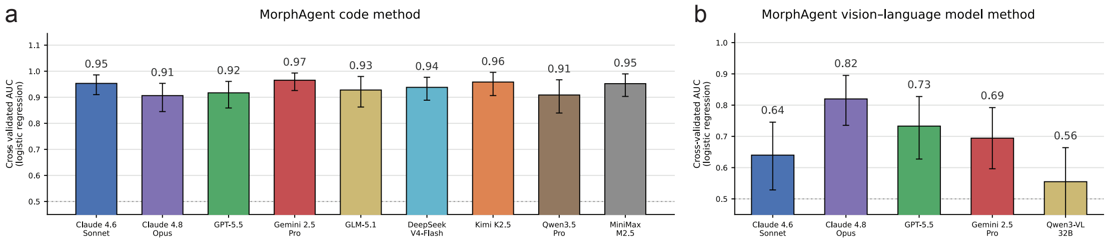


**Supplementary Fig. 20.** Cross-backbone comparison of Tau-genotype classification using MorphAgent features. For each backbone, MorphAgent independently redesigned the feature cards from the same dataset context and biological task prompt, after which the resulting features were quantified and evaluated using the same downstream classification protocol. **(a)** Area under the receiver operating characteristic curve (AUC; mean ± 95% bootstrap confidence interval) for distinguishing wild-type (WT; n=69) from P301S+S320F double-mutant (n=44) cells using logistic regression on independently generated code-based MorphAgent features. Nine backbones completed the feature-design and evaluation pipeline: Claude 4.6 Sonnet, Claude 4.8 Opus, GPT-5.5, Gemini 2.5 Pro, GLM-5.1, DeepSeek V4-Flash, Kimi K2.5, Qwen3.5 Pro and MiniMax M2.5. **(b)** As in (a), but using independently designed and VLM-scored MorphAgent features. Five backbones completed this pipeline: Claude 4.6 Sonnet, Claude 4.8 Opus, GPT-5.5, Gemini 2.5 Pro and Qwen3-VL-32B-Instruct.

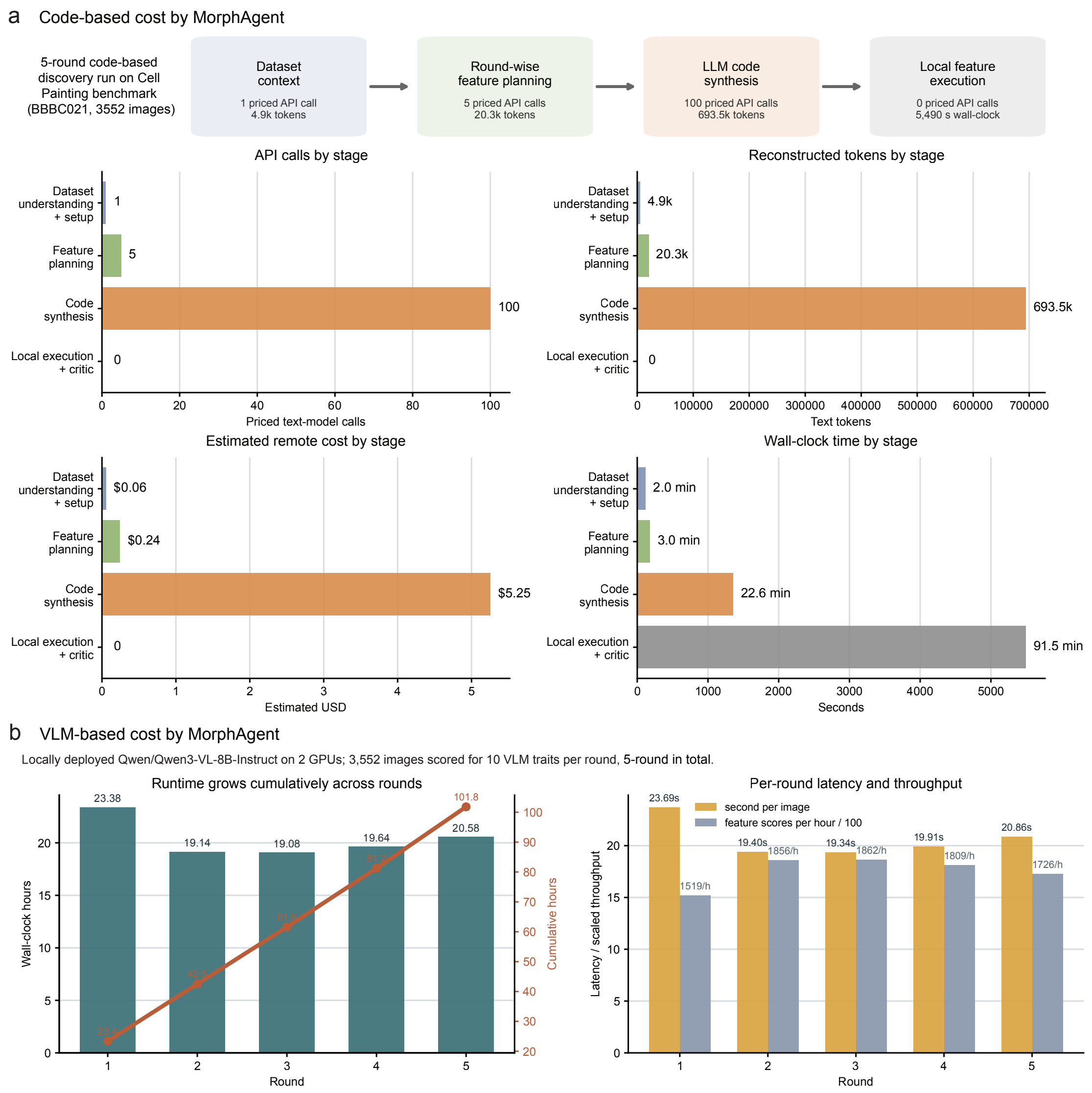


**Supplementary Fig. 21.** Computational and economic cost of MorphAgent feature discovery and VLM-based phenotype scoring. **(a)** Resource use for a representative five-round code-based MorphAgent run on 3,552 BBBC021 images. The workflow was divided into dataset setup, round-wise feature planning, LLM-based code synthesis and local feature execution. The run contained 106 priced text-model calls and 718,748 tokens (620,702 input and 98,046 output), corresponding to an estimated API cost of approximately US$5.55 under the stated pricing schedule. Code synthesis accounted for most remote-model costs, whereas local execution contributed substantially to wall-clock time. **(b)** Runtime and throughput of locally deployed Qwen3-VL-8B-Instruct on two NVIDIA GeForce RTX 4090 GPUs. Scoring 3,552 images for ten traits per round over five rounds produced 177,600 feature scores in 101.8 h. Per-round latency was approximately 19–24 s per image, corresponding to roughly 1,500–1,900 feature scores per hour.

**Algorithm 1.** Iterative feature construction pipeline of MorphAgent. Pseudocode describing dataset initialization, knowledge aggregation, segmentation precomputation, multi-round feature planning, execution through code-based or VLM-based pathways, automated validation, and iterative feedback with optional expert intervention. Candidate features are proposed in rounds, instantiated as scalar readouts, evaluated, and fed back into subsequent iterations to refine the final phenomic feature space.

**Prompt 1.** Prompt templates and interfaces used in MorphAgent. This compendium summarizes the prompt structures used for expert knowledge extraction, deep research summarization, literature aggregation, iterative feature planning, code synthesis and repair, VLM-based semantic scoring, validator generation, and expert intervention. Together, these templates define the interface through which MorphAgent integrates dataset context, biological knowledge, and iterative feedback during feature construction.

**Supplementary Table 1.** Summary of biological imaging datasets analyzed in this study. Characteristics of the six experimental datasets utilized for benchmarking, phenotypic discovery, and cross-modality inference are summarized. The table details the specific role of each dataset (e.g., perturbation profiling or mutant discrimination), sample sizes, microscopy modalities, and the availability of paired transcriptomic profiles.

**Supplementary Table 2.** Expert-curated 16-feature reference panel. Expert-curated image features derived from SIM super resolution imaging data for quantifying Tau localization, microtubule association and condensate formation. For each image-derived feature, the operational definition is paired with the subcellular structure or physiological process represented by the measurement.

**Feature Cards:** Three PDF files (**Supplementary feature list 1, Supplementary feature list 2, Supplementary feature list 3**) provide the feature name, quantification pathway (code or VLM) and biological description of each feature used in the analyses. **Supplementary feature list 4** records the assignment of each Tau feature to its morphology-defined biological process and cellular compartments. **Supplementary feature lists 5, 6** record the nuclear-associated features in MorphAgent and CellPorfiler.
