## Supplementary material for "Biologically grounded cell profiling across microscopy modalities": Algorithm

**Algorithm 1:** MorphAgent Pipeline (Part I: Setup, Initialization & Planning)

**Inputs** : user_query; data_root (dataset directory or project directory containing `dataset/`); description_path (optional); results_dir (optional); metadata_path (optional; for statistical validation); enable_expert_knowledge, enable_deep_research, enable_rag; paddlex_device (optional); method $\in$ {`code`, `vlm`, `both`}; temperature (LLM planning temperature); multigpu (optional; for VLM parallelization)

**Hyperparameters:** num_rounds $R$; features_per_iteration $K$; target_feature_count $T$ (used as planning context); illumination_correction_enabled and its parameters; segmentation backend and parameters (Cellpose-SAM); code_max_cycles (ReAct), code_timeout, code_error_rate_threshold; VLM model, max_images, resize_max

**Outputs** : $\mathbf{X}_{\text{all}}$ (all extracted features, persisted as `features.csv`); $F_{\text{all}}$ (names of all extracted features; columns of `features.csv` excluding `sample_id`); $F_{\text{HQ}}$ (validated subset; extracted from `round_r/feature_analysis_summary.json` when `metadata_path` is provided)

`// 0. Resolve paths and initialize run directory`

Resolve `data_root`; if `data_root/dataset` exists then set dataset directory $\leftarrow$ `data_root/dataset`

Create `results_dir` if not provided: `dataset_parent/results/run_{timestamp}`

Locate dataset description file: use `description_path` if provided else search common names (e.g., `dataset_index.txt`, `README.md`)

`// 1. Enumerate samples`

$\mathcal{D} \leftarrow$ list all non-hidden subdirectories under dataset directory (sorted)

Assert $|\mathcal{D}| > 0$

`// 2. Dataset understanding (LLM-assisted)`

Generate a structured dataset description by combining: (i) directory structure scan; (ii) raw description text (if available)

Store dataset description text in `results_dir/dataset_description.txt` (or equivalent)

`// 3. Knowledge aggregation (optional, cached to results)`

**if** *enable_expert_knowledge* **then**

  Extract and summarize expert materials from `project_root/expert_knowledge/`; save to `expert_knowledge_summary.txt`

**if** *enable_deep_research* **then**

  Extract text from PDFs in `project_root/deep_research/` using PaddleX; summarize; save to `deep_research_summary.txt`

**if** *enable_rag* **then**

  Extract text from `project_root/RAG/` (PDF/XML), batch by token budget; summarize with MoA/profiling focus; save to `rag_knowledge_summary.txt`

`// 4. Global segmentation precomputation (optional but enabled by default)`

Infer segmentation channels from dataset description (heuristic; may be overridden by feature requirements)

**foreach** $s \in \mathcal{D}$ **do**

  Select primary image file(s) for $s$; run Cellpose-SAM segmentation if not cached

  Save masks to `s/segmentation/{cyto,nuclei,cytoplasm}.tif`

Write segmentation summary (including mask ordering) to `results_dir/segmentation_summary.json`

`// 5. Initialize multi-round bookkeeping`

Initialize `features.csv` with column `sample_id` and $N$ rows (one per $s \in \mathcal{D}$)

Set $F_{\text{all}} \leftarrow \emptyset$, $F_{\text{HQ}} \leftarrow \emptyset$

Set `previous_analysis_summary` $\leftarrow$ `None`

**Algorithm 2:** MorphAgent Pipeline (Part II: Execution)

```
// 6. Multi-round feature propose-and-verify (Start)
for $r \leftarrow 1$ to $R$ do
    // 6.1 Prepare planning context (avoid duplicates; leverage prior analysis)
    Load existing feature names from features.csv into $F_{\text{all}}$
    Build planning context including: dataset description, knowledge summaries, $F_{\text{all}}$, and
      previous_analysis_summary
    Restrict available methods based on method argument
    // 6.2 Feature planning (LLM returns strict JSON list)
    Run planning graph on a representative sample (typically the first sample) to obtain feature plan
      $\mathcal{P}_r = \{f_{r,j}\}_{j=1}^{K}$
    Save round_r/feature_plan.json
    // 6.3 Preprocess secondary slices for VLM (first round only)
    if $r = 1$ then
        foreach $s \in \mathcal{D}$ do
            Ensure s/slices/ exists; if missing, generate PNG slices from primary TIFF
            if illumination_correction_enabled then
                Compute ICF (global or channel-wise) and apply during slice generation
            Write s/slices/channel_mapping.json when the data is multi-channel 2D
    // 6.4 Execute VLM features (batch per sample)
    Let $\mathcal{V}_r \leftarrow \{f \in \mathcal{P}_r : f.\texttt{method} = \texttt{vlm}\}$
    if $|\mathcal{V}_r| > 0$ then
        foreach $s \in \mathcal{D}$ (optionally in parallel across GPUs) do
            Select VLM input images for $s$: prefer secondary slices/ else fall back to primary files
            if $\exists f \in \mathcal{V}_r$ requiring segmentation then
                Load segmentation masks if available; otherwise run per-sample segmentation
            Call VLM once to score all features in $\mathcal{V}_r$; obtain $\{(f, \hat{y}_{s,f})\}$
            Log prompt, image list, and model outputs to round_r/features/vlm_batch/...
        Append VLM feature columns to features.csv (preserving sample order)
    // 6.5 Execute code features (ReAct code synthesis + batch execution)
    Let $\mathcal{C}_r \leftarrow \{f \in \mathcal{P}_r : f.\texttt{method} = \texttt{code}\}$
    foreach $f \in \mathcal{C}_r$ do
        if $f$ requires segmentation then
            Ensure per-sample segmentation masks exist; record per-sample image/mask relative
              paths
        // Generate code + execute with retries
        Initialize cycle counter $c \leftarrow 0$
        repeat
            Generate Python function extract(img, *segmentation_masks) using LLM prompts
              with dataset + knowledge context
            Execute extract on all samples using primary image file selection; collect values and
              errors
            if error rate $\leq$ code_error_rate_threshold then
                break
            Use LLM-based fixer to propose (i) dependency installation steps and (ii) guidance; apply
              and regenerate
            $c \leftarrow c + 1$
        until $c$ = code_max_cycles
        Append code feature column to features.csv
        Save round_r/features/{feature_name}/extract.py and logs
    // Loop continues in Part III...
```

**Algorithm 3:** MorphAgent Pipeline (Part III: Validation)

`// ... Continuation of Loop` $r$

`// 6.6 Statistical feature validation (optional; drives` $F_{\mathrm{HQ}}$ `and next-round planning)`

**if** *metadata_path is provided* **then**

  Run automated analysis: LLM generates an analysis protocol (JSON) and executable code

  Compute per-feature variability metrics and cross-modality alignment metrics using metadata

  Write `round_r/feature_analysis.csv`, `feature_analysis.json`, `feature_analysis_summary.json`, and `feature_analysis_report.md`

  Update `previous_analysis_summary` with: high-variance features, stable features, high-alignment features, and recommendations

  Update $F_{\mathrm{HQ}}$ according to criteria in `feature_analysis_summary.json`

Save `round_r/round_results.json`

`// End of Loop` $r$

`// 7. Final outputs`

Let $\mathbf{X}_{\mathrm{all}} \leftarrow$ values in `features.csv` excluding `sample_id`

Output $\mathbf{X}_{\mathrm{all}}$, $F_{\mathrm{all}}$ (column names), and $F_{\mathrm{HQ}}$ (validated subset)
