## Supplementary material for "Biologically grounded cell profiling across microscopy modalities": Prompt

### Prompts for MorphAgent (Templates and Interfaces)

#### Knowledge acquisition and summarization prompts

**Overview.** MorphAgent aggregates three complementary knowledge sources—**expert knowledge**, **deep research**, and **RAG literature**—and converts each into a structured textual summary emphasizing *visual appearance descriptions* that can guide segmentation, code-based feature implementation, and VLM scoring.

##### (i) Expert knowledge extraction and consolidation

**Automation.** The system scans a project-level `expert_knowledge/` folder for supported files (text, images, PDFs), summarizes each item, and then synthesizes a unified summary.

###### Prompt.

```
You are an expert knowledge extraction assistant. Your task is to extract key information from
    expert-provided documents to guide image feature extraction, segmentation, and feature
    generation.

Please extract:
1) Important image features and morphological indicators (priorities; what to focus on vs ignore)
    .
2) Appearance descriptions of important features (CRITICAL):
   - shape, size, intensity, texture, spatial distribution
   - how to recognize them in images
   - guidance for segmentation, coding, and VLM scoring
3) Biological background knowledge and related concepts
4) Recommendations and pitfalls for feature extraction

Output should be clear, structured, and particularly detailed in describing appearances.
```

###### Inputs and filled variables.

- `file_path.name`: filename for provenance.
- `text` or inferred image info (when the expert artifact is an image/PDF).

**Output contract.** Free-form structured English text with explicit appearance descriptions and prioritization, saved as `expert_knowledge_summary.txt`.

##### (ii) Deep research summarization (automated literature reading)

**Automation (supplementary).** In the manuscript setting, deep research is run as a fully automated literature-reading stage: (a) retrieve candidate papers (see RAG stage below), (b) parse PDFs using Paddle-based document understanding, and (c) run an automated "deep research" agent to synthesize findings into feature-oriented guidance. We provide an implementation using the DeepResearch module in `/data3/yez/MorphAgent_github/deep_research_module`.

**Prompt.**

```
You are an expert scientific literature analyst. Your task is to extract key information from
    Deep Research PDFs to guide image feature extraction.

Please extract:
1) Important findings and conclusions relevant to image-derived phenotypes
2) Important image features / morphological indicators
3) Appearance descriptions of important features (CRITICAL):
   - what the feature looks like in images
   - visual morphology, spatial distribution, texture, intensity patterns
   - how to recognize it (color, shape, size, location, texture)
   - how it should guide segmentation, coding, and VLM scoring
4) Methods and technical details relevant to imaging/analysis
5) Recommendations and pitfalls for feature extraction

Output should be clear, structured, and particularly detailed in describing appearances.
```

**Inputs and filled variables.**

- A batch of PDF texts: `{pdf_name, pdf_text}` pairs (text may be truncated to meet token budgets).
- Optional device spec for document parsing (e.g., `gpu:0`).

**Output contract.** A unified deep-research summary in English, saved as `deep_research_summary.txt`, emphasizing feature appearances and operational guidance.

##### (iii) RAG literature aggregation (MoA + profiling focus)

**Automation (supplementary).** For pharmacological datasets (e.g., BBBC021), MorphAgent can automatically build a literature corpus by downloading open-access PMC XML and/or PDFs for each compound (or mechanism) and then summarizing these documents with explicit focus on MoA and morphological profiling. We provide an automated acquisition pipeline based on `Build-PMC-OA` (download + conversion).

**Prompt.**

```
You are a scientific literature analysis expert focusing on (1) drug mechanisms of action (MoA)
    and (2) morphological profiling.

Your task: extract key information directly related to:
1) MoA: molecular targets, pathways, inhibition/activation modes, cellular process-level
    mechanisms
2) Morphological profiling: Cell Painting/high-content profiling, CellProfiler features,
    phenotypes, profile matching, MoA-oriented matching
```

```
Very Important:
- Only extract information directly related to MoA and morphological profiling
- For each piece of information, label evidence strength (Strong/Moderate/Weak)
- Describe morphological features in detail (visual appearance, spatial distribution, etc.)
- These descriptions will guide segmentation, coding, and VLM scoring

Output should be clear and structured.
```

**Inputs and filled variables.**

- `document_texts`: parsed text from PDFs and/or PMC XML files.
- `document_names` and `document_types` for provenance.
- Automatic batching based on token budget (documents are split into batches; batch summaries are merged by a second-stage prompt).

**Output contract.** A unified RAG knowledge summary in English, saved as `rag_knowledge_summary.txt`, explicitly structured around MoA and profiling, with evidence grading.

#### Feature planning prompt (iterative proposal)

**Purpose.** In each iteration, MorphAgent uses an LLM to propose a small set of new scalar features tailored to the dataset format and the scientific query. The prompt enforces (i) strict JSON output, (ii) explicit method choice (`code` vs `vlm`), (iii) segmentation requirement flags, and (iv) non-duplication with previously extracted features.

##### Inputs (filled variables)

**Dataset and imaging context.**

- `{dataset_description}`: LLM-generated dataset description (dimension type, channel mapping, file organization).
- `{channel_information}`: expanded channel mapping (for multi-channel) and image ordering explanation.
- `{dataset_image_format}`: extracted/augmented format notes (2D multi-channel or 3D z-stack).
- `{cell_context_info}`: single-cell vs multi-cell context inferred from the user query.

**Task context and constraints.**

- `{user_query}`: user-defined scientific objective.
- `{target_feature_count}`, `{features_per_iteration}`: target size and per-round budget.
- `{available_methods}` and `{method_instructions}`: method restriction (`code`/`vlm`/both).

**Iteration memory (critical for non-duplication and refinement).**

- `{previous_analysis_guidance}`: distilled guidance from prior round quality analysis (high-variance features, stable features, high-alignment features, recommendations).
- `{all_existing_feature_names}` (embedded inside guidance): the complete set of previously extracted feature names (must avoid duplicates).

**Knowledge sources (optional).**

- `{deep_research_info}`: deep-research summary emphasizing visual appearances.
- `{rag_knowledge_info}`: RAG summary emphasizing MoA and profiling.
- `{expert_knowledge_info}`: expert summary emphasizing prioritized traits and appearances.

**Prompt (excerpted header + output contract).** The full template is long; below we include the high-signal structure and the strict output contract used by the system.

```
You are an expert in computational image analysis and feature engineering.
Your task is to design scalar image features for the given dataset.

1. Dataset Information
{dataset_description}
{channel_information}

2. User Query and Task Context
User Query: {user_query}
{cell_context_info}

2.5. Previous Round Analysis Results (Multi-Round Execution)
{previous_analysis_guidance}

3. Knowledge-Guided Feature Design
{deep_research_info}
{rag_knowledge_info}
{expert_knowledge_info}

4. Requirements for New Features
Design {features_per_iteration} new scalar features...

5. Segmentation Requirement
For each proposed feature, determine whether segmentation is needed...

6. Method Choice (code vs vlm)
Available methods: {available_methods}
... choose method in {"code","vlm"} and justify ...

7. Output Format (strict JSON)
Return only a JSON array:
[
  {
    "name": "short_feature_name_in_snake_case",
    "description": "...",
    "category": "morphology"|"intensity"|"texture"|"distribution"|"spatial"|"other",
    "needs_segmentation": true|false,
    "segmentation_prompt": "..."|null,
```

```
    "method": "code"|"vlm",
    "method_rationale": "..."
  }
]

CRITICAL NAMING REQUIREMENT:
- if method == "vlm", name MUST start with "vlm_"
- if method == "code", name MUST NOT start with "vlm_"
```

##### Expected output

**Output contract.** The LLM must return a syntactically valid JSON array with exactly the specified keys. Each entry is a feature specification used downstream to route execution:

- `method=code`: triggers code synthesis and batch execution on *primary* image files.
- `method=vlm`: triggers VLM scoring on *secondary* slices (preferred) with fallback to primary files.
- `needs_segmentation=true`: triggers segmentation checks and provides masks to downstream executors.

#### Coding agent prompts and ReAct-style recovery

**Purpose.** For features assigned to `method=code`, MorphAgent synthesizes Python code implementing a single scalar-valued function `extract(img, *segmentation_masks)`. The agent is constrained to operate on *primary* image files (original data), optionally using segmentation masks. Execution is instrumented with an error-driven repair loop (ReAct-style) that combines (i) code planning, (ii) code generation, (iii) batch execution, and (iv) error reflection that proposes environment fixes and high-level guidance for regeneration.

##### (i) Code planning prompt (optional chain-of-thought plan)

**Inputs (filled variables).**

- Feature spec: `{feature_name}`, `{feature_description}`, `{feature_category}`
- Image statistics: `{image_shape}`, `{image_dtype}`, `{image_min}`, `{image_max}`
- Segmentation interface: `{segmentation_files_info}`, `{segmentation_mask_order}`, `{segmentation_statistics}`
- Channel information: `{channel_information}`

**Prompt.**

```
You are an expert Python engineer for scientific image analysis.
Create a detailed implementation plan.

Feature name: {feature_name}
Feature description: {feature_description}
Feature category: {feature_category}

```

```
Input interface:
- img: numpy array, shape={image_shape}, dtype={image_dtype}, range=[{image_min},{image_max}]
- segmentation masks (optional):
{segmentation_files_info}
{segmentation_mask_order}
{segmentation_statistics}
```

**Output contract.** A natural-language plan (not executed directly) that is appended into the code-generation prompt as an "Implementation Plan" section.

##### (ii) Code generation prompt (strict function interface)

**Critical constraint.** The code must implement exactly one function `extract(img, *segmentation_masks)` and *must not* perform file I/O or orchestration. All imports must be inside the function.

**Key filled variables (non-exhaustive but high-signal).**

- Dataset context: `{dataset_description}`, `{dataset_image_format}`, `{channel_information}`, `{cell_context_info}`
- Feature spec: `{feature_name}`, `{feature_description}`, `{feature_category}`
- Execution interface: `{image_shape}`, `{image_dtype}`, `{image_min}`, `{image_max}`
- Segmentation interface: `{segmentation_files_info}`, `{segmentation_mask_order}`, `{segmentation_statistics}`
- Per-sample paths (when segmentation required): `{sample_paths_info}` (relative image/-mask paths for multiple samples)
- Knowledge sources: `{deep_research_info_code}`, `{rag_knowledge_info_code}`, `{expert_knowledge_info_code}`
- Fix guidance (only after failures): `{feature_guidance_message}`

**Prompt (header + core rules).**

```
You are a senior Python engineer for scientific image analysis.
You MUST write a SINGLE function:
  def extract(img, *segmentation_masks):
that computes a scalar feature from an image.

CRITICAL: The img parameter is loaded from PRIMARY FILES only.
Do NOT use slices/ or other derived data for code-based features.

Function Input Interface:
1) img: numpy array (shape={image_shape}, dtype={image_dtype}, range min={image_min}, max={
    image_max})
2) segmentation masks (optional, auto-provided):
{segmentation_files_info}
{segmentation_mask_order}
{segmentation_statistics}

Allowed libraries:
```

```
- numpy, scipy.ndimage
- optionally skimage.measure / skimage.filters for morphology
- optionally cv2 (limited API set)

Important rules:
- NO file I/O, NO plotting, NO external executables
- ALL imports MUST be inside extract()
- Handle dimensionality according to dataset description; if unexpected, return 0.0
- Always return float(...)
```

**Expected output contract.** Return *only* the function definition (no markdown, no extra text). The orchestrator saves it as `round_r/features/<feature>/extract.py` and executes it on all samples.

#### (iii) Reflection / fix prompt (ReAct-style recovery)

**Purpose.** When batch execution fails, the system summarizes representative errors and asks a fixer prompt to propose (a) an installation script for missing dependencies and (b) high-level guidance to improve robustness. The guidance is injected into the next code-generation attempt.

**Prompt.**

```
You are a helper agent that improves environment readiness and communicates high-level guidance
    to the code generation LLM.

Feature: {feature_name} (method={method})
Description: {feature_description}
Existing code:
{existing_code}

Error summary:
{error_summary}

CRITICAL: Environment Context
The code runs in a specific conda environment (typically 'micro_v4_test').
When generating install_script, you may:
1) Provide plain pip/conda commands, or
2) Explicitly wrap with: conda run -n micro_v4_test ...
```

**Expected output contract.** The fixer returns a JSON object with:

- `install_script`: shell commands to install missing packages (optional).
- `guidance_message`: concise guidance injected into the next generation attempt.

#### (iv) ReAct loop (control flow summary)

At runtime, the orchestrator repeats: generate → execute on all samples → if failure rate is high, reflect/fix → regenerate, up to a maximum number of cycles (default: 3).

#### VLM prompts (single-feature and batch scoring)

**Purpose.** For features assigned to `method=vlm`, MorphAgent uses a vision-language model (VLM) to score semantically rich morphological traits. The VLM prompt explicitly encodes dataset format, channel mapping, image ordering, and knowledge summaries, and enforces a strict output contract: free-text analysis followed by a final-line JSON score.

##### Inputs (filled variables)

**Dataset and image context.**

- `{dataset_description}`: includes explicit data organization rules (primary vs secondary).
- `{dataset_image_format}`: dimension notes (2D multi-channel vs 3D).
- `{channel_information}`: channel mapping and biological meaning.
- `{image_list_description}`: ordered list of provided images with per-slice semantics (critical for 2D multi-channel and 3D z-stacks).

**Knowledge sources.**

- `{deep_research_info}`, `{rag_knowledge_info}`, `{expert_knowledge_info}`

**Feature specification.**

- Single-feature: `{feature_name}`, `{feature_description}`, `{feature_category}`
- Batch: `{features_list_text}`, `{num_features}`

##### Single-feature VLM scoring prompt.

```
You are an expert in image analysis and computational microscopy.

Dataset Information
{dataset_description}

Data Path Selection (CRITICAL)
- Primary files (original data): direct files in the sample directory
- Secondary files (derived data): files in subdirectories like slices/
- For VLM features: prefer secondary (e.g., slices/*.png); if not available, may use primary
{dataset_image_format}
{channel_information}
{image_list_description}

Knowledge Sources for Feature Guidance
{deep_research_info}{rag_knowledge_info}{expert_knowledge_info}

TASK
Evaluate ONE feature:
- Name: {feature_name}
- Description: {feature_description}
- Category: {feature_category}

CONTINUOUS SCORING REQUIREMENTS
```

```
- Output a continuous score in [0,100] with fine-grained values (avoid anchoring).

OUTPUT FORMAT
(1) Detailed natural-language analysis (with spatial references: slices, channels, regions)
(2) FINAL line: pure JSON
{"score": <float>}
```

##### Batch VLM scoring prompt.

```
TASK
Evaluate MULTIPLE features simultaneously (total: {num_features}):
{features_list_text}

OUTPUT FORMAT
(1) Detailed natural-language analysis for EACH feature
(2) FINAL line: pure JSON mapping feature_name -> score
{"feature_name_1": <float>, ..., "feature_name_N": <float>}
```

##### Expected output contract

The final line must be valid JSON with numeric scores in [0,100]. The system parses this JSON to populate `features.csv`. Any deviation from the contract is treated as failure and may trigger retry logic.

#### Iteration memory (ReAct) and metadata-aware refinement

**Purpose.** MorphAgent is designed as an iterative system: each round proposes new features, executes them on all samples, and (when metadata is available) performs feature-quality analysis to construct a compact "memory" that guides the next round. This mechanism prevents duplicate features, encourages exploration around high-performing features, and ties feature discovery to external metadata (e.g., treatment labels, MoA, dose, or transcriptomic embeddings).

##### (i) What is stored as iteration memory?

At the beginning of round $r > 1$, the planner receives a structured object `previous_analysis_summary` that may include:

- `all_existing_feature_names`: complete list of previously extracted feature names from `features.csv` (hard constraint: avoid duplicates).
- `previous_features_summary`: a compact summary from the previous round (feature names, counts, and analysis highlights).
- `cumulative_analysis`: optional aggregated summaries across rounds.

##### (ii) How metadata is used (when provided)

When `metadata_path` is provided, MorphAgent runs an automated analysis module after each round:

- It evaluates *feature variability* (e.g., variance, standard deviation, coefficient of variation).

- It evaluates *cross-modality alignment* between each feature and metadata fields, selecting appropriate tests based on field type (categorical vs continuous vs embeddings).

The output is summarized into a JSON memory object (saved as `feature_analysis_summary.json`) and a human-readable report (saved as `feature_analysis_report.md`).

##### (iii) Memory injection into the next planning prompt (ReAct-style guidance)

The planning prompt includes a dedicated section that is automatically populated with `previous_analysis_guida`[illegible] The injected guidance is structured as:

- **High-performing features**: high-variance features and/or features with strong metadata alignment.
- **Low-performing features**: stable/low-variance features to avoid replicating.
- **Strong recommendation**: generate *related* features forming a "feature family" around successful concepts, while ensuring non-duplication of names.

**Injected text.**

```
High-Performing Features:
  ... list of feature names ...
Critical Guidance:
  - Generate RELATED new features based on these successful features.
Low-Performing Features:
  ... list of feature names ...
All Previously Extracted Features (MUST avoid duplicate feature names):
  ... exhaustive feature list ...
```

##### (iv) Outputs of the iteration mechanism

The iteration mechanism produces two outputs that are used in the manuscript:

- **All extracted features** $F_{all}$ and the corresponding matrix $\mathbf{X}_{all}$ (persisted as `features.csv`).
- **Validated features** $F_{HQ}$: extracted from `feature_analysis_summary.json` (e.g., high-variance set, high-alignment set). The exact criteria can be reported as thresholds in the Supplement (e.g., top-$m$ by variance; correlation $> \tau$).

**Note.** The memory mechanism is intentionally lightweight: it summarizes the empirical evidence from prior rounds into a compact guidance block, rather than carrying forward full raw statistics. This design keeps prompts short enough for stable generation while preserving the key signals needed for iterative refinement.

#### Prompts for feature-quality analysis (protocol, code, report)

**Purpose.** When a metadata table is provided, MorphAgent runs an automated feature-quality analysis after each round. This module is itself LLM-driven: it (i) plans an analysis protocol conditioned on dataset description and metadata schema, (ii) generates executable Python analysis code from the protocol, and (iii) generates a structured summary and narrative report that feeds back into the next feature-planning iteration.

#### (i) Analysis protocol planning prompt (STRICT JSON)

**Inputs.**

- `description_text`: dataset description (includes metadata semantics).
- `metadata_columns`: list of metadata fields (names).
- `metadata_sample`: one example row (types/ranges).
- `num_features`: number of features currently in `features.csv`.

**Prompt.**

```
You are an expert in computational biology and feature analysis.
Design a comprehensive analysis protocol for evaluating image features.

You MUST return a STRICT JSON protocol.
For each feature, evaluate TWO aspects:
1) Feature variability (variance, std, coefficient of variation, group-wise variance)
2) Cross-modality alignment with metadata (correlation, separability, ANOVA, effect size)

Output JSON structure:
{
  "variability_metrics": [...],
  "cross_modality_metrics": [...],
  "metadata_field_classification": {
    "categorical_labels": [...],
    "continuous_embeddings": [...],
    "batch_info": [...],
    "biological_perturbations": [...]
  },
  "analysis_workflow": [...]
}
Return ONLY valid JSON.
```

**Output contract.** A valid JSON object saved as `feature_analysis_protocol.json`. The protocol must be uniform for all features and computable with numpy/pandas/scipy only.

#### (ii) Analysis code generation prompt (Python-only)

**Inputs.** The protocol JSON (above).

**Prompt.**

```
You are an expert Python programmer specializing in scientific data analysis.
Generate a complete, executable function:
  def run_feature_analysis(features, feature_names, metadata_df):
that:
- processes ALL features,
- computes all protocol metrics,
- handles missing values and edge cases,
- uses only numpy, pandas, scipy,
- returns {"results": DataFrame, "summary": dict}.
Return ONLY the Python code.
```

**Output contract.** Plain Python code saved as `feature_analysis_code.py` and executed by the analysis executor.

##### (iii) Report generation prompt (JSON summary + markdown report)

**Inputs.** A compact summary of the analysis outputs (metric names, number of features, summary statistics) plus the dataset description.

**Prompt.**

````
You must generate TWO outputs:
1) JSON Summary:
{
  "high_variance_features": [...],
  "stable_features": [...],
  "high_correlation_features": [
    {"feature": "...", "metadata_field": "...", "correlation": 0.85, "metric_type": "..."}
  ],
  "recommendations": [...]
}
2) Markdown Report:
- Executive summary
- Variability analysis
- Cross-modality alignment analysis
- Key findings and recommendations

Return in the format:
```json
{JSON_SUMMARY}
```
```markdown
{MARKDOWN_REPORT}
```
````

**Output contract.** The module writes:

- `feature_analysis_summary.json` (machine-readable memory for the next iteration)
- `feature_analysis_report.md` (human-readable narrative)
