## Supplementary feature list 1 for "Biologically grounded cell profiling across microscopy modalities"

### MorphAgent BBBC021 feature catalog

467 MorphAgent features for BBBC021 Cell Painting (MCF7). Method: vlm or code.

| # | Feature name | Method | Description |
|---|---|---|---|
| 1 | **tubulin intensity total** | **code** | Computes the total integrated intensity of the Tubulin channel across the entire image. This serves as a proxy for total microtubule polymer mass, which is directly affected by taxanes (stabilizers) and vinca alkaloids (destabilizers). |
| 2 | **actin fiber alignment index** | **code** | Estimates the alignment of actin filaments by calculating the anisotropy of the structure tensor in the Actin channel. High alignment suggests stress fiber formation, while low alignment suggests disorganized cytoskeleton. |
| 3 | **cell roundness mean** | **code** | Calculates the average roundness (4 * pi * area / perimeter^2) of segmented cell bodies based on the Actin/Tubulin signal. Rounding is a key feature of mitotic arrest and detachment, while flattening indicates senescence. |
| 4 | **dapi texture entropy** | **code** | Measures the entropy of the pixel intensity distribution within the nucleus (DAPI channel). High entropy corresponds to heterogeneous chromatin texture (e.g., condensation, fragmentation), which is relevant for detecting DNA damage or apoptosis. |
| 5 | **tubulin actin overlap fraction** | **code** | Calculates the fraction of Tubulin signal that spatially overlaps with Actin signal (using thresholded masks). This quantifies the extent of cytoskeletal interplay and cell spreading. |
| 6 | **cell confluence ratio** | **code** | Calculates the ratio of the image area covered by cells (Actin/Tubulin mask) to the total image area. This measures cell growth inhibition or cytotoxicity. |
| 7 | **dapi intensity std** | **code** | Calculates the standard deviation of pixel intensities within segmented nuclei. High variability suggests chromatin condensation (bright spots) mixed with darker regions, typical of apoptotic or mitotic cells. |
| 8 | **actin radial distribution** | **code** | Measures the distribution of Actin intensity relative to the cell center (e.g., ratio of cortical actin to internal actin). Changes in this distribution reflect cytoskeletal reorganization events like rounding or spreading. |
| 9 | **tubulin texture contrast** | **code** | Computes the Haralick Contrast feature for the Tubulin channel. This measures the local intensity variation, helping to distinguish between smooth, diffuse tubulin staining and sharp, fibrous microtubule networks. |
| 10 | **cytoplasm nucleus area ratio** | **code** | Calculates the average ratio of cytoplasmic area (Cell area - Nuclear area) to nuclear area. This N:C ratio is a fundamental cellular parameter that changes during differentiation, senescence (increased ratio), and cancer progression. |
| 11 | **nucleus area mean** | **code** | Calculates the mean area (in pixels) of segmented nuclei using the DAPI channel. Changes in nuclear size can indicate cell cycle arrest, swelling, or pyknosis (shrinkage) associated with apoptosis. |
| 12 | **dapi texture haralick contrast mean** | **code** | Calculates the mean Haralick contrast of the DAPI channel within nuclear boundaries. This quantifies chromatin condensation and texture heterogeneity, which are key markers of DNA damage and early apoptosis. |
| 13 | **micronuclei count mean** | **code** | Counts the number of small, distinct DAPI-positive objects located within the cytoplasm but outside the main nucleus, averaged per cell. This is a direct indicator of genomic instability and chromosomal damage (e.g., from Vinca alkaloids). |
| 14 | **tubulin intensity skewness mean** | **code** | Computes the skewness of the pixel intensity distribution in the Tubulin channel for each cell. High skewness can indicate the presence of bright, bundled microtubules (e.g., Taxol effect) versus a more diffuse distribution. |
| 15 | **tubulin fiber alignment index** | **code** | Measures the anisotropy or directional coherence of structures in the Tubulin channel using structure tensors. High alignment indicates bundling or stress response, distinguishing between different microtubule poisons. |
| 16 | **tubulin radial concentration ratio** | **code** | Calculates the ratio of Tubulin intensity in the perinuclear region versus the cell periphery. This helps identify phenotypes like monoastral spindles (central concentration) versus spread interphase networks. |
| 17 | **actin cell area mean** | **code** | Measures the mean area of the entire cell defined by the Actin cytoskeleton. This captures gross morphological changes such as cell shrinking (apoptosis) or enlargement/flattening (senescence). |
| 18 | **actin eccentricity mean** | **code** | Measures the mean eccentricity (0=circle, 1=line) of the cell shape defined by Actin. This quantifies cell elongation, relevant for detecting migratory phenotypes or stress responses. |
| 19 | **actin edge intensity ratio** | **code** | Calculates the ratio of Actin intensity at the cell boundary (cortex) relative to the cell center. This can identify cortical actin ring formation or loss of stress fibers. |
| 20 | **nucleus cytoplasm area ratio** | **code** | Calculates the ratio of nuclear area (DAPI) to cytoplasmic area (Actin/Tubulin). The N:C ratio is a classic cytological metric that changes during differentiation, cancer progression, and drug response. |
| 21 | **cell neighbor count mean** | **code** | Calculates the mean number of neighboring cells within a fixed radius for each cell. This quantifies cell density and clustering, which can be affected by contact inhibition or cytotoxic cell loss. |
| 22 | **nucleus count** | **code** | Counts the total number of nuclei detected in the DAPI channel. This serves as a primary proxy for cell proliferation or cytotoxicity (cell loss). |
| 23 | **tubulin fiber gradient magnitude mean** | **code** | Computes the mean gradient magnitude of the Tubulin (green) channel within cell boundaries. High values indicate sharp, distinct microtubule fibers (bundling), while low values suggest diffuse or depolymerized tubulin. |
| 24 | **actin boundary roughness mean** | **code** | Measures the roughness of the cell boundary (perimeter squared / (4 * pi * area)) derived from the Actin channel. High roughness can indicate membrane blebbing or invasive protrusions (invadopodia). |
| 25 | **tubulin intensity mean** | **code** | Measures the average pixel intensity in the Tubulin (green) channel across the entire image. Changes in overall tubulin intensity can indicate microtubule stabilization (e.g., Taxol) or destabilization (e.g., Vinca alkaloids). |

| | | | |
|---|---|---|---|
| 26 | **actin intensity cv** | **code** | Calculates the coefficient of variation (std/mean) of pixel intensities in the Actin (red) channel. High variance may indicate stress fiber formation or cytoskeletal disruption, contrasting with diffuse staining. |
| 27 | **dapi tubulin correlation** | **code** | Computes the Pearson correlation coefficient between the DAPI (nucleus) and Tubulin (microtubule) channels pixel-wise. Changes in this correlation can indicate gross morphological rearrangements or co-localization shifts during mitosis. |
| 28 | **cell count** | **code** | Counts the total number of nuclei detected in the DAPI channel. A reduction in cell count is a direct proxy for cytotoxicity or cytostasis. |
| 29 | **nuclear eccentricity std** | **code** | Computes the standard deviation of nuclear eccentricity across the image. A high standard deviation suggests a heterogeneous population containing both round (interphase) and elongated (mitotic or deformed) nuclei. |
| 30 | **dapi intensity kurtosis** | **code** | Calculates the kurtosis of the pixel intensity distribution in the DAPI channel. High kurtosis indicates a distribution with heavy tails or peaks, potentially reflecting bright condensed chromatin foci amidst a darker background. |
| 31 | **cell skeleton branch points** | **code** | Skeletons the Actin channel mask and counts the number of branch points. This serves as a proxy for the complexity of the actin cytoskeletal network. |
| 32 | **tubulin radial distribution** | **code** | Measures the radial intensity profile of Tubulin relative to the centroid of the nearest nucleus. This quantifies whether microtubules are concentrated perinuclearly or extended to the cell periphery. |
| 33 | **actin tubulin intensity ratio** | **code** | Computes the ratio of total Actin intensity to total Tubulin intensity. Shifts in this ratio can indicate specific disruption of one cytoskeletal component relative to the other. |
| 34 | **tubulin texture entropy** | **code** | Computes the entropy of the texture in the Tubulin channel. This feature captures the complexity of the microtubule network; low entropy might suggest depolymerization (diffuse signal) while high entropy could indicate bundling or stabilization (e.g., Taxol effect). |
| 35 | **actin tubulin correlation** | **code** | Measures the Pearson correlation coefficient between pixel intensities of the Actin (red) and Tubulin (green) channels. This quantifies the spatial overlap and co-localization of the cytoskeleton components, which can be disrupted by drugs affecting cell shape. |
| 36 | **nuclear actin ratio** | **code** | Calculates the ratio of total integrated intensity in the DAPI channel to the Actin channel. This feature provides a gross measure of the nuclear-to-cytoplasmic ratio, often altered in cell cycle arrest or hypertrophy. |
| 37 | **cell area mean** | **code** | Calculates the mean total cell area using the Actin channel to define the cell boundary. This helps identify phenotypes like senescence (enlarged cells) or cytotoxic shrinkage. |
| 38 | **dapi contrast haralick** | **code** | Computes the Haralick Contrast texture feature for the DAPI channel. This measures the local intensity variation, capturing chromatin texture patterns associated with different cell cycle states or drug effects. |
| 39 | **tubulin intensity kurtosis** | **code** | Computes the kurtosis of the pixel intensity distribution in the Tubulin channel. High kurtosis indicates a distribution with heavy tails, suggesting the presence of very bright structures (e.g., bundles) against a dark background. |
| 40 | **dapi tubulin overlap fraction** | **code** | Measures the fraction of total Tubulin intensity that spatially overlaps with the segmented nuclear region. This can indicate microtubule encroachment into the nuclear area or nuclear envelope breakdown. |
| 41 | **nuclear roundness mean** | **code** | Measures the mean roundness of nuclei. Deviations from circularity can indicate apoptosis (fragmentation/blebbing) or mitotic catastrophe. |
| 42 | **dapi intensity entropy** | **code** | Measures the Shannon entropy of pixel intensities within segmented nuclei. High entropy indicates heterogeneous chromatin texture (e.g., condensation or punctate foci), while low entropy suggests uniform staining. |
| 43 | **tubulin fiber texture contrast** | **code** | Measures the Haralick contrast in the Tubulin channel. High contrast often correlates with microtubule bundling (thick, rigid fibers) induced by Taxanes, as opposed to a fine meshwork. |
| 44 | **cell eccentricity mean** | **code** | Calculates the mean eccentricity of segmented cells (based on Actin/cytoskeleton). This quantifies cell elongation, distinguishing between rounded (mitotic/dead) and elongated (mesenchymal/migratory) phenotypes. |
| 45 | **micronuclei count per cell** | **code** | Counts small, detached DAPI-positive objects near the main nucleus. Micronuclei are a specific indicator of genomic instability and mitotic slippage (e.g., Vinca alkaloids). |
| 46 | **tubulin radial distribution ratio** | **code** | Calculates the ratio of Tubulin intensity in the perinuclear region versus the cell periphery. This spatial feature helps identify phenotypes like monoastral spindles (central concentration) versus normal spreading. |
| 47 | **nuclear cytoplasmic ratio** | **code** | Computes the ratio of the total nuclear area (DAPI) to the total cytoplasmic area (Actin - DAPI). This is a fundamental cellular metric often altered in cancer progression and drug response. |
| 48 | **dapi solidity mean** | **code** | Measures the solidity (area / convex hull area) of nuclei. Low solidity indicates irregular boundaries, invaginations, or blebbing associated with nuclear stress or apoptosis. |
| 49 | **tubulin intensity variance** | **code** | Calculates the variance of pixel intensities in the Tubulin channel. High variance suggests distinct structures (bundles/fibers), while low variance suggests diffuse depolymerization. |
| 50 | **actin fiber orientation coherence** | **code** | Measures the global coherence of actin fiber orientation using structure tensor analysis. High coherence suggests aligned stress fibers, while low coherence indicates a disorganized cytoskeleton. |
| 51 | **dapi granulometry mean** | **code** | Performs a granulometric analysis (opening operations with increasing structuring element sizes) on the DAPI channel to estimate the average size of bright chromatin speckles. This quantifies chromatin condensation patterns. |
| 52 | **micronuclei count approx** | **code** | Counts the number of small, distinct DAPI-positive objects that are detached from the main nuclei. An increase in micronuclei is a hallmark of genomic instability and mitotic slippage. |
| 53 | **actin cortical intensity ratio** | **code** | Calculates the ratio of Actin intensity at the cell boundary (cortex) versus the cell interior. This distinguishes cells with strong cortical actin rings from those with diffuse or stress-fiber dominant actin. |

| | | | |
|---|---|---|---|
| 54 | **dapi haralick contrast mean** | **code** | Computes the mean Haralick contrast texture feature for the DAPI channel within segmented nuclei. High contrast may indicate chromatin condensation or punctate nuclear bodies. |
| 55 | **tubulin intensity mass mean** | **code** | Calculates the mean total integrated intensity of the Tubulin channel per cell. This proxies the total abundance of polymerized microtubules, which can be affected by stabilizing (Taxanes) or destabilizing (Vinca alkaloids) drugs. |
| 56 | **tubulin skeleton length mean** | **code** | Skeletonizes the Tubulin signal within cells and measures the total length of the skeleton. This quantifies the complexity and density of the microtubule network. |
| 57 | **actin compactness mean** | **code** | Computes the compactness (perimeter squared divided by area) of the cell body. This feature captures complex shape changes, such as the formation of protrusions, invadopodia, or membrane ruffling. |
| 58 | **nucleus to cell area ratio mean** | **code** | Calculates the mean ratio of nuclear area (DAPI) to total cell area (Actin) for each cell. This N/C ratio is a fundamental cytological metric that changes during differentiation, cancer progression, and drug response. |
| 59 | **tubulin actin correlation mean** | **code** | Calculates the Pearson correlation coefficient between pixel intensities of Tubulin and Actin channels within each cell. This measures the spatial colocalization and structural integrity of the cytoskeleton components. |
| 60 | **cell neighbor distance mean** | **code** | Calculates the mean distance to the nearest neighbor for each cell. This captures spatial patterns like clustering (colonies) vs scattering, which can be affected by drugs influencing cell motility or contact inhibition. |
| 61 | **tubulin radial intensity gradient mean** | **code** | Measures the average gradient of tubulin intensity radiating from the nucleus center to the cell periphery. This feature helps identify phenotypes like monoastral spindles (centralized tubulin) versus normal interphase networks. |
| 62 | **actin edge intensity ratio mean** | **code** | Calculates the ratio of Actin intensity at the cell boundary (cortex) versus the cell interior. This distinguishes cells with strong cortical actin rings from those with diffuse or stress-fiber-rich actin. |
| 63 | **dapi peak intensity count mean** | **code** | Detects and counts local intensity maxima (peaks) within each segmented nucleus in the DAPI channel. This estimates the number of heterochromatin foci or nucleoli, which can change under stress. |
| 64 | **tubulin haralick contrast global** | **code** | Calculates the global Haralick contrast texture feature for the Tubulin channel. This measures the local intensity variation, helping to distinguish between diffuse microtubules and distinct, high-contrast bundles. |
| 65 | **tubulin intensity kurtosis global** | **code** | Computes the kurtosis of the pixel intensity distribution for the entire Tubulin channel. High kurtosis indicates a distribution with heavy tails or peaks, potentially corresponding to bright microtubule bundles against a dark background. |
| 66 | **dapi tubulin overlap fraction mean** | **code** | Measures the fraction of the Tubulin signal that spatially overlaps with the nuclear mask. High overlap may indicate mitotic spindles (which form around chromosomes) or cytoskeletal collapse onto the nucleus. |
| 67 | **cell count total** | **code** | Counts the total number of nuclei in the image. This is a primary readout for cytotoxicity (cell death) or proliferation inhibition. |
| 68 | **actin texture entropy mean** | **code** | Computes the mean entropy of the Actin texture within segmented cells. High entropy indicates a complex, disordered cytoskeletal network, while low entropy may suggest diffuse or uniform staining. |
| 69 | **tubulin moment of inertia mean** | **code** | Calculates the mean moment of inertia of the Tubulin intensity relative to the cell centroid. This measures how dispersed the microtubule network is from the cell center. |
| 70 | **dapi high intensity area fraction mean** | **code** | Calculates the fraction of the nuclear area that exceeds a high intensity threshold (e.g., >90th percentile of image). This quantifies the extent of bright, condensed chromatin regions. |
| 71 | **cell perimeter area ratio mean** | **code** | Computes the mean ratio of perimeter squared to area for segmented cells (Actin). This is a measure of boundary complexity or compactness, sensitive to cell spreading and membrane ruffling. |
| 72 | **tubulin intensity skewness** | **code** | Measures the skewness of the pixel intensity histogram in the Tubulin channel. A highly skewed distribution may indicate bright, sparse structures (like bundles or spindles) against a dark background. |
| 73 | **dapi tubulin distance mean** | **code** | Calculates the average Euclidean distance between the centroid of the nucleus and the weighted centroid of the tubulin signal within a cell. This captures cell polarity and the symmetry of the microtubule network. |
| 74 | **dapi peak intensity ratio** | **code** | Calculates the ratio of the maximum pixel intensity to the mean pixel intensity within the nucleus. A high ratio suggests the presence of bright nuclear foci (e.g., condensed chromatin or micronuclei). |
| 75 | **tubulin radial distribution slope** | **code** | Calculates the gradient of Tubulin intensity radiating from the nuclear centroid to the cell periphery. This helps distinguish phenotypes where microtubules are clustered perinuclearly (e.g., collapsed) versus extended to the edge. |
| 76 | **dapi boundary intensity mean** | **code** | Measures the mean intensity of DAPI signal at the nuclear boundary (rim). A bright rim can indicate marginalization of chromatin, a specific feature of certain apoptotic or necrotic processes. |
| 77 | **actin texture contrast mean** | **code** | Calculates the Haralick Contrast texture feature for the Actin channel. High contrast indicates sharp transitions typical of distinct stress fibers, while low contrast may indicate diffuse actin staining. |
| 78 | **tubulin polarity index** | **code** | Measures the spatial distribution of tubulin intensity relative to the nucleus centroid. It calculates the vector sum of intensity-weighted positions in the Tubulin channel centered on the nucleus, quantifying the degree of polarization of the microtubule network, relevant for migration or spindle formation. |
| 79 | **actin fiber alignment** | **code** | Analyzes the local orientation of actin filaments in the Actin channel using structure tensors or gradient histograms. A high score indicates strongly aligned stress fibers (common in static, adherent cells), while a low score indicates a disorganized meshwork (common in motile or rounded cells). |
| 80 | **nuclear circularity median** | **code** | Calculates the median circularity (4*pi*area/perimeter^2) of nuclei. Deviations from circularity can indicate nuclear blebbing, fragmentation (apoptosis), or irregular shapes associated with nuclear envelope stress. |

| | | | |
|---|---|---|---|
| 81 | **actin cortical enrichment** | **code** | Measures the ratio of Actin intensity at the cell boundary (cortex) versus the cell interior. Changes in cortical actin are relevant for cell motility and adhesion phenotypes. |
| 82 | **nuclear actin exclusion** | **code** | Quantifies the average intensity of Actin signal within the nuclear region relative to the cytoplasmic region. In healthy cells, actin is largely excluded from the nucleus; an increase might indicate nuclear envelope rupture or segmentation errors. |
| 83 | **dapi intensity integrated** | **code** | Sums the total DAPI intensity across all nuclei. This is a proxy for total DNA content in the field of view, which correlates with cell number and ploidy levels. |
| 84 | **actin puncta prominence** | **vlm** | Visually scores how prominent high-intensity actin puncta appear within cells (e.g., focal adhesions or actin aggregates). Higher values indicate more conspicuous punctate actin structures against the surrounding cytoskeleton. |
| 85 | **tubulin texture haralick contrast** | **code** | Computes the Haralick Contrast texture feature on the Tubulin (green) channel. This quantifies the local intensity variations, which can capture the difference between smooth, intact microtubule networks and disrupted, bundled, or fragmented microtubules caused by taxanes or vinca alkaloids. |
| 86 | **nucleus cytoplasm intensity ratio actin** | **code** | Calculates the ratio of mean Actin intensity inside the nucleus vs. the cytoplasm. While Actin is primarily cytoplasmic, changes in this ratio can indicate segmentation issues or specific nuclear actin involvement. |
| 87 | **dapi granularity spectrum mean** | **code** | Computes the mean power spectrum intensity at high frequencies for the DAPI channel. This acts as a proxy for nuclear granularity/heterochromatin condensation without explicit texture feature selection. |
| 88 | **cytoskeleton radial distribution tubulin** | **code** | Measures the radial intensity profile of Tubulin from the nucleus center outwards. This quantifies whether microtubules are concentrated perinuclearly or extend to the cell periphery. |
| 89 | **cell orientation coherence** | **code** | Measures the standard deviation of the orientation angles of the major axes of all cells in the image. Low deviation implies cells are aligned in a similar direction (e.g., due to flow or contact guidance), while high deviation implies random orientation. |
| 90 | **tubulin branching density** | **code** | Applies a skeletonization algorithm to the Tubulin channel and counts the number of branch points per unit area. This quantifies the complexity of the microtubule network. |
| 91 | **nuclear shape circularity mean** | **code** | Computes the mean circularity (4*pi*area/perimeter^2) of nuclei. This metric helps distinguish between healthy, round nuclei and irregular, blebbed, or fragmented nuclei often seen in apoptosis or drug-induced stress. |
| 92 | **tubulin intensity mean per cell** | **code** | Calculates the mean intensity of the Tubulin channel within segmented cell boundaries. This quantifies the overall abundance of microtubules, which can be affected by stabilizing or destabilizing drugs. |
| 93 | **actin cell eccentricity mean** | **code** | Measures the mean eccentricity of the cell boundary defined by Actin staining. This quantifies cell elongation, differentiating between flattened, mesenchymal-like morphologies and rounded, amoeboid or mitotic phenotypes. |
| 94 | **nucleus cytoplasm area ratio mean** | **code** | Calculates the mean ratio of nuclear area to cytoplasmic area (N:C ratio) for each cell. This is a classic cytological metric that changes during differentiation, cell cycle arrest, and neoplastic transformation. |
| 95 | **dapi tubulin pixel correlation** | **code** | Computes the Pearson correlation coefficient between DAPI and Tubulin pixel intensities across the whole image (or per cell). This measures the spatial overlap of DNA and microtubules, which is relevant for detecting mitotic spindles. |
| 96 | **actin texture gabor energy mean** | **code** | Applies a bank of Gabor filters to the Actin (red) channel and calculates the mean energy. This captures the prevalence of linear structures like stress fibers at various orientations. |
| 97 | **nuclear boundary solidity mean** | **code** | Calculates the mean solidity (area / convex hull area) of nuclei. Low solidity indicates irregular, lobulated, or fragmented nuclear boundaries, often associated with nuclear envelope stress or apoptosis. |
| 98 | **cell dispersion index** | **code** | Calculates the nearest-neighbor distance statistics for cell centroids across the image. This quantifies spatial clustering or dispersion, reflecting cell density and potential contact inhibition or loss thereof. |
| 99 | **tubulin radial intensity gradient** | **code** | Measures the rate of change of tubulin intensity radiating from the nuclear centroid. This quantifies the spatial distribution of the microtubule network (perinuclear accumulation vs. peripheral extension). |
| 100 | **tubulin intensity mass displacement** | **code** | Calculates the distance between the geometric centroid of the cell and the intensity-weighted centroid of the Tubulin channel. Large displacements indicate asymmetric microtubule organization, relevant for polarization or defects in spindle formation. |
| 101 | **actin texture gabor energy** | **code** | Computes the energy response of Gabor filters applied to the Actin channel. This quantifies the presence and strength of linear structures like stress fibers, distinguishing them from diffuse cortical actin. |
| 102 | **cell compactness mean** | **code** | Measures the mean compactness (perimeter squared divided by area) of cells defined by Actin. This shape factor differentiates between spread-out, irregular cells and compact, round cells (e.g., mitotic or rounded). |
| 103 | **nucleus cell centroid distance** | **code** | Measures the Euclidean distance between the centroid of the nucleus and the centroid of the whole cell. This quantifies cell polarity and symmetry; loss of polarity can indicate cytoskeletal disruption. |
| 104 | **tubulin radial distribution mean** | **code** | Measures the average distance of tubulin signal intensity from the nuclear centroid. This helps distinguish between perinuclear clustering (e.g., collapsed cytoskeleton) and peripheral distribution. |
| 105 | **nucleus intensity entropy** | **code** | Measures the Shannon entropy of pixel intensities within segmented nuclei in the DAPI channel. This quantifies the texture complexity of chromatin; high entropy may indicate fragmented or condensed chromatin associated with DNA damage or apoptosis. |
| 106 | **cell circularity mean** | **code** | Calculates the mean circularity (4 * pi * area / perimeter^2) of the entire cell boundary defined by Actin. This distinguishes between spread-out, mesenchymal-like cells and rounded, potentially detached or mitotic cells. |
| 107 | **tubulin total intensity mean** | **code** | Measures the average total integrated intensity of the Tubulin channel per cell. This quantifies the abundance of microtubules, which can be significantly altered by microtubule stabilizers (e.g., Taxol) or destabilizers. |

| | | | |
|---|---|---|---|
| 108 | **actin fiber alignment coherence** | **code** | Analyzes the local orientation of actin filaments to determine a coherence score. High coherence indicates parallel stress fibers, while low coherence suggests a disorganized meshwork, relevant for detecting cytoskeletal reorganization. |
| 109 | **tubulin texture correlation** | **code** | Computes the Haralick correlation texture feature for the Tubulin channel. This measures the linear dependency of gray levels on neighboring pixels, capturing the structural regularity of the microtubule network (e.g., bundles vs. diffuse). |
| 110 | **cell extent mean** | **code** | Calculates the mean extent (ratio of object area to bounding box area) of segmented cells. This shape descriptor helps differentiate between spread-out, irregular cells (low extent) and compact, geometric shapes (high extent). |
| 111 | **actin tubulin overlap pearson** | **code** | Computes the Pearson correlation coefficient between pixel intensities of the Actin and Tubulin channels within cell masks. This measures the spatial colocalization of the two cytoskeletal networks, which may decouple under specific drug treatments. |
| 112 | **nucleus boundary roughness** | **code** | Measures the roughness of the nuclear boundary (perimeter^2 / (4 * pi * area) normalized or similar fractal dimension proxy). Irregular boundaries can indicate nuclear envelope stress or early fragmentation. |
| 113 | **tubulin radial profile slope** | **code** | Calculates the slope of the intensity profile of Tubulin radiating from the nucleus center to the cell periphery. A steep negative slope indicates perinuclear concentration (e.g., monoastral spindles), while a flat slope indicates diffuse distribution. |
| 114 | **actin intensity skewness** | **code** | Measures the skewness of the pixel intensity distribution in the Actin channel. High skewness suggests the presence of sparse, bright structures (like stress fibers or foci) against a darker background. |
| 115 | **tubulin intensity mean cell** | **code** | Measures the mean intensity of the Tubulin (green) channel within the cell boundary. This quantifies the overall abundance of microtubules, which can be affected by stabilizers (e.g., Taxol) or destabilizers. |
| 116 | **actin area mean** | **code** | Calculates the mean area of the cell cytoplasm defined by the Actin channel. This captures cell spreading versus rounding, a key morphological response to cytoskeletal toxins or mitotic arrest. |
| 117 | **tubulin distribution asymmetry** | **code** | Measures the displacement between the weighted center of mass of the Tubulin signal and the geometric centroid of the cell. High asymmetry can indicate polarized microtubule organization or formation of localized structures like nuclear caps. |
| 118 | **actin edge sharpness mean** | **code** | Calculates the mean gradient magnitude at the boundary of the segmented cell in the Actin channel. Sharp edges may indicate tense, spread cells, while diffuse edges might suggest membrane blebbing or retraction. |
| 119 | **nucleus actin correlation** | **code** | Computes the Pearson correlation coefficient between pixel intensities of DAPI and Actin channels within the cell area. This measures the spatial overlap of the nucleus and cytoskeleton, which can change during mitosis or cell spreading. |
| 120 | **tubulin texture contrast mean** | **code** | Calculates the mean Haralick contrast of the Tubulin channel within segmented cell boundaries. High contrast correlates with distinct microtubule bundling (e.g., Taxol effect), while low contrast indicates diffuse tubulin. |
| 121 | **dapi haralick entropy mean** | **code** | Calculates the mean Haralick Entropy of the DAPI channel within nuclei. Entropy measures the randomness of the texture; high entropy suggests complex, non-uniform chromatin structure. |
| 122 | **tubulin skeleton branch length mean** | **code** | Skeletons the tubulin network and calculates the mean length of branches. This quantifies the continuity of the microtubule network, which is disrupted by destabilizers. |
| 123 | **cell neighbor adjacency count** | **code** | Calculates the average number of immediate neighbors for each cell. This measures local cell crowding, which can influence morphology and drug response. |
| 124 | **nucleus compactness mean** | **code** | Computes the mean compactness (perimeter squared divided by area) of nuclei. This shape factor helps distinguish between smooth, round nuclei (healthy/interphase) and irregular or lobulated nuclei (apoptotic/mitotic catastrophe). |
| 125 | **nucleus cytoplasm ratio mean** | **code** | Calculates the mean ratio of nuclear area to cytoplasmic area for each cell. This is a classic cytological metric for cell health and differentiation. |
| 126 | **actin intensity mean** | **code** | Measures the mean average intensity of the Actin (red) channel within segmented cells. High intensity may indicate stress fiber formation or accumulation, while low intensity suggests depolymerization. |
| 127 | **tubulin haralick contrast mean** | **code** | Computes the mean Haralick Contrast of the Tubulin channel within cells. High contrast corresponds to sharp, distinct fibers (bundling), while low contrast indicates diffuse or depolymerized tubulin. |
| 128 | **actin haralick correlation mean** | **code** | Computes the mean Haralick Correlation of the Actin channel within cells. Measures the linear dependency of gray levels, reflecting the organized, repetitive nature of stress fibers versus disordered actin. |
| 129 | **dapi tubulin pearson correlation** | **code** | Computes the Pearson correlation coefficient between pixel intensities of the DAPI and Tubulin channels globally. This measures the spatial overlap or exclusion of nuclei and microtubules. |
| 130 | **nucleus displacement mean** | **code** | Measures the average distance between the centroid of the nucleus and the centroid of the cell body. Large displacements can indicate polarization, motility, or specific phenotypes like 'nuclear caps'. |
| 131 | **cell solidity mean** | **code** | Computes the mean solidity (area / convex hull area) of the cell boundary. This feature detects surface irregularities like membrane blebbing (low solidity) versus smooth, spread-out morphologies (high solidity), indicative of apoptotic or necrotic processes. |
| 132 | **tubulin radial distribution gradient** | **code** | Analyzes the radial intensity profile of Tubulin from the nucleus center to the cell periphery. This scalar summarizes whether microtubules are concentrated perinuclearly (e.g., collapsed) or extend to the cell edge. |
| 133 | **tubulin texture contrast glcm** | **code** | Calculates the GLCM contrast of the Tubulin channel. High contrast correlates with microtubule bundling (thick, distinct fibers often seen with Taxanes), while low contrast suggests a diffuse or depolymerized network. |

| | | | |
|---|---|---|---|
| 134 | **nuclear fragmentation index** | **code** | Counts the number of small, bright DAPI objects relative to the total number of nuclei. This serves as a proxy for the apoptotic index, detecting nuclear fragmentation (karyorrhexis). |
| 135 | **tubulin spatial entropy** | **code** | Calculates the entropy of the spatial distribution of tubulin intensity. High entropy implies a diffuse, disordered distribution (depolymerization), while low entropy implies organized structures (spindles or bundles). |
| 136 | **nuclear intensity cv mean** | **code** | Computes the mean Coefficient of Variation (std/mean) of DAPI intensity within each nucleus. High variation correlates with chromatin condensation or punctate patterns like gamma-H2AX foci indicative of DNA damage. |
| 137 | **cell form factor mean** | **code** | Calculates the mean form factor (4 * pi * Area / Perimeter^2) of segmented cells. Values close to 1 indicate perfect circles (mitotic rounding), while lower values indicate elongated or irregular shapes (mesenchymal phenotype or stress fibers). |
| 138 | **tubulin texture energy mean** | **code** | Calculates the mean Haralick Energy (angular second moment) of the Tubulin channel within segmented cells. High energy indicates textural uniformity, while low energy suggests complex, heterogeneous patterns like microtubule bundling or disruption. |
| 139 | **actin cortical ratio mean** | **code** | Calculates the ratio of Actin intensity at the cell periphery (cortex) versus the cell center. This feature helps distinguish cells with strong cortical rings (often rounded) from those with pervasive stress fibers (spread out). |
| 140 | **dapi tubulin correlation global** | **code** | Computes the Pearson correlation coefficient between pixel intensities of the DAPI and Tubulin channels globally. High correlation might indicate mitotic cells where tubulin (spindle) condenses around chromosomes, versus interphase where they are spatially distinct. |
| 141 | **tubulin skeleton density global** | **code** | Measures the total length of the skeletonized Tubulin network normalized by the total foreground area. This quantifies the complexity and density of the microtubule network, relevant for detecting network disruption or stabilization. |
| 142 | **actin orientation coherency global** | **code** | Measures the global coherency of Actin fiber orientation using structure tensor analysis. High coherency indicates aligned stress fibers, while low coherency suggests a disorganized actin meshwork. |
| 143 | **tubulin gradient magnitude mean** | **code** | Calculates the mean magnitude of the image gradient (e.g., Sobel filter) in the Tubulin channel. This serves as a proxy for the sharpness and definition of microtubule fibers. |
| 144 | **tubulin haralick entropy mean** | **code** | Computes the entropy of the texture in the Tubulin channel. High entropy suggests a complex, disordered cytoskeletal network, while low entropy may indicate loss of structure or diffuse staining. |
| 145 | **actin intensity mean global** | **code** | Measures the mean pixel intensity of the Actin channel across the entire image. This quantifies the overall abundance of the actin cytoskeleton. |
| 146 | **dapi texture entropy mean** | **code** | Computes the entropy of the pixel intensity distribution within the nucleus (DAPI channel). High entropy indicates heterogeneous chromatin texture (e.g., condensation), while low entropy suggests uniform staining. |
| 147 | **nuclear form factor mean** | **code** | Computes the mean form factor (4 * pi * Area / Perimeter^2) of nuclei. Values close to 1 indicate perfect circles, while lower values indicate irregular shapes, capturing phenotypes like nuclear blebbing or fragmentation described in the RAG knowledge. |
| 148 | **dapi intensity mad** | **code** | Calculates the Median Absolute Deviation (MAD) of pixel intensities within the DAPI channel across the entire image. This serves as a robust measure of texture heterogeneity in chromatin structure, potentially reflecting condensation or DNA damage responses. |
| 149 | **tubulin fiber alignment strength** | **code** | Measures the anisotropy or alignment strength of the tubulin channel using structure tensor analysis. High values indicate parallel bundling of microtubules (a Taxane phenotype), while low values indicate a disordered meshwork. |
| 150 | **tubulin skeleton network length mean** | **code** | Applies skeletonization to the tubulin channel of segmented cells and calculates the mean total length of the skeleton per cell. This quantifies the complexity and intactness of the microtubule network, which is disrupted by Vinca alkaloids. |
| 151 | **actin cell extent mean** | **code** | Calculates the mean 'extent' (ratio of object area to bounding box area) of segmented cells in the Actin channel. This helps distinguish between spread-out, irregular mesenchymal-like shapes and compact, rounded cells. |
| 152 | **actin radial intensity gradient** | **code** | Measures the gradient of actin intensity from the cell center to the periphery. This distinguishes cells with strong cortical actin rings (peripheral) from those with diffuse or stress-fiber dominant (central) actin distributions. |
| 153 | **tubulin actin correlation** | **code** | Computes the Pearson correlation coefficient between Tubulin and Actin channels. This metric captures the degree of overlap between the microtubule network and the actin cytoskeleton, which may decouple under specific drug treatments. |
| 154 | **tubulin intensity cv** | **code** | Calculates the coefficient of variation (std/mean) of pixel intensities in the Tubulin (green) channel within the cell body. This measures the heterogeneity of the microtubule network, distinguishing between diffuse tubulin (depolymerized) and bundled/structured microtubules. |
| 155 | **dapi haralick contrast** | **code** | Calculates the Haralick Contrast texture feature for the DAPI channel within segmented nuclei. This quantifies the local intensity variations, capturing subtle changes in chromatin structure that may not be reflected in simple intensity statistics. |
| 156 | **tubulin perinuclear ratio** | **code** | Calculates the ratio of Tubulin intensity in a ring immediately surrounding the nucleus versus the rest of the cell. This captures the collapse of the microtubule network around the nucleus, a phenotype seen with certain destabilizing drugs. |
| 157 | **cell area mean actin** | **code** | Calculates the mean area of segmented cells based on the Actin channel. Significant increases in cell area can indicate senescence (flattened, enlarged cells), while decreases may indicate rounding or shrinkage. |
| 158 | **actin cell eccentricity** | **code** | Measures the mean eccentricity of cells defined by the Actin channel. This distinguishes between elongated, fibroblast-like morphologies and rounded cells (e.g., in mitotic arrest or non-adherent states). |

| | | | |
|---|---|---|---|
| 159 | **actin texture entropy** | **code** | Calculates the entropy of the pixel intensities in the Actin channel. Higher entropy indicates a more disordered, complex cytoskeletal network, whereas lower entropy may indicate uniform staining or loss of structure. |
| 160 | **nucleus to cytoplasm area ratio** | **code** | Calculates the average ratio of nuclear area (DAPI) to total cell area (Actin) for segmented cells. This N/C ratio is a fundamental cellular metric that changes during differentiation, cancer progression, and drug response. |
| 161 | **tubulin actin pearson correlation** | **code** | Calculates the correlation between cytoskeletal channels (Tubulin/Green and Actin/Red). Changes in this correlation reflect gross cytoskeletal reorganization, such as cell rounding where both signals colocalize at the periphery. |
| 162 | **nuclear displacement index** | **code** | Measures the distance between the centroid of the nucleus and the centroid of the cell body (Actin). A high displacement indicates cell polarity or asymmetric morphology, while zero indicates a centered nucleus. |
| 163 | **global cell confluence** | **code** | Calculates the fraction of the total image area covered by cells (based on a combined mask of Actin and Tubulin). This serves as a proxy for cell density and proliferation/death rates. |
| 164 | **nucleus haralick contrast mean** | **code** | Computes the mean Haralick contrast of the DAPI texture within segmented nuclei. This quantifies chromatin condensation patterns, distinguishing between smooth interphase nuclei and textured mitotic or apoptotic nuclei. |
| 165 | **tubulin fiber alignment index mean** | **code** | Measures the anisotropy or directional coherence of tubulin structures within each cell. High values indicate bundled or parallel microtubules (e.g., Taxol effect), while low values indicate a meshwork. |
| 166 | **tubulin intensity cv global** | **code** | Calculates the coefficient of variation (std/mean) of pixel intensities in the Tubulin channel across the entire image. This measures the heterogeneity of the microtubule network; stabilized bundles (Taxanes) create high contrast/variance, while depolymerization (Vinca) creates diffuse signal. |
| 167 | **actin intensity skewness global** | **code** | Measures the skewness of the pixel intensity distribution in the Actin channel globally. A shift in skewness can indicate a change in the ratio of background to filamentous actin structures. |
| 168 | **tubulin cell entropy mean** | **code** | Computes the mean texture entropy of tubulin intensity within each cell. Higher entropy indicates more heterogeneous microtubule organization. |
| 169 | **actin radial distribution cv mean** | **code** | Measures the variation of Actin intensity as a function of distance from the cell center (nucleus). This captures the spatial organization of the cytoskeleton (e.g., cortical ring vs. diffuse). |
| 170 | **nucleus boundary roughness mean** | **code** | Calculates the roughness of the nuclear boundary (perimeter^2 / (4 * pi * area) normalized or similar metric focusing on high-frequency boundary changes). This can indicate nuclear envelope stress or early fragmentation. |
| 171 | **cytoplasm actin texture correlation mean** | **code** | Measures the Haralick correlation of Actin texture within the cytoplasm (excluding the nucleus). This assesses the linear dependency of gray levels, reflecting the structure of actin stress fibers. |
| 172 | **tubulin mean intensity cytoplasm** | **code** | Calculates the average intensity of the Tubulin (green) signal within the cytoplasmic region (cell mask minus nuclear mask). This quantifies the abundance of the microtubule network, which can be affected by tubulin-stabilizing or destabilizing drugs. |
| 173 | **actin mean intensity cytoplasm** | **code** | Calculates the average intensity of the Actin (red) signal within the cytoplasm. Changes in actin abundance or polymerization status are key indicators of cytoskeletal remodeling or cell motility inhibition. |
| 174 | **actin texture correlation mean** | **code** | Calculates the average GLCM Correlation for the Actin channel within cells. This measures the linear dependency of gray levels on those of neighboring pixels, capturing the directional structure of stress fibers versus diffuse cortical actin. |
| 175 | **dapi tubulin correlation cell mean** | **code** | Calculates the Pearson correlation coefficient between DAPI (blue) and Tubulin (green) pixel intensities within each cell. High correlation may indicate mitotic stages where tubulin (spindle) and DNA (chromosomes) are spatially colocalized. |
| 176 | **nuclear actin center distance mean** | **code** | Measures the mean Euclidean distance between the centroid of the nucleus and the centroid of the total cell (actin/tubulin). This quantifies cell polarity, as migratory cells often have the nucleus displaced towards the rear. |
| 177 | **tubulin skeleton branch density** | **code** | Skeletonizes the Tubulin (green) signal and calculates the density of branch points per unit area. This quantifies the complexity of the microtubule network, distinguishing between dense networks and sparse or bundled fibers. |
| 178 | **tubulin texture correlation mean** | **code** | Calculates the Haralick texture correlation of the Tubulin channel within cells. This measures the linear dependency of gray levels, which helps distinguish between fibrous (high correlation) and diffuse (low correlation) microtubule structures. |
| 179 | **micronucleus density** | **code** | Counts the number of small, detached DAPI-positive objects (micronuclei) relative to the total image area or cell count. Micronuclei are a hallmark of genomic instability and DNA damage. |
| 180 | **actin intensity cv mean** | **code** | Calculates the Coefficient of Variation (std/mean) of actin intensity within each cell. This captures the patchiness or concentration of actin into stress fibers versus a diffuse cortical distribution. |
| 181 | **dapi boundary intensity gradient** | **code** | Measures the mean gradient magnitude of the DAPI signal at the nuclear boundary. Sharp gradients indicate a well-defined nuclear envelope, while diffuse gradients may suggest nuclear envelope breakdown (prometaphase). |
| 182 | **cell perimeter mean** | **code** | Measures the mean perimeter length of segmented cells. An increased perimeter relative to area (high complexity) can indicate membrane ruffling, blebbing, or the formation of protrusions like invadopodia. |
| 183 | **cell shape form factor mean** | **code** | Computes the mean form factor (4 * pi * Area / Perimeter^2) of segmented cells. Values close to 1 indicate a perfect circle (e.g., rounded mitotic cells), while lower values indicate elongation or complex shapes (e.g., mesenchymal phenotype). |
| 184 | **actin intensity max mean** | **code** | Calculates the mean of the maximum pixel intensity in the Actin channel for each cell. High peak intensities often correspond to the presence of dense stress fibers or focal adhesions. |

| | | | |
|---|---|---|---|
| 185 | **actin radial distribution ratio** | **code** | Measures the ratio of Actin intensity in the outer 20% of the cell radius versus the inner 50%. This quantifies cortical actin enrichment versus diffuse cytoplasmic actin, relevant for cell adhesion and motility phenotypes. |
| 186 | **nuclear shape roundness mean** | **code** | Computes the mean roundness (4 * pi * area / perimeter^2) of nuclei. This scalar feature helps distinguish between healthy, round nuclei and the irregular, lobulated, or fragmented nuclei often seen in apoptosis or drug toxicity. |
| 187 | **actin total intensity mean** | **code** | Measures the average total integrated intensity of the Actin channel per cell. Changes in actin polymerization levels are indicative of cytoskeletal remodeling, stress fiber formation, or cell death processes. |
| 188 | **tubulin spatial distribution kurtosis** | **code** | Calculates the kurtosis of the pixel intensity distribution in the Tubulin channel for the whole image. High kurtosis can indicate the presence of bright bundles against a dark background (bundling phenotype). |
| 189 | **dapi contrast mean** | **code** | Calculates the local contrast in the DAPI channel. High contrast suggests distinct heterochromatin/euchromatin domains or nuclear speckles, which may be altered by drugs affecting DNA organization. |
| 190 | **tubulin fiber length mean** | **code** | Estimates the mean length of microtubule filaments after skeletonization of the Tubulin channel. Shortened fibers may indicate destabilization (e.g., Vinca alkaloids), while long, bundled fibers suggest stabilization (e.g., Taxanes). |
| 191 | **tubulin intensity radial cv mean** | **code** | Calculates the coefficient of variation of tubulin intensity measured in concentric rings from the nucleus center to the cell boundary. This captures the spatial distribution of microtubules, distinguishing between diffuse cytoplasmic staining and concentration around the nucleus (e.g., in rounded mitotic cells). |
| 192 | **actin intensity kurtosis mean** | **code** | Measures the kurtosis of the actin intensity distribution within cells. High kurtosis indicates the presence of extreme intensity values (e.g., bright stress fibers or focal adhesions against a dark background), while low kurtosis implies a more uniform distribution. |
| 193 | **actin tubulin overlap fraction** | **code** | Calculates the fraction of Tubulin signal that spatially overlaps with high Actin signal. This can indicate the coordination between the microtubule network and the actin cortex, which is relevant for cell motility and shape. |
| 194 | **tubulin fiber texture energy** | **code** | Computes the texture energy (e.g., from GLCM or laws texture energy measures) of the Tubulin channel globally. This quantifies the 'roughness' or prominence of microtubule fibers, distinguishing between diffuse tubulin (depolymerized) and strong fibrous networks (stabilized/bundled). |
| 195 | **nuclear shape eccentricity mean** | **code** | Measures the mean eccentricity of nuclei, indicating how elongated they are. This helps distinguish between normal interphase nuclei (oval), mitotic nuclei (often rounder or irregular depending on phase), and distorted nuclei due to mechanical stress or drug effects. |
| 196 | **micronucleus object ratio** | **code** | Calculates the ratio of small, detached DAPI objects (micronuclei) to normal-sized nuclei. Micronuclei are a hallmark of genomic instability and mitotic slippage, which are specific outcomes of drugs like Vincristine or Methotrexate mentioned in the knowledge base. |
| 197 | **cell spread area mean** | **code** | Measures the mean total area of the cell (cytoplasm + nucleus) using Actin or Tubulin channels. Significantly enlarged spread areas are a primary morphological signature of senescence, a phenotype noted in the RAG for certain drug treatments. |
| 198 | **cell shape solidity mean** | **code** | Calculates the mean solidity (area / convex hull area) of cells. Low solidity indicates irregular, ruffled, or blebbing edges, which can be a sign of membrane instability, motility (invadopodia), or apoptotic blebbing. |
| 199 | **actin skeleton branch density** | **code** | Skeletonizes the Actin channel and measures the density of branch points. This quantifies the complexity of the actin network, distinguishing between dense meshworks and simplified or collapsed cytoskeletons. |
| 200 | **nucleus cytoplasm tubulin ratio** | **code** | Ratio of mean Tubulin intensity in the nuclear region vs the cytoplasmic region. While Tubulin is cytoplasmic, signal bleed or specific transport defects might alter this ratio, or nuclear envelope breakdown in mitosis would equalize it. |
| 201 | **tubulin fiber anisotropy mean** | **code** | Analyzes the Tubulin (green) channel to measure the directional consistency (anisotropy) of microtubule fibers. High anisotropy indicates aligned bundles (stabilization), while low anisotropy suggests a disorganized meshwork (destabilization). |
| 202 | **cellular foreground fraction** | **code** | Calculates the fraction of the total image area occupied by cellular material (foreground). This serves as a proxy for cell confluence and can indicate cytotoxicity (cell loss) or cytostatic effects (growth inhibition). |
| 203 | **actin fiber texture contrast** | **code** | Computes the Haralick Contrast of the Actin channel within segmented cells. High contrast indicates distinct stress fibers, while low contrast suggests a diffuse or depolymerized actin cortex. |
| 204 | **tubulin spatial moment variance** | **code** | Calculates the variance of the spatial distribution of tubulin intensity relative to the cell centroid (Second Moment). This describes how spread out or concentrated the microtubule network is. |
| 205 | **nearest neighbor distance cv** | **code** | Computes the coefficient of variation (CV) of the distance to the nearest neighbor for all nuclei. High CV implies clustering (heterogeneity), while low CV implies a regular grid-like or uniform distribution. |
| 206 | **actin area coverage ratio** | **code** | Calculates the ratio of pixels exceeding a threshold in the Actin channel to the total image area. This measures cell confluence and spreading, which are affected by cytoskeletal drugs. |
| 207 | **tubulin texture contrast global** | **code** | Calculates the Haralick contrast of the Tubulin channel across the entire image. High contrast corresponds to sharp, defined microtubules (or bundles), while low contrast indicates diffuse or depolymerized tubulin. |
| 208 | **actin fiber texture energy** | **code** | Measures the texture energy of the Actin (red) channel using Laws' texture energy measures or similar filter banks. This feature captures the prominence of cytoskeletal structures like stress fibers, which appear as high-frequency textural patterns compared to diffuse cortical actin. |

| | | | |
|---|---|---|---|
| 209 | **nuclear area cv** | **code** | Calculates the coefficient of variation (std dev / mean) of the areas of segmented nuclei. This scalar captures the heterogeneity of nuclear sizes in the population, which is relevant for detecting mixed populations of normal, micronucleated, or enlarged senescent cells. |
| 210 | **actin cortical enrichment ratio** | **code** | Measures the ratio of Actin (red) intensity at the cell boundary (cortex) versus the cell center. A high ratio indicates cortical actin rings, often seen in rounded cells or specific adhesion states. |
| 211 | **dapi boundary roughness** | **code** | Calculates the mean roughness (ratio of perimeter to convex hull perimeter) of nuclear boundaries. Increased roughness can indicate nuclear blebbing or fragmentation associated with apoptosis. |
| 212 | **tubulin skeleton density** | **code** | Applies a skeletonization algorithm to the thresholded Tubulin channel (Channel 1) and calculates the density of skeleton pixels. This proxies the abundance of polymerized microtubule fibers versus diffuse background signal. |
| 213 | **dapi intensity skewness** | **code** | Calculates the skewness of the pixel intensity histogram for the DAPI channel (Channel 0). A shift towards higher skewness indicates a subset of very bright pixels, characteristic of chromatin condensation or formation of bright foci. |
| 214 | **perinuclear tubulin intensity** | **code** | Measures the mean intensity of Tubulin (Channel 1) in a ring-shaped region immediately surrounding each segmented nucleus. This specifically targets phenotypes like the 'nuclear cap' or perinuclear accumulation of organelles/tubulin. |
| 215 | **actin texture contrast** | **code** | Computes the Haralick Contrast feature on the Actin channel (Channel 2). This measures the local intensity variation, helping to distinguish between cells with distinct stress fibers (high contrast) and those with diffuse cortical actin (low contrast). |
| 216 | **n c ratio mean** | **code** | Computes the average Nucleus-to-Cytoplasm area ratio. This is a critical cellular metric that changes during cell cycle progression and in response to drugs affecting cell growth vs. division. |
| 217 | **dapi glcm contrast mean** | **code** | Calculates the mean contrast from the Gray Level Co-occurrence Matrix (GLCM) of the nuclear region. This texture feature quantifies local variations in DNA density, effectively capturing the 'graininess' of chromatin which changes under drug-induced stress or DNA damage. |
| 218 | **tubulin intensity kurtosis mean** | **code** | Calculates the kurtosis of the pixel intensity distribution within the tubulin channel for each cell. High kurtosis indicates a distribution with heavy tails or peaks, suggesting the presence of bright, distinct structures (like bundles or spindles) against a dark background, versus a diffuse microtubule network. |
| 219 | **tubulin fiber alignment coherence mean** | **code** | Uses structure tensor analysis to measure the local coherence of tubulin fibers. High coherence indicates parallel bundling of microtubules (e.g., taxane stabilization), while low coherence suggests a disordered or depolymerized network. |
| 220 | **channel correlation dapi tubulin** | **code** | Computes the Pearson correlation coefficient between pixel intensities of the DAPI and Tubulin channels within the cell mask. High correlation may indicate mitotic stages where chromosomes and spindle microtubules are spatially coincident, or nuclear accumulation of tubulin. |
| 221 | **actin radial intensity gradient mean** | **code** | Measures the gradient of actin intensity from the cell center (nucleus) to the periphery. This quantifies the distribution of actin, distinguishing between cells with strong cortical actin rings and those with diffuse cytoplasmic actin or stress fibers. |
| 222 | **dapi area coefficient of variation** | **code** | Calculates the coefficient of variation (std/mean) of nuclear areas across the entire image. A high value indicates high heterogeneity in nuclear size, suggesting a population with mixed cell cycle stages, micronuclei, or apoptotic fragmentation. |
| 223 | **tubulin center of mass displacement mean** | **code** | Measures the distance between the geometric center of the nucleus and the intensity-weighted center of mass of the tubulin network. Large displacements indicate cell polarity or asymmetric microtubule organization typical of migrating or polarized cells. |
| 224 | **dapi nearest neighbor distance mean** | **code** | Computes the average distance from each nucleus to its nearest neighbor. This spatial feature quantifies cell clustering or confluence, which can be affected by contact inhibition or drugs causing cell detachment. |
| 225 | **actin texture lacunarity mean** | **code** | Calculates the lacunarity of the actin channel, a fractal dimension measure that quantifies the 'gappiness' or heterogeneity of the cytoskeletal meshwork. This helps characterize the complexity of the actin organization. |
| 226 | **actin intensity std global** | **code** | Computes the standard deviation of pixel intensities in the Actin (red) channel for the whole image. This measures the heterogeneity of the cytoskeleton; stress fibers create high variance, while diffuse actin results in lower variance. |
| 227 | **dapi intensity cv** | **code** | Calculates the Coefficient of Variation (std/mean) of pixel intensities within the DAPI channel across the entire image. This provides a global measure of nuclear heterogeneity and cell density variations. |
| 228 | **dapi actin distance mean** | **code** | Measures the Euclidean distance between the centroid of the nucleus and the centroid of the cell (Actin). Large offsets indicate cell polarity or asymmetric spreading. |
| 229 | **dapi granularity mean** | **code** | Measures the granularity of the nuclear channel using morphological opening operations with increasing structuring element sizes. Specifically targets the detection of nuclear foci or chromatin condensation. |
| 230 | **cell elongation mean** | **code** | Calculates the average ratio of the major axis to the minor axis of the fitted ellipse for each cell. This quantifies cell shape changes, such as the transition from a spread, mesenchymal shape to a rounded morphology or an elongated stress phenotype. |
| 231 | **actin intensity std** | **code** | Calculates the standard deviation of intensity in the Actin channel within cells. High standard deviation may indicate the presence of distinct structures like stress fibers, while low std suggests diffuse actin. |
| 232 | **tubulin haralick correlation mean** | **code** | Computes the Haralick Correlation for the Tubulin channel. This measures the linear dependency of gray levels, which can capture the directional structure of microtubule networks. |
| 233 | **cell orientation std** | **code** | Calculates the standard deviation of the orientation angles of cells in an image. Low variance indicates alignment (e.g., in flow or specific tissue-like structures), while high variance indicates random orientation. |
| 234 | **cytoplasm intensity entropy actin** | **code** | Computes the Shannon entropy of pixel intensities in the Actin channel within the cytoplasm (excluding the nucleus). This captures the complexity and disorder of the cytoskeletal network. |

| | | | |
|---|---|---|---|
| 235 | **tubulin actin intensity ratio mean** | **code** | Calculates the ratio of total Tubulin intensity to total Actin intensity per cell. This captures the balance between the microtubule and actin cytoskeletons, which can be disrupted by specific inhibitors. |
| 236 | **tubulin fiber alignment anisotropy** | **code** | Measures the anisotropy of texture in the Tubulin channel, indicating how strongly the microtubules are aligned in a specific direction. This is relevant for detecting microtubule bundling (taxanes) or disruption (vinca alkaloids). |
| 237 | **cell nucleus elongation** | **code** | Measures elongation of segmented nuclei from nuclear shape (eccentricity / aspect). Higher values indicate more elongated nuclei. |
| 238 | **tubulin intensity cv mean** | **code** | Calculates the mean Coefficient of Variation (std/mean) of Tubulin intensity within cells. High variance often indicates the presence of bright, discrete structures (bundles/spindles) against a dark background. |
| 239 | **nuclear fragmentation count** | **code** | Counts the number of small, detached nuclear bodies (micronuclei) in the image. An increase in this count is a direct indicator of genomic instability or mitotic slippage (e.g., Vinca alkaloids). |
| 240 | **tubulin fiber skeleton length mean** | **code** | Applies skeletonization to the Tubulin channel within each cell and calculates the mean total length of the skeleton per cell. This quantifies the extent of the microtubule network, distinguishing between polymerized networks and depolymerized diffuse signals. |
| 241 | **nucleus cell area ratio mean** | **code** | Calculates the mean ratio of nuclear area (DAPI) to total cell area (Actin/Tubulin). The N:C ratio is a fundamental cellular metric that changes during cell cycle progression and in response to drugs affecting cell growth vs division. |
| 242 | **tubulin texture haralick contrast mean** | **code** | Computes the mean Haralick Contrast of the Tubulin channel within cells. This measures the local intensity variation, distinguishing between smooth, diffuse tubulin staining and high-contrast fibrous networks. |
| 243 | **cell orientation coherence global** | **code** | Calculates the global coherence of local gradients in the Tubulin channel using the Structure Tensor. This measures how aligned the microtubule fibers are across the entire field of view, relevant for detecting stress fiber alignment. |
| 244 | **tubulin radial distribution peak** | **code** | Analyzes the radial intensity profile of Tubulin from the cell center to periphery. A peak near the center suggests perinuclear accumulation (e.g., collapsed network), while a uniform distribution suggests a healthy network. |
| 245 | **nucleus shape form factor mean** | **code** | Measures the mean form factor (4*pi*area/perimeter^2) of nuclei. This scalar quantifies nuclear circularity, which helps distinguish between normal interphase nuclei (high circularity) and lobulated or fragmented nuclei seen in apoptosis or specific drug-induced stress. |
| 246 | **tubulin texture gabor energy mean** | **code** | Measures the mean energy of Gabor filter responses on the Tubulin channel. This quantifies the strength and directionality of microtubule fibers, helping to detect phenotypes like microtubule bundling (high energy/directionality) or depolymerization (low energy/diffuse signal). |
| 247 | **actin cortical intensity ratio mean** | **code** | Calculates the ratio of actin intensity at the cell periphery (cortical) versus the cell interior. This feature identifies cytoskeletal reorganization, such as the formation of cortical actin rings or stress fibers, which are relevant to cell motility and adhesion. |
| 248 | **actin texture haralick contrast mean** | **code** | Computes the mean Haralick contrast of the Actin channel. This quantifies the local intensity variations, effectively measuring the 'roughness' of the actin cytoskeleton, which increases with the formation of distinct stress fibers or punctate structures. |
| 249 | **cell nucleus area ratio mean** | **code** | Calculates the mean ratio of nuclear area to total cell area (N:C ratio). This is a critical biological parameter that changes with cell differentiation, neoplastic transformation, and response to cytotoxic drugs. |
| 250 | **cell neighbors count mean** | **code** | Calculates the mean number of adjacent neighbors for each cell. This captures local cell density and clustering patterns, which can be affected by drugs that inhibit contact inhibition or cause cell detachment. |
| 251 | **tubulin intensity radial distribution** | **code** | Measures the ratio of Tubulin intensity in the perinuclear region versus the cell periphery. This can identify phenotypes like perinuclear microtubule collapse or broad distribution in spreading cells. |
| 252 | **cell shape eccentricity mean** | **code** | Computes the mean eccentricity of segmented cells (0=circle, 1=line) based on the Actin channel. This quantifies cell elongation, helping to distinguish between rounded mitotic cells and elongated mesenchymal-like phenotypes. |
| 253 | **cyto nucleus area ratio mean** | **code** | Calculates the mean ratio of cytoplasmic area to nuclear area (N:C ratio). Changes in this ratio are indicative of cell growth, hypertrophy, or specific developmental states and are altered by many compounds. |
| 254 | **tubulin texture haralick entropy mean** | **code** | Computes the mean Haralick entropy of the Tubulin channel within cells. High entropy suggests a disordered or complex microtubule network, while low entropy might indicate diffuse staining or distinct bundling. |
| 255 | **tubulin spatial moment inertia** | **code** | Calculates the moment of inertia of the Tubulin intensity distribution relative to the cell centroid. This quantifies how dispersed the microtubule network is; relevant for distinguishing bundled vs. spread networks. |
| 256 | **cell local density knn dist** | **code** | Computes the mean distance to the k-nearest neighbors (e.g., k=3) for each cell centroid. This measures local cell density, which can indicate contact inhibition or toxicity-induced cell loss. |
| 257 | **nuclear margin intensity ratio** | **code** | Calculates the ratio of DAPI intensity at the nuclear boundary (rim) versus the center. This can detect marginalization of chromatin, a feature of early apoptosis. |
| 258 | **tubulin distribution radial cv** | **code** | Measures the coefficient of variation of Tubulin intensity in radial bins extending from the nucleus. This captures the spatial heterogeneity of the microtubule network (e.g., perinuclear accumulation vs. uniform spreading). |
| 259 | **actin intensity total** | **code** | Sums the total intensity of the Actin channel across the image. A drastic reduction might indicate cell loss or actin depolymerization, while an increase could suggest stress fiber formation or cell spreading. |

| | | | |
|---|---|---|---|
| 260 | **tubulin fiber linearity index** | **code** | Analyzes the Tubulin channel to measure the linearity or alignment of microtubule fibers using structure tensor analysis. This distinguishes between the disordered meshwork of normal cells and the rigid bundles induced by Taxanes. |
| 261 | **actin boundary roughness** | **code** | Measures the roughness of the cell perimeter (perimeter / convex perimeter). This captures membrane ruffling and irregularities associated with motility or cytoskeletal disruption. |
| 262 | **tubulin texture angular second moment mean** | **code** | Computes the average Angular Second Moment (ASM) from the Gray Level Co-occurrence Matrix (GLCM) of the Tubulin channel within cells. ASM measures textural uniformity; high values indicate uniform intensity (diffuse tubulin), while low values suggest structured fibers. |
| 263 | **actin polarity vector magnitude mean** | **code** | Calculates the magnitude of the vector between the cell centroid and the intensity-weighted centroid of the Actin channel. This quantifies cell polarity and the asymmetry of the cytoskeleton, relevant for migration and adhesion phenotypes. |
| 264 | **dapi tubulin correlation coefficient** | **code** | Computes the Pearson correlation coefficient between DAPI and Tubulin pixel intensities within the cell area. This measures the degree of spatial overlap or exclusion between the nucleus and the microtubule network. |
| 265 | **cell perimeter roughness mean** | **code** | Measures the ratio of the cell perimeter to the perimeter of the convex hull of the cell. This captures membrane complexity, such as blebbing (apoptosis/pyroptosis) or filopodia extensions, versus smooth boundaries. |
| 266 | **nuclear intensity edge center ratio** | **code** | Calculates the ratio of mean DAPI intensity at the nuclear periphery (edge) versus the nuclear center. This can detect chromatin margination, a specific feature of early apoptosis. |
| 267 | **nuclear intensity integrated std** | **code** | Measures the standard deviation of the integrated intensity (sum of pixel values) of nuclei across the image. High variance suggests a heterogeneous population, potentially indicating mixed cell cycle states or subpopulations undergoing apoptosis/pyroptosis. |
| 268 | **nucleus tubulin correlation** | **code** | Calculates the Pearson correlation coefficient between the DAPI (nucleus) and Tubulin (microtubule) channels on a pixel-by-pixel basis. This measures the spatial overlap and exclusion patterns, which can change during mitosis (spindle formation) or cell rounding. |
| 269 | **tubulin intensity spatial distribution ratio** | **code** | Calculates the ratio of tubulin intensity in the perinuclear region versus the cell periphery. This can quantify the collapse of the microtubule network around the nucleus (e.g., due to destabilizers) versus a spread-out network. |
| 270 | **dapi granularity spectrum slope** | **code** | Calculates the slope of the power spectrum of the DAPI channel image. This characterizes the distribution of granular sizes in the chromatin, potentially capturing texture changes associated with condensation or DNA damage foci. |
| 271 | **tubulin radial distribution entropy** | **code** | Calculates the entropy of the radial distribution of tubulin intensity centered on the nucleus. This quantifies the disorder in the radial arrangement of microtubules, distinguishing organized radial arrays from disordered networks. |
| 272 | **cytoskeleton polarity vector magnitude** | **code** | Calculates the magnitude of the vector between the nuclear centroid and the cell centroid (based on Actin/Tubulin). High values indicate polarized cells (migratory phenotype), while low values indicate symmetric/round cells. |
| 273 | **micronucleus count mean** | **code** | Counts the average number of small, detached DAPI-positive objects per cell. Micronuclei are a hallmark of genomic instability and mitotic catastrophe, often induced by DNA-damaging agents. |
| 274 | **tubulin radial intensity cv** | **code** | Calculates the coefficient of variation of Tubulin intensity in radial bins extending from the nucleus center. High variation implies directional structures (like spindles), while low variation implies a uniform halo. |
| 275 | **actin gradient magnitude mean** | **code** | Computes the mean magnitude of the image gradient in the Actin channel. This serves as a proxy for cytoskeletal complexity and edge sharpness, differentiating between smooth and ruffled/spiky cell margins. |
| 276 | **microtubule fiber alignment index** | **code** | Measures the anisotropy of the Tubulin channel using structure tensors or gradient orientation histograms. High alignment indicates microtubule bundling (e.g., Taxol effect), while low alignment suggests a diffuse or depolymerized network. |
| 277 | **nuclear shape irregularity index** | **code** | Computes the mean compactness (perimeter squared divided by area) of segmented nuclei. This quantifies deviations from a perfect circle, capturing nuclear blebbing, fragmentation, or irregular shapes associated with apoptosis or mitotic catastrophe. |
| 278 | **dapi intensity kurtosis global** | **code** | Calculates the kurtosis of the pixel intensity histogram for the DAPI channel across the entire image. High kurtosis indicates a heavy tail of very bright pixels, serving as a proxy for the presence of condensed mitotic chromosomes or pyknotic nuclei. |
| 279 | **micronuclei detection count** | **code** | Counts the number of small, distinct DAPI-positive objects that are separated from the main nuclei. An increase in this count is a marker of genomic instability or mitotic slippage (e.g., Vinca alkaloids). |
| 280 | **cytoskeleton nucleus area ratio** | **code** | Calculates the mean ratio of the Actin cell area to the DAPI nuclear area per cell (N:C ratio). Changes in this ratio track cell spreading (senescence) versus shrinkage (apoptosis) or rounding. |
| 281 | **tubulin texture lacunarity** | **code** | Measures the lacunarity (gappiness) of the Tubulin network. High lacunarity indicates large gaps or a disrupted network (depolymerization), while low lacunarity suggests a dense, uniform meshwork. |
| 282 | **nuclear distribution clustering coefficient** | **code** | Uses spatial statistics (e.g., nearest neighbor distance or Ripley's K) on nuclear centroids to quantify whether cells are clustered (colonies) or randomly distributed. This can reflect contact inhibition or toxicity-induced sparseness. |
| 283 | **dapi tubulin intensity correlation** | **code** | Calculates the Pearson correlation coefficient between pixel intensities of DAPI and Tubulin channels. This helps identify if tubulin is spatially excluded from the nucleus (interphase) or overlapping/collapsed (mitosis/toxicity). |
| 284 | **actin edge gradient magnitude** | **code** | Computes the mean gradient magnitude at the boundaries of segmented cells in the Actin channel. This measures the sharpness of the cell cortex, distinguishing between defined cortical actin and diffuse/blebbing edges. |

| | | | |
|---|---|---|---|
| 285 | **total cellular fluorescence actin** | **code** | Calculates the integrated density (sum of pixel values) of the Actin channel across all segmented cells. This proxies total protein content or cell mass, which can increase in senescent cells. |
| 286 | **nucleus tubulin center offset mean** | **code** | Calculates the Euclidean distance between the centroid of the nucleus and the intensity-weighted centroid of the Tubulin signal within the cell. Large offsets may indicate asymmetric polarization or 'nuclear cap' formation. |
| 287 | **dapi contrast haralick mean** | **code** | Computes the mean Haralick Contrast feature for the DAPI channel. This texture feature is sensitive to sharp intensity transitions, such as those found in nuclear foci or condensed chromosomes. |
| 288 | **tubulin fiber texture strength** | **code** | Calculates the texture contrast (e.g., Haralick contrast) of the Tubulin (green) channel within the cell cytoplasm. This quantifies the difference between diffuse tubulin and distinct microtubule bundling or polymerization phenotypes. |
| 289 | **actin fiber texture strength** | **code** | Calculates the texture contrast of the Actin (red) channel. This helps differentiate between cells with strong stress fibers (high contrast) and those with diffuse or cortical actin (low contrast/smooth). |
| 290 | **dapi actin pearson correlation** | **code** | Calculates the Pearson correlation between DAPI (blue) and Actin (red) intensities. This can serve as a negative control or detect unusual phenotypes where actin collapses onto the nucleus (e.g., nuclear cap formation). |
| 291 | **dapi texture entropy global** | **code** | Calculates the entropy of the pixel intensity distribution in the DAPI channel across the entire image. High entropy indicates complex, heterogeneous chromatin texture, while low entropy suggests homogeneity. |
| 292 | **tubulin spatial anisotropy global** | **code** | Measures the global spatial anisotropy of the Tubulin channel using structure tensors. High anisotropy indicates aligned fibers (bundles), while low anisotropy indicates a random meshwork. |
| 293 | **tubulin intensity skewness global** | **code** | Measures the skewness of the intensity histogram of the Tubulin channel. Positive skewness indicates a background-dominated image with sparse bright structures (fibers), while lower skewness suggests diffuse staining. |
| 294 | **tubulin spatial frequency mean** | **code** | Measures the mean spatial frequency of the Tubulin channel (e.g., using Laplacian variance). High values indicate fine, detailed microtubule networks; low values indicate blurred or diffuse structures. |
| 295 | **nucleus max intensity mean** | **code** | Calculates the mean of the maximum pixel intensity within each segmented nucleus. This is sensitive to bright nuclear bodies or DNA damage foci (like gamma-H2AX if it were a channel, or bright DAPI spots). |
| 296 | **actin intensity mean per cell** | **code** | Calculates the mean intensity of the Actin channel within segmented cell boundaries. Changes in actin abundance or polymerization status are captured by this intensity metric. |
| 297 | **cell roundness factor** | **code** | Computes the form factor (4*pi*area/perimeter^2) of the whole cell boundary defined by Actin. High roundness is a strong indicator of mitotic arrest or detachment (e.g., Vinca alkaloids). |
| 298 | **nuclear intensity variance** | **code** | Calculates the variance of pixel intensities within segmented nuclei. High variance can indicate chromatin condensation or punctate staining patterns associated with DNA damage. |
| 299 | **nucleus to cytoplasm ratio** | **code** | Calculates the ratio of the total nuclear area (DAPI) to the total cytoplasmic area (Actin/Tubulin minus DAPI). This N/C ratio is a fundamental cellular metric often altered in cancer and drug response. |
| 300 | **actin polarity index** | **code** | Measures the displacement between the centroid of the nucleus (DAPI) and the centroid of the cell body (Actin). A large displacement vector magnitude indicates polarized cells, relevant for migration or adhesion studies. |
| 301 | **dapi boundary irregularity** | **code** | Measures the irregularity of the nuclear boundary (e.g., perimeter^2 / area) in the DAPI channel. Highly irregular boundaries (blebbing) are a hallmark of apoptosis or nuclear envelope stress. |
| 302 | **cytoskeleton radial distribution** | **code** | Measures the radial intensity profile of Tubulin/Actin from the nuclear center to the cell periphery. Changes in this profile capture phenotypes like cell spreading vs. rounding up. |
| 303 | **total image intensity ratio actin dapi** | **code** | Calculates the ratio of total integrated intensity of Actin to DAPI for the whole image. This global metric can indicate cell size relative to DNA content (hypertrophy vs. atrophy). |
| 304 | **dapi texture contrast mean** | **code** | Measures the mean Haralick contrast of the DAPI channel within segmented nuclei. High contrast suggests heterogeneous chromatin condensation, a hallmark of early apoptosis or specific mitotic phases. |
| 305 | **actin radial distribution gradient** | **code** | Measures the gradient of actin intensity from the cell center (nucleus) to the cell periphery. This quantifies the distribution of the cytoskeleton (e.g., cortical actin vs. stress fibers). |
| 306 | **nucleus major axis orientation entropy** | **code** | Calculates the entropy of the orientation angles of the major axes of all nuclei. Low entropy implies alignment (e.g., flow or stress), while high entropy implies random orientation. |
| 307 | **actin intensity distribution skewness** | **code** | Calculates the skewness of the pixel intensity histogram for the Actin channel. A shift in skewness can indicate a transition from diffuse cytoplasmic actin (uniform) to stress fibers or cortical rings (high intensity subsets). |
| 308 | **nuclear major axis orientation coherence** | **code** | Calculates the coherence (alignment) of the major axes of all nuclei in an image. High coherence might indicate flow or alignment due to external forces, while random orientation is expected in static culture; deviations might suggest organized tissue-like structures or artifacts. |
| 309 | **cytoplasm area mean** | **code** | Calculates the mean area of the cytoplasm (Cell Area - Nuclear Area). Changes in cytoplasmic volume are key indicators of cell health, swelling (necrosis/pyroptosis), or shrinkage (apoptosis). |
| 310 | **dapi glcm entropy mean** | **code** | Calculates the mean entropy of the Gray-Level Co-occurrence Matrix (GLCM) for the DAPI channel within nuclei. This quantifies chromatin texture heterogeneity; high entropy suggests condensed or fragmented chromatin often seen in apoptosis or mitotic arrest. |
| 311 | **actin glcm contrast mean** | **code** | Calculates the GLCM contrast of the Actin channel. High contrast indicates the presence of distinct structural features like stress fibers or cortical rings, as opposed to a uniform actin distribution. |
| 312 | **dapi actin correlation mean** | **code** | Computes the Pearson correlation between DAPI and Actin channels. This can capture phenotypes where actin collapses onto the nucleus or where the nucleus is displaced, altering the typical spatial exclusion. |

| | | | |
|---|---|---|---|
| 313 | **actin major axis length mean** | **code** | Calculates the mean length of the major axis of the ellipse fitting the actin cytoskeleton of segmented cells. This quantifies cell elongation, helping to distinguish between flattened, mesenchymal-like phenotypes and rounded, amoeboid or mitotic cells. |
| 314 | **nuclear haralick entropy mean** | **code** | Calculates the mean Haralick entropy (from GLCM) of the texture within segmented nuclei. This quantifies the randomness of chromatin organization; condensed chromatin (mitosis/apoptosis) typically shows different entropy levels compared to euchromatin in healthy cells. |
| 315 | **tubulin structure tensor coherence** | **code** | Computes the coherence of the structure tensor in the Tubulin channel, averaged over the image. This measures the degree of local orientation alignment (anisotropy) of microtubule fibers; high coherence suggests aligned bundles, while low coherence suggests a disorganized meshwork. |
| 316 | **nuclear integrated intensity cv** | **code** | Calculates the Coefficient of Variation (std/mean) of the integrated DAPI intensity across the population of nuclei. This measures the heterogeneity of DNA content, reflecting the distribution of cells across cell cycle phases (G1 vs G2/M) or the presence of aneuploidy. |
| 317 | **dapi intensity mass displacement** | **code** | Calculates the distance between the geometric centroid and the intensity-weighted centroid of each nucleus. This scalar captures asymmetric chromatin distribution, which can occur during nuclear cap formation or early karyorrhexis. |
| 318 | **tubulin local binary pattern energy** | **code** | Computes the energy of the Local Binary Pattern (LBP) histogram for the Tubulin channel. LBP is robust to illumination changes and captures local micro-texture; energy reflects the uniformity of the microtubule network texture. |
| 319 | **actin intensity radial gradient** | **code** | Measures the gradient of Actin intensity from the cell center (nucleus) to the cell periphery. This distinguishes cells with cortical actin rings from those with diffuse or perinuclear actin accumulation. |
| 320 | **actin tubulin correlation r** | **code** | Computes the Pearson correlation coefficient between Actin and Tubulin pixel intensities within each cell. This measures the degree of colocalization or spatial coordination between the two cytoskeletal networks. |
| 321 | **tubulin fiber texture contrast mean** | **code** | Calculates the Haralick contrast of the Tubulin channel within cell boundaries. High contrast corresponds to sharp, distinct microtubule fibers (bundling), while low contrast suggests diffuse or depolymerized tubulin. |
| 322 | **tubulin actin manders overlap mean** | **code** | Computes Manders' overlap coefficient for Tubulin and Actin channels. This quantifies the fraction of the cytoskeleton that is spatially co-localized, reflecting the structural integrity and organization of the cell's framework. |
| 323 | **nucleus orientation disorder** | **code** | Calculates the standard deviation of the major axis orientations of nuclei in the image. High disorder suggests random orientation, while low disorder might indicate alignment due to flow or contact guidance. |
| 324 | **cell area actin mean** | **code** | Calculates the mean area of the entire cell body defined by the Actin channel. This captures cell spreading or shrinkage, which are key indicators of cytoskeletal disruption or senescence (enlarged, flattened cells). |
| 325 | **nucleocytoplasmic ratio mean** | **code** | Calculates the mean ratio of nuclear area (DAPI) to cytoplasmic area (Actin minus DAPI) per cell. The N/C ratio is a fundamental cellular metric often altered in cancer progression and response to cytotoxic drugs. |
| 326 | **tubulin texture gabor energy** | **code** | Measures the mean Gabor filter energy in the Tubulin channel within cell boundaries. This texture feature specifically targets the detection of linear structures, quantifying the difference between diffuse tubulin and distinct microtubule bundles. |
| 327 | **cell polarity nucleus actin offset** | **code** | Measures the distance between the centroid of the nucleus (DAPI) and the centroid of the cell body (Actin). This vector magnitude quantifies cell polarity and asymmetry, which can be disrupted by cytoskeletal drugs. |
| 328 | **tubulin intensity cv intracell** | **code** | Computes the mean Coefficient of Variation (std/mean) of Tubulin intensity within individual cells. This quantifies the heterogeneity of the microtubule network (e.g., high variance in bundled/aggregated states vs low in diffuse states). |
| 329 | **tubulin radial distribution kurtosis** | **code** | Measures the kurtosis of the radial intensity distribution of tubulin relative to the cell center. High kurtosis indicates tubulin concentrated in a specific ring or spot (e.g., monoastral spindle), while low kurtosis implies diffuse distribution. |
| 330 | **actin edge to center intensity ratio** | **code** | Calculates the ratio of Actin intensity at the cell periphery (cortical actin) versus the cell center. This helps distinguish cells with strong cortical rings (rounded/mitotic) from those with basal stress fibers. |
| 331 | **dapi nuclear count** | **code** | Counts the total number of distinct nuclei identified in the DAPI channel. This serves as a proxy for cell count and proliferation, which is fundamental for assessing cytotoxicity or growth inhibition. |
| 332 | **dapi nuclear area mean** | **code** | Calculates the average area of segmented nuclei in the DAPI channel. Changes in nuclear size can indicate cell cycle arrest (enlargement in G2/M) or apoptosis (shrinkage/pyknosis). |
| 333 | **dapi nuclear circularity mean** | **code** | Measures the average circularity (4*pi*area/perimeter^2) of nuclei. Deviations from circularity can signal nuclear blebbing, fragmentation, or mitotic deformation. |
| 334 | **tubulin radial intensity slope** | **code** | Measures the gradient of Tubulin intensity radiating outwards from the nuclear centroid. A steep drop-off indicates perinuclear accumulation, while a flat slope indicates diffuse distribution. |
| 335 | **tubulin intensity cv cytoplasm** | **code** | Computes the coefficient of variation (std/mean) of pixel intensities in the Tubulin channel within the cytoplasmic region (excluding the nucleus). This quantifies the texture heterogeneity of the microtubule network, distinguishing between diffuse tubulin (depolymerized) and bundled/rigid filaments (stabilized). |
| 336 | **dapi boundary sharpness mean** | **code** | Measures the gradient magnitude at the boundary of the nucleus. A sharp boundary suggests an intact nuclear envelope, while a diffuse boundary may indicate nuclear envelope breakdown (NEBD) during prometaphase. |
| 337 | **nucleus to cytoplasm ratio mean** | **code** | Calculates the mean ratio of nuclear area to cytoplasmic area (Cell Area - Nuclear Area). A classic cytological metric that changes drastically during cell cycle progression or toxic response. |

| | | | |
|---|---|---|---|
| 338 | **tubulin anisotropy mean** | **code** | Measures the mean anisotropy of the Tubulin channel texture within cells. High anisotropy indicates strongly aligned fibers (e.g., Taxol-induced bundles), while low anisotropy indicates isotropic/diffuse signal. |
| 339 | **actin orientation coherence mean** | **code** | Computes the coherence of the structure tensor (gradient orientation) in the Actin channel. This measures how aligned the stress fibers are; high coherence indicates parallel stress fibers, low coherence indicates disorganized cytoskeleton. |
| 340 | **nucleus tubulin distance mean** | **code** | Calculates the Euclidean distance between the centroid of the nucleus and the weighted centroid of the Tubulin signal. A large shift may indicate cell polarization or displacement of the microtubule organizing center. |
| 341 | **nuclear intensity mean** | **code** | Computes the mean pixel intensity of the DAPI channel within segmented nuclear regions. This serves as a proxy for DNA content (ploidy) and chromatin condensation levels. |
| 342 | **tubulin haralick contrast** | **code** | Computes the Haralick Contrast texture feature for the Tubulin channel. This measures the local intensity variation, capturing the 'roughness' or distinctness of microtubule structures versus background. |
| 343 | **dapi tubulin distance center mass** | **code** | Calculates the distance between the center of mass of the DAPI signal (nucleus) and the Tubulin signal (cytoskeleton) for each cell. Large offsets can indicate cell polarization or asymmetric spreading. |
| 344 | **actin intensity std cytoplasm** | **code** | Measures the standard deviation of Actin intensity specifically within the cytoplasmic region (excluding the nucleus). High variance may indicate stress fibers or focal adhesions, while low variance suggests diffuse actin. |
| 345 | **nuclear shape roughness** | **code** | Measures the roughness of the nuclear boundary (perimeter^2 / (4 * pi * area) - 1, or similar metric). High roughness is associated with nuclear blebbing or irregularities seen in cellular stress. |
| 346 | **nuclear cap presence** | **vlm** | Detects the presence of 'nuclear caps'—asymmetric clustering of organelles or nuclear envelope expansion—which distinguishes oxidative stress from general S-phase arrest. |
| 347 | **microtubule bundling severity** | **vlm** | Scores the visual severity of microtubule bundling, looking for thick, rigid, high-intensity fibers compared to a fine meshwork. This is a key phenotype for taxane-class drugs. |
| 348 | **cell blebbing index** | **vlm** | Estimates the extent of membrane blebbing (bulges in the plasma membrane), a key feature of apoptosis and pyroptosis. |
| 349 | **tubulin mesh complexity** | **vlm** | Rates the visual complexity and density of the microtubule meshwork in interphase cells. |
| 350 | **anaphase bridge detection** | **vlm** | Detects the presence of anaphase bridges (DNA strands connecting separating chromosomes), a sign of chromosome segregation errors. |
| 351 | **nucleolar prominence** | **vlm** | Visually assesses the distinctness or prominence of nucleoli within the nucleus (often visible as dark holes in DAPI or specific texture). |
| 352 | **vacuolization index** | **vlm** | Detects the presence of cytoplasmic vacuoles (holes in the cytoplasmic staining), often a sign of cellular stress or death. |
| 353 | **polarized actin distribution** | **vlm** | Assesses whether actin is polarized to one side of the cell (indicative of migration) or evenly distributed. |
| 354 | **membrane blebbing index** | **vlm** | Visually estimates the degree of plasma membrane blebbing, a key feature of pyroptosis and apoptosis. This involves recognizing balloon-like protrusions on the cell surface. |
| 355 | **actin stress fiber alignment** | **vlm** | Visually scores the alignment and prominence of actin stress fibers. |
| 356 | **nuclear fragmentation severity** | **vlm** | Visually assesses the severity of nuclear fragmentation (karyorrhexis) or micronuclei formation, which are key indicators of apoptosis or genomic instability. |
| 357 | **multinucleation presence** | **vlm** | Visually identifies cells containing multiple nuclei within a single cytoplasm boundary. This phenotype indicates cytokinesis failure or mitotic slippage. |
| 358 | **cytoskeletal disruption index** | **vlm** | Visually rates the overall chaos or disruption of the actin/tubulin network compared to a healthy organized meshwork. |
| 359 | **membrane blebbing severity** | **vlm** | Visually assesses the presence of plasma membrane blebbing, a feature of apoptosis and pyroptosis, visible as irregular protrusions in the Actin/Tubulin channels. |
| 360 | **actin stress fiber abundance** | **vlm** | Visually scores the abundance of actin stress fibers. High values indicate numerous thick, parallel contractile fibers; low values correspond to cortical or diffuse actin without prominent stress fibers. |
| 361 | **nuclear shape irregularity** | **vlm** | Scores how irregular, lobulated, elongated, or non-elliptical the segmented nuclei appear in the blue DAPI channel. High values indicate distorted nuclear contours rather than smooth oval nuclei. This may capture nuclear stress, chromatin remodeling, mitotic defects, or compound-induced morphological abnormalities. |
| 362 | **apoptotic body presence** | **vlm** | Detects the presence of apoptotic bodies (small, condensed, fragmented cell remnants). Indicates late-stage apoptosis. |
| 363 | **spindle polarity defects** | **vlm** | Identifies multipolar spindles or other polarity defects beyond monoastrality. Relevant for various mitotic inhibitors. |
| 364 | **nuclear texture heterogeneity** | **vlm** | Visually assesses the heterogeneity of nuclear staining (e.g., granular vs smooth). Provides a semantic readout of nuclear texture heterogeneity. |
| 365 | **lamellipodia presence** | **vlm** | Detects the visual presence of broad, sheet-like actin protrusions (lamellipodia) at cell edges. This indicates active migration and intact actin polymerization machinery, which may be inhibited by actin-targeting drugs. |
| 366 | **monoastral spindle presence** | **vlm** | Visually estimates the presence and prevalence of monoastral spindles (rosette-like chromosome arrangement with radiating microtubules), a specific phenotype of Eg5 inhibition mentioned in the RAG knowledge. |
| 367 | **micronuclei presence** | **vlm** | Detects the presence of micronuclei—small, distinct DAPI-positive bodies near the main nucleus. These are markers of genomic instability or mitotic slippage. |

| | | | |
|---|---|---|---|
| 368 | **multinucleation count** | **vlm** | Visually counts cells that appear to have multiple nuclei within a single cytoplasmic boundary. Multinucleation is a specific sign of cytokinesis failure, often induced by drugs like Cytochalasin or Taxanes. |
| 369 | **invadopodia like structures** | **vlm** | Visually identifies actin-rich protrusions extending from the cell body, resembling invadopodia. This relates to cell motility and metastatic potential. |
| 370 | **anaphase bridge presence** | **vlm** | Visually detects DNA bridges connecting separating nuclei (anaphase bridges). This indicates DNA replication stress or telomere fusion/resolution defects. |
| 371 | **cell rounding severity** | **vlm** | Scores the visual degree to which segmented cells appear round rather than spread or elongated, using the actin and tubulin channels together with the cell/cytoplasm mask. High values indicate compact, circular cells consistent with mitotic arrest or cytotoxic rounding phenotypes. This is relevant for distinguishing compound effects that alter adhesion, cytoskeleton, or cell-cycle state. |
| 372 | **overall sample debris** | **vlm** | Visually estimates the amount of cellular debris (small, irregular fragments) in the image. High debris correlates with extensive cell death and lysis. |
| 373 | **nuclear molding** | **vlm** | Visually detects nuclear molding, where nuclei deform against each other or against vacuoles. This indicates crowding or high plasticity. |
| 374 | **actin intensity std mean** | **code** | Calculates the standard deviation of pixel intensities in the Actin channel per cell. High variance often corresponds to the presence of distinct structures like stress fibers, whereas low variance implies a diffuse cytoskeleton. |
| 375 | **tubulin radial distribution cv** | **code** | Measures the coefficient of variation of Tubulin intensity profiles radiating from the nucleus center to the cell periphery. This captures spatial organization, such as perinuclear clustering vs. uniform distribution. |
| 376 | **nuclear intensity std** | **code** | Computes the standard deviation of pixel intensities within segmented nuclei in the DAPI channel. High variation may indicate chromatin condensation or fragmentation associated with apoptosis or DNA damage. |
| 377 | **actin dapi distance mean** | **code** | Measures the mean distance between the centroid of the nucleus and the centroid of the cell body (defined by Actin). Large offsets can indicate cell polarization or motility. |
| 378 | **mitotic cell fraction** | **code** | Estimates the fraction of cells in the image that are in mitosis, defined by high DAPI intensity and high circularity. This quantifies mitotic arrest or proliferation rate. |
| 379 | **cytoskeleton area mean** | **code** | Calculates the mean area of the cytoskeleton (Tubulin/Actin) excluding the nuclear region. This specifically measures cytoplasmic spreading. |
| 380 | **tubulin spatial concentration** | **code** | Measures how concentrated the Tubulin signal is relative to the cell centroid (e.g., second moment of intensity). This helps distinguish between diffuse microtubules and concentrated structures like mitotic spindles. |
| 381 | **nuclear shape factor form factor** | **code** | Computes the form factor (4*pi*area / perimeter^2) of nuclei. Values close to 1 indicate perfect circles (e.g., healthy or rounded nuclei), while lower values indicate irregular shapes (e.g., blebbing or stress). |
| 382 | **tubulin fiber anisotropy global** | **code** | Measures the global anisotropy of the Tubulin channel using structure tensors. High anisotropy indicates aligned microtubule bundles (e.g., Taxane effect), while low anisotropy suggests a disorganized or depolymerized network. |
| 383 | **actin texture haralick contrast** | **code** | Computes the Haralick Contrast from the Gray Level Co-occurrence Matrix (GLCM) of the Actin channel. This quantifies the local intensity variation, capturing the roughness or distinctness of cytoskeletal filaments. |
| 384 | **actin boundary tortuosity mean** | **code** | Measures the tortuosity (perimeter / major_axis_length) of the cell boundary defined by Actin. High tortuosity indicates membrane ruffling or irregular protrusions (e.g., invadopodia). |
| 385 | **nucleus centroid displacement** | **code** | Calculates the distance between the geometric centroid of the nucleus and the geometric centroid of the whole cell. Large displacement indicates cell polarization or asymmetric spreading. |
| 386 | **dapi boundary intensity ratio** | **code** | Calculates the ratio of DAPI intensity at the nuclear boundary vs. the nuclear center. High boundary intensity can indicate chromatin margination, a feature of early apoptosis. |
| 387 | **actin tubulin correlation mean** | **code** | Calculates the Pearson correlation coefficient between Actin and Tubulin pixel intensities within each cell. Disruption of the cytoskeleton often leads to a decoupling of these two structural networks. |
| 388 | **tubulin spatial moment mean** | **code** | Calculates the second spatial moment of tubulin intensity relative to the cell center. This describes how spread out the microtubule network is (e.g., concentrated perinuclearly vs. extending to the periphery). |
| 389 | **nuclear boundary intensity gradient** | **code** | Measures the average intensity gradient magnitude at the boundary of segmented nuclei. Sharp gradients indicate well-defined nuclear envelopes, while diffuse gradients might suggest envelope breakdown. |
| 390 | **dapi tubulin overlap pearson** | **code** | Calculates the Pearson correlation coefficient between pixel intensities of DAPI and Tubulin channels within the cell. This measures the degree of spatial overlap; overlap increases during mitosis when the nuclear envelope breaks down and microtubules access the chromosomes. |
| 391 | **actin spatial radial distribution ratio** | **code** | Measures the ratio of Actin intensity in the outer 10% of the cell radius (cortex) versus the inner region. This distinguishes cells with strong cortical actin rings (often rounded/mitotic) from those with stress fibers throughout the cytoplasm. |
| 392 | **cellular intensity ratio actin tubulin** | **code** | Calculates the ratio of total Actin intensity to total Tubulin intensity per cell. This feature captures the balance between the cytoskeleton components, which can be disrupted differentially by actin-targeting vs. tubulin-targeting drugs. |
| 393 | **dapi boundary irregularity index** | **code** | Measures the irregularity of the nuclear boundary (perimeter^2 / (4 * pi * area) - 1, or similar metric). Irregular, lobulated nuclei are signs of nuclear stress, senescence, or specific drug toxicities. |
| 394 | **actin skeleton length total** | **code** | Skeletonizes the Actin channel and sums the total length of the resulting skeleton. This proxies the complexity and density of the actin filament network. |
| 395 | **nucleus actin centroid displacement** | **code** | Measures the distance between the centroid of the nucleus and the centroid of the cell (defined by Actin). This quantifies cell polarity and the centering of the nucleus. |

| | | | |
|---|---|---|---|
| 396 | **actin intensity mass displacement** | **code** | Measures the mean distance between the geometric centroid of the cell and the intensity-weighted centroid of the Actin channel. This quantifies cellular polarity and asymmetric cytoskeleton distribution, which can be affected by motility-modulating drugs. |
| 397 | **tubulin fiber alignment mean** | **code** | Estimates the coherence of local gradients in the Tubulin channel within cells to quantify fiber alignment. High alignment suggests bundling (e.g., Taxol effect), while low alignment indicates a meshwork or depolymerization. |
| 398 | **cell size heterogeneity** | **code** | Computes the standard deviation of cell areas within the image. A high value indicates a heterogeneous population (e.g., mixture of giant senescent cells and normal cells), which is a hallmark of certain drug responses. |
| 399 | **tubulin fiber anisotropy** | **code** | Quantifies the directionality and alignment of microtubule structures in the Tubulin channel. High anisotropy suggests ordered spindles or bundles, while low anisotropy indicates a disorganized meshwork or depolymerization. |
| 400 | **nucleus tubulin center offset** | **code** | Calculates the Euclidean distance between the centroid of the nucleus and the centroid of the tubulin intensity distribution. This measures cellular polarity and symmetry, which can be disrupted by drugs affecting cytoskeletal organization. |
| 401 | **tubulin skeleton length** | **code** | Computes the total length of the skeletonized microtubule network within a cell. This proxies the density and complexity of the microtubule network, which is reduced by depolymerizing agents like Vinca alkaloids. |
| 402 | **nucleus major axis length** | **code** | Measures the length of the major axis of the ellipse fitting the nucleus. Elongated nuclei can indicate specific cell cycle phases or deformation due to mechanical stress or drug effects. |
| 403 | **cell neighbor count** | **code** | Counts the number of adjacent cells touching or within a small radius of the target cell. This captures local cell density and confluence, which can influence cell morphology and drug response. |
| 404 | **dapi intensity std dev** | **code** | Calculates the standard deviation of pixel intensities in the DAPI channel. High standard deviation correlates with the presence of bright heterochromatin foci or apoptotic bodies against a darker nuclear background. |
| 405 | **nuclear shape irregularity mean** | **code** | Measures the mean irregularity of nuclear shapes (e.g., 1 - circularity) in the DAPI channel. Irregular nuclear envelopes can indicate DNA damage, nuclear fragmentation, or cellular stress. |
| 406 | **tubulin radial distribution peak mean** | **code** | Analyzes the radial intensity profile of Tubulin starting from the nuclear centroid. A sharp peak near the center suggests perinuclear collapse or monoastral spindles, while a flat profile indicates a spread network. |
| 407 | **actin filament contrast mean** | **code** | Calculates the local contrast (e.g., standard deviation or Haralick contrast) of the Actin channel. High contrast indicates distinct stress fibers, while low contrast suggests diffuse cortical actin. |
| 408 | **dapi chromatin heterogeneity mean** | **code** | Measures the entropy or intensity variance of the DAPI signal within nuclei. High heterogeneity correlates with chromatin condensation (e.g., in apoptosis or mitosis). |
| 409 | **total cellular occupancy fraction** | **code** | Calculates the fraction of the image area covered by cells (Actin/Tubulin foreground). This serves as a proxy for cell confluence and growth inhibition. |
| 410 | **dapi dna content mean** | **code** | Computes the mean integrated intensity (sum of pixel values) of the DAPI channel per nucleus. This proxies for DNA content, helping distinguish G1, S, and G2/M cell cycle phases. |
| 411 | **cell boundary solidity mean** | **code** | Measures the mean solidity (Area / Convex Hull Area) of cell boundaries defined by Actin. Low solidity indicates rough edges, blebbing, or dendritic protrusions. |
| 412 | **cell clustering index** | **code** | Calculates the average nearest-neighbor distance between nuclear centroids, normalized by cell density. Measures whether cells are clustered (colonies) or randomly distributed. |
| 413 | **dapi nuclear boundary intensity ratio** | **code** | Calculates the ratio of mean intensity at the nuclear boundary to the mean intensity of the nuclear center. This can help identify 'nuclear cap' phenotypes or marginal chromatin condensation. |
| 414 | **actin star morphology count** | **code** | Counts the number of cells exhibiting a 'star-like' or highly spiked actin morphology. This phenotype can be associated with specific cytoskeletal disruptors or cellular contraction. |
| 415 | **dapi tubulin center displacement** | **code** | Measures the distance between the centroid of the nucleus (DAPI) and the centroid of the tubulin signal (cell body). Large displacements can indicate cell polarization or asymmetric cytoskeletal collapse. |
| 416 | **actin intensity kurtosis** | **code** | Computes the kurtosis of the Actin channel intensity distribution. High kurtosis indicates a heavy-tailed distribution, potentially reflecting the formation of bright actin stress fibers or aggregates against a lower background. |
| 417 | **actin cortical ratio** | **code** | Measures the ratio of Actin intensity at the cell periphery (cortex) versus the cell center. This quantifies cytoskeletal reorganization, such as cortical ring formation or stress fiber loss. |
| 418 | **actin haralick contrast mean** | **code** | Computes the Haralick Contrast texture feature for the Actin channel. This measures the local intensity variation, capturing the difference between smooth actin staining and distinct stress fibers. |
| 419 | **tubulin texture entropy global** | **code** | Calculates the entropy of the pixel intensity distribution in the Tubulin channel. Higher entropy suggests a more disordered or complex microtubule network, whereas lower entropy might indicate depolymerization or diffuse signal. |
| 420 | **cell density local mean** | **code** | Calculates the mean distance to the nearest k-neighbors for each nucleus. This measures local cell crowding and clustering, which can indicate contact inhibition or colony formation patterns. |
| 421 | **tubulin radial profile slope mean** | **code** | Measures the gradient of Tubulin intensity radiating from the nuclear centroid. A steep drop-off might indicate perinuclear clustering, while a flat profile indicates diffuse distribution. |
| 422 | **actin spatial moment displacement mean** | **code** | Calculates the distance between the geometric centroid of the cell and the weighted centroid (center of mass) of the Actin intensity. This quantifies cell polarity and asymmetric cytoskeletal distribution. |
| 423 | **micronucleus fraction** | **code** | Calculates the ratio of small, detached nuclear objects (micronuclei) to normal-sized nuclei. This is a direct indicator of genomic instability or mitotic slippage mentioned in the knowledge base. |

| | | | |
|---|---|---|---|
| 424 | **nucleus cell centroid distance mean** | **code** | Measures the Euclidean distance between the centroid of the nucleus and the centroid of the cell (cytoplasm). This captures cell polarity and asymmetry, which can be affected by cytoskeletal disruption. |
| 425 | **tubulin fiber texture entropy** | **code** | Calculates the entropy of the texture in the Tubulin (green) channel within cell boundaries. High entropy suggests a disordered microtubule network, while low entropy may indicate stabilization or bundling. |
| 426 | **nucleus cytoplasm centroid distance** | **code** | Calculates the Euclidean distance between the centroid of the nucleus and the centroid of the whole cell (cytoplasm). This measures cell polarity and asymmetry. |
| 427 | **dapi tubulin correlation r** | **code** | Computes the Pearson correlation coefficient between DAPI and Tubulin pixel intensities within the cell. This measures the degree of spatial overlap, identifying phenotypes where tubulin invades the nuclear space or collapses onto it. |
| 428 | **dapi texture gabor mean** | **code** | Applies a bank of Gabor filters to the DAPI channel and calculates the mean response. This captures specific frequency and orientation patterns in chromatin texture that simple statistics might miss. |
| 429 | **cell orientation entropy** | **code** | Measures the entropy of the distribution of cell orientations (major axis angles). High entropy indicates random orientation; low entropy indicates alignment (e.g., due to flow or contact guidance). |
| 430 | **nuclear intensity integrated median** | **code** | Calculates the median integrated intensity (sum of pixel values) per nucleus in the DAPI channel. This correlates with DNA content, helping to distinguish G1 (2N) from G2/M (4N) cells. |
| 431 | **actin area fraction** | **code** | Calculates the percentage of the image covered by Actin signal (above a threshold). This serves as a proxy for cell confluence and spreading, distinct from simple cell count. |
| 432 | **tubulin actin correlation global** | **code** | Computes the Pearson correlation coefficient between pixel intensities of the Tubulin and Actin channels. This measures the spatial overlap and co-regulation of the two cytoskeletal networks. |
| 433 | **nuclear nearest neighbor distance median** | **code** | Calculates the median distance from each nucleus to its nearest neighbor. This quantifies spatial clustering, which can indicate colony formation or contact inhibition effects. |
| 434 | **actin per nucleus ratio** | **code** | Calculates the ratio of total Actin area to the number of nuclei. This provides an average 'cell size' estimate without requiring difficult single-cell segmentation in dense clusters. |
| 435 | **tubulin fiber skeleton length** | **code** | Estimates the total length of microtubule fibers by skeletonizing the Tubulin channel. This is sensitive to the integrity of the microtubule network versus diffuse staining. |
| 436 | **dapi intensity skewness global** | **code** | Measures the skewness of the DAPI intensity histogram. A highly skewed distribution indicates a sparse signal (distinct nuclei on dark background), while lower skewness might indicate background noise or nuclear swelling/diffusion. |
| 437 | **actin texture homogeneity global** | **code** | Computes the GLCM homogeneity of the Actin channel. High homogeneity implies a smooth, uniform actin distribution, while low homogeneity suggests distinct structures like stress fibers or cortical rings. |
| 438 | **tubulin intensity heterogeneity** | **code** | Measures the coefficient of variation (std/mean) of pixel intensities in the Tubulin channel within the cytoplasmic region. High heterogeneity may indicate microtubule bundling or disruption compared to a smooth network. |
| 439 | **nuclear eccentricity variance** | **code** | Computes the variance of nuclear eccentricity across the cell population. High variance suggests a mix of normal and aberrant (e.g., elongated or condensed) nuclear shapes. |
| 440 | **actin perimeter area ratio** | **code** | Calculates the ratio of cell perimeter to cell area based on the Actin channel. High ratios indicate complex, protrusive cell shapes (e.g., filopodia), while low ratios indicate smooth, round shapes. |
| 441 | **dapi entropy** | **code** | Computes the Shannon entropy of the intensity histogram of the DAPI channel. Entropy quantifies the complexity or heterogeneity of nuclear staining patterns. Higher entropy indicates more varied nuclear intensities, potentially reflecting heterogeneous DNA content, chromatin condensation states, or mitotic activity, all of which are relevant when assessing drug effects on MCF-7 cells. |
| 442 | **cell aspect ratio mean** | **code** | Calculates the mean aspect ratio (major axis / minor axis) of segmented cells. This quantifies cell elongation, helping to identify phenotypes like fibroblast-like stretching or rounding up due to mitotic arrest. |
| 443 | **actin radial distribution mean** | **code** | Measures how Actin intensity varies with distance from the nucleus center (e.g., ratio of peripheral to perinuclear intensity). This captures spatial organization phenotypes like cortical actin rings vs. stress fibers. |
| 444 | **nucleus tubulin intensity ratio mean** | **code** | Computes the ratio of Tubulin intensity inside the nucleus region versus the cytoplasm. While Tubulin is cytoplasmic, this metric can detect segmentation errors or specific biological events where the nuclear envelope breaks down. |
| 445 | **dapi bright spot peak count** | **code** | Detects and counts local intensity maxima (peaks) within the nuclear region that exceed a relative threshold. This serves as a proxy for quantifying nuclear foci (e.g., DNA damage repair sites) mentioned in the MoA knowledge. |
| 446 | **actin texture haralick entropy** | **code** | Computes the Haralick entropy of the Actin channel texture. High entropy indicates a complex, disordered cytoskeletal network, while low entropy may suggest diffuse staining or loss of structural integrity. |
| 447 | **tubulin skeleton branch count** | **code** | Skeletons the Tubulin structure and counts the number of branch points. This quantifies the complexity of the microtubule network, distinguishing between dense meshworks and sparse or bundled arrays. |
| 448 | **tubulin spatial radial cv mean** | **code** | Calculates the coefficient of variation of Tubulin intensity in radial bands extending from the nuclear centroid. This specifically targets the 'monoastral spindle' phenotype (Eg5 inhibition), where tubulin is concentrated centrally rather than distributed in a bipolar spindle or network. |
| 449 | **tubulin texture fiber alignment mean** | **code** | Measures the anisotropy or directional coherence of the Tubulin channel using structure tensors or gradient histograms. High alignment scores correspond to the 'microtubule bundling' phenotype (Taxanes), where fibers become rigid and parallel. |
| 450 | **cell nucleus cytoplasm ratio mean** | **code** | Computes the ratio of nuclear area to total cell (cytoplasmic) area. Variations in N/C ratio are indicative of cell cycle stage, differentiation status, or pathological states like multinucleation. |

| | | | |
|---|---|---|---|
| 451 | **channel correlation tubulin actin** | **code** | Calculates the Pearson correlation between Tubulin and Actin channels. This measures the overall integrity and co-organization of the cytoskeleton; disruption by drugs like Vinca alkaloids may uncouple these networks. |
| 452 | **dapi intensity cv inter cell** | **code** | Calculates the coefficient of variation of integrated DAPI intensity *between* cells in a field. High inter-cell variance suggests a heterogeneous population with cells arrested in different cell cycle phases (e.g., G1 vs G2/M block). |
| 453 | **actin spot count mean** | **code** | Counts the number of high-intensity actin spots (e.g., focal adhesions or aggregates) per cell using a Laplacian of Gaussian (LoG) blob detector. An increase may indicate cytoskeletal rigidification or aggregation. |
| 454 | **cell area std dev** | **code** | Calculates the standard deviation of cell areas across the image. A high standard deviation indicates a heterogeneous population, potentially containing a mix of giant senescent cells and small apoptotic fragments. |
| 455 | **actin skeleton branch points** | **code** | Skeletons the Actin channel intensity and counts the number of branch points per cell. This quantifies the complexity and interconnectivity of the actin network. |
| 456 | **nuclear integrated intensity sum** | **code** | Sums the total integrated intensity of DAPI across all nuclei in the image. This serves as a proxy for total DNA content, which can reflect cell count (confluence) as well as ploidy changes if normalized by cell number. |
| 457 | **tubulin fiber alignment** | **code** | Measures the directional alignment of tubulin fibers in the green channel using structure tensor analysis or gradient orientation. High alignment may indicate microtubule bundling (e.g., Taxol effect), while disorder suggests depolymerization. |
| 458 | **cell count normalized** | **code** | Counts the number of segmented nuclei and normalizes by the image area. This provides a measure of cell density, which is a direct proxy for cytotoxicity or cytostatic effects of the drugs. |
| 459 | **actin perimeter mean** | **code** | Calculates the mean perimeter of cell boundaries defined by Actin. An increase in perimeter relative to area (fractal dimension) can indicate membrane ruffling or the formation of protrusions like invadopodia. |
| 460 | **nucleus area median** | **code** | Calculates the median area of segmented nuclei based on the DAPI channel. Changes in nuclear size are key indicators of cellular states such as senescence (enlarged) or apoptosis (pyknosis/shrinkage). |
| 461 | **nucleus chromatin texture entropy** | **code** | Measures the entropy of pixel intensities within segmented nuclei in the DAPI channel. High entropy indicates heterogeneous chromatin texture, which correlates with DNA damage or condensation stages. |
| 462 | **tubulin radial intensity decay** | **code** | Measures the rate of decay of Tubulin intensity as a function of distance from the nucleus center. This captures the spatial distribution of microtubules, relevant for detecting phenotypes like monoastral spindles where tubulin is centrally concentrated. |
| 463 | **actin cell eccentricity median** | **code** | Calculates the median eccentricity (elongation) of cell boundaries defined by the Actin channel. This feature captures gross morphological changes such as the transition from epithelial to mesenchymal-like shapes or rounding due to mitotic arrest. |
| 464 | **actin intensity skewness mean** | **code** | Computes the skewness of the actin intensity distribution within segmented cells. This helps distinguish between cells with uniform actin staining and those with bright cortical rings or localized stress fibers. |
| 465 | **cell packing density local** | **code** | Estimates the local density of cells by calculating the mean distance to the nearest neighbor for each nucleus. This feature relates to cell proliferation rates and contact inhibition. |
| 466 | **cytoplasm to nucleus area ratio** | **code** | Calculates the ratio of cytoplasmic area (Cell area minus Nucleus area) to Nuclear area. This N:C ratio is a classic cytological metric that changes with cell cycle progression and senescence. |
| 467 | **actin tubulin overlap manders** | **code** | Computes the Manders' Overlap Coefficient between Actin (red) and Tubulin (green) channels within the cell mask. This quantifies the spatial relationship between the two major cytoskeletal components. |
