## Supplementary feature list 2 for "Biologically grounded cell profiling across microscopy modalities"

### MorphAgent HSC feature catalog

25 MorphAgent mitochondrial features for HSC age-state profiling. Method: vlm or code.

| # | Feature name | Method | Description |
|---|---|---|---|
| 1 | **mitochondria background signal heterogeneity** | **code** | Measures the standard deviation of intensities in the background region (pixels not covered by any mitochondrial label) after masking out the cell area if available, yielding a scalar that reflects residual background noise and possible diffuse mitochondrial signal. Higher values indicate more heterogeneous non-organelle signal, which might correspond to cytosolic mitochondrial remnants or segmentation under/over-calls. This feature is useful to quantify image quality and subtle diffuse mitochondrial morphology not captured by object masks. |
| 2 | **mito perimeter intensity gradient mean** | **code** | For each mitochondrial instance, samples intensities just inside and just outside its perimeter (e.g., one-pixel inward and outward normal directions) and computes the average inward-minus-outward intensity difference across all boundary pixels, then averages this value across objects weighted by perimeter length. This feature measures the typical sharpness of the mitochondrial signal edge relative to local background, capturing whether mitochondria have crisp boundaries or more diffuse, low-contrast edges in label-free imaging. It is relevant because sharper, high-contrast organelle boundaries may reflect distinct refractive index differences or packing states associated with particular metabolic or stress conditions. |
| 3 | **background mito signal to noise** | **code** | Estimates a signal-to-noise ratio by computing the mean mitochondrial intensity inside the cell mask (union of labels) and dividing by the standard deviation of intensities in a surrounding background ring (e.g., a 20-pixel dilation of the cell mask minus the mask). This quantifies how cleanly the mitochondrial signal stands out from retained background noise in each label-free image. Variations in this ratio per cell can reflect differences in imaging quality or intrinsic scattering properties that should be accounted for in downstream models. |
| 4 | **mitochondria neighbor intensity autocorrelation** | **code** | Computes a local Moran's I or similar spatial autocorrelation statistic on mitochondrial intensities within the cell mask, using 8-connected pixel neighborhoods, and returns the resulting scalar. Higher positive values indicate clustered high-intensity mitochondria regions, while values near zero indicate random or checkerboard-like patterns. This feature captures mesoscale patchiness of mitochondrial activity, which may relate to localized ROS hotspots and autophagic activity in HSCs. |
| 5 | **mitochondria texture anisotropy glcm** | **code** | Captures directional organization of mitochondrial texture by computing gray-level co-occurrence matrix (GLCM) features at multiple orientations (e.g., 0°, 45°, 90°, 135°) and returning the coefficient of variation of GLCM contrast across angles. High anisotropy indicates that intensity patterns are more structured along specific directions (e.g., aligned tubules), whereas low values indicate isotropic, punctate distributions, which may distinguish different mitochondrial architectures in HSCs. |
| 6 | **mitochondria long tubule fraction** | **code** | Calculates the fraction of mitochondrial objects whose length scale exceeds a high-length threshold, defined for each object as the square root of its area or its major-axis length in pixels. This yields the proportion of the mitochondrial population forming long tubules rather than small fragments, providing a scalar measure of network fusion that complements average aspect ratio and count. |
| 7 | **mitochondrial area fraction** | **code** | Measures the fraction of the 512x512 image area occupied by mitochondrial signal, defined as the total number of pixels assigned to non-zero labels in the mitochondria label map divided by the total image pixel count. This reflects how much of the cell area is filled with mitochondria and can distinguish metabolically active cells with dense networks from cells with sparse mitochondria. |
| 8 | **mitochondria local autocorrelation length** | **code** | Computes the 2D spatial autocorrelation function of mitochondrial intensity within the cell and estimates the correlation length as the radius at which autocorrelation decays to 1/e of its maximum. This length scale summarizes the typical size of coherent mitochondrial intensity patches, distinguishing fine punctate patterns from extended networks. It is relevant as a scale-aware measure of mitochondrial organization linked to stem cell metabolic wiring. |
| 9 | **mito intensity autocorrelation length** | **code** | Computes the 2D spatial autocorrelation function of the label-free intensity restricted to mitochondrial pixels and estimates the characteristic correlation length (e.g., the distance at which the autocorrelation decays to 1/e of its maximum). A longer autocorrelation length indicates larger, smoother patches of similar intensity, while a shorter length reflects fine-grained intensity texture within mitochondria. This is relevant for summarizing whether mitochondrial-related signal is organized into large homogeneous domains or highly speckled patterns, which may map to differences in ultrastructural organization or metabolic heterogeneity. |
| 10 | **mitochondria to cell intensity dynamic range ratio** | **code** | Computes the dynamic range (e.g., 95th minus 5th percentile) of pixel intensities within mitochondrial objects and divides it by the dynamic range in the entire cell mask. This normalizes mitochondrial contrast to overall cell-level contrast, quantifying how strongly the mitochondria-related morphology channel stands out relative to residual background and other structures. A higher ratio suggests more sharply defined mitochondrial structures, potentially reflecting stronger labeling of active mitochondrial regions in these label-free HSC images. |
| 11 | **background noise robust snr** | **code** | This feature estimates a robust signal-to-noise ratio by taking the median intensity of mitochondrial pixels (those with nonzero labels) as signal and the median absolute deviation (MAD) of intensities in background pixels (cell mask complement) as noise, then forming their ratio. It quantifies how strongly the mitochondrial morphology signal stands out above retained background noise while being robust to outliers. This reflects the overall quality and contrast of the label-free mitochondrial signal per cell. |
| 12 | **mito intensity geodesic variation** | **code** | Constructs a graph where mitochondrial pixels are connected if they are neighbors within the same instance, computes the geodesic distance (shortest path length) along this graph from the brightest mitochondrial pixel to all other mitochondrial pixels, and then calculates the coefficient of variation of intensities as a function of geodesic distance. The final scalar is the slope of a linear fit of intensity CV versus distance. A strongly positive slope indicates that intensity becomes more heterogeneous along mitochondrial branches from bright hubs, potentially reflecting localized hotspots of mitochondrial activity, whereas a flat slope suggests more uniform internal signal. This is relevant for capturing intra-network heterogeneity in the label-free mitochondrial morphology signal. |

| | | | |
|---|---|---|---|
| 13 | **mitochondria local entropy mean** | **code** | Applies a local entropy filter (e.g., Shannon entropy in a sliding window) to the mitochondrial intensity image within the cell mask, then averages the entropy values over all cell pixels. This summarizes the typical local complexity and texture of mitochondrial signal: higher entropy reflects fine-grained, irregular patterns (e.g., many small fragments), while lower entropy suggests smoother, more uniform structures (e.g., extended tubules). It offers a texture-based view of mitochondrial remodeling beyond simple object counts. |
| 14 | **mitochondrial radial anisotropy index** | **code** | Within the segmented cell, converts pixel coordinates to polar (radius, angle) around the cell centroid and constructs a radial mitochondrial intensity profile along multiple angular sectors. It then computes the coefficient of variation across sector-wise radial profiles, quantifying directional anisotropy of the mitochondrial distribution (e.g., aligned along one axis vs. isotropic). Higher anisotropy can reveal polarized mitochondrial deployment associated with asymmetric stem cell fate decisions. |
| 15 | **mitochondria network continuity score** | **code** | Quantifies how network-like versus fragmented the mitochondrial morphology is within a cell. High values indicate long, continuous tubules and branched interconnected networks; low values indicate mostly isolated, round or punctate mitochondrial objects. |
| 16 | **mitochondria length width skewness** | **code** | For each mitochondria instance in the label map, this feature computes the major/minor axis ratio (length/width) and then takes the skewness of that per-object distribution over the whole cell. It quantifies whether the population of mitochondria is biased toward elongated (high axis ratio) or spherical shapes, beyond simple mean aspect ratio. A strongly positive skew indicates a minority of highly elongated mitochondria on a background of mostly round objects, which may reflect specific mitochondrial remodeling states. |
| 17 | **mitochondria intensity entropy** | **code** | Computes the Shannon entropy of the mitochondrial intensity histogram within the cell mask. Higher entropy indicates a broad, heterogeneous range of intensities (potentially corresponding to mixed mitochondrial states), while lower entropy indicates a more uniform mitochondrial signal. This captures global heterogeneity in the label-free mitochondrial morphology signal beyond simple mean or variance. |
| 18 | **mito clustered vs isolated object fraction** | **code** | Identifies mitochondrial instances that are in spatial clusters by dilating each object slightly and counting how many have another instance within a small distance versus those that remain isolated; the feature is the fraction of total mitochondrial area contributed by clustered objects. This captures whether mitochondria tend to form spatially clustered aggregates versus being dispersed as isolated fragments throughout the cell. Such clustering can reflect organelle crowding or localized metabolic hubs, which may be relevant for distinguishing different HSC states. |
| 19 | **mitochondria intensity gini inside cell** | **code** | This feature computes the Gini coefficient of mitochondrial signal intensities for all pixels within the cell mask (excluding background). It quantifies how uneven the mitochondrial-related label-free signal is across the cell: a high Gini indicates a few very bright mitochondrial regions amid dim background, whereas a low Gini indicates more homogeneous signal. Intensity heterogeneity can reflect heterogeneous mitochondrial mass, membrane potential, or ROS within single HSCs. |
| 20 | **mitochondria area fraction in perinuclear band** | **code** | Measures the fraction of total mitochondrial area that resides in a fixed-width perinuclear band around the cell’s nucleus. A nucleus mask (from an appropriate channel or prior segmentation) defines the band (e.g., 2–5 µm radius), and the overlap of mitochondrial labels with this band is summed and divided by total mitochondrial area. This captures how strongly mitochondria are concentrated around the nucleus versus more peripheral, reflecting shifts in metabolic organization. |
| 21 | **mitochondria intensity entropy within cell** | **code** | Computes the Shannon entropy of the intensity histogram for pixels inside the mitochondrial region of the cell, using a fixed number of bins (e.g., 64). Higher entropy indicates a broader, more heterogeneous distribution of mitochondrial intensities, potentially reflecting mixed populations of highly and weakly active mitochondria, while lower entropy suggests more uniform signal. This feature captures global intensity heterogeneity tied to mitochondrial functional states. |
| 22 | **mitochondrial clusteredness by nearest neighbor** | **code** | Quantifies how spatially clustered mitochondrial objects are within the cell by comparing mean nearest-neighbor distances between object centroids to a uniform spatial null. Higher values indicate stronger intra-cell mitochondrial clusteredness. |
| 23 | **mitochondrial network fragmentation score** | **code** | Summarizes mitochondrial fragmentation on a continuum from fused/network-like to punctate. Low scores correspond to long, continuous, branched tubules; high scores correspond to fragmented, punctate mitochondria. |
| 24 | **mitochondrial network compactness** | **vlm** | Visually scores how compact versus dispersed the mitochondrial network appears within the cell. Higher values indicate a more compact, centrally concentrated mitochondrial organization; lower values indicate a more dispersed network. |
| 25 | **mitochondrial neighbor intensity variogram slope** | **code** | Estimates the slope of the empirical semi-variogram of mitochondrial intensity between neighboring pixels at increasing spatial lags within the cell. Captures how quickly intensity similarity decays with distance. |
