## Supplementary feature list 3 for "Biologically grounded cell profiling across microscopy modalities"

### MorphAgent Tau feature catalog

301 MorphAgent features for Tau SIM imaging of SH-SY5Y cells. Method: vlm or code.

| # | Feature name | Method | Description |
|---|---|---|---|
| 1 | **tau filament fragmentation score** | **code** | Calculates the number of disjoint skeletal segments in 'mask_filament' normalized by the total skeletal length. A higher score indicates fragmented, discontinuous neurites typical of cytoskeletal breakdown. |
| 2 | **tau filament texture anisotropy** | **code** | This feature captures directional texture within filament regions by computing the ratio of GLCM-based contrast along the dominant filament orientation versus the perpendicular orientation, restricted to mask_filament.tif. For each cell, a local structure tensor or orientation histogram within filament pixels yields a dominant angle; gray-level co-occurrence matrices are then computed along this angle and its orthogonal counterpart to estimate directional contrast, and their ratio defines the anisotropy. High anisotropy suggests well-aligned, directionally coherent filaments, while low anisotropy reflects more isotropic, disorganized filament texture. |
| 3 | **tau soma granularity scale ratio** | **code** | This feature measures the ratio of fine- to coarse-scale granularity of Tau within the soma by computing wavelet- or Laplacian-of-Gaussian-based granularity energy at two spatial scales (e.g., small sigma vs. large sigma) restricted to the mask_cell minus mask_nucleus region, and taking the small-scale energy divided by large-scale energy. On the 2D MIP, a low ratio indicates dominance of larger, NFT-like aggregates, whereas a high ratio reflects predominantly fine puncta or small condensates. It is intended to distinguish early granular Tau pathology from late-stage large inclusions. |
| 4 | **tau aggregation scale median radius** | **code** | Across all Tau-positive objects derived from the union of droplet, filament, and bundle masks, this feature characterizes the dominant physical scale of Tau aggregates. Each connected object is approximated by an equivalent-area disk and its radius is computed; the feature is the median of these radii. This provides a robust measure of whether Tau pathology in a cell is dominated by nanoscale puncta, intermediate droplets, or large inclusion bodies. |
| 5 | **tau intensity cell vs extracellular ratio** | **code** | This feature computes the ratio between the mean Tau intensity inside the cell mask (mask_cell.tif) and the mean Tau intensity in the area outside the cell but within the image field. It quantifies how strongly Tau signal is confined to the labeled neuron versus diffuse or extracellular/background signal, which is relevant for distinguishing true intracellular pathology from nonspecific fluorescence or extracellular deposits. A higher ratio suggests more cell-confined Tau, whereas a lower ratio may indicate extracellular Tau or ghost-like tangles outside the cell mask. |
| 6 | **tau cell body enrichment index** | **code** | This feature measures the enrichment of Tau fluorescence in the whole cell body relative to the surrounding background by computing the ratio of mean Tau intensity inside the cell mask (mask_cell.tif) to the mean intensity in a thin ring just outside the cell. It captures whether the neuron shows globally elevated Tau signal within the soma and proximal processes, which is a hallmark of pathological mis-localization away from axon-only dominance. A higher value indicates a stronger global Tau accumulation in the patient cell. |
| 7 | **tau cell to background intensity ratio** | **code** | This feature computes the mean Tau fluorescence intensity inside the cell mask divided by the mean intensity outside the cell mask (background) in the 2D MIP. It quantifies how strongly Tau is enriched in the patient neuron relative to its local background, providing a basic measure of overall Tau burden per cell. Higher values may indicate globally elevated Tau accumulation in the cell. |
| 8 | **tau droplet intensity texture contrast** | **code** | Inside each droplet in mask_droplet.tif, this feature computes a local gray-level co-occurrence matrix (GLCM) contrast on the Tau MIP and then averages the contrast across droplets. Higher contrast indicates more internal intensity heterogeneity (e.g., core–shell architecture or sub-structures), whereas low contrast corresponds to homogeneous, liquid-like droplets. This captures subtle maturation of Tau condensates into heterogeneous aggregates. |
| 9 | **nuc background ratio** | **code** | Fluorescence intensity ratio of nucleus to background. Background is defined as the region outside the whole cell mask. This ratio indicates nuclear Tau enrichment relative to the image background. |
| 10 | **axon like filament linear intensity gradient** | **code** | For each filament object inside the cell mask, this feature extracts its skeleton, parameterizes the longest geodesic path, samples Tau intensity along this path, and fits a linear model of intensity versus path distance. The per-filament slopes are then averaged to yield an overall linear gradient index, capturing whether Tau intensity tends to decline along filaments (healthy axon-like behavior) or becomes flat/irregular (disrupted distal–proximal gradient in pathology). More negative mean slopes correspond to preserved proximal-to-distal decay of Tau signal. |
| 11 | **tau droplet texture contrast** | **code** | This feature measures fine-scale heterogeneity of Tau within droplet regions by computing a gray-level co-occurrence matrix (GLCM) contrast over the Tau intensities restricted to mask_droplet.tif. For each droplet pixel neighborhood, GLCM contrast is computed at a chosen distance and orientation, and the mean contrast over all droplet pixels is reported. Higher values reflect sharper local intensity changes and more internally structured droplets, potentially indicating maturation into fibrillar aggregates. |
| 12 | **tau cell to background dynamic range** | **code** | This feature computes, at the image level, the ratio between the 95th percentile Tau intensity inside all cell masks (mask_cell) and the 95th percentile Tau intensity in background regions outside any cell mask. It reflects how much brighter Tau-positive structures are relative to the local background, integrating effects of both expression level and aggregation contrast. Reduced dynamic range could indicate diffuse, low-contrast Tau or imaging issues, whereas increased range corresponds to strong, localized Tau aggregates visible in super-resolution microscopy. |
| 13 | **tau intra cell intensity skewness** | **code** | This feature computes the skewness of the Tau intensity distribution within the cell mask (mask_cell), capturing how heavy-tailed the per-cell intensity histogram is. High positive skewness reflects a small number of very bright pixels on a dimmer background, consistent with focal aggregates, whereas low skewness corresponds to more uniform Tau distribution. It is independent of absolute expression level and focuses on distribution shape. |

| | | | |
|---|---|---|---|
| 14 | **tau droplet internal texture entropy** | **code** | Within each droplet (mask_droplet), this feature computes a gray-level co-occurrence matrix (GLCM) entropy on the Tau channel pixels (e.g., using a fixed offset and quantization), then averages this entropy over droplets within the cell. It reflects how structurally homogeneous or heterogeneous each Tau droplet's interior is: low entropy corresponds to uniform, solid droplets, whereas high entropy indicates internal substructure or nanoscale heterogeneity projected into the MIP. The scalar output is the mean GLCM entropy across droplets. |
| 15 | **tau radial decay from cell centroid** | **code** | This feature summarizes how Tau intensity decays with distance from the cell centroid by fitting a linear regression of mean Tau intensity in concentric radial bins versus radial distance within the cell mask and taking the regression slope. A steep negative slope suggests soma-centric Tau concentration that fades toward neurites, while a flatter or positive slope indicates relatively uniform or distal-enriched Tau distribution, relevant to assessing redistribution of Tau from axon-like processes back toward the soma. |
| 16 | **tau bundle filament intensity contrast** | **code** | Within the bundle mask (mask_bundle.tif), this feature computes the mean Tau intensity in bundle pixels and the mean Tau intensity in adjacent filament pixels (mask_filament.tif overlapping cell but outside bundles), then takes their ratio. It quantifies how much brighter bundled fibrillar aggregates are compared with surrounding individual filaments on the 2D MIP, a proxy for higher-order Tau packing. |
| 17 | **tau cell edge to nucleus intensity gradient** | **code** | Within the cell mask, this feature computes the radial gradient of Tau intensity from the cell edge toward the nuclear boundary by averaging Tau intensity in concentric normalized distance bands between the cell perimeter and nuclear mask, then fitting a line to intensity versus normalized distance. The slope of this line quantifies whether Tau tends to be enriched at the cortex/neurites (negative slope) or near the perinuclear region (positive slope). This captures coarse Tau relocalization from axonal/neuritic regions toward the soma and perinuclear area in pathological states. |
| 18 | **tau cell cytoplasmic texture entropy** | **code** | Within each cell's cytoplasmic region (mask_cell minus mask_nucleus), this feature computes the gray-level co-occurrence matrix (GLCM) entropy of Tau intensities at a fixed pixel offset and direction, then averages entropy across cells. It characterizes how heterogeneous and 'granular' the cytoplasmic Tau distribution is in the 2D MIP, with higher entropy reflecting more complex, speckled patterns typical of diffuse aggregates and mixed fibrillar/clustered states. Lower entropy corresponds to smoother, more uniform Tau distributions. |
| 19 | **tau cellwide granularity ratio** | **code** | This feature applies a multiscale spot-enhancing filter (e.g., Laplacian of Gaussian at two Tau-relevant scales) to the Tau MIP within the cell mask and computes the ratio of total LoG response at a small scale (puncta/nanocluster-sized) to that at a larger scale (droplet-sized). It quantifies whether Tau signal is dominated by fine punctate granules versus larger aggregates across the whole cell. Shifts in this ratio can reflect progression from diffuse/oligomeric Tau toward larger inclusion bodies. |
| 20 | **cellwise tau aggresome compactness** | **code** | For each cell, this feature measures how compact the largest Tau aggregate is by computing the ratio of the area of its convex hull to its actual area using high-intensity Tau objects within mask_cell.tif. First, an adaptive threshold on Tau intensity inside each cell mask identifies aggregate objects; the largest object per cell is selected, and its convex hull area divided by its pixel area is computed, then averaged across cells. Values close to 1 indicate a compact, convex aggresome-like inclusion, whereas higher values indicate more irregular, ramified aggregates. |
| 21 | **tau neurite radial decay irregularity** | **code** | For each cell, compute the distance transform from the nucleus center over the cell mask and sample Tau intensity along multiple radial lines extending into neuritic regions, then fit a smooth decay model (e.g., exponential) and quantify the residual variance normalized by the model fit. This feature quantifies how irregular the radial decay of Tau intensity is from soma toward neurites, detecting disruptions of the normal axon-dominant gradient expected for non-pathological Tau. |
| 22 | **tau pathological mislocalization score** | **vlm** | This feature is a visual scoring of how strongly Tau appears mislocalized from thin axon-like processes into the soma and thicker dendritic processes in the 2D Tau MIP for each segmented neuron. It should reflect the expert notion of pathology by jointly considering the relative brightness and presence of compact aggregates in the soma and dendrites versus more physiological, fine linear signal along axon-like processes. |
| 23 | **tau cellwide texture entropy in tau positive region** | **code** | Restricting to Tau-positive pixels within the cell mask (e.g., above a low percentile threshold of Tau intensity), this feature computes gray-level co-occurrence matrix (GLCM) entropy on the 2D MIP. Higher entropy reflects more complex, heterogeneous Tau texture (mixture of diffuse, filamentous, and droplet patterns), whereas lower entropy indicates more uniform distributions, helping distinguish structured aggregates from homogeneous background. |
| 24 | **bundle width** | **code** | Width of the thickest microtubule bundle in pixels. Computed from the bundle mask (mask_bundle.tif) by analyzing connected components, computing the distance transform for each component, and taking twice the maximum distance transform value as the width of the thickest bundle. |
| 25 | **intra cell tau texture entropy contrast** | **code** | Within the cell mask on the Tau MIP, compute a gray-level co-occurrence matrix (GLCM) at a fixed pixel distance (e.g., 3–5 pixels) and orientation-averaged, then derive GLCM entropy and contrast as texture measures of Tau distribution. Subtract the entropy measured within the nucleus mask from the entropy within the cytoplasmic cell region (cell minus nucleus) to obtain an entropy contrast value. This feature emphasizes how much more texturally complex Tau patterns are in the cytoplasm versus near the nuclear region, which is relevant to distinguishing diffuse vs granular Tau pathology. |
| 26 | **tau intensity entropy within cell** | **code** | Restricted to the cell mask, this feature computes the Shannon entropy of the Tau intensity histogram on the 2D MIP. It reflects the diversity of intensity levels inside the neuron: lower entropy corresponds to relatively homogeneous staining, whereas higher entropy indicates coexistence of dark regions, intermediate diffuse signal, and very bright aggregates, as expected in heterogeneous Tau pathology. |
| 27 | **single filament cyto ratio** | **code** | Fluorescence intensity ratio of single filament Tau intensity to cytoplasmic Tau intensity. This measures the enrichment of Tau on individual microtubule filaments relative to the diffuse cytoplasmic pool. |
| 28 | **tau soma pathology staging score** | **vlm** | For each neuron, visually inspect the Tau channel within the soma to assign a scalar pathology stage that integrates multiple cues: overall soma brightness, presence and size of discrete Tau inclusions, and textural coarse-graining (from diffuse to coarse granular to dense inclusions). The score should increase from near-normal diffuse low Tau to severe pathology with large, dense soma aggregates, providing an interpretable staging of soma Tau pathology from a single number. |

| | | | |
|---|---|---|---|
| 29 | **tau cellwide intensity entropy** | **code** | This feature computes the Shannon entropy of the Tau intensity histogram within the cell mask (mask_cell) in the 2D MIP. It reflects the complexity and spread of Tau intensity levels across the cell: low entropy suggests a relatively uniform low or high signal, while high entropy suggests a mix of background, intermediate, and bright aggregates, characteristic of heterogeneous aggregation states. |
| 30 | **tau cell compartment mislocalization score** | **code** | This feature scores Tau mislocalization by combining three normalized ratios: soma-to-cell mean intensity, droplets-to-cell mean intensity, and filaments-to-cell mean intensity, each z-scored across the dataset and summed for each cell. It yields a single scalar that increases when Tau is shifted from uniformly thin filaments into soma-dense, droplet-rich patterns, which are characteristic of pathological redistribution in tauopathy. It leverages all provided masks (cell, droplet, filament) when present. |
| 31 | **tau soma radial distribution skewness** | **code** | Within the soma region (mask_cell minus mask_filament and mask_droplet, approximating the compact cell body), compute the radial distance of each pixel from the soma centroid and quantify the skewness of Tau intensity as a function of radius by fitting a radial histogram (intensity-weighted). This scalar measures whether Tau is biased toward the periphery versus the center of the soma, potentially reflecting perinuclear versus cortical accumulation patterns of Tau inclusions in 2D MIPs. |
| 32 | **tau intensity cell to droplet enrichment ratio** | **code** | This feature computes the ratio between the mean Tau intensity inside all droplet masks and the mean Tau intensity in the remaining cell area (cell mask minus droplet mask) on the 2D MIP. It quantifies how enriched Tau is in punctate droplet-like aggregates relative to the diffuse cytoplasmic pool, which is relevant for distinguishing cells with strong sequestration of Tau into discrete inclusions from those with more homogeneous distribution. |
| 33 | **tau cellwide granularity scale ratio** | **code** | On the Tau MIP restricted to mask_cell, this feature computes multiscale granularity by convolving with Gaussian or ring filters at two biologically motivated scales (e.g., small scale ~1–2 pixels for fine puncta vs larger scale ~4–8 pixels for droplet/bundle-scale structures) and taking the ratio of their mean filter responses. This scalar captures the balance between fine-grained punctate Tau versus larger aggregates within the cell. A high ratio indicates dominance of larger-scale granules or bundles over fine puncta, consistent with aggregation progression. |
| 34 | **tau droplet enrichment ratio** | **code** | Calculates the ratio of mean Tau intensity inside mask_droplet.tif to the mean intensity of the surrounding cytoplasm (mask_cell.tif excluding droplets). This measures the partition coefficient of Tau into phase-separated condensates. |
| 35 | **tau droplet intensity enrichment** | **code** | Calculates the ratio of mean intensity inside `mask_droplet.tif` regions relative to the mean intensity of the rest of the cell (`mask_cell.tif` excluding droplets). Measures how concentrated Tau is within these phase-separated droplets. |
| 36 | **tau droplet intensity contrast** | **code** | Measures the contrast of droplets relative to the immediate background (local cell cytoplasm). High contrast indicates dense, compact aggregates rather than diffuse pooling. |
| 37 | **intranuclear tau texture entropy** | **code** | This feature quantifies the textural complexity of Tau signal inside nuclei by computing the gray-level co-occurrence matrix (GLCM) entropy within mask_nucleus.tif on the Tau MIP and averaging across all nuclei. After masking Tau intensities to nuclear regions, a GLCM is computed at a fixed pixel offset and angle, and the entropy of the normalized co-occurrence distribution is used as the per-nucleus measure. Elevated entropy reflects more heterogeneous, punctate, or structured intranuclear Tau, which could indicate aberrant nuclear Tau interactions. |
| 38 | **tau intra cell texture entropy** | **code** | This feature calculates the gray-level co-occurrence matrix (GLCM) entropy of the Tau channel restricted to the cell mask, averaging entropy over multiple orientations and a defined pixel offset. It characterizes the local intensity texture within the neuron, distinguishing smooth diffuse Tau distributions from highly heterogeneous, granular or honeycomb-like patterns. Higher entropy suggests more complex, disordered Tau organization associated with pathological aggregation. |
| 39 | **bundle filament alignment coherence** | **code** | Within the union of filament and bundle masks (mask_filament.tif ∪ mask_bundle.tif), estimate local orientation at each pixel using a structure tensor or similar method, and compute the mean resultant vector length of orientations restricted to the bundle mask versus the filament-only regions. The feature is defined as coherence_bundle − coherence_filament, where coherence is the circular resultant length. A higher value indicates that thick bundles contain more uniformly aligned Tau compared to thinner filaments, reflecting more ordered, NFT-like fibrillar tracts. |
| 40 | **tau texture in cell body entropy** | **code** | This feature computes the gray-level co-occurrence matrix (GLCM) entropy of the Tau intensity within the cell mask, using several pixel offsets averaged together. It quantifies the complexity and granularity of the Tau distribution in the soma, where higher entropy corresponds to heterogeneous, punctate or tangle-like patterns, and lower entropy corresponds to smoother, diffusely distributed Tau. |
| 41 | **tau cell to background intensity contrast** | **code** | This feature computes the ratio between the mean Tau intensity inside the cell mask and the mean Tau intensity in an annular background region immediately surrounding the cell on the 2D MIP. It reflects how strongly Tau is enriched in the patient neuron relative to its local background, independent of absolute exposure settings, and can indicate overall Tau accumulation within the cell. |
| 42 | **tau droplet shape anisotropy index** | **code** | For all droplet objects in mask_droplet, this feature computes the mean eccentricity (or major-to-minor axis ratio) across droplets, weighted by droplet area. It quantifies whether Tau droplets are predominantly round (low anisotropy) or elongated/rod-like (high anisotropy), thereby distinguishing spherical inclusions from more fibrillar or rod-like condensates. This could relate to different Tau aggregation states or maturation stages. |
| 43 | **tau soma vs neurite dominance score** | **vlm** | This feature uses a model to score, on a continuous scale, whether Tau signal visually appears predominantly localized in the soma versus in the neurites (axonal/dendritic processes) in the MIP. It captures a global impression of Tau redistribution from axon-dominant to soma/dendrite-dominant patterns, integrating intensity, area, and morphology that may not be fully summarized by simple ratios. A higher score indicates stronger soma-dominant Tau accumulation, consistent with advanced Tau pathology. |
| 44 | **tau filament radial redistribution index** | **code** | This feature measures how Tau filament intensity redistributes from perinuclear to peripheral regions by computing the fraction of filament intensity within an inner ring (near the nucleus mask) versus an outer ring (near the cell boundary). A higher value indicates Tau filaments concentrated near the cell center, while a lower value indicates redistribution toward distal neurites. |

| | | | |
|---|---|---|---|
| 45 | **filament orientation coherence** | **code** | Analyzes the orientation of structures in 'mask_filament' using structure tensor analysis or gradient orientation. It outputs a scalar measuring how aligned the filaments are. Healthy microtubules are often highly aligned in neurites, while tangles may be disorganized. |
| 46 | **tau filament alignment within cell** | **code** | Within the cell mask, compute the principal orientation of all filament objects from mask_filament.tif (e.g., via structure tensor or regionprops orientation on their skeleton) and calculate the magnitude of the mean resultant vector of these orientations. This yields a scalar between 0 and 1 that quantifies how aligned Tau-positive filaments are: values near 1 indicate a preferred common direction, while lower values indicate disordered or tangled filament orientations. |
| 47 | **tau filament linear fraction** | **code** | Within the Tau-positive filament mask, this feature estimates the fraction of filament area that is part of locally line-like, high-aspect-ratio structures versus bulkier, blob-like regions. It can be implemented by skeletonizing mask_filament.tif, reconstructing a band around the skeleton, and computing the area of Tau signal within this band divided by the total Tau-positive filament area. Lower values suggest loss of smooth, linear neuritic Tau and emergence of thicker aggregates along the neurites. |
| 48 | **bundle filament alignment score** | **code** | This feature measures how well individual filament objects are locally aligned with nearby bundle structures, quantifying the degree to which filaments are organized into higher-order bundles. For each filament object in mask_filament.tif, its main orientation (from regionprops) is compared to the local orientation of the nearest bundle object in mask_bundle.tif, and the average absolute cosine of orientation differences is computed across filaments. Higher values indicate more coherent alignment of filaments with bundles, reflecting more ordered fibrillar architecture. |
| 49 | **tau perisomatic enrichment index** | **code** | This feature quantifies Tau enrichment in a narrow perisomatic ring by computing the mean Tau intensity in a band of fixed pixel width just outside the soma boundary (dilated mask_cell minus mask_cell) and dividing it by the mean intensity in more distal neuritic regions (cell mask minus the dilated soma). It captures whether Tau accumulates preferentially near the soma–neurite interface, which can correspond to early mislocalization or proximal axonal pathology. The band width is defined in pixels but can be mapped to microns if pixel size is known. |
| 50 | **tau droplet intensity compaction score** | **code** | For each Tau droplet in mask_droplet.tif, this feature computes the ratio of its 90th percentile Tau intensity to its 50th percentile intensity and then averages this ratio across all droplets. A higher score indicates that droplets have a bright, compact core relative to their median signal, consistent with more condensed pathological Tau inclusions. |
| 51 | **tau bundle soma overlap fraction** | **code** | This feature computes, for each cell, the fraction of total bundle area (from mask_bundle.tif) that lies within the cell body region defined by mask_cell minus mask_nucleus. It summarizes how much thick Tau bundle signal is confined to the soma versus neurites, reflecting soma-centered neurofibrillary tangle-like pathology. Values near 1 indicate bundles largely restricted to the soma. |
| 52 | **tau bundle cytoplasmic infiltration ratio** | **code** | This feature measures the fraction of total Tau bundle area (mask_bundle) that lies outside the nucleus but inside the cell, divided by the bundle area inside putative neuritic regions (mask_filament). It captures whether large Tau bundles are mislocalized into the soma/cytoplasm versus remaining confined to neuritic filaments. Higher values indicate greater somatic/cytoplasmic infiltration of bundled Tau. |
| 53 | **tau mislocalization score** | **vlm** | This feature is a visual score of how severely Tau is mislocalized from axonal-like thin processes into the cell body and thicker dendrite-like processes in the 2D MIP of the Tau channel. It should reflect the qualitative shift from an axon-dominant, thin linear pattern to strong soma-centered and thick neurite Tau accumulation, which is characteristic of pathological Tau redistribution. |
| 54 | **tau inside vs outside cell intensity ratio** | **code** | This feature computes the ratio between the mean Tau intensity inside the cell mask and the mean Tau intensity in a thin ring just outside the cell mask, both on the 2D Tau MIP. It captures how strongly Tau is retained within the segmented neuron versus dispersed in the local extracellular/background space, which is relevant for distinguishing compact intracellular pathology from diffuse leakage or ghost-like aggregates. |
| 55 | **tau pathology severity score** | **vlm** | A visual assessment score (1-5) of the overall Tau pathology, considering factors like somatic brightness, neurite fragmentation, and aggregate presence. This integrates multiple visual cues into a single severity metric. |
| 56 | **tau bundle alignment coherence** | **code** | Measures the orientation coherence of structures within the 'bundle' mask using structure tensor analysis. Pathological bundles (like NFTs) may show different alignment properties compared to healthy cytoskeletal arrangements. |
| 57 | **tau droplet vs filament pattern bias** | **vlm** | This feature estimates, for a given cell, the relative dominance of droplet-like versus filamentous Tau structures in the Tau MIP, returning a scalar where negative values indicate predominantly droplet-like patterns, positive values indicate predominantly filamentous/bundle-like patterns, and values near zero indicate a mixed state. It probes whether Tau pathology manifests mainly as punctate droplets or as extended fibers and tangles. |
| 58 | **droplet number** | **code** | Number of Tau protein liquid-liquid phase separation (LLPS) droplets in the cell. Computed by counting the number of connected components in the droplet mask (mask_droplet.tif). Each connected component represents one droplet. |
| 59 | **tau soma inclusion complexity score** | **vlm** | This feature visually scores the complexity of Tau-positive inclusions within the cell soma, considering whether they appear as a single homogeneous droplet, a few simple blobs, or a highly irregular, multi-lobed, or fibrillar network filling the soma. It reflects progression from early condensate-like inclusions to more advanced NFT-like structures. Higher scores correspond to more complex, irregular, and network-like soma Tau inclusions. |
| 60 | **tau nuclear enrichment zscore** | **code** | This feature quantifies Tau mislocalization into the nucleus by comparing the mean Tau intensity inside the nuclear mask to the mean and standard deviation of Tau intensity in the cytoplasmic (cell minus nucleus) region, reported as a z-score. A high positive z-score indicates abnormally enriched nuclear Tau relative to the cell’s own cytoplasm, independent of global brightness. This is relevant for detecting aberrant Tau partitioning into nuclear compartments in patient neurons. |

| | | | |
|---|---|---|---|
| 61 | **tau filament alignment with cell axis** | **code** | This feature measures the average alignment of Tau-positive filaments with the main elongation axis of the cell in the MIP. First, the cell's major axis is obtained from mask_cell.tif, then each filament object in mask_filament.tif is fitted with an ellipse or principal direction; the feature is the mean absolute cosine of the angle between filament orientation and the cell major axis, weighted by filament length. Values near 1 indicate predominantly axon-like, longitudinal filaments, whereas lower values indicate more disordered or misoriented filament organization consistent with disrupted microtubule-associated Tau. |
| 62 | **tau droplet size tail heaviness** | **code** | Using mask_droplet.tif, this feature measures the heaviness of the droplet size tail by computing the ratio between the mean area of the largest 10% of droplets and the median droplet area. A large ratio indicates the presence of a few very large Tau droplets compared to the typical droplet size, suggestive of coalescing inclusions or advanced aggregation, whereas a ratio near 1 indicates a more uniform droplet population. This emphasizes rare, large Tau condensates that may be biologically important yet underrepresented by simple averages. |
| 63 | **tau cellwide radial aggregation gradient** | **code** | This feature measures the radial gradient of Tau aggregation from the nucleus outward by dividing the cell mask into a set of concentric rings centered on the nuclear centroid and computing a Spearman correlation between ring radius and ring-wise mean Tau intensity or droplet density. It captures whether Tau aggregates tend to concentrate perinuclearly, at intermediate cytoplasmic distances, or toward the cell periphery, which may change with disease stage. A negative correlation implies perinuclear enrichment, while a positive correlation indicates peripheral bias. |
| 64 | **tau droplet cytoplasmic enrichment score** | **code** | This feature calculates the fraction of droplet area lying inside the cell mask but outside the nuclear mask, divided by the total droplet area in the image. It estimates how specifically Tau droplets are enriched in the cytoplasmic/neuritic compartment rather than nucleus or extracellular space. Higher values suggest cytoplasm-targeted condensates, aligning with stress granule–like Tau recruitment. |
| 65 | **tau droplet soma area fraction** | **code** | Among all Tau-positive droplets within a cell, this feature measures the fraction of total droplet area that lies inside the soma region (cell mask minus nucleus) on the 2D MIP. It captures whether Tau droplets preferentially accumulate in the soma as opposed to neurites, which is relevant to soma-centered Tau pathology. Higher values indicate soma-dominated droplet pathology. |
| 66 | **perinuclear tau droplet enrichment index** | **code** | This feature measures how enriched Tau-positive droplets are near the nucleus by computing the fraction of droplet area that lies within a fixed-distance band surrounding the nuclear mask divided by the fraction expected if droplets were uniformly distributed within the cell (droplet area in band divided by band area over cell area). Values greater than one indicate preferential perinuclear localization of Tau condensates. |
| 67 | **tau extranuclear droplet fraction** | **code** | This feature calculates the fraction of Tau droplets that lie completely outside the nuclear mask, defined as the number of droplet objects whose pixels do not intersect mask_nucleus.tif divided by the total number of droplets in the cell. It reflects whether Tau condensates preferentially form in the cytoplasm rather than the nucleus in these 2D MIPs. Shifts in this fraction may indicate altered subcellular compartmentalization of pathological Tau assemblies. |
| 68 | **tau intracellular vs extracellular droplet fraction** | **code** | This feature estimates the fraction of Tau-positive droplets that are intracellular versus extracellular (ghost tangle-like) by comparing droplet locations to the cell mask. For all droplet objects in mask_droplet.tif, those whose centroid lies inside mask_cell.tif are counted as intracellular; the feature is defined as intracellular_count / total_droplet_count. Lower fractions indicate more droplets occurring outside labeled cells, consistent with ghost tangles and late-stage neurodegeneration. |
| 69 | **tau droplet soma proximity fraction** | **code** | For each Tau droplet (connected component in mask_droplet.tif), this feature measures the minimum Euclidean distance from the droplet centroid to the cell boundary (mask_cell.tif), counting droplets that lie within a fixed distance threshold (e.g., 2 μm in pixels) of the soma edge, and returns the fraction of all droplets that are peri-somatic. It summarizes whether Tau droplets preferentially localize close to the cell body versus being dispersed in distal processes, which may reflect early versus late aggregation patterns. |
| 70 | **droplet tau granularity balance** | **code** | For Tau-positive droplets (liquid-like condensates), this feature measures whether Tau signal within droplets is dominated by many small puncta or by fewer large aggregates. Using the droplet mask as an ROI, Tau signal is thresholded and connected components within droplets are identified; the feature is defined as the ratio of total area in small components (e.g., area below a scale threshold) to total area in large components. A shift from small- to large-structure dominance may reflect maturation of stress-granule-like Tau condensates into more solid aggregates. |
| 71 | **tau filament alignment entropy** | **code** | This feature estimates the orientation distribution of Tau-positive filament pixels (from mask_filament.tif) using a structure-tensor or gradient-based orientation map and then computes the Shannon entropy of the orientation histogram. Low entropy indicates well-aligned filaments (e.g., axon-like) whereas high entropy indicates disordered or tangled filament orientations. This is relevant for capturing transition from ordered axonal Tau to disorganized tauopathy-associated filaments. |
| 72 | **tau high intensity cluster count** | **code** | Counts high-intensity Tau clusters within the cell mask. Higher values indicate more numerous bright Tau foci or microclusters. |
| 73 | **tau filament alignment consistency** | **code** | Within each cell, this feature measures the consistency of Tau filament orientations by computing the structure-tensor-based orientation for pixels in mask_filament, then calculating the circular variance of these orientations and finally averaging the inverse variance (alignment consistency) across cells. High alignment consistency corresponds to long, coherently oriented filaments typical of ordered axonal microtubule-associated Tau, while low consistency indicates disorganized or tangled filament bundles consistent with pathological NFTs. This captures nanoscale ordering changes in super-resolution Tau filaments using only the 2D MIP. |
| 74 | **tau droplet compactness index** | **code** | For each labeled droplet in mask_droplet, this feature computes 4π·area/perimeter² and then averages over all droplets in the image, yielding a scalar compactness index of Tau-positive droplets. Values near 1 indicate round, compact inclusions, whereas lower values indicate elongated or irregular droplets. This quantifies whether patient Tau droplets tend to be spherical condensates or more deformed structures. |

| | | | |
|---|---|---|---|
| 75 | **tau radial enrichment toward cell center** | **code** | Using the cell mask, this feature computes the radial profile of Tau intensity from the cell centroid to the boundary and then calculates a normalized inner-versus-outer enrichment score (e.g., mean intensity in the inner 50% of radius divided by mean intensity in the outer 50%). Higher values indicate Tau concentrating toward the cell center (somatic enrichment), whereas lower values indicate peripheral or neuritic enrichment, aligning with transitions from axon-dominant to soma-dominant Tau pathology. |
| 76 | **tau pathology morphotype score** | **vlm** | This feature asks a model to visually rate the dominant Tau pathology morphotype in each cell on a continuous scale, where low values correspond to predominantly smooth axonal-like filaments with minimal droplets, intermediate values to mixed fibrillar and droplet patterns, and high values to dense, tangle-like bundles and large droplets within soma and proximal neurites. It integrates subtle cues such as filament beading, droplet clustering, and overall re-localization patterns that are difficult to fully capture with handcrafted metrics. The output is a single scalar severity score per cell, averaged if multiple cells are present. |
| 77 | **tau cell peripheral enrichment index** | **code** | Restricting to the cell mask (mask_cell), this feature computes a radial profile of Tau intensity from the cell centroid to the cell boundary and reports the ratio of mean intensity in the outermost radial decile to the mean intensity in the innermost decile. Values greater than 1 indicate peripheral or neurite-enriched Tau, whereas values closer to or below 1 suggest soma-centric Tau accumulation, which is characteristic of certain pathological redistribution patterns. |
| 78 | **tau cell edge to center intensity ratio** | **code** | Within the cell mask, compute the mean Tau intensity in an inner core region (obtained by eroding the cell mask by a fixed radius, e.g., 10–20% of the effective cell radius) and in a peripheral ring region (cell minus eroded core), then calculate the peripheral_mean / core_mean ratio. This feature quantifies whether Tau is preferentially enriched at the cell periphery (e.g., in neurite roots and cortical cytoplasm) versus centrally, which can change with Tau relocalization and aggregate formation. |
| 79 | **tau droplet nuclear proximity enrichment** | **code** | For each Tau droplet in mask_droplet.tif, this feature computes its shortest distance to the nuclear boundary and compares the distribution of distances to a randomized baseline (e.g., by shuffling droplet positions within the cell mask) to obtain a z-score of observed mean distance. It quantifies whether Tau droplets preferentially cluster near the nucleus versus being uniformly distributed in the cytoplasm, which may indicate stress granule-like perinuclear recruitment. A negative z-score denotes enrichment near the nucleus. |
| 80 | **tau texture contrast inside vs outside nucleus** | **code** | This feature compares fine-scale Tau texture between the nucleus and the surrounding cytoplasm by computing a GLCM contrast (or similar second-order texture statistic) within the nuclear mask and in a cytoplasmic ring (cell minus nucleus), then taking their ratio. A strong change in contrast can indicate abnormal intranuclear speckling or loss of structured axonal texture in the cytoplasm that is associated with disease-related redistribution. |
| 81 | **tau droplet edge enrichment ratio** | **code** | For each droplet in mask_droplet, the mean Tau intensity in a one-pixel-wide ring at the droplet boundary is divided by the mean Tau intensity in the droplet interior (eroded region), and the feature is the median of this ratio across droplets. It captures whether Tau within droplets is shell-enriched (ring brighter than core) versus homogeneously distributed, potentially distinguishing different condensate states or maturation stages. Ratios significantly above one indicate edge-enriched Tau organization. |
| 82 | **tau neurite radial decay consistency** | **code** | For each labeled neurite-like object in the cell mask (cell minus nucleus), this feature computes the correlation between Tau intensity and distance from the soma boundary along the neurite skeleton, then averages the correlation coefficients over neurites. A strongly negative and consistent correlation corresponds to the normal proximal-to-distal Tau decay along processes, whereas loss or reversal of this decay indicates pathological redistribution and distal clustering. |
| 83 | **tau filament orientation dispersion** | **code** | This feature measures the dispersion of filament orientations by computing the orientation of each filament fragment (e.g., via regionprops or local structure tensor within mask_filament.tif) and then calculating the circular variance of these angles. Low dispersion indicates highly aligned neuritic Tau, while high dispersion reflects a more disorganized or tangled network of Tau-positive filaments. Changes in this metric can capture neurite disorganization associated with pathology. |
| 84 | **tau intensity gradient along principal filament** | **code** | This feature quantifies how Tau intensity varies from the putative proximal to distal end along the longest Tau-positive process by sampling Tau intensity along the skeleton of the longest filament/bundle and fitting a linear regression of intensity versus arc-length position. The regression slope (normalized by mean intensity) is reported as the feature, with more negative values indicating a strong proximal-to-distal decay similar to normal axonal Tau gradients and near-zero or positive slopes indicating loss or reversal of this gradient in pathology. |
| 85 | **tau filament linear anisotropy** | **code** | This feature estimates the directional coherence of Tau filaments by computing, over pixels within the filament mask, the structure-tensor-based orientation field and then taking the mean anisotropy (difference between largest and smallest eigenvalues divided by their sum). It measures how strongly filaments align along a dominant direction versus forming disordered, tangled networks. Lower anisotropy indicates more disorganized, tangle-like filament arrangements, whereas higher anisotropy reflects more axon-like, aligned filaments. |
| 86 | **tau peripheral enrichment** | **code** | Measures whether Tau fluorescence is enriched toward the cell periphery versus the center, using a radial intensity profile within the cell mask. Higher values indicate stronger peripheral Tau enrichment. |
| 87 | **soma to extracellular tau intensity shift** | **code** | This feature compares Tau enrichment inside versus outside the segmented neuron by computing the log2 ratio of mean Tau intensity within the cell mask to the mean Tau intensity in a pericellular background ring (a fixed-width band outside the cell mask, clipped to the image). It quantifies whether Tau is predominantly intracellular or has leaked/accumulated in extracellular space, which may indicate presence of ghost-tangle-like extracellular aggregates in advanced pathology. |
| 88 | **tau nuclear exclusion ratio** | **code** | The ratio of mean intensity within 'mask_nucleus' to the mean intensity of the surrounding soma (mask_cell minus mask_filament). Healthy cells exclude Tau from the nucleus (low ratio); pathology may lead to leakage or nuclear inclusions. |
| 89 | **tau peripheral enrichment index** | **code** | Within the cell mask, this feature computes a radial profile of Tau intensity from the cell centroid to the boundary and reports the ratio of mean intensity in the outermost 25% of the cell radius to that in the innermost 25%. It measures whether Tau is preferentially enriched at the cell periphery and neurite bases versus the perinuclear region, which may reflect redistribution of Tau from axon-like core structures toward distal or membrane-proximal compartments. It uses distance transforms on mask_cell.tif over the 2D Tau MIP. |

| | | | |
|---|---|---|---|
| 90 | **tau filament directional coherence** | **code** | Considering all filament pixels in mask_filament, this feature estimates the directional coherence of Tau-positive filaments by computing the structure-tensor-based orientation at each filament pixel and then calculating the circular variance of these orientations. Low circular variance (high coherence) indicates filaments predominantly aligned along one or few directions (e.g., organized axonal tracks), whereas high circular variance reflects disordered, tangled filament orientations typical of advanced NFT-like structures in 2D. The feature is reported as 1 minus circular variance so that higher values indicate more coherent alignment. |
| 91 | **tau filament beading score** | **code** | For all filament pixels (mask_filament), this feature estimates a beading score by computing the coefficient of variation (standard deviation divided by mean) of Tau intensity along the filament skeleton. Elevated local intensity variation along filaments corresponds to a beaded or punctate Tau distribution rather than a smooth, uniform filament, a hallmark of pathological fragmentation. |
| 92 | **tau intensity cell to nucleus ratio** | **code** | This feature computes the ratio between mean Tau intensity inside the full cell mask (mask_cell) and mean Tau intensity inside the nuclear mask (mask_nucleus) for each cell, then averages across cells in the image. It captures whether Tau is preferentially enriched in the cytoplasmic/neuritic compartment versus the nucleus, which is expected in normal axon-dominant Tau localization. A relative increase in nuclear-proximal intensity (lower ratio) may correlate with loss of polarized neuritic distribution or altered subcellular trafficking in patient neurons. |
| 93 | **tau cell body enrichment ratio** | **code** | This feature computes the mean Tau intensity inside the cell mask and divides it by the mean Tau intensity in an annular pericellular background region (a fixed-width ring around the cell). It quantifies how strongly Tau is enriched inside the patient neuron relative to its local background on the 2D MIP Tau channel. A high ratio indicates abnormally strong intracellular Tau accumulation, relevant to pathological Tau mislocalization. |
| 94 | **tau cellwide granularity multiscale entropy** | **code** | On the full 2D Tau MIP within the cell mask, this feature applies multi-scale band-pass (e.g., Laplacian-of-Gaussian) filters at several spot-size scales and computes the Shannon entropy of the filter response histograms, then averages over scales. Higher values correspond to richer, multi-scale granular patterns of Tau (e.g., mixed fine puncta and coarse inclusions) suggestive of heterogeneous aggregation states in patient neurons. |
| 95 | **tau droplet enrichment index** | **code** | This feature measures how enriched Tau is within droplet regions by taking the mean Tau intensity inside the droplet mask divided by the mean Tau intensity in the rest of the cell (cell mask excluding droplet mask). It quantifies preferential Tau accumulation in phase-separated or granular droplets versus diffuse cytosolic Tau. High values indicate strong Tau concentration in droplets compared to the bulk cell. |
| 96 | **tau punctate pattern along processes** | **vlm** | Visually scores whether Tau along neurite/process structures appears punctate versus smooth. Higher values indicate a more punctate pattern along processes. |
| 97 | **tau cell internal intensity coefficient of variation** | **code** | For each cell, this feature computes the coefficient of variation (standard deviation divided by mean) of Tau intensities inside the cell mask and then averages this value across cells in the image. It captures intra-cellular heterogeneity of Tau distribution on the 2D MIP, distinguishing cells with smooth, uniform Tau from those with highly heterogeneous, speckled, or inclusion-rich patterns. Higher values reflect more heterogeneous intra-cell Tau patterns consistent with mixed diffuse and aggregated states. |
| 98 | **tau cell level intensity heterogeneity** | **code** | Within each cell mask, compute the coefficient of variation (standard deviation divided by mean) of Tau intensity, then average this coefficient over all cells. This feature captures intra-cellular heterogeneity of Tau signal, distinguishing cells with smooth diffuse Tau from those with a patchwork of bright inclusions and dark areas. High heterogeneity suggests strong subcellular clustering of Tau consistent with pathological aggregation. |
| 99 | **tau cellwide multiscale granularity ratio** | **code** | This feature applies a series of bandpass (difference-of-Gaussians) filters at two spatial scales corresponding to small droplets (e.g., ~0.2–0.4 µm) and larger inclusions (e.g., ~0.6–1.0 µm), computes the average absolute response at each scale within the cell mask, and returns the ratio small_scale_response / large_scale_response. It quantifies whether Tau granularity is dominated by small oligomer-like puncta or by larger inclusions and bundles, providing a coarse index of aggregation stage within the single-cell MIP. |
| 100 | **cell tau compartmental polarization index** | **code** | This feature quantifies how polarized Tau intensity is towards the neurite (cell) periphery versus the nucleus within a single-cell MIP. Using the cell and nucleus masks, it computes the mean Tau intensity in a peripheral ring region (cell mask minus a nucleus-centered eroded core) and divides it by the mean Tau intensity in the perinuclear region. Lower values indicate Tau concentrating towards the nucleus/core (consistent with soma-centered aggregates), while higher values indicate a more peripheral/neuritic bias, relevant to detecting Tau mislocalization from neurites into soma. |
| 101 | **tau background corrected cell intensity cv** | **code** | This feature estimates a global background Tau level from pixels outside all cell masks, subtracts this background from Tau intensities inside each cell, and computes the coefficient of variation of background-corrected mean cell intensities across all cells in the image. It measures inter-cell heterogeneity in Tau accumulation while reducing the influence of field-to-field illumination differences. High values indicate strong cell-to-cell variability in Tau load, which may reflect heterogeneous uptake or aggregation. |
| 102 | **tau cytoplasmic granularity multi scale** | **code** | This feature quantifies cytoplasmic Tau granularity by applying a bank of band-pass filters (e.g., difference-of-Gaussians) at several spatial scales within the cell mask excluding nucleus and then computing the scale at which filter response energy is maximal. The resulting scalar is the response-weighted average spatial scale of granules, reflecting whether Tau is predominantly diffuse, small-droplet-like, or forming larger puncta. This is relevant for capturing transitions from smooth distribution to granule-rich stress-granule-like patterns. |
| 103 | **tau pathology stage score** | **vlm** | A visual assessment score (1-5) estimating the stage of Tau pathology based on distribution: 1=Strictly Axonal, 2=Distal Axon Accumulation, 3=Diffuse Somatic, 4=Mature NFT (flame/globose), 5=Ghost Tangle/Cell Death. |

| | | | |
|---|---|---|---|
| 104 | **tau filament end to end alignment score** | **code** | This feature quantifies how coherently aligned Tau filaments and bundles are within the cell by computing the orientation of each skeleton segment in the filament+bundle mask and then measuring the concentration of the resulting orientation distribution (e.g., resultant vector length on the unit circle). It distinguishes parallel, tract-like Tau organization from disordered, tangled networks. Higher scores indicate predominantly parallel-aligned filaments, whereas lower scores indicate more isotropic, tangled arrangements. |
| 105 | **tau filament alignment score** | **code** | This feature measures how directionally aligned the Tau filaments are within the cell by computing the orientation of each filament object in mask_filament.tif and calculating the circular variance of these orientations, then taking 1 – variance as an alignment score. Values near 1 indicate filaments share a common direction (e.g., ordered axon-like bundles), while lower values indicate a disordered filament network, as expected in pathological tangle formation. |
| 106 | **tau droplet vs fibril morphology bias** | **vlm** | Using the Tau MIP together with droplet, filament, and bundle masks, this feature prompts a model to estimate a continuous score from 0 to 1 indicating whether Tau pathology appears more liquid-droplet–like (round, condensate-like puncta; score near 0) versus fibrillar/NFT-like (elongated, bundled, thread-like structures; score near 1). This captures the qualitative balance between condensate-driven and fibrillar Tau assemblies, which may correspond to different mechanistic states. |
| 107 | **tau soma vs neurite intensity balance** | **code** | This feature calculates the ratio of mean Tau intensity in the soma region to mean Tau intensity in neurites (cell mask minus soma and nucleus). It directly quantifies the shift from a neurite/axon-dominant Tau distribution towards soma-dominant accumulation, which is a hallmark of Tau pathology in neurons. |
| 108 | **tau peripheral to central cell intensity ratio** | **code** | Using the cell mask, this feature erodes the mask to define a central core and subtracts it to obtain a thin peripheral ring, then computes the ratio of mean Tau intensity in the periphery to that in the core. It captures whether Tau is enriched near the cell boundary (e.g., in neurite bases) or concentrated centrally around soma/nucleus, which may shift with disease progression. This provides a simple descriptor of radial Tau redistribution within the cell. |
| 109 | **tau soma vs process fractal dimension** | **code** | This feature compares the complexity of Tau-positive structures within the soma versus in neuritic processes. Within mask_cell, tau-positive pixels above a fixed or adaptive threshold are binarized, and box-counting fractal dimension is computed separately inside an eroded soma core (proximal to mask_nucleus) and in the remaining cell processes; the feature is the difference (process FD minus soma FD). A large positive difference suggests more ramified, filamentous Tau in processes relative to more compact structures in soma, whereas reduced difference indicates pathological spreading of complex aggregates into the soma. |
| 110 | **tau droplet median eccentricity** | **code** | Within each droplet mask, this feature computes the eccentricity (ellipse-based shape elongation) of each droplet object and then takes the median eccentricity across all droplets in the image. It distinguishes between predominantly round, condensate-like Tau droplets (low eccentricity) and more elongated or irregular droplets that may reflect fibrillar coalescence or morphological maturation. Changes in this median can report shifts in droplet shape populations under different patient or treatment conditions. |
| 111 | **tau intranuclear enrichment zscore** | **code** | This feature quantifies whether Tau signal is abnormally present in the nucleus by computing the mean Tau intensity inside the nucleus mask, subtracting the mean intensity in an annular perinuclear cytoplasmic ring, and dividing by the cytoplasmic standard deviation to yield a z-score. Positive values indicate intranuclear Tau enrichment relative to nearby cytoplasm, a potentially aberrant localization in patient cells. |
| 112 | **tau filament curvature mean** | **code** | This feature computes the mean curvature of Tau filaments by fitting polylines to their skeletons and estimating local curvature along each filament, then averaging curvature values weighted by arc length. Lower curvature corresponds to straighter, possibly more tensioned fibers, whereas higher curvature indicates more tortuous or coiled filaments, which may be associated with advanced Tau pathology. |
| 113 | **tau cell interior vs periphery intensity gradient** | **code** | Within mask_cell, this feature converts the cell region to a normalized distance map (0 at boundary, 1 at center) and computes the correlation between the Tau intensity and the distance-to-boundary values. A positive correlation indicates stronger Tau in the perinuclear/interior region; a negative or flat correlation points to peripheral or uniformly distributed Tau. This gradient summarizes whether Tau is centrally concentrated (somatic) or more peripheral (neuritic) in a single scalar. |
| 114 | **tau peripheral ring heterogeneity score** | **code** | This feature measures spatial heterogeneity of Tau intensity along the cell perimeter, sensitive to localized distal axonal accumulations in 2D MIPs. A thin ring is constructed at the cell edge (e.g., by eroding the cell mask), divided into angular sectors around the nucleus or cell centroid, and the coefficient of variation of mean Tau intensity across sectors is computed. Higher values indicate patchy, sector-specific Tau hotspots at the periphery, consistent with focal distal clusters rather than uniform axonal labeling. |
| 115 | **soma tau heterogeneity index** | **code** | This feature measures the heterogeneity of Tau intensity within the soma by computing the coefficient of variation (standard deviation divided by mean) of Tau pixel intensities inside the cell mask but outside the nucleus (mask_cell.tif minus mask_nucleus.tif, restricted to soma region if available). It specifically reflects how patchy or inhomogeneous Tau is within the soma: low values indicate diffuse, relatively uniform Tau, whereas high values correspond to mixed bright inclusions and dark background typical of aggregated pathology within the cell body. |
| 116 | **tau cytoplasmic heterogeneity in cell mask** | **code** | Within the cell mask (mask_cell.tif), compute the coefficient of variation (standard deviation divided by mean) of Tau intensity after excluding the nucleus region. This measures how patchy or heterogeneous the cytoplasmic Tau distribution is on the MIP, independent of overall brightness. Higher heterogeneity suggests a mixture of bright inclusions and darker cytoplasm, consistent with granular or beaded pathology rather than diffuse, uniform labeling. |
| 117 | **tau intensity cv within cell mask** | **code** | This feature computes the coefficient of variation (standard deviation divided by mean) of Tau fluorescence intensity within the cell mask on the 2D MIP. It captures how heterogeneous the Tau signal is inside each cell, distinguishing smooth, diffuse distributions from strongly heterogeneous, punctate or filament-rich cells. Higher values may indicate pathological redistribution or aggregation of Tau within the neuron. |

| | | | |
|---|---|---|---|
| 118 | **tau soma to periphery intensity ratio** | **code** | Within the cell mask, this feature compares mean Tau intensity in a central 'soma-like' region (obtained by eroding the cell mask by a fixed fraction of its equivalent radius) to the mean intensity in the complementary peripheral band. It approximates the redistribution of Tau from processes toward the soma in pathological states, even when explicit axon/dendrite masks are not available. |
| 119 | **tau cell compartmental redistribution score** | **code** | This feature quantifies Tau mislocalization from the cell body to the periphery by computing the difference between the mean Tau intensity in a thin peripheral ring of the cell mask and the mean intensity in an inner core region. Specifically, within mask_cell.tif, an inner core is obtained by eroding the cell mask (e.g. 20–30% of the equivalent radius), and a peripheral ring is the cell mask minus this eroded core; the feature is (peripheral_mean / core_mean). A value >1 indicates relative enrichment of Tau at the cell edge vs. the cell center, which may reflect pathological redistribution toward neurites and away from the soma. This uses only the 2D Tau MIP and the provided cell mask. |
| 120 | **tau distal filament cluster asymmetry** | **code** | This feature captures preferential Tau clustering toward filament tips versus their bases. For each labeled filament in mask_filament.tif, Tau intensity is projected onto its skeleton and the cumulative intensity in the distal 25% of arc-length is divided by that in the proximal 25%; the feature is the average log-ratio across filaments. Positive values indicate distal-biased Tau accumulation, consistent with early distal neurite pathology. |
| 121 | **tau intensity skewness in droplets** | **code** | This feature pools all Tau pixel intensities inside the droplet mask for each cell, computes the skewness of the intensity distribution, and then averages skewness across cells. Positive skewness indicates a distribution dominated by many moderately bright pixels with a tail of very bright cores, consistent with sharply peaked inclusions; near-zero or negative skewness suggests more uniform or plateau-like droplet interiors. This intensity-shape statistic complements simple mean intensity by capturing how sharply Tau concentrates within droplets. |
| 122 | **tau pattern pathology grade** | **vlm** | This feature is a visual pathology grade (e.g., on a continuous 0–1 scale) assigned by a model that inspects the single-cell Tau MIP and corresponding segmentation masks to judge the overall Tau pattern as axon-dominant and smooth versus soma/droplet/bundle-dominated with beaded, tangled morphologies. It aggregates nuanced cues such as beading along neurites, presence of tangle-like bundles in the soma, droplet condensates, and loss of normal axonal gradients into a single scalar severity score. This is intended to capture complex tauopathic phenotypes that are difficult to fully encode with hand-crafted features. |
| 123 | **tau cell edge to center gradient** | **code** | Within the cell mask, this feature quantifies the radial gradient of Tau intensity from the cell center toward the boundary. It can be computed by defining the cell centroid, binning pixels by normalized distance from centroid to cell edge, fitting a line to mean Tau intensity versus radial distance, and using the slope as the feature. Positive or less negative slopes indicate peripheral enrichment (e.g., neurite-dominated Tau), whereas negative slopes indicate central or somatic enrichment. |
| 124 | **tau soma boundary enrichment index** | **code** | For each cell, construct two concentric bands: an inner band adjacent to the inner side of the cell boundary and an outer band just inside the nucleus (if present), then compute the ratio of mean Tau intensity in the peripheral band to that in the inner-somatic band and average across cells. This feature measures perisomatic enrichment of Tau along the cell membrane relative to central soma, which may reflect early mislocalization or membrane-associated aggregation. Values greater than one indicate Tau concentrated near the soma edge. |
| 125 | **tau radial variogram slope** | **code** | Estimates the slope of the radial intensity variogram of Tau across the cell, summarizing how Tau intensity covariance changes with distance from the cell center. |
| 126 | **tau peripheral ring enrichment** | **code** | This feature measures the relative enrichment of Tau intensity in a thin peripheral ring of the cell compared to the cell interior by defining an inner cell mask (eroded version of mask_cell) and a peripheral ring mask (mask_cell minus eroded mask) and computing the mean Tau intensity in the ring divided by mean intensity in the interior. On 2D MIP images, high values indicate Tau accumulation near the plasma membrane and distal neurites, while low values indicate soma-centered Tau. It is meant to capture outward spreading of Tau pathology. |
| 127 | **tau pattern smooth vs beaded score** | **vlm** | This feature asks a model to visually score, on a standardized scale, how beaded versus smooth the Tau-positive filament and bundle structures appear within each cell on the 2D MIP. It captures the presence of periodic swellings or bead-like interruptions along neurites, which are characteristic of pathological Tau redistribution but difficult to robustly quantify with simple filters or 1D profiles in arbitrary orientations. |
| 128 | **tau filament linear alignment score** | **code** | Within mask_filament, this feature computes the average coherence of local structure tensors (or elongatedness of fitted ellipses) across all filament objects, yielding a scalar between 0 and 1 summarizing how well Tau filaments align along a dominant direction. Higher values indicate long, well-aligned fibrils, whereas lower values correspond to more fragmented, disordered or radially oriented filamentous Tau pathology. |
| 129 | **radial tau gradient from nucleus to cell edge** | **code** | This feature measures the radial gradient of Tau intensity from the nuclear centroid to the cell periphery by sampling mean Tau intensity in a series of concentric level-set bands between mask_nucleus and mask_cell and fitting a linear slope of intensity versus normalized radius. It captures whether Tau is enriched near the perinuclear region or towards distal processes in the 2D MIP, which is relevant for detecting somatic versus neuritic redistribution of Tau. A positive or negative slope reflects a bias of Tau accumulation towards center or periphery of the neuron. |
| 130 | **tau beaded neurite severity** | **vlm** | This feature scores the severity of 'beaded' Tau-positive neurites, defined as neurite segments where Tau intensity appears as a string of bright beads separated by thinner, dimmer segments rather than as smooth, continuous lines. It aggregates across the cell to estimate how prevalent and pronounced such beaded patterns are, which reflect fragmented or pearl-on-a-string-like Tau pathology along processes. Higher scores correspond to more frequent and more pronounced beading along Tau-positive filaments in the 2D MIP. |
| 131 | **beaded vs smooth filament pattern score** | **vlm** | This feature is a visual score capturing whether Tau-positive neurites appear predominantly smooth (continuous, uniform intensity) or beaded (string-of-pearls pattern of bright puncta separated by narrower segments) when viewed in the 2D Tau MIP with the provided filament and bundle masks overlaid. The score rates the overall pattern on a continuous scale, for example from 0 (completely smooth, no apparent beading) to 1 (strongly beaded along most Tau-positive processes), integrating subtle context like periodicity, local swellings, and interruptions that are difficult to formalize algorithmically. |

| | | | |
|---|---|---|---|
| 132 | **tau intranuclear invasion fraction** | **code** | Using mask_nucleus and mask_cell, this feature calculates the fraction of nuclear area that is Tau-positive above a robust intensity threshold (e.g., Otsu on Tau within the cell), reporting the Tau-positive nuclear area divided by the total nuclear area. While nuclear Tau is not canonical, intranuclear signal or penetration can indicate severe mislocalization or segmentation artifacts; this scalar captures such events systematically across cells. Higher values indicate more extensive intranuclear Tau signal in the MIP. |
| 133 | **nuclear to cytoplasmic tau intensity ratio** | **code** | This feature calculates the ratio of mean Tau intensity inside the nuclear mask (mask_nucleus) to the mean Tau intensity in the cytoplasmic compartment, defined as the cell mask minus the nuclear mask. It quantifies Tau mislocalization into or out of the nucleus, which may reflect altered nuclear-cytoplasmic trafficking or stress-related nuclear changes in patient neurons. A shift in this ratio compared to controls could indicate pathological redistribution of Tau. |
| 134 | **intranuclear tau invasion index** | **code** | This feature measures potential mislocalization of Tau into the nucleus by computing the ratio of mean Tau intensity inside the nuclear mask to the mean Tau intensity in the surrounding perinuclear cytoplasm (cell mask minus nucleus). It is designed to capture abnormal Tau accumulation within the nucleus relative to the rest of the cell, which could represent a distinct pathological mislocalization phenotype. |
| 135 | **nuclear to cytoplasmic tau ratio** | **code** | This feature computes the ratio of mean Tau intensity within the nucleus mask to the mean Tau intensity in the perinuclear cytoplasm (cell mask minus nucleus mask) on the 2D MIP Tau image. It captures whether Tau signal has abnormally accumulated in or around the nucleus compared to the surrounding cytoplasm, which may reflect advanced mislocalization from axonal compartments toward soma/nuclear vicinity. Values substantially above 1 flag cells with disproportionately high nuclear/perinuclear Tau relative to the rest of the cell. |
| 136 | **tau nuclear enrichment ratio** | **code** | This feature computes the mean Tau intensity inside the nucleus mask divided by the mean Tau intensity in the cytoplasmic cell mask excluding nucleus. On 2D Tau MIP images, it quantifies whether Tau is abnormally enriched in the nuclear region, which can indicate mislocalization and advanced cellular stress in patient neurons. Values >1 reflect disproportionate nuclear accumulation relative to the surrounding soma. |
| 137 | **tau soma vs extranuclear intensity ratio** | **code** | This feature calculates the ratio between mean Tau intensity inside the nuclear mask and mean Tau intensity in the surrounding cytoplasmic cell region (cell mask minus nucleus) on the 2D MIP. It captures redistribution of Tau from axonal/cytoplasmic compartments into the soma/nuclear region, which is indicative of pathological mislocalization and stress-related Tau accumulation. |
| 138 | **nuclear tau intrusion score** | **vlm** | Visually scores the degree to which Tau fluorescence appears to intrude into the nuclear region. Higher values indicate visible Tau-positive signal within the segmented nucleus rather than strong nuclear exclusion. |
| 139 | **tau nucleus to cytoplasm intensity shift** | **code** | This feature measures the shift of Tau signal from cytoplasm toward the nuclear region by computing the ratio of mean Tau intensity inside the nucleus mask to mean Tau intensity in the cytoplasmic compartment (cell mask minus nucleus mask). It is specifically designed for single Tau-channel images and uses mask_nucleus.tif and mask_cell.tif. Elevated values may indicate abnormal Tau accumulation in perinuclear or nuclear-proximal regions associated with pathological mislocalization. |
| 140 | **tau nuclear enrichment index** | **code** | This feature measures the ratio of mean Tau intensity inside the nucleus mask (mask_nucleus.tif) to the mean Tau intensity in the cytoplasmic shell (cell mask minus nucleus). It quantifies whether Tau signal is abnormally enriched in or around the nucleus, which can reflect mislocalization to the soma-associated compartment in pathological states. |
| 141 | **tau droplet maturity score** | **vlm** | This feature reflects the perceived 'maturity' of Tau-positive droplets, distinguishing small, diffuse, fuzzy condensates from larger, sharply bounded, dense inclusions. The model visually evaluates droplet regions in the Tau MIP (using the droplet mask if present) and produces a scalar score where higher values indicate more numerous and more mature, inclusion-like droplets. |
| 142 | **tau beaded filament pattern score** | **vlm** | This feature is a visual score estimating how strongly Tau-positive filaments within the cell exhibit a bead-like ("string of pearls") pattern, where bright puncta are periodically distributed along otherwise continuous filaments. It aims to capture the transition from smooth, uniform axonal Tau to fragmented, punctate Tau along neurites, which is characteristic of emerging Tau pathology. |
| 143 | **tau pattern pathology score** | **vlm** | This feature is a scalar score (e.g., on a fixed scale) that rates how closely the Tau MIP image of a segmented neuron matches a pathological pattern, defined visually as loss of axon-dominant Tau, somatic and dendritic Tau enrichment, presence of bright soma inclusions, and disorganized filament/bundle patterns. It integrates subtle, multi-scale visual cues across the Tau channel and masks (cell, nucleus, droplet, filament, bundle) that may be difficult to capture fully with hand-crafted descriptors. |
| 144 | **tau filament alignment anisotropy in cell** | **code** | This feature estimates the dominant orientation distribution of Tau filaments within the cell by computing the structure tensor orientation histogram in the intersection of mask_filament.tif and mask_cell.tif, then returning 1 minus the circular entropy of this orientation histogram. Higher values indicate strongly aligned filaments, while lower values indicate isotropic, disorganized Tau filament orientations, which may reflect loss of normal axonal alignment. |
| 145 | **tau soma to neurite intensity ratio** | **code** | Calculates the ratio of mean Tau fluorescence intensity within the cell body (soma) to the mean intensity in the neurites (filaments). This is a primary indicator of pathological Tau mislocalization from axons to the soma. |
| 146 | **tau soma to neurite intensity imbalance** | **code** | This feature quantifies the imbalance of Tau between soma and neurites by computing (mean Tau intensity in nucleus-dilated soma region) divided by mean Tau intensity in the cell periphery outside that soma region, using mask_nucleus.tif and mask_cell.tif. It serves as a proxy for somatic mislocalization of Tau: values above 1 indicate that Tau is more concentrated near the soma and nucleus compared to neurites, reflecting pathological relocalization from axons into the cell body. |
| 147 | **bundle compactness in cell** | **vlm** | Visually scores compactness of Tau bundle structures within the cell. Higher values indicate more compact, tightly packed bundles. |

| | | | |
|---|---|---|---|
| 148 | **tau soma vs process visual severity** | **vlm** | This feature is a visually assessed severity score (e.g., on a 0–1 scale) of Tau mislocalization from processes (axons/dendrites) into the soma, based on the relative brightness and concentration of Tau in the soma compared to neurites in the 2D Tau MIP. The score judges whether the signal appears axon-dominant (physiological) or soma-dominant with bright inclusions (pathological), integrating subtle shape and context cues beyond simple intensity thresholds. This captures a high-level phenotype strongly linked to Tau pathology progression. |
| 149 | **tau inclusion maturity score** | **vlm** | Visual assessment of the 'maturity' of somatic inclusions, grading from 0 (diffuse haze) to 10 (dense, structured NFT with clear boundaries). This captures the structural evolution described in the literature that is hard to quantify with simple intensity thresholds. |
| 150 | **peri nuclear tau ring enhancement** | **code** | This feature quantifies Tau enrichment in a thin ring around the nucleus by computing the mean Tau intensity in a band defined as a fixed-width dilation of the nuclear mask minus the nucleus itself, divided by the mean Tau intensity in the remainder of the soma. It captures whether Tau preferentially accumulates at the perinuclear cytoplasm, a pattern often seen when Tau relocates into the soma. Elevated values indicate a pronounced perinuclear Tau ring relative to bulk somatic cytoplasm. |
| 151 | **tau nucleus periphery enrichment index** | **code** | This feature quantifies Tau intensity enrichment at the nuclear periphery by comparing mean Tau intensity in a narrow ring around the nucleus (e.g., a dilated nucleus mask minus the nucleus mask) to the mean Tau intensity in the remainder of the cytoplasm (cell minus dilated nucleus). It measures whether Tau aggregates preferentially accumulate in a perinuclear band, which is consistent with certain stress- or SG-related Tau phenotypes in neurons. If a nucleus mask is missing, the feature is set to NaN. |
| 152 | **perinuclear tau enrichment index** | **code** | For each nucleus, construct a perinuclear ring region within the cell (e.g., 2–4 pixels outside the nuclear boundary but inside the cell mask), compute the mean Tau intensity in this ring, and divide it by the mean Tau intensity in the remaining cytoplasm (cell minus nucleus minus ring). This index quantifies perinuclear Tau enrichment, capturing patterns where Tau accumulates near the nucleus, which can be associated with stress-related reorganization of Tau. Values above 1 indicate preferential perinuclear localization. |
| 153 | **tau nuclear peripheral shell enrichment** | **code** | This feature measures how strongly Tau accumulates in a thin shell around the nucleus relative to the rest of the cell cytoplasm. Within mask_cell.tif, the region within a fixed distance (e.g. 2–3 pixels) outside mask_nucleus.tif is defined as the perinuclear shell, and the remaining cytoplasmic area is the non-perinuclear cytoplasm; the feature is the ratio of mean Tau intensity in the perinuclear shell to the mean Tau intensity in the non-perinuclear cytoplasm. Elevated values indicate Tau clustering near the nucleus, consistent with certain stress or aggregation phenotypes. |
| 154 | **nuclear adjacent tau enrichment ratio** | **code** | This feature quantifies whether Tau accumulates preferentially in a narrow cytoplasmic shell immediately surrounding the nucleus, as expected for soma-centered inclusions. A perinuclear rim is defined by dilating the nucleus mask within the cell and subtracting the nucleus; the mean Tau intensity in this rim is divided by the mean Tau intensity in the remaining cytoplasm (cell minus rim minus nucleus). Ratios above one indicate perinuclear Tau enrichment, a potential indicator of neurofibrillary tangle-like soma pathology. |
| 155 | **tau perinuclear ring intensity contrast** | **code** | This feature measures whether Tau forms a perinuclear ring by comparing Tau intensity in a thin shell around the nucleus to the Tau intensity in the rest of the cytoplasm. A perinuclear shell is defined by dilating the nucleus mask by a fixed distance and subtracting the nucleus; the mean Tau intensity in this shell divided by the mean Tau intensity in the remaining cytoplasm provides the contrast. Values greater than 1 indicate perinuclear enrichment of Tau. |
| 156 | **tau nuclear boundary accumulation score** | **code** | This feature quantifies Tau accumulation around the nuclear boundary by defining a narrow ring (e.g., 3–5 pixels thick) around the nuclear mask (inside the cell) and computing its mean Tau intensity normalized by the mean Tau intensity in the rest of the cytoplasm. A value greater than 1 indicates preferential Tau enrichment at the perinuclear rim, a spatial pattern that may precede or accompany formation of intracellular Tau inclusions. It is computed on the 2D Tau MIP using mask_nucleus.tif and mask_cell.tif. |
| 157 | **tau nuclear ring enrichment** | **code** | This feature measures the mean Tau intensity in a thin perinuclear ring (e.g., a fixed-width band obtained by dilating the nucleus mask and subtracting the nucleus) divided by the mean Tau intensity in the remaining cell cytoplasm. It quantifies whether Tau accumulates preferentially around the nucleus, which can reflect pathological mislocalization toward the soma and perinuclear compartment. Higher values indicate a pronounced perinuclear Tau rim relative to the rest of the cell. |
| 158 | **perinuclear tau ring completeness score** | **vlm** | Visually scores how completely Tau fluorescence forms a continuous annular band around the nucleus. Higher values indicate a more continuous and uniform perinuclear Tau ring. |
| 159 | **nuclear peri tau enrichment index** | **code** | Using the nucleus mask (mask_nucleus.tif), construct a perinuclear ring by dilating the nucleus by a fixed distance (e.g., 1–2 µm in pixels) and subtracting the original nucleus mask, then compute the mean Tau intensity in this ring divided by the mean Tau intensity in the remainder of the cell (mask_cell minus dilated nucleus). This index quantifies whether Tau preferentially accumulates in a perinuclear shell versus more distal regions of the cell. Elevated values may correspond to soma-centered or perinuclear Tau pathology rather than distal axonal enrichment. |
| 160 | **perinuclear tau ring intensity ratio** | **code** | This feature computes the ratio of mean Tau intensity in a perinuclear ring (a band within a fixed pixel distance outside the nuclear mask but inside the cell mask) to the mean Tau intensity in the remaining cytoplasm. It quantifies whether Tau preferentially accumulates around the nucleus, forming a perinuclear shell often associated with pathological clustering and stress-related reorganization. Elevated values point to perinuclear enrichment of Tau. |
| 161 | **reticular perinuclear tau accumulation** | **vlm** | Visually scores reticular Tau accumulation in the perinuclear region. Higher values indicate stronger reticular perinuclear Tau enrichment around the nucleus. |
| 162 | **tau cytoplasm to neurite intensity ratio** | **code** | This feature quantifies redistribution of Tau from neurites into the soma by comparing mean Tau intensity in the perinuclear cytoplasm to mean Tau intensity in neurites. The cytoplasm mask is derived as cell minus nucleus, and neurites are approximated as cell area beyond a distance band around the nucleus. Higher ratios indicate soma-enriched Tau, consistent with pathological mislocalization. |

| | | | |
|---|---|---|---|
| 163 | **tau nuclear peripheral enrichment ratio** | **code** | This feature compares mean Tau intensity in a thin ring surrounding the nucleus (a band within the cell mask at a fixed distance outside mask_nucleus) to mean Tau intensity in the remainder of the cell cytoplasm. It is computed on the 2D Tau MIP and captures whether Tau abnormally accumulates at the perinuclear region, which may relate to stress granule-associated pathology and perinuclear aggregation. Values greater than 1 indicate preferential Tau enrichment near the nuclear periphery. |
| 164 | **tau perinuclear intensity enrichment** | **code** | This feature computes the mean Tau intensity in a perinuclear ring (e.g., 2–4 pixels outside mask_nucleus.tif but inside mask_cell.tif) divided by the mean Tau intensity in the remaining cytoplasm. It quantifies perinuclear accumulation of Tau, which can be associated with stress-related reorganization or early aggregation around the nucleus. |
| 165 | **tau intra neurite beading index** | **code** | This feature quantifies how strongly Tau intensity along each filament/bundle segment fluctuates in a bead-like (punctate) manner rather than remaining smooth. For each connected filament or bundle object inside the cell mask, Tau intensity is sampled along its skeleton, local peaks are detected, and the variance of peak spacing and normalized peak-to-trough contrast is computed; the feature is the length-weighted average of this beading score across all filaments and bundles in the cell. It is relevant because pathological Tau in neurites often appears as irregular 'beaded' structures instead of smooth axonal labeling. |
| 166 | **tau beaded neurite prevalence** | **vlm** | This feature assesses how prevalent "beaded" Tau-positive neurites are in the neuron, defined visually as processes where Tau signal appears as a chain of bright, enlarged puncta separated by thinner segments along a neurite. The metric inspects the Tau 2D MIP together with neurite masks (cell and filament/bundle masks) and outputs a scalar score reflecting the fraction of neurite length showing clear beaded morphology, which is a hallmark of degenerating Tau-positive axons and dendrites. |
| 167 | **tau cell body compaction index** | **code** | This feature measures how compact Tau fluorescence is within the cell mask by computing the ratio between the area of the convex hull of all Tau-positive pixels (above Otsu threshold) inside mask_cell.tif and the total Tau-positive area. A value close to 1 indicates that Tau signal fills a single compact region in the soma, whereas smaller values indicate fragmented or dispersed patterns. This is relevant for distinguishing diffuse Tau from compact inclusion-like aggregates in patient neurons. |
| 168 | **tau subcellular distribution pattern score** | **vlm** | This feature uses a model to qualitatively rate the overall subcellular distribution pattern of Tau in the MIP image—whether Tau appears predominantly axonal, redistributed to soma and dendrites, or concentrated in large somatic inclusions—mapped to a continuous score (e.g., 0 for normal axon-dominant, 1 for strongly soma/dendrite-dominant with large inclusions). It integrates multiple cues such as relative brightness of soma versus processes, presence of somatic droplets or bundles, and loss of clear axonal gradients into a single scalar that reflects Tau mislocalization severity. Such holistic judgments are directly aligned with expert descriptions of Tau pathology progression. |
| 169 | **tau extracellular like fraction** | **code** | This feature estimates the fraction of high-intensity Tau signal lying outside the provided cell mask but within the field of view by thresholding Tau intensity and computing the ratio of suprathreshold area outside mask_cell.tif to the total suprathreshold area. It is intended to approximate the abundance of ghost tangle–like or extracellular Tau deposits relative to cell-associated Tau in patient samples. |
| 170 | **tau soma granularity entropy** | **code** | Measures the entropy of the texture within the cell body (soma) mask after applying a high-pass filter. This quantifies the transition from diffuse cytoplasmic signal (smooth) to granular oligomers or aggregates (rough) characteristic of pathology. |
| 171 | **tau droplet radial distribution from cell center** | **code** | This feature calculates the mean normalized radial position of Tau droplets by taking each droplet centroid's distance from the cell centroid, dividing by the maximum distance to the cell boundary, and averaging over droplets. Values near 0 indicate droplets clustered centrally, whereas values near 1 indicate droplets biased toward the cell periphery, capturing spatial redistribution of aggregates within neuronal soma and processes. |
| 172 | **tau droplet to fibril transition likelihood** | **vlm** | This feature estimates, from visual inspection, how likely the observed Tau morphology reflects a droplet-to-fibril transition state, characterized by bright round droplets connected to or budding off thin filamentous or bundled structures within the cell. Higher scores indicate more frequent and clearer examples of droplets feeding or transforming into linear aggregates in the MIP image. |
| 173 | **tau beaded filament severity score** | **vlm** | This feature asks a model to visually rate, on a continuous scale, the severity of beaded or 'pearls-on-a-string' appearance along Tau-positive neurites within each image, aggregating to a single score (e.g., 0–1 or 0–100). It targets a subtle morphological transition from smooth linear Tau labeling to fragmented, bead-like chains that is challenging to capture with simple filters but critical for staging Tau pathology. Higher scores correspond to more extensive and pronounced beaded filament morphology across cells in the field. |
| 174 | **tau filament beading visual score** | **vlm** | This feature visually scores the degree of beaded or fragmented appearance along Tau-positive neurites, judging whether filaments appear smooth and continuous versus exhibiting prominent bead-like swellings or breaks. It directly targets the expert-described 'beaded' tauopathy morphology that may be challenging to encode purely with local skeleton metrics. |
| 175 | **tau pathology stage estimate** | **vlm** | This feature represents a categorical-to-scalar mapping (e.g., 0–3) of the overall Tau pathology stage for the neuron in the 2D Tau MIP, integrating intensity localization (axon-dominant vs soma/dendrite-dominant), presence of droplets, filaments, and bundles. The model classifies the cell into stages such as normal-like, early mislocalization (diffuse somatodendritic increase), intermediate droplet/filament formation, or advanced bundled fibrillar aggregates, then outputs a corresponding scalar stage score. This condenses complex spatial and morphological patterns into a single progression metric. |
| 176 | **tau cytoplasm enriched cluster fraction** | **code** | Within the cell mask but outside the nuclear mask, this feature thresholds the Tau MIP (e.g., using Otsu or a high-percentile threshold), finds all suprathreshold connected components, and returns the fraction of cytoplasmic area occupied by components larger than a given area threshold. It is designed to capture the presence of sizable cytoplasmic Tau inclusions (NFT-like deposits) rather than diffuse low-level signal. An increased fraction indicates more space taken up by large somatic Tau clusters. |

| | | | |
|---|---|---|---|
| 177 | **tau soma hotspot fraction** | **code** | This feature measures the extent to which Tau forms discrete high-intensity hotspots within the soma rather than being smoothly distributed. Inside mask_cell.tif intersected with mask_nucleus.tif's complement (soma cytoplasm), an adaptive or robust global threshold (e.g. Otsu or mean+2σ) is applied to the Tau channel; the feature is the fraction of soma cytoplasmic area above this threshold. Larger values indicate that a substantial portion of the soma cytoplasm is occupied by high-intensity accumulations, consistent with neurofibrillary tangle-like inclusions. |
| 178 | **tau cellwide radial heterogeneity** | **code** | This feature measures how radially heterogeneous Tau intensity is within the cell by computing radial profiles from the cell centroid outwards (within mask_cell) and taking the coefficient of variation of mean intensities across radial bins. High values indicate strong central vs peripheral differences (e.g., soma-dominated or peripheral aggregate patterns), whereas low values indicate radially uniform signal. It captures coarse spatial redistribution of Tau within the cell. |
| 179 | **tau pattern stage score** | **vlm** | This feature is a visual estimate of the overall Tau pathology stage within a single neuronal cell on a continuous scale based on the relative prominence of axonal, somatic, dendritic, droplet, filament, and bundle patterns in the 2D Tau MIP and segmentation overlays. The model is prompted to integrate cues such as axon-dominant smooth distribution versus soma/dendrite-dominant granular aggregates, presence of beaded processes, and large intracellular bundles or extracellular ghost-like structures, and to map these visual patterns onto a scalar stage index from 'normal-like' to 'advanced aggregated' Tau pathology. |
| 180 | **tau extranuclear ghost like fraction** | **code** | This feature estimates the fraction of high-intensity Tau aggregates that lie outside any cell mask, serving as a proxy for ghost tangle–like extracellular remnants. High-intensity Tau objects are identified as connected components above a global or cell-adaptive threshold; the fraction is computed as the total area of such objects that do not overlap mask_cell.tif divided by the total area of all high-intensity Tau objects in the field of view. Larger values indicate more Tau aggregates that are not associated with intact cell bodies. |
| 181 | **single filament bundle ratio** | **code** | Fluorescence intensity ratio of single filament to bundle. Computed as the mean Tau intensity on single filaments divided by the mean Tau intensity on bundles. This indicates the relative Tau decoration between thin filaments and thick bundles. |
| 182 | **tau soma inclusion maturity score** | **vlm** | This feature visually grades the maturity of somatic Tau inclusions on a continuous scale, considering factors such as inclusion size, density, and internal complexity (e.g., small faint granules vs large dense NFT-like bodies with internal structure). It summarizes how advanced Tau pathology appears in the soma of each neuron, beyond simple area or intensity, by integrating visual cues of inclusion compactness, texture, and organization. Higher scores indicate larger, denser, more complex NFT-like inclusions, while lower scores correspond to absent or only fine diffuse somatic Tau. |
| 183 | **soma tau texture homogeneity** | **code** | Within each nucleus-associated soma region (cell mask around a nucleus), this feature computes gray-level co-occurrence matrix (GLCM) homogeneity of the Tau channel and then averages these homogeneity values across somas. It distinguishes smooth, diffuse Tau distributions (high homogeneity) from coarse, heterogeneous textures produced by dense inclusions and mixed fibrils (low homogeneity). Changes in this feature are indicative of transition from normal low-Tau soma to heavily aggregated, NFT-like soma patterns. |
| 184 | **tau aggregation morphology complexity** | **vlm** | This feature uses a model to assess the morphological complexity of Tau aggregates within a single cell on the MIP, integrating visual cues like the coexistence of droplets, thin fibers, and thick bundles, as well as beaded or tangle-like structures. The model should output a scalar index that increases when multiple distinct aggregate morphologies and higher-order structures (e.g., tangled bundles, ghost-like deposits) are present, reflecting a more advanced and heterogeneous aggregation phenotype. |
| 185 | **tau nuclear exclusion score** | **code** | Measures the degree to which Tau is excluded from the nucleus, calculated as (Mean Cell Intensity - Mean Nuclear Intensity) / Mean Cell Intensity using the provided 'mask_cell' and 'mask_nucleus'. A lower score indicates pathological nuclear invasion of Tau. |
| 186 | **tau inside vs outside cell ghost fraction** | **code** | This feature computes the fraction of total high-intensity Tau area that lies outside the cell mask, by thresholding the Tau MIP to obtain a global Tau-positive mask, computing Tau-positive area inside versus outside the cell mask, and reporting outside_area / (inside_area + outside_area). It serves as a proxy for ghost tangle–like extracellular Tau aggregates that are no longer contained within intact neuronal boundaries. Larger fractions suggest more extracellular or cell-detached Tau pathology. |
| 187 | **tau bundle complex network score** | **vlm** | This feature measures the visual complexity of the Tau bundle network, reflecting how extensively bundles branch, intersect, and form mesh-like structures versus remaining as a few simple, straight tracts. a model inspects the Tau channel within the bundle mask and outputs a scalar score indicating the degree of network-like, tangled architecture consistent with advanced bundle pathology. |
| 188 | **tau morphology stage score** | **vlm** | This feature asks a model to assign a scalar score reflecting the apparent Tau pathology stage of each neuron in the 2D Tau MIP, based on droplet, filament, and bundle morphology within the provided masks (e.g., predominantly diffuse/filamentous vs. dominated by large compact bundles or mixed morphologies). It is designed to capture a high-level, integrative assessment of Tau aggregation state that accounts for subtle shape, texture, and spatial context beyond simple geometric statistics. |
| 189 | **per cell tau spatial gini index** | **code** | This feature measures how unevenly Tau fluorescence is distributed within each cell by computing a Gini coefficient on per-pixel Tau intensities inside mask_cell.tif, then averaging the coefficient across cells in the image. Tau intensities within each cell mask are flattened, sorted, and used to compute the Gini inequality index; values near 0 indicate uniform distribution, while higher values indicate strong spatial concentration into bright subregions. Increased inequality corresponds to focal Tau aggregation rather than diffuse cytosolic presence. |
| 190 | **tau droplet vs filament dominance** | **vlm** | This feature assesses whether the Tau aggregates in a neuron appear predominantly droplet-like (round, liquid-like condensates) or filament/bundle-like (elongated fibrils), returning a continuous score where one extreme represents purely droplet-dominated and the other purely filament/bundle-dominated cells. It captures a conceptual shift in aggregation mode that is difficult to fully encode with simple morphometric thresholds. |

| | | | |
|---|---|---|---|
| 191 | **tau nuclear to cytoplasmic intensity contrast** | **code** | This feature computes the difference between mean Tau intensity inside the nucleus mask and mean Tau intensity in the cytoplasm (cell mask minus nucleus mask), normalized by their sum: (I_nucleus – I_cytoplasm) / (I_nucleus + I_cytoplasm + ε). It quantifies Tau mislocalization toward or away from the nucleus, which may reflect advanced soma pathology where Tau aggregates distort or envelop the nucleus. Values near zero indicate balanced distribution, positive values indicate nuclear enrichment, and negative values indicate cytoplasmic bias. |
| 192 | **tau pathological morphology score** | **vlm** | This feature is a visual score of how strongly the Tau signal in the 2D MIP exhibits pathological patterns such as large soma-centered inclusions, dense dendritic aggregates, and beaded neurites, relative to a healthy axon-dominated, diffuse Tau distribution. a model would rate each cell on a continuous scale (e.g., 0–1) based on the overall morphology and spatial distribution of Tau aggregates within the provided masks. |
| 193 | **tau extracellular aggregate fraction** | **code** | This feature estimates the fraction of high-intensity Tau aggregates that lie outside the cell mask by identifying connected components of suprathreshold Tau signal in the whole image and computing the ratio of aggregate area that has no overlap with mask_cell. It approximates the burden of ghost tangle–like or extracellular Tau deposits relative to intracellular Tau. |
| 194 | **tau neurite pearling index** | **code** | Within the filament+bundle masks representing neurites, this feature measures the degree of bead-like Tau distribution by computing the coefficient of variation of Tau intensity along the skeletonized neurite centerlines: high local peak-to-trough variability along the skeleton yields a higher index. It captures the transition from smooth axonal Tau to pathological 'beaded' or fragmented Tau distributions described in Tauopathy. |
| 195 | **tau soma texture anisotropy** | **code** | Restricted to the soma region (cell mask minus nucleus), this feature computes Gabor-filter responses across multiple orientations and compares the variance of responses between orientations to derive an anisotropy index. A more anisotropic texture indicates Tau organized into oriented fibrils or honeycomb-like meshes within the soma, as opposed to isotropic diffuse staining. |
| 196 | **tau filament alignment index inside cell** | **code** | This feature measures the mean orientation coherence of Tau-positive filaments within the cell by computing the orientation of each filament object in mask_filament (via regionprops) and then calculating 1 minus the circular variance of these angles, restricted to filament pixels inside mask_cell. It summarizes how uniformly aligned filamentous Tau structures are, distinguishing ordered, axon-like patterns from more disorganized filament bundles often associated with pathology. Lower coherence suggests a loss of ordered microtubule-associated alignment in patient neurons. |
| 197 | **tau perinuclear enrichment score** | **code** | This feature quantifies how strongly Tau accumulates in a perinuclear ring relative to the rest of the cytoplasm by comparing mean intensity in a thin band surrounding the nucleus (morphological dilation of mask_nucleus within mask_cell) to mean intensity in the remaining cytoplasmic cell area. Elevated scores suggest perinuclear condensation or early inclusion formation. The score is normalized as (I_ring – I_cytoplasm) / (I_ring + I_cytoplasm). |
| 198 | **tau radial redistribution index** | **code** | This feature quantifies how much Tau signal redistributes from the perinuclear region toward the cell periphery, reflecting soma-to-process or process-to-soma shifts. Using mask_cell.tif and mask_nucleus.tif, a normalized distance transform from the nucleus boundary to the cell boundary is computed, and Tau intensity is averaged in inner (perinuclear) versus outer (peripheral) radial bands; the feature is defined as (outer-band mean intensity minus inner-band mean intensity) divided by their sum. Positive values indicate peripheral enrichment, negative values indicate perinuclear enrichment. |
| 199 | **tau aggregation stage estimate** | **vlm** | This feature prompts a model to assign a continuous stage score reflecting the progression of Tau aggregation in each cell—from predominantly diffuse axonal Tau with minimal somatic involvement, through early somatic/dendritic diffuse enhancement, to prominent somatic NFTs and dense neuritic bundles. The scalar can be obtained by mapping model outputs over ordered textual stage descriptions to a numeric scale. It is designed to capture complex, multi-feature morphological trajectories of Tau pathology that are difficult to encode in a single analytic formula. |
| 200 | **tau intra cell intensity gini** | **code** | This feature computes the Gini coefficient of Tau pixel intensities within the cell mask, summarizing how unequal the distribution of Tau signal is across the cell area (0 = perfectly uniform, values approaching 1 = highly concentrated in few pixels). It captures the transition from diffuse Tau distribution to highly focal aggregates that occupy a small fraction of the cell area but dominate the signal, as observed in pathological Tau aggregation. It uses all Tau pixels inside mask_cell.tif on the 2D MIP. |
| 201 | **tau intra cellular heterogeneity gini** | **code** | This feature measures the inequality of Tau intensity distribution within the cell mask by computing the Gini coefficient over all Tau pixel intensities inside mask_cell.tif. A low Gini coefficient corresponds to relatively uniform Tau distribution, while a high coefficient indicates strong spatial heterogeneity with a few very bright regions (potential aggregates) dominating the intensity. This complements simple variance by emphasizing aggregate-dominated intensity profiles characteristic of advanced Tau pathology. |
| 202 | **tau cell edge enrichment score** | **code** | This feature evaluates how much Tau accumulates near the cell boundary by computing the mean Tau intensity in a narrow band along the inside of the cell mask edge and dividing it by the mean intensity in the remaining interior. Elevated edge enrichment may indicate mislocalization of Tau toward neurite origins or membrane-proximal condensates. |
| 203 | **tau cell edge enrichment index** | **code** | Restricted to the cell mask (mask_cell.tif), this feature computes the ratio of mean Tau intensity in a thin peripheral band along the cell boundary (e.g., 3–5 pixels inside the mask border) to the mean Tau intensity in the interior region. It quantifies whether Tau is enriched at the cortical cytoplasm or along neurite membranes versus more centrally distributed, which may reflect changes in trafficking or aggregation at the cell periphery. Edge-enriched patterns suggest abnormal relocation of Tau toward membranes or processes. |
| 204 | **tau pathology severity visual score** | **vlm** | This feature represents a visual severity score summarizing how advanced Tau pathology appears within the neuron in the 2D MIP Tau channel, integrating cues such as presence and density of large inclusions, beaded neurites, and global redistribution from fine filaments to dense aggregates. a model would rate the cell on a continuous scale (e.g., 0 to 1) corresponding to minimal, moderate, or severe Tau aggregation. This high-level feature is directly relevant for ranking patient neurons by pathological burden. |

| | | | |
|---|---|---|---|
| 205 | **tau droplet maturity assessment** | **vlm** | This feature uses a model to inspect Tau-positive droplets (from mask_droplet) and rate, on a continuous scale from 0 to 1, how advanced or 'mature' the droplet population appears (e.g., many small, spherical and isolated droplets vs. larger, irregular, coalesced condensates and emerging fibrillar structures), then averages this rating across droplets and cells. It is designed to capture morphologic transitions in Tau condensates that precede or accompany fibril and NFT formation, reflecting phase separation and coarsening that are hard to parameterize exhaustively. Higher scores indicate more irregular, coalesced, and advanced-looking Tau condensates. |
| 206 | **intracellular tau polarity relative to nucleus** | **code** | This feature quantifies how polar the Tau distribution is around the nucleus by computing the magnitude of the vector sum of Tau intensity–weighted position vectors from the nuclear centroid, normalized by total Tau intensity. Using mask_nucleus.tif and mask_cell.tif, all Tau-positive pixels inside the cell are assigned vectors pointing from the nuclear centroid; summing and normalizing yields a dimensionless polarity score between 0 (perfectly symmetric Tau distribution) and 1 (all Tau concentrated in one directional sector). High polarity may indicate asymmetric Tau mislocalization toward a specific pole or neurite. |
| 207 | **tau nuclear peripheral enrichment** | **code** | Using mask_nucleus.tif, this feature computes the ratio of mean Tau intensity in a thin annulus at the nuclear periphery (generated by dilating the nuclear mask by a fixed pixel radius and subtracting the original mask) to the mean Tau intensity inside the nuclear interior. It detects whether Tau accumulates preferentially at the nuclear envelope versus diffusely within the nucleus, which could indicate aberrant nuclear-associated Tau pathology. |
| 208 | **tau intra cell intensity heterogeneity** | **code** | Within mask_cell.tif, this feature computes the coefficient of variation (standard deviation divided by mean) of Tau intensity at the pixel level. It summarizes how heterogeneous Tau distribution is across the cell area, distinguishing cells with smooth, diffuse Tau from those with highly punctate or filamentous patterns that create strong intra-cell variability. |
| 209 | **tau pattern soma vs neurite pathology score** | **vlm** | This feature is a score (e.g., from 0 to 1) quantifying how strongly the Tau pattern in a cell matches a pathological phenotype where soma and neurites show abnormal Tau accumulation compared to an axon-dominant, healthy pattern. The score assesses the overall visual appearance of Tau in the provided Tau MIP and segmentation overlays, including subtle aspects such as beaded neurites, diffuse soma brightening, and disruption of normal axonal gradients. The scalar score summarizes this visual judgment into a single dimension for downstream analysis. |
| 210 | **tau intranuclear to perinuclear intensity ratio** | **code** | This feature measures the mean Tau intensity inside the nucleus mask divided by the mean Tau intensity in a concentric perinuclear ring (e.g., dilated nucleus minus nucleus, clipped to the cell mask). It tests for aberrant nuclear Tau accumulation or nuclear exclusion by comparing intranuclear signal to that in the immediate perinuclear cytoplasm, which may reflect stress-related mislocalization or transport defects. |
| 211 | **tau nucleus to cytoplasm leakage index** | **code** | This feature computes the ratio of mean Tau intensity inside mask_nucleus to mean Tau intensity in the perinuclear cytoplasm (cell mask minus nucleus, restricted to a narrow ring) on the 2D MIP. It captures abnormal nuclear entry or accumulation of Tau versus cytoplasmic localization around the nucleus, which can reflect perturbations of nucleocytoplasmic trafficking in Tau pathology. Elevated ratios indicate stronger nuclear Tau leakage relative to perinuclear cytoplasm. |
| 212 | **tau intra cellular radial heterogeneity** | **code** | This feature quantifies how radially heterogeneous the Tau intensity is within the cell by computing the coefficient of variation of mean Tau intensity across concentric radial bands from the nuclear centroid to the cell periphery. On the 2D MIP, the cell is divided into several equal-area shells, and the variation of shell-wise mean intensities is used as the metric. High heterogeneity suggests strong proximal–distal gradients or patchy Tau redistribution, which are hallmarks of mislocalization from axons to soma and dendrites. |
| 213 | **tau intranuclear signal enrichment** | **code** | This feature measures whether Tau signal abnormally invades the nucleus by computing the mean Tau intensity inside mask_nucleus divided by the mean Tau intensity in the perinuclear cytoplasm (mask_cell dilated minus mask_nucleus, constrained to the cell). Values significantly above 1 suggest nuclear Tau enrichment relative to nearby cytoplasm. Such aberrant intranuclear localization can reflect atypical Tau pathology or imaging artifacts that need to be quantified. |
| 214 | **tau cellwide granularity entropy** | **code** | This feature quantifies the diversity of Tau granular structures across the entire cell by computing a multiscale granularity spectrum (e.g., area opened or bandpass-filtered responses at several radii) and then taking the Shannon entropy of the normalized spectrum. It reflects how many different characteristic spot/cluster sizes are present, from small puncta to larger aggregates, capturing the complexity of Tau aggregation states within a cell. Higher entropy indicates a richer mix of granule scales and thus more heterogeneous Tau pathology. |
| 215 | **global tau pathology severity estimate** | **vlm** | Given the full Tau channel MIP and all region masks for a single neuron, this feature has a model provide a scalar severity estimate of Tau pathology based on visual cues such as redistribution from axon to soma/dendrites, presence and size of bright inclusions, fragmentation of filaments, and any ghost-like extracellular aggregates. It condenses multiple complex morphological aspects into a single expert-like pathology score. |
| 216 | **tau filament local beading score** | **code** | This feature quantifies the degree of beading along Tau filaments by first skeletonizing mask_filament, then sampling Tau intensity along each skeleton and counting local intensity peaks; it returns the mean number of peaks per unit skeleton length. Higher values indicate a more pronounced bead-like pattern along neurites, reminiscent of pathologic fragmentation and varicosities. |
| 217 | **tau pattern normal vs pathological score** | **vlm** | This feature is a scalar score reflecting how consistent the overall Tau pattern within the cell MIP appears with a normal axon-dominant, smooth distribution versus a pathological pattern with soma/neurite aggregates, droplets, or beaded filaments. The model is prompted to visually assess the relative Tau intensities in soma, nucleus, neurites, droplets, filaments, and bundles (using overlays of the provided masks) and output a single continuous score where low values correspond to normal-like patterns and high values to strongly pathological Tau redistribution and aggregation. |
| 218 | **tau cell polarization asymmetry** | **code** | This feature characterizes how polarized the Tau distribution is within each cell by comparing Tau intensity in opposite half-cells. For each cell, the principal axis is found from mask_cell.tif, the cell is split into two halves along this axis, and the absolute difference in mean Tau intensity between the two halves, normalized by the total, is computed and averaged over cells. High asymmetry indicates strong polarization (e.g., Tau concentrated toward one pole or neurite), while low values suggest more uniform or diffuse distribution. |

| | | | |
|---|---|---|---|
| 219 | **tau pathology stage visual score** | **vlm** | This feature represents a model–derived scalar score summarizing the overall Tau pathology stage per image by visually assessing the 2D Tau MIP and the provided segmentation overlays. The model should jointly consider Tau localization (axon-dominant vs soma/dendrite-rich), presence of compact bundles versus diffuse filaments, droplet-like granules, and neuritic beading to assign a continuous severity score (e.g., from minimal/normal-like to advanced NFT-rich pathology). It integrates multiple subtle morphological cues that are hard to capture fully with hand-crafted features. |
| 220 | **tau cell body pathology index** | **code** | This feature quantifies how strongly Tau aggregates within the cell body by combining the fraction of cell-mask area occupied by high-intensity Tau clusters and their relative brightness. On the 2D Tau MIP, Tau intensities within the cell mask are thresholded (e.g., Otsu or high quantile) to define suprathreshold clusters, and the feature is computed as (suprathreshold area / cell area) multiplied by (mean suprathreshold intensity / mean cell intensity). Higher values indicate more extensive and brighter somatic Tau inclusions, consistent with pathological somatic accumulation. |
| 221 | **tau cellwide intensity fractal dimension** | **code** | This feature estimates the fractal dimension of the Tau-positive pattern within the cell, reflecting how space-filling and complex the Tau distribution is. Using mask_cell as a support, the Tau channel is binarized at a fixed percentile threshold, and a box-counting method is applied across multiple scales to estimate the fractal dimension of the Tau-positive set. Higher fractal dimension indicates more space-filling, complex networks of Tau (e.g., widespread fibrils and aggregates) rather than sparse, simple labeling. |
| 222 | **tau cellwide structure tensor anisotropy** | **code** | This feature computes the mean anisotropy of the 2D structure tensor of the Tau MIP signal across the cell mask, summarizing how strongly Tau intensity patterns are locally oriented (filamentous) versus isotropic (droplet-like). High anisotropy indicates predominance of aligned filaments or bundles, while low anisotropy suggests more isotropic, granular aggregates and diffuse patterns. It can thus differentiate filamentous versus droplet-dominated Tau morphologies at the cell scale. |
| 223 | **tau morphology progression score** | **vlm** | This feature asks a model to visually score the overall Tau morphology in the cell on a continuous scale (e.g., 0–1) representing progression from normal axon-dominant, fine punctate Tau to severe pathology with soma-centered NFTs, dense bundles, and extensive droplets, using all provided segmentation masks as context. The model integrates spatial relationships between nucleus, soma, droplets, filaments, and bundles on the 2D MIP to estimate how advanced the Tau aggregation phenotype appears. It is designed to capture complex gestalt patterns that are difficult to formulate as simple handcrafted metrics. |
| 224 | **tau morphology pathology score** | **vlm** | This feature is a visual score estimating the overall severity of Tau pathology in a single-cell MIP, integrating expert-like assessments of diffuse versus punctate signal, presence of filaments and bundles, and abnormal nuclear or perinuclear accumulation. It aims to mimic an expert neuropathologist's holistic judgment of Tau aggregation stage from the single-channel Tau image and its masks. |
| 225 | **tau cell polarization score** | **code** | This feature computes the Tau-intensity-weighted centroid within the cell mask and measures its distance from the cell's geometric centroid, normalized by the cell's equivalent radius. It quantifies how polarized the Tau distribution is within the 2D MIP—e.g., biased toward one side of the soma/neurites versus evenly distributed. Loss or reversal of normal axon-directed polarization and emergence of somatodendritic Tau can manifest as changes in this score. |
| 226 | **tau cell polarization moment** | **code** | This feature computes a vector from the cell centroid to the intensity-weighted Tau centroid and reports its magnitude normalized by the effective cell radius (square root of cell area divided by $\pi$). It measures how polarized Tau distribution is within the cell, for example biased toward one side along a main neurite versus isotropically spread. Larger normalized displacement suggests strong Tau polarization (e.g., axon-biased), whereas smaller values may reflect diffuse, soma-centered pathology. |
| 227 | **cell level tau polarity score** | **code** | For each cell, this feature computes the intensity-weighted centroid of Tau signal within the cell mask and the geometric centroid of the cell, measures the distance between these centroids, normalizes by an effective cell radius, and then averages this normalized distance over all cells. It quantifies how polarized the Tau distribution is within cells, with high scores reflecting strong asymmetry (e.g., Tau concentrated toward a single axon or dendritic pole) and low scores reflecting more symmetric, soma-centered or globally redistributed Tau seen in pathology. This helps capture shifts from axon-polarized to soma-diffuse Tau localization. |
| 228 | **axon like filament length to cell diameter ratio** | **code** | For each cell, this feature computes the ratio of the longest geodesic length of a Tau filament/bundle (approximated via skeletonization of mask_filament $\cup$ mask_bundle) to the effective cell diameter (square root of cell mask area). It captures how far the longest Tau-positive process extends relative to cell size, serving as a proxy for preservation or loss of long axon-like Tau distribution in pathological states where Tau retracts into soma and proximal processes. |
| 229 | **tau mislocalization severity score** | **vlm** | This feature is a visual, cell-level score of Tau mislocalization severity across the image, integrating the relative prominence of axonal-like linear Tau, somatic inclusions, and dendrite-like network enrichment seen in the single Tau-channel MIP. a model is prompted to qualitatively rate each cell (e.g., from 0 = predominantly axonal, diffuse Tau to 1 = strong somatic inclusions and dendritic filling) based on the provided masks and raw image, and the per-cell scores are averaged into a scalar. Higher values correspond to more advanced mislocalization, complementing low-level quantitative ratios. |
| 230 | **tau droplet morphology complexity** | **vlm** | This feature is a score of the morphological complexity of Tau droplets in the 2D MIP, considering aspects such as size heterogeneity, shape (round vs elongated), clustering, and proximity to filaments and bundles. It aims to distinguish simple, small spherical droplets from complex, coalesced, or irregular condensates that may correspond to different aggregation states. Higher scores indicate more heterogeneous, irregular, or clustered droplet morphologies. |
| 231 | **tau axon dominance visual index** | **vlm** | This feature visually estimates the degree to which Tau fluorescence is dominated by thin, linear axon-like processes versus soma and dendritic regions. a model inspects the Tau MIP (with cell context) and assigns a continuous index, where high values indicate predominantly smooth, axonal line-like signal and low values indicate substantial somatic/dendritic accumulation or loss of clear axon-dominant patterning. |
| 232 | **tau aggregate maturity visual score** | **vlm** | This feature assigns a visual maturity score to Tau pathology within each cell, ranging from predominantly diffuse or small puncta to large, complex, neurofibrillary tangle-like bundles occupying the soma and processes. It should integrate aggregate size, compactness, branching, and impact on overall cell shape in the 2D MIP to capture the stage of Tau aggregation. |

| | | | |
|---|---|---|---|
| 233 | **tau soma aggregation severity score** | **vlm** | This feature provides a visual severity score (e.g., 0–1) for Tau aggregation within the soma by evaluating the Tau MIP restricted to the cell body (mask_cell minus neurite regions). It considers the number, size, and brightness of intracytoplasmic Tau inclusions relative to diffuse background, corresponding to progression from diffuse enrichment to large, dense NFTs-like aggregates. The score summarizes overall somatic aggregation burden per cell. |
| 234 | **tau neurite beading index** | **code** | Restricted to neuritic filament regions (mask_filament within mask_cell), this feature projects Tau intensity along skeletonized neurites and counts the number of pronounced local intensity peaks per unit neurite length, then averages this peak density across all cells. A high beading index corresponds to a pronounced “string-of-beads” appearance—alternating bright swellings and thin connections—that is characteristic of degenerating Tau-positive neurites, whereas healthy neurites show smoother, more uniform profiles. The feature thus numerically captures neurite beading severity from the 2D MIP. |
| 235 | **tau distribution pattern score soma vs processes** | **vlm** | This feature asks a model to visually rate how strongly Tau signal is concentrated in the soma versus neuronal processes (filaments and bundles) on the 2D MIP. The model should consider whether Tau appears axon-dominant (bright along a thin process, weak in soma) versus soma/dendrite-dominant (bright globular soma and thick processes) and output a scalar score on a continuous scale, where higher scores indicate more soma/process-enriched pathology-like patterns. |
| 236 | **tau subcellular mislocalization score** | **vlm** | Using the Tau channel MIP alongside cell and nucleus masks, this feature prompts a model to rate the degree of Tau mislocalization from axonal/neuritic regions into the soma and nucleus on a continuous scale. It reflects expert-like judgment of whether Tau is in an axon-dominant, mixed, or soma-dominant pattern considering morphology and relative intensities in different compartments. |
| 237 | **tau axon dominance score** | **vlm** | This feature is a visual estimate of how strongly Tau signal is dominated by axonal-like thin processes versus somatic and dendritic compartments in a single-cell MIP. It asks the model to judge whether Tau appears mainly as smooth, elongated linear signal along one or few thin neurites, with relatively low somatic and dendritic brightness, corresponding to a healthy axon-dominant distribution. |
| 238 | **tau pattern soma beaded vs diffuse score** | **vlm** | Within the soma region defined by mask_cell.tif excluding mask_nucleus.tif, this feature asks a model to rate on a continuous scale how strongly Tau signal appears as beaded or punctate aggregates lined up along short trajectories versus as smoothly diffuse staining without distinct beads. The score captures subtle gestalt differences in intracytoplasmic Tau appearance that link to early versus advanced granular pathology and are difficult to formalize in simple numerical features. |
| 239 | **tau nucleus peripheral enrichment score** | **code** | Using the nucleus mask, define an inner core (eroded nucleus) and a peripheral rim (nucleus minus inner core), then compute (rim_mean - core_mean) / (rim_mean + core_mean + ε) of Tau intensity. This normalized contrast quantifies whether Tau preferentially accumulates at the nuclear periphery versus the nuclear interior in 2D MIP, which may reflect stress-related or pathological redistribution. |
| 240 | **tau soma granularity spectrum ratio** | **code** | Within the cell mask excluding the nucleus (soma cytoplasm), this feature computes the ratio of high-frequency to low-frequency granularity energy by applying Laplacian-of-Gaussian filters at two scales (e.g., small sigma for fine granules and larger sigma for coarse structures) and summing squared responses. It captures whether somatic Tau is predominantly fine-grained (diffuse) or dominated by coarse, punctate or droplet-like aggregates. An increased ratio of coarse-to-fine granularity indicates progression toward pronounced Tau inclusions in the soma. |
| 241 | **dendritic tau beading severity** | **vlm** | This feature visually assesses how prominent Tau beading is along dendritic processes, focusing on whether Tau signal forms a smooth continuous line, mild irregular thickenings, or pronounced bead-like swellings along dendrites. Using the Tau MIP and the mask_droplet.tif and mask_filament.tif within the dendritic portions of the cell, the model is prompted to assign a scalar severity score reflecting the degree of bead-like fragmentation of Tau along dendrites. Higher scores indicate more severe dendritic beading, a key morphological marker of pathological Tau. |
| 242 | **tau soma aggregate complexity index** | **vlm** | This feature uses a model to assign a continuous score to the visual complexity of Tau aggregates within the soma, ranging from absent or a single homogeneous bright inclusion (low complexity) to multiple, irregularly shaped, intertwined filamentous or honeycomb-like aggregates (high complexity). It aims to capture qualitative transitions from diffuse or simple inclusions to tangled, NFT-like architectures that are challenging to quantify purely by low-level regionprops and texture metrics. |
| 243 | **tau axon vs dendrite pattern score** | **vlm** | This feature asks a model to visually assess, across all neurons in the image, how strongly Tau fluorescence appears confined to thin, relatively unbranched axon-like processes versus being redistributed into thicker, highly branched dendritic trees and somata. The model returns a scalar score (e.g., from 0 to 1) representing the degree of axon-dominant, gradient-like Tau patterning (high score) versus widespread dendrite/soma enrichment and beaded neuritic pathology (low score). This captures complex, multi-scale morphology and topology that are difficult to fully encode with simple skeleton and intensity statistics. |
| 244 | **tau ghost tangle likelihood** | **vlm** | This feature estimates, on a 0–1 scale, how likely the image contains Tau-positive ghost tangle–like structures: bright Tau aggregates that are clearly outside any intact cell outline (mask_cell.tif) and not associated with obvious nuclei. A high score corresponds to visually distinct extracellular Tau inclusions or debris that resemble remnants of degenerating neurons, while a low score indicates Tau aggregates are confined to intact cells. This reflects late-stage pathology where extracellular Tau-rich remnants accumulate. |
| 245 | **bundle sparseness along the skeleton** | **vlm** | Visually scores how sparse versus compact Tau-positive bundles appear along their skeleton. Higher values indicate more sparse, fragmented bundle architecture. |
| 246 | **tau pathology pattern severity score** | **vlm** | On the Tau MIP and provided segmentation overlays, this feature asks a model to rate from 0 to 1 the overall severity of Tau pathology in the neuron based on visual hallmarks such as soma-centered inclusions, dendritic beading, filament disorganization, and droplet or bundle abundance. The score should reflect a holistic visual assessment of how closely the cell resembles advanced patient-like Tau aggregation patterns described in the domain knowledge. |
| 247 | **tau pathology morphology severity score** | **vlm** | This feature is a scalar score summarizing overall Tau pathology morphology in the 2D MIP Tau channel, integrating visual cues such as presence of large soma inclusions, beaded neurites, distal clusters, and disorganized filament networks relative to the provided masks. Each cell is rated each cell on an ordinal or continuous scale from normal axon-dominant Tau distribution to severe mislocalization and aggregation. This complements coded features by capturing complex, composite patterns described in Tauopathy literature that are difficult to formalize algorithmically. |

| | | | |
|---|---|---|---|
| 248 | **tau aggregation stage score** | **vlm** | Visually grades the Tau distribution pattern into stages (e.g., 0=Axonal, 1=Diffuse Somatic, 2=Dense NFT) based on the textual descriptions of Braak staging equivalents. |
| 249 | **tau intranuclear puncta intensity fraction** | **code** | Within the nuclear mask (mask_nucleus), this feature detects small high-intensity Tau puncta by applying a local threshold or Laplacian-of-Gaussian spot detector and reports the fraction of total nuclear Tau intensity contained in these puncta versus diffuse background. A higher fraction indicates more discrete intranuclear Tau accumulations, which may correspond to aberrant nuclear inclusions or stress-related condensates. |
| 250 | **tau nucleus to cell edge distance weighted intensity** | **code** | This feature computes a distance-weighted average Tau intensity within the cell, where each pixel's intensity is weighted by its normalized distance from the nuclear centroid to the cell boundary (e.g., 0 at the nucleus center, 1 at the cell edge). It summarizes whether Tau signal is biased toward central (perinuclear) or peripheral (neuritic) regions along the soma-to-neurite continuum. Lower values imply Tau concentrated near the nucleus, while higher values indicate Tau biased toward the cell periphery and processes. |
| 251 | **axon like tau skeleton fragmentation index** | **code** | Within the cell mask, this feature identifies Tau-positive skeletons that are long and thin (proxy for axon-like processes) and quantifies fragmentation by counting the number of skeleton endpoints per unit skeleton length. Skeletons are derived from thresholded Tau within non-nuclear cell regions, filtered by high aspect ratio and minimal thickness to approximate axon-like segments; the fragmentation index is defined as total endpoints divided by total skeleton length. Higher values indicate more beaded, broken, or discontinuous Tau along putative axon-like processes, consistent with axonal degeneration or beading pathology. |
| 252 | **tau filament fragmentation index** | **code** | Computes the number of disconnected components in the filament mask (`mask_filament.tif`) normalized by the total filament length. High values indicate neuritic beading or fragmentation, a key sign of cytoskeletal breakdown. |
| 253 | **tau cellwide granularity multiscale index** | **code** | This feature measures Tau granularity by applying a series of band-pass or difference-of-Gaussians filters at several spatial scales across the entire cell mask and computing the normalized sum of absolute filter responses. It summarizes the presence of punctate and beaded patterns at multiple sizes, distinguishing smooth Tau distributions from ones dominated by puncta and small aggregates. Higher values indicate stronger multiscale granularity in Tau signal. |
| 254 | **tau cell body vs total intensity fraction** | **code** | This feature computes the fraction of total Tau fluorescence contained within the cell body relative to the total Tau intensity across the whole cell mask. For each sample, Tau intensity is summed over the cell mask and over the nucleus+perinuclear soma region (approximated by a dilation of the nucleus within the cell mask), and the ratio soma_sum / cell_sum is reported. Elevated values indicate Tau mislocalization from neurites into the soma, a hallmark of pathological Tau redistribution in patient neurons. |
| 255 | **cytoplasmic tau granularity at droplet scale** | **code** | Restricted to the cytoplasmic region (cell mask minus nucleus), this feature applies a bank of Laplacian-of-Gaussian filters tuned to the median droplet radius (estimated from droplet mask areas) and computes the normalized variance of the filtered Tau response. It quantifies how strongly the Tau signal is punctated at the characteristic droplet scale, independent of explicitly segmented droplets, thereby capturing diffuse nanocluster-like granularity versus smoother backgrounds. Higher values indicate more pronounced droplet-scale granularity of Tau within the cytoplasm. |
| 256 | **tau extranuclear nucleus overlap fraction** | **code** | Quantify the fraction of Tau-positive pixels (e.g., above a robust global threshold) that lie within the nucleus mask versus outside it, then specifically report the fraction inside the nucleus. Although Tau is primarily cytoplasmic/axonal, mislocalization or strong overlap with nuclear regions on the 2D MIP could reflect aberrant localization or segmentation inconsistencies. This feature captures the degree of apparent nuclear Tau presence relative to extranuclear Tau. |
| 257 | **tau intensity of a single filament** | **code** | Average fluorescence intensity of Tau protein on single microtubule filaments. Computed by taking the mean intensity of the fluorescence image within the filament mask (mask_filament.tif) excluding the bundle mask regions. This isolates Tau signal on fine individual filaments. |
| 258 | **tau multiscale laplacian peak density** | **code** | Estimates multiscale Laplacian peak density of Tau intensity texture, capturing fine-scale intensity peaks that are better resolved at higher optical resolution. |
| 259 | **tau extranuclear fraction in cell** | **code** | This feature measures the fraction of Tau signal in the cell that is located outside the nucleus by computing (sum Tau intensity in cell mask minus sum in nucleus mask) divided by sum in cell mask. It quantifies the degree to which Tau is redistributed from nuclear-proximal regions to the extranuclear soma and neurites, which may increase with pathological mislocalization and aggregation. It operates on the 2D Tau MIP using mask_cell.tif and mask_nucleus.tif. |
| 260 | **tau extranuclear fraction** | **code** | This feature calculates the fraction of total cellular Tau intensity that lies outside the nucleus, by dividing the sum of Tau intensities in the cell mask excluding nucleus by the sum within the entire cell mask. It captures the degree of nuclear versus cytoplasmic Tau localization, which can shift in certain pathological states and is complementary to purely intensity-based or morphological features. |
| 261 | **tau intranuclear leakage fraction** | **code** | This feature computes the fraction of total cellular Tau signal that lies within nuclear masks by summing Tau intensity inside mask_nucleus and dividing by the sum inside mask_cell, aggregated over all cells. It captures abnormal intranuclear Tau accumulation or leakage relative to total cellular Tau in 2D MIP Tau images. Non-negligible values may indicate severe mislocalization or imaging artifacts; near-zero values are expected in typical Tau pathology localized to cytoplasm and neurites. |
| 262 | **intranuclear tau fraction** | **code** | Compute the fraction of total cellular Tau signal (sum of intensities within the cell mask) that lies inside the nuclear mask. This scalar captures abnormal nuclear accumulation of Tau, which should be minimal in typical axon-dominant distributions but may increase in stressed or pathologic conditions. It is robust to absolute intensity scaling since it uses a within-cell fraction. |
| 263 | **tau intranuclear infiltration fraction** | **code** | This feature measures the fraction of total cell Tau signal that falls inside the nucleus mask, computed as the sum of Tau intensities in mask_nucleus divided by the sum in mask_cell. Although Tau is typically cytosolic, pathological conditions may show aberrant intranuclear Tau, so higher values could indicate abnormal nuclear invasion. The feature is defined as NaN or zero when no nucleus mask is present. |

| | | | |
|---|---|---|---|
| 264 | **cellwide tau granularity high frequency fraction** | **code** | This feature computes the proportion of Tau signal power residing in high-frequency components by applying a Laplacian-of-Gaussian filter at a small spatial scale to the Tau MIP and then taking the ratio of the sum of squared filtered values to the sum of squared original intensities, restricted to the cell mask. It quantifies fine-grained punctate or beaded texture versus smooth diffuse signal within the neuron, which is relevant for distinguishing early granular pathology from homogeneous Tau accumulation. A higher fraction suggests more punctate, granular Tau morphology in patient cells. |
| 265 | **tau beaded neurite fraction visual** | **vlm** | This feature visually estimates the fraction of neurite length within a cell that exhibits a beaded (string-of-beads) Tau pattern rather than smooth linear Tau along neurites, using the 2D Tau MIP and the neurite portions of mask_filament and mask_bundle. It directly reflects the presence of discrete Tau puncta along neurites indicative of pathological fragmentation or droplet formation along processes. The output is a scalar score between 0 and 1 summarizing how much of the neurite network appears beaded. |
| 266 | **tau droplet fraction of cell tau** | **code** | This feature measures the fraction of total Tau signal within the cell that is contained in droplet-like regions, computed as the sum of Tau intensities inside mask_droplet divided by the sum inside mask_cell. It captures how much of the Tau pool has condensed into punctate or droplet-like aggregates, which is a hallmark of pathological phase-separated condensates. Larger values indicate a higher degree of dropletization relative to diffuse cytosolic Tau. |
| 267 | **tau soma to cell intensity ratio** | **code** | This feature computes the mean Tau intensity inside the nucleus-aware soma region (cell mask minus nucleus mask) divided by the mean Tau intensity over the entire cell mask. It captures the relative enrichment of Tau in the soma cytoplasm versus the whole cell, which is expected to increase when Tau relocates from axons/neurites into the soma in pathological states. Values >1 indicate soma-biased Tau distribution, while values near or below 1 indicate a more even or neurite-dominated distribution. |
| 268 | **tau cell body to neurite intensity ratio** | **code** | This feature computes the mean Tau intensity inside the cell mask but outside the nucleus (approximating soma) divided by the mean Tau intensity in the neuritic compartment, defined as the cell mask minus the soma (eroded cell mask) and nucleus. It quantifies the redistribution of Tau from neurites into the soma, which is expected to increase with pathological Tau mislocalization in patient neurons on 2D Tau MIP images. |
| 269 | **tau droplet somatic occupancy fraction** | **code** | This feature measures the fraction of the soma area occupied by Tau-positive droplets, computed as the area of droplet mask pixels that lie inside the cell mask but outside the nucleus, divided by the somatic area. It quantifies how extensively Tau droplets invade the soma, distinguishing sparse, focal inclusions from widespread dropletization. Larger occupancy suggests more severe droplet-based Tau condensation within the somatic compartment. |
| 270 | **tau droplet area fraction in cell** | **code** | This feature measures the fraction of the cell area occupied by Tau-positive droplet regions, computed as the total area of mask_droplet intersected with mask_cell divided by the area of mask_cell. It quantifies how much of the soma and processes are filled by compact Tau droplets, which is relevant to assessing the burden of intracytoplasmic Tau inclusions at the single-cell level. |
| 271 | **normalized tau droplet area fraction** | **code** | This feature measures the fraction of the cell area occupied by Tau-positive droplets, defined as the total area of droplet mask pixels divided by the cell mask area. It quantifies the burden of punctate Tau aggregation normalized to cell size, allowing comparison across cells with different sizes or imaging fields. A higher fraction indicates more extensive droplet-like aggregation within the neuron. |
| 272 | **tau droplet burden per cell area** | **code** | This feature measures the total Tau-positive droplet area (in pixels) normalized by the total cell area within the cell mask on the 2D MIP. Droplets are defined by the user-provided droplet mask; their area fraction within the cell quantifies the burden of compact Tau condensates relative to cell size. Increased normalized droplet burden is expected in cells with higher degrees of Tau condensation or phase-separated aggregates. |
| 273 | **tau droplet load per cell area** | **code** | This feature measures the total Tau signal contained in all segmented Tau-positive droplets (sum of pixel intensities within mask_droplet) normalized by the total area of the cell mask. It quantifies how much Tau is sequestered into discrete droplet-like condensates per unit cytoplasmic area. Increased values may reflect enhanced phase separation or stress-granule-like Tau condensation in diseased neurons. |
| 274 | **tau droplet coverage in cell** | **code** | This feature measures the fraction of the cell area occupied by Tau-positive droplets by taking the area of the droplet mask within the cell mask divided by the total cell mask area. It quantifies the overall burden of discrete Tau condensates or stress-granule-like droplets in each neuron and is relevant to Tau aggregation and phase separation phenotypes. If no droplet mask is present, the feature is set to zero or NaN according to a predefined policy. |
| 275 | **tau droplet cell relative area** | **code** | This feature measures the fraction of the cell area occupied by Tau droplets by taking the total area of mask_droplet.tif intersected with mask_cell.tif and dividing by the total mask_cell area. It summarizes the burden of droplet-like Tau condensates per neuron on the 2D MIP, which is relevant to quantifying condensation-dominated pathology across patient cells. |
| 276 | **tau intranuclear hotspot fraction** | **code** | This feature assesses whether there are focal Tau hotspots inside the nucleus, which would be unexpected in healthy neurons. Within mask_nucleus.tif, a relative threshold (e.g., top 5% of nuclear Tau intensity) is applied and the fraction of nuclear area exceeding this threshold is computed. Higher values indicate abnormal intranuclear Tau clustering or leakage. |
| 277 | **tau intracellular granularity at multiple scales** | **code** | Within the cell mask, this feature applies a bank of Laplacian-of-Gaussian filters at several spatial scales and computes, for each scale, the normalized variance of the filter responses, then aggregates them (e.g., as a weighted sum) into a single scalar. This multi-scale granularity score captures how punctate versus diffuse the Tau distribution is across nanocluster-sized, droplet-sized, and larger inclusion scales. Higher values indicate richer multiscale granule structure consistent with stress granules, droplets, and fibrillar aggregates. |
| 278 | **tau droplet nuclear proximity index** | **code** | For each Tau droplet in mask_droplet.tif, this feature computes the minimum Euclidean distance from the droplet centroid to the nuclear mask, and then reports the mean inverse distance over all droplets (1 / (distance + ε)). Higher values indicate that droplets cluster near or within the perinuclear region, while lower values reflect droplets located in distal processes or cell periphery. This index is intended to reflect cell body–centered Tau aggregation versus more neuritic aggregation. |

| | | | |
|---|---|---|---|
| 279 | **tau cytoplasmic texture coarseness** | **code** | Restricted to the cytoplasmic region (mask_cell minus mask_nucleus), this feature computes a multi-scale coarseness descriptor such as Tamura coarseness or the average local variance of Tau intensity over several window sizes, averaged within the cytoplasm. It captures the transition from fine, smooth Tau distribution to coarse, patchy patterns as droplets, filaments, and bundles accumulate. Increased coarseness is expected in cells with advanced Tau aggregation. |
| 280 | **tau axon guard fraction** | **code** | This feature estimates the fraction of total cellular Tau intensity that lies in a thin inner band along the cell boundary (e.g., a 2–3 pixel-wide erosion of the cell mask) relative to the total Tau intensity within the whole cell mask. It captures how much Tau is concentrated at the periphery where axon and neurite origins tend to reside in MIPs, serving as a proxy for axon-dominant vs soma-centered Tau localization. Lower values may indicate loss of normal axonal enrichment and pathological redistribution toward the soma interior. |
| 281 | **soma tau texture heterogeneity** | **code** | This feature measures fine-scale heterogeneity of Tau within the soma by computing a gray-level co-occurrence matrix (GLCM) based contrast or entropy metric on the Tau channel restricted to the cell mask but excluding nucleus and filament/bundle regions, and normalizing by the somatic Tau mean intensity. Higher values indicate a more granular, rough texture consistent with intracytoplasmic aggregates, whereas lower values indicate smoother, diffuse somatic Tau distribution. |
| 282 | **tau droplet fragmentation index** | **code** | Within the droplet mask, this feature quantifies how fragmented Tau-positive droplets are by computing (number of connected droplet objects) divided by (total droplet mask area). Higher values indicate many small Tau droplets rather than fewer larger condensates, reflecting changes in condensate nucleation versus coarsening dynamics. |
| 283 | **tau texture granularity in droplet free cytoplasm** | **code** | Within the cell mask but excluding any droplet, filament, and bundle masks, compute a mid-scale granularity or bandpass-filtered variance (e.g., via Laplacian-of-Gaussian at a defined sigma) of Tau intensity. This quantifies subresolution or diffuse Tau texture in droplet-free cytoplasm on the MIP, capturing early diffuse pathology distinct from resolved aggregates. |
| 284 | **axon like tau gradient strength** | **code** | Within the cell mask, this feature approximates an axon-like process by finding the longest skeleton path in the filament+bundle masks and measuring the monotonicity of Tau intensity along that path. Tau intensity is sampled along the skeleton and a linear fit of intensity versus arc length is performed; the feature is the signed $R^2$-weighted slope magnitude, with high negative values indicating a smooth proximal-to-distal decay reminiscent of healthy axonal Tau gradients and low or positive values suggesting gradient disruption. |
| 285 | **tau background cytoplasmic granularity scale2** | **code** | Restricted to the non-droplet, non-filament, non-bundle cytoplasmic Tau signal (mask_cell minus all provided sub-masks), this feature calculates a Laplacian-of-Gaussian granularity measure at an intermediate spatial scale (e.g., sigma corresponding to ~0.3–0.5 µm) and averages the filter response. It captures mid-scale granular puncta in the diffuse cytoplasm, distinguishing smooth diffuse Tau from finely granular stress-granule-like or oligomeric patterns. |
| 286 | **tau soma texture coarseness** | **code** | Within each cell's soma region (mask_cell minus all neuritic filament and bundle masks if those are available), this feature computes a multi-scale local variance (e.g., average variance in sliding windows of several sizes) and aggregates them into a coarseness index, then averages across cells. Healthy somatic Tau is expected to be low and relatively smooth, while pathological inclusions yield coarse, high-variance textures corresponding to dense NFTs and irregular fibril networks. This texture-based coarseness scalar complements simple intensity metrics by being sensitive to sub-soma heterogeneity. |
| 287 | **tau intracellular granularity multiscale** | **code** | Within the cell mask, this feature quantifies Tau granularity by computing the variance of the Laplacian-of-Gaussian–filtered Tau image at two or more spatial scales (e.g., sigma corresponding to small puncta vs larger inclusions) and then summing these variances. It captures the presence of multi-scale granular structure in Tau signal, from small droplets to larger aggregates, beyond what simple mean intensity reveals. Higher values indicate more pronounced punctate or aggregated patterns consistent with stress granules or Tau inclusions. |
| 288 | **droplet tau fraction and clumpiness** | **code** | This feature measures the fraction of total cellular Tau signal confined to droplet regions and modulates it by a clumpiness index. First, the sum of Tau intensity within mask_droplet is divided by the sum within mask_cell; then this fraction is multiplied by the Gini coefficient of droplet-region intensities, yielding higher values when Tau is both droplet-enriched and concentrated into a few very bright droplets. This aims to distinguish diffuse low-grade droplet formation from highly aggregated, pathologic Tau droplets in 2D MIPs. |
| 289 | **tau cellwide granularity lowpass minus highpass** | **code** | This feature captures the balance between coarse (large-scale) and fine (small-scale) Tau granularity over the entire cell. Within mask_cell.tif, Gaussian-smoothed versions of the Tau MIP at two spatial scales (e.g., $\sigma \approx 0.3$ µm and $\sigma \approx 1.0$ µm) are produced, and the global variance at the coarse scale minus the variance at the fine scale is computed. Larger values indicate dominance of broad, diffuse accumulations over fine puncta, consistent with progression from oligomers to larger inclusions. |
| 290 | **nuclear to cytoplasmic tau intensity gradient** | **code** | This feature computes the radial gradient of Tau intensity from the nucleus boundary outward into the cytoplasm by sampling concentric distance bins from the nuclear mask up to the cell boundary and fitting a line to intensity versus distance. The fitted slope, normalized by mean cellular intensity, quantifies whether Tau is concentrated near the nucleus (positive or flat slope) or depleted around it (negative slope), probing nuclear-adjacent accumulation versus peripheral localization. |
| 291 | **tau peripheral filament density gradient** | **code** | This feature captures how Tau filament density changes from the cell center toward the periphery, reflecting potential distal accumulation. Within mask_cell.tif, a distance transform from the cell centroid is used to divide the cell into concentric radial bins (e.g. 4–6 bins); for each bin, the density of filament pixels from mask_filament.tif (filament area divided by bin area) is computed, and a linear regression of density versus normalized radius is fit; the feature is the slope of this regression. A positive slope indicates enrichment of filaments in the outer cell regions relative to the center. |
| 292 | **tau filament length density in cell** | **code** | This feature measures the total skeletonized filament length within the cell, normalized by the cell area, using mask_filament and mask_cell on the Tau MIP. It approximates how densely Tau filaments occupy the cell, with higher values indicating abundant filamentous Tau structures relative to cell size. This is relevant to distinguishing filament-rich NFTs from more droplet-like aggregation phenotypes. |

| | | | |
|---|---|---|---|
| 293 | **tau bundle soma coverage ratio** | **code** | This feature measures how extensively Tau bundles cover neuronal cell bodies by computing, for each cell, the fraction of soma area (mask_cell.tif minus mask_nucleus.tif) that overlaps with mask_bundle.tif, then averaging these fractions over all cells. Larger coverage indicates that bundles are invading or enveloping somata, a phenotype reminiscent of neurofibrillary tangle formation within cell bodies. |
| 294 | **tau subcellular radial variance** | **code** | Within each cell, this feature measures how Tau intensity varies as a function of radial distance from the nuclear centroid to the cell boundary by computing the variance of mean Tau intensity across a fixed number of concentric radial bins. High values indicate strong radial heterogeneity (e.g., perinuclear condensates or edge-enriched filaments), whereas low values correspond to more uniform distributions. This captures loss of normal axon-distal gradients when only a 2D MIP is available. |
| 295 | **axon like filament length density** | **code** | Within the cell mask, this feature estimates the density of long, thin Tau-positive filaments that resemble axonal segments by selecting filament objects from mask_filament with high elongation (major/minor axis ratio above a threshold) and length above a minimum, then summing their skeleton lengths and normalizing by cell area. Higher values correspond to preservation or proliferation of elongated axon-like Tau structures, while lower values may indicate fragmentation or loss of continuous Tau-decorated neurites. |
| 296 | **tau filament linear density within cell** | **code** | This feature measures the total filament skeleton length per unit cell area by skeletonizing the filament mask (mask_filament) and dividing the total skeleton length by the cell mask area. It quantifies how densely Tau-positive filaments occupy the cell in 2D MIP, reflecting the extent of fibrillar Tau organization versus more compact aggregates. Higher values indicate a more extended filamentous network, while lower values suggest loss of filaments or consolidation into droplets/bundles. |
| 297 | **tau filament length per cell area** | **code** | This feature computes the total skeletonized length of Tau filaments (from mask_filament.tif after thinning) divided by the cell area, giving a length-per-area density of filamentous Tau. It reflects the extent of fibrillar Tau pathology normalized for cell size, distinguishing filament-rich from filament-poor cells. |
| 298 | **tau filament skeleton length per cell** | **code** | This feature skeletonizes the Tau filament mask (mask_filament.tif) and sums the skeleton length in pixels, then normalizes by the total cell area. It captures how much filamentous Tau structure exists per unit cell area, reflecting the extent of microtubule-aligned or fibrillar Tau organization in the 2D MIP. |
| 299 | **tau filament endpoints per cell area** | **code** | After skeletonizing the filament mask within each cell, this feature counts the number of filament endpoints (vertices with degree 1) and divides by the cell area, then averages this density across cells. It reflects how fragmented or beaded the filament network is: many endpoints per area correspond to short, broken filaments or bead-like chains, while fewer endpoints indicate longer continuous fibrils. This is relevant for differentiating early fragmented pathology from later continuous filament bundles. |
| 300 | **tau droplet density per cell area** | **code** | Counts the number of discrete droplet objects normalized by the total cell area. This provides a density metric for aggregation load independent of cell size. |
| 301 | **tau droplet count per cell area** | **code** | This feature counts the number of distinct Tau-positive droplets within the droplet mask (mask_droplet.tif) and normalizes by the cell area from mask_cell.tif, yielding a droplet density per unit cell area. It reflects the abundance of discrete Tau droplet-like aggregates, which may correspond to stress-granule-like or early oligomeric accumulations in patient neurons. The feature is defined on the 2D Tau MIP using connected-component analysis of the droplet mask. |
