## Supplementary feature list 4 for "Biologically grounded cell profiling across microscopy modalities"

### MorphAgent Tau feature catalog

400 MorphAgent features for Tau SIM imaging of SH-SY5Y cells (code + VLM). Organized as the Fig. 5c three-level hierarchy: **Morphology-defined biological process** → **Subcellular architecture** → **Feature**. Process names follow the current Fig. 5d labels; each process includes the summary description from the functional-clustering table (row-matched). Architecture names follow the original scale-based vocabulary, with approximate structural scales from the size-based clustering table. 363 features are assigned to the hierarchy; 37 additional catalog features are listed at the end.

| **Level 1** Morphology-defined biological process | **Level 2** Subcellular architecture | **Level 3** Feature name · method · description |
|---|---|---|

#### Morphology-defined biological process · Tau mislocalization

**Description.** This feature class quantifies enrichment, exclusion, annular or sectorized distribution of Tau signal around the soma, perinuclear zone, and nuclear boundary, as well as intensity differences across the nucleocytoplasmic interface. Biologically, it reflects redistribution of Tau from its normal axonal/microtubule-associated localization toward somatic and perinuclear compartments.

52 features · 4 subcellular architectures

##### Subcellular architecture · Nuclear and perinuclear Tau architecture (27 features)

**Approximate scale.** Organized around a nucleus ~5-10 um in diameter; the perinuclear annulus/shell is typically sub-um to several um thick, with circumferential length ~10-30+ um.

| # | Feature name | Method | Description |
|---|---|---|---|
| 1 | cytoplasm nuclear tau enrichment ratio | code | Measures the ratio of mean Tau fluorescence in the cytoplasm to mean Tau fluorescence in the nucleus, using the 2D Tau MIP and the provided cell and nucleus masks. The cytoplasm is defined as mask_cell minus mask_nucleus, with a small numerical offset to avoid division by zero. This captures abnormal cytoplasmic Tau accumulation while controlling for nuclear/background fluorescence and may reflect Tau mislocalization severity. |
| 2 | nuclear ring tau azimuthal heterogeneity | code | Measures how uneven Tau signal is around the nucleus by sampling Tau intensity in a narrow cytoplasmic annulus immediately outside the nuclear mask and binning it by angle around the nuclear centroid. The scalar is the robust dispersion of angular-bin intensities, such as IQR divided by median. It complements nuclear-boundary gradient features by focusing on tangential asymmetry of perinuclear Tau accumulation rather than radial edge sharpness. |
| 3 | perinuclear low tau crescent eccentricity | code | Measures the shape anisotropy of low-Tau regions immediately surrounding the nucleus. Within a perinuclear cytoplasmic annulus, pixels below a local Tau percentile are identified, connected components are combined or summarized, and their intensity-free inertia eccentricity is computed. A high value indicates crescent-like nuclear shadowing or one-sided Tau exclusion, whereas a low value indicates more isotropic low-Tau annular gaps. |
| 4 | perinuclear tau annular gap length cv | code | Measures how unevenly Tau-depleted angular gaps are distributed around the nucleus within a narrow perinuclear cytoplasmic annulus. The annulus is divided into angular bins, bins below a robust Tau threshold are grouped into contiguous low-Tau runs, and the coefficient of variation of run lengths is returned. This captures whether perinuclear exclusion occurs as many small gaps or a few large shadow-like sectors, distinct from simply measuring the fraction of shadowed angles. |
| 5 | perinuclear tau annulus radial phase lag | code | Construct concentric cytoplasmic annuli from the nuclear boundary toward the cell edge and identify the angular sector with maximum Tau signal in each annulus. The feature is the circular dispersion of angular phase shifts between neighboring annuli. It captures whether perinuclear and outer cytoplasmic Tau hotspots are radially aligned, twisted, or spatially decoupled. |
| 6 | perinuclear tau cap angular sharpness | code | Measures how narrowly concentrated the strongest perinuclear Tau signal is around the nucleus in the 2D MIP. Using the nucleus and cell masks, Tau intensity is sampled in a thin cytoplasmic annulus outside the nucleus, binned by angle, and the angular full-width or circular peakedness of the brightest cap is converted to a scalar sharpness score. This complements perinuclear cap strength by distinguishing a broad circumferential enrichment from a tight asymmetric Tau cap. |
| 7 | perinuclear tau cap resultant strength | code | This feature analyzes Tau intensity in a narrow cytoplasmic band surrounding the nucleus and computes the circular resultant length of angular Tau mass around the nuclear centroid. Values near zero indicate Tau distributed evenly around the nucleus, while high values indicate a one-sided perinuclear Tau cap. It complements nuclear-boundary fragmentation metrics by measuring directional concentration of perinuclear Tau rather than patch count. |
| 8 | perinuclear tau closing gap recovery | code | Measures how much fragmented high Tau signal in the perinuclear cytoplasm can be reconnected by a small morphological closing operation. A locally thresholded high-Tau mask is formed in a perinuclear annulus, closed with a disk-shaped structuring element, and the added area is divided by the original high-Tau annular area. Higher values indicate broken ring-like or mesh-like Tau patterns with small gaps near the nucleus. |

| # | Feature name | Method | Description |
|---|---|---|---|
| 9 | perinuclear tau enrichment ratio | code | Measures the ratio of mean Tau intensity in the inner cytoplasmic zone near the nucleus to mean Tau intensity in the more distal cell region, using the 2D MIP Tau image with mask_cell and mask_nucleus. The zones can be defined by distance transform within the cell mask excluding the nucleus, for example comparing the closest third versus farthest third of cytoplasmic pixels. This captures pathological somatic/perinuclear Tau accumulation while normalizing against distal neuritic Tau signal. |
| 10 | perinuclear tau ring width halfmax | code | This feature estimates the radial thickness of a Tau-enriched perinuclear ring by computing the radial Tau profile outward from the nucleus and measuring the width over which intensity remains above half of the local perinuclear peak-to-cytoplasm contrast. A broad ring may indicate diffuse somatic Tau accumulation, whereas a narrow ring may indicate sharper compartmentalization. It is distinct from angular perinuclear features because it focuses on radial spread rather than azimuthal phase or heterogeneity. |
| 11 | perinuclear tau shadow sector fraction | code | Measures whether the perinuclear cytoplasm contains a broad angular sector depleted of Tau signal. In a narrow cytoplasmic ring around the nucleus, angular sectors are scored as low Tau if their median intensity is below the cytoplasmic median, and the largest contiguous low-Tau sector is divided by the full circle. This may reflect asymmetric nuclear exclusion, aggregate displacement, or polarized somatic Tau accumulation. |
| 12 | perinuclear tau shell angular mad | code | Measures robust angular heterogeneity of Tau in the immediate perinuclear cytoplasmic shell. The shell is defined as the inner portion of cytoplasm adjacent to the nuclear mask, divided into angular sectors around the nuclear centroid, and the feature is the median absolute deviation of sector Tau intensity normalized by the sector median. This captures asymmetric perinuclear Tau caps or crescent-like accumulation without reducing the pattern to a simple mean enrichment ratio. |
| 13 | perinuclear tau shell enrichment | code | Computes the mean Tau intensity in a cytoplasmic shell immediately surrounding the nucleus divided by the mean Tau intensity in the remaining cytoplasm. The shell can be generated by dilating the nucleus within the cell mask on the 2D MIP. Perinuclear Tau enrichment may capture early somatic accumulation or NFT-like Tau organization around a nuclear exclusion zone. |
| 14 | perinuclear tau signal fraction | code | Measures the fraction of total Tau fluorescence inside the cell mask that lies in a perinuclear cytoplasmic annulus, defined as pixels inside mask_cell, outside mask_nucleus, and within a fixed dilation distance from the nuclear boundary. The input is the 2D Tau MIP, so the feature summarizes projected somatic/perinuclear Tau accumulation rather than 3D localization. Increased values may reflect abnormal somatic Tau enrichment, a key signature of Tau mislocalization. |
| 15 | tau perinuclear peak offset normalized | code | Measures where the cytoplasmic Tau radial intensity profile reaches its maximum relative to the nuclear boundary and cell boundary. The cytoplasm is binned by normalized distance from the nucleus outward, the mean or robust percentile Tau intensity is computed per bin, and the peak bin distance is returned as a normalized scalar. This distinguishes tight perinuclear accumulation from broader cytoplasmic or peripheral Tau redistribution, relevant to somatic Tau mislocalization and aggregate positioning. |
| 16 | tau puncta perinuclear enrichment ratio | code | Detects small bright Tau puncta in the cytoplasm and compares puncta density in an inner perinuclear shell with puncta density in the remaining cytoplasm. The scalar is an area-normalized inner-to-outer puncta ratio, optionally using integrated puncta intensity instead of count when puncta segmentation is noisy. It may reflect stress-granule-like or early aggregate-like Tau recruitment near the soma and nucleus. |
| 17 | nuclear bipolar tau cap score | vlm | Measures whether Tau signal forms two opposing enriched caps or lobes around the nucleus rather than a single crescent or uniform perinuclear ring. A high value indicates a bipolar perinuclear Tau arrangement with enrichment on roughly opposite nuclear sides. This may reflect polarized cytoplasmic trafficking, nuclear displacement forces, or asymmetric aggregate compartmentalization. |
| 18 | nuclear clearance shadow orientation score | vlm | Scores whether the nucleus creates an asymmetric Tau-poor shadow or clearance zone that is directionally biased within the surrounding cytoplasmic Tau field. A high score indicates a clear one-sided nuclear-adjacent depletion or shadowing pattern rather than uniform perinuclear exclusion. This may reflect spatial displacement of Tau aggregates or cytoplasmic organization around the nucleus. |
| 19 | perinuclear cytoplasm thickness imbalance score | vlm | Scores how uneven the cytoplasmic space is around the nucleus within the segmented cell mask. A high score indicates that the nucleus lies close to one cell boundary or that cytoplasm is compressed on one side and expanded on another, regardless of whether the displacement is caused by Tau aggregates. This complements Tau-mass displacement features by focusing on cell-and-nucleus geometry rather than the location of bright Tau signal alone. |
| 20 | perinuclear tau band width irregularity score | vlm | Scores how uneven the apparent thickness of Tau signal is around the nucleus within the 2D MIP cell mask. A high value indicates that the perinuclear Tau band alternates between thick swollen sectors and thin or absent sectors, rather than forming a uniform halo. This may reflect asymmetric somatic Tau accumulation or localized aggregate growth near the nucleus. |
| 21 | perinuclear tau crescent bias score | vlm | Estimates how strongly Tau fluorescence forms a one-sided crescent or cap around the nucleus within the 2D MIP cell mask. This differs from generic nuclear displacement by focusing on asymmetric perinuclear Tau wrapping rather than centroid shift, and may reflect polarized somatic aggregate growth or cytoplasmic crowding around the nucleus. |

| # | Feature name | Method | Description |
|---|---|---|---|
| 22 | perinuclear tau crowding score | vlm | A continuous score measuring whether Tau signal forms dense, asymmetric, or ring-like accumulations immediately around the nucleus in the 2D MIP. High values indicate cytoplasmic Tau crowding, nuclear encasement, or apparent displacement by nearby aggregates. This may reflect advanced somatic Tau aggregation and altered subcellular organization. |
| 23 | perinuclear tau enrichment score | vlm | Measures the visual prominence of Tau fluorescence in the cytoplasmic rim surrounding the nucleus in the 2D Tau MIP. The scalar score should be high when Tau forms a bright annular, crescent-like, or asymmetric perinuclear accumulation compared with the rest of the cell. This may capture early or localized somatic Tau redistribution around the nucleus. |
| 24 | perinuclear tau ring completeness score | vlm | Scores how completely Tau fluorescence forms a continuous annular band around the nucleus in the 2D MIP, using the nucleus and cell masks to focus on perinuclear cytoplasm. Unlike a one-sided crescent measure, this feature emphasizes circumferential continuity and uniformity of the perinuclear Tau ring. A high score may reflect somatic Tau redistribution and perinuclear aggregation patterns linked to neuronal stress or Tau handling pathways. |
| 25 | perinuclear tau sector compaction score | vlm | Scores whether perinuclear Tau is concentrated into one or a few compact angular sectors around the nucleus instead of forming a diffuse or uniform perinuclear distribution. The VLM should inspect the Tau MIP within the cytoplasmic region immediately outside the nucleus mask and estimate the degree of sector-like compaction. This may capture pathological perinuclear Tau crowding linked to somatic accumulation and nuclear stress. |
| 26 | perinuclear tau sector edge synchrony score | vlm | Measures whether bright perinuclear Tau sectors have visually aligned, coherent radial edges around the nucleus in the 2D Tau MIP. A high score indicates that multiple perinuclear Tau patches appear organized into coordinated sector boundaries, while a low score indicates scattered or unrelated perinuclear deposits. This captures the organization of perinuclear Tau domains rather than their total enrichment or compaction. |
| 27 | perinuclear to peripheral grain size transition score | vlm | Scores whether Tau texture changes in apparent grain size from the perinuclear cytoplasm to the outer cell periphery in the 2D MIP. High values indicate a clear transition, such as fine perinuclear granules becoming coarser peripheral clumps or the reverse, rather than uniform texture across the cell. This complements radial intensity features by focusing on texture scale rather than brightness alone. |

#### Subcellular architecture · Nuclear and nucleocytoplasmic compartmentalization scale (9 features)

**Approximate scale.** Typical mammalian nucleus diameter approximately 5-10 um.

| # | Feature name | Method | Description |
|---|---|---|---|
| 28 | cytoplasmic tau nuclear exclusion ratio | code | Measures the mean Tau intensity in the cell mask excluding the nucleus divided by the mean Tau intensity inside the nucleus mask, using the 2D Tau MIP. Tau is expected to be largely cytoplasmic, and stronger cytoplasmic enrichment relative to the nucleus can reflect somatic Tau accumulation while controlling for local image brightness. This feature is robustly scalar and uses the provided cell and nucleus masks. |
| 29 | nuclear exclusion front sharpness cv | code | Sample short outward radial profiles from the nuclear boundary into the cytoplasm and identify the strongest local Tau intensity rise along each profile. The feature is the coefficient of variation of these boundary-front sharpness values around the nucleus. It measures whether Tau exclusion or accumulation near the nucleus is uniform or occurs in localized sharp fronts, relevant to perinuclear Tau redistribution. |
| 30 | nuclear tau exclusion index | code | Measures how strongly Tau signal is excluded from the nucleus by comparing mean Tau intensity inside the nucleus with mean Tau intensity in the surrounding cytoplasm. The value can be computed as cytoplasmic mean divided by nuclear mean after background correction. Abnormal nuclear leakage or poor cytoplasmic compartmentalization may reflect altered Tau localization or cell state. |
| 31 | nucleus to edge tau onoff transition density | code | Casts radial rays from the nuclear centroid to the cell boundary and samples Tau intensity along each ray after robust binarization into Tau-high and Tau-low states. The feature is the average number of high-to-low or low-to-high state transitions per unit cytoplasmic path length. It captures radial fragmentation and alternating Tau bands along nucleus-to-membrane trajectories, complementing total variation by counting discrete on/off structural changes. |
| 32 | tau nuclear clearance depth integral | code | This feature measures the integrated Tau intensity deficit immediately outside the nucleus compared with the broader cytoplasmic Tau baseline. It samples a normalized perinuclear zone and integrates how far below the cytoplasmic median the Tau signal remains, producing a scalar index of nuclear-adjacent Tau depletion or exclusion. It complements sharpness-based nuclear exclusion measures by quantifying the depth and spatial extent of the low-Tau clearance zone. |
| 33 | droplet nuclear avoidance zone score | vlm | Measures whether Tau droplets are visually excluded from a clear zone immediately surrounding the nucleus. A high score indicates that droplets occupy cytoplasm farther from the nuclear boundary while leaving a relatively droplet-free perinuclear gap. This captures droplet spatial organization relative to nuclear geometry, complementary to droplet abundance or clustering. |
| 34 | nuclear tau exclusion clarity score | vlm | Scores how clearly Tau signal is excluded from the nuclear mask relative to the surrounding cytoplasm in the 2D MIP. High values indicate a sharp dark nucleus surrounded by Tau-positive cytoplasm, while low values indicate weak cytoplasmic contrast, apparent nuclear overlap, or obscuration by dense Tau signal. This may reflect somatic Tau accumulation patterns and nuclear compression or masking without duplicating perinuclear crowding. |

| # | Feature name | Method | Description |
|---|---|---|---|
| 35 | nuclear tau intrusion score | vlm | Scores the degree to which Tau fluorescence appears to intrude into the nuclear mask region in the 2D MIP. Low values indicate strong nuclear exclusion, while high values indicate visible Tau-positive islands, haze, or projections within the segmented nucleus. This is distinct from boundary contact because it focuses on apparent intranuclear Tau signal rather than pressure at the nuclear edge. |
| 36 | nucleus to edge tau clearance asymmetry score | vlm | Measures whether Tau-negative cytoplasmic clearance space between the nucleus and cell boundary is unevenly distributed around the nucleus. A high score indicates that one side of the nucleus has a broad Tau-poor corridor or void while the opposite side is crowded by Tau structures. This may capture asymmetric cytoplasmic reorganization or inclusion pressure without directly duplicating nuclear displacement metrics. |

##### Subcellular architecture · Cell boundary, cortical, peripheral-shell, and extracellular-neighborhood scale (8 features)

**Approximate scale.** Set against a whole-cell diameter of ~10-100 um; the measured region is typically a thin band or boundary arc inside/outside the cell edge.

| # | Feature name | Method | Description |
|---|---|---|---|
| 37 | aggregate nuclear proximity weighted intensity | code | Measures whether segmented Tau aggregates are preferentially located near the nucleus by weighting aggregate Tau intensity by inverse distance to the nuclear boundary. The aggregate region is the union of available droplet, filament, and bundle masks within mask_cell, evaluated on the 2D Tau MIP. Higher values indicate perinuclear or soma-associated aggregate burden, which is relevant to neurofibrillary tangle-like somatic Tau pathology. |
| 38 | nuclear boundary tau gradient peak | code | Quantifies the sharp Tau intensity transition across the nuclear boundary in the 2D MIP by comparing Tau signal in a thin cytoplasmic rim just outside the nucleus to a thin rim just inside the nucleus. Strong positive values indicate intense perinuclear Tau accumulation or exclusion of Tau from the nucleus, both potentially related to somatic aggregate organization. This differs from a simple cytoplasm/nucleus ratio because it focuses specifically on the immediate nuclear interface. |
| 39 | nuclear cytoplasm tau interface contrast cv | code | Samples paired narrow bands immediately inside and outside the nuclear boundary and computes the coefficient of variation of the local Tau contrast around the nucleus. High values indicate patchy, asymmetric perinuclear Tau accumulation or uneven nuclear exclusion, while low values indicate a more uniform nuclear-cytoplasmic interface. This may capture nuclear-adjacent Tau redistribution patterns not summarized by mean perinuclear enrichment alone. |
| 40 | nuclear tau aggregate contact arc fraction | code | This feature estimates the fraction of the nuclear boundary that lies adjacent to high-intensity Tau aggregate signal. High-Tau objects are detected within the cell, and the nuclear perimeter pixels within a small distance of those objects are counted as contacted arc length. It may reflect nuclear encasement, perinuclear aggregate accumulation, or soma-filling tangle-like pathology. |
| 41 | nuclear tau boundary patch fragmentation | code | Detects high-Tau pixels adjacent to the nuclear boundary and counts the number of separated contact patches along the boundary, normalized by nuclear perimeter. Unlike total contact arc fraction, this feature emphasizes whether nuclear-associated Tau is continuous or fragmented into many discrete patches. It may reflect differences between smooth perinuclear accumulation and punctate aggregate contact with the nucleus. |
| 42 | filament nuclear tangential wrapping score | vlm | Scores the extent to which Tau filaments curve tangentially around the nucleus instead of pointing radially toward or away from it. A high value indicates arc-like or crescent-shaped filaments following the nuclear contour. This captures a geometric relationship between filament orientation and nuclear position in the 2D MIP. |
| 43 | nuclear boundary tau contact pressure score | vlm | Measures the apparent extent to which bright Tau inclusions directly contact, hug, or indent the nuclear boundary in the 2D MIP. This is complementary to nuclear Tau exclusion and nuclear displacement because it focuses on local aggregate-nucleus interface pressure rather than global nuclear position or intensity inside the nucleus. |
| 44 | nuclear proximity tau object size gradient score | vlm | Measures whether Tau-positive objects such as droplets, short filaments, or compact aggregates systematically change size with distance from the nucleus. A high score indicates a visible nuclear-proximity gradient in object scale, independent of whether objects are more abundant near or far from the nucleus. This may capture perinuclear maturation or spatially regulated Tau aggregation in the single-cell MIP. |

##### Subcellular architecture · Whole-cell geometry, global polarity, and spatial-distribution scale (8 features)

**Approximate scale.** Approximately 10-100+ um; larger if the full neurite arbor is included in the 2D projection.

| # | Feature name | Method | Description |
|---|---|---|---|
| 45 | tau centroid to nucleus axis alignment | code | This feature measures the alignment between the vector from the nucleus centroid to the Tau intensity-weighted centroid and the major axis of the segmented cell. The output can be the absolute cosine of this angle, optionally weighted by the normalized Tau-centroid displacement magnitude. It captures whether Tau redistribution is directed along the main neuritic/cellular axis rather than merely being off-center. |
| 46 | tau half mass distance from nucleus | code | Computes the normalized distance from the nuclear boundary at which 50% of cytoplasmic Tau intensity mass is accumulated, using the 2D MIP Tau image and nucleus/cell masks. A small value indicates perinuclear Tau concentration, while a larger value indicates Tau spreading toward the cell periphery or neurites. This captures a continuous redistribution pattern distinct from simple radial entropy or gradient slope. |

| # | Feature name | Method | Description |
|---|---|---|---|
| 47 | tau intensity centroid offset from nucleus | code | Measures the distance between the Tau intensity-weighted centroid within the cell mask and the nucleus centroid, normalized by the square root of cell area. A larger offset indicates asymmetric Tau localization, such as polarized enrichment in a process or eccentric aggregate burden. This captures spatial redistribution of Tau in the 2D single-cell MIP rather than total brightness alone. |
| 48 | tau mass nucleus cell axis third moment | code | Quantifies asymmetric Tau mass placement along the anatomical axis from the nucleus centroid to the whole-cell centroid. Each cytoplasmic pixel is projected onto this axis and weighted by Tau intensity, and the normalized third central moment of the projected Tau mass distribution is reported. This captures directional somatic polarization or aggregate displacement not captured by simple centroid offset alone. |
| 49 | tau nucleus distance intensity mutual information | code | Computes the mutual information between binned Tau intensity and normalized distance from the nucleus within the cell mask. High values indicate that Tau intensity is strongly organized by nuclear distance, even if the relationship is non-monotonic or ring-like. This captures radial compartmental structure distinct from simple radial slopes, peak locations, or mean perinuclear ratios. |
| 50 | tau radial distance kurtosis about nucleus | code | Treats Tau intensity within the cell as mass and computes the intensity-weighted kurtosis of pixel distances from the nuclear centroid. High values indicate heavy-tailed radial Tau placement, such as simultaneous perinuclear and far-peripheral aggregates, while lower values indicate a more compact radial distribution. This adds a fourth-moment radial descriptor distinct from radial skewness, half-mass distance, or earth-mover summaries. |
| 51 | tau radial intensity slope from nucleus | code | Computes the robust slope or Spearman correlation between Tau intensity and distance from the nuclear boundary within the cell mask, excluding nuclear pixels. A negative or near-zero slope indicates Tau concentrated near the soma/perinuclear region, whereas a positive slope suggests more distal neuritic enrichment. This provides a continuous spatial localization feature complementary to simple compartment intensity ratios. |
| 52 | tau puncta perinuclear sg clustering score | vlm | Scores whether small, bright Tau puncta form stress-granule-like clusters preferentially around the nucleus within the cytoplasm in the 2D Tau MIP. The feature emphasizes localized perinuclear puncta enrichment rather than uniform puncta burden, providing a Tau-only proxy for cytoplasmic condensate recruitment or impaired granule clearance. Higher values indicate dense perinuclear clustering of discrete Tau puncta with relatively less dispersed cytoplasmic distribution. |

#### Morphology-defined biological process · Global Tau spatial redistribution

**Description.** This feature class describes global reorganization of Tau signal relative to the nucleus, cell boundary, major cellular axis, radial/angular sectors, and geodesic distance. It captures changes in cell polarity, soma-to-periphery or soma-to-process compartmentalization, and the directionality of Tau transport.

35 features · 3 subcellular architectures

##### Subcellular architecture · Nuclear and perinuclear Tau architecture (5 features)

**Approximate scale.** Organized around a nucleus ~5-10 um in diameter; the perinuclear annulus/shell is typically sub-um to several um thick, with circumferential length ~10-30+ um.

| # | Feature name | Method | Description |
|---|---|---|---|
| 53 | cytoplasmic tau angular polarity | code | Measures how directionally polarized the Tau signal is within the cytoplasm of the 2D MIP, using the intensity-weighted circular resultant length of cytoplasmic pixels around the nuclear centroid. A value near 0 indicates Tau is distributed evenly around the nucleus, while higher values indicate lopsided accumulation in one angular sector. This complements centroid displacement by capturing angular concentration rather than only the displacement distance of the intensity center. |
| 54 | tau angular second harmonic strength | code | This feature measures the strength of bipolar or bar-like Tau asymmetry around the nucleus. Within the cell mask, Tau intensity is accumulated as a function of angle around the nuclear centroid, and the normalized amplitude of the second Fourier harmonic is returned. It is designed to capture elongated two-sided Tau organization distinct from simple one-sided polarity. |
| 55 | tau angular sector entropy deficit | code | Divides the cytoplasm into angular sectors around the nuclear centroid and computes one minus the normalized entropy of Tau mass across sectors. Values near zero indicate angularly uniform Tau, while larger values indicate polarized or one-sided Tau accumulation. This is related to angular heterogeneity but distinct from harmonic strength because it captures any angular concentration pattern, not only a second-order symmetry. |
| 56 | tau shape axis mass laterality | code | This feature computes the principal long axis of the segmented cell in the 2D MIP and compares Tau mass on the two opposite sides of that axis-aligned cell coordinate system. The scalar is the absolute normalized difference in Tau integrated intensity between the two longitudinal halves. It measures whether Tau accumulation is biased toward one cellular pole or neuritic side, complementing angular entropy-style features with a cell-shape-referenced asymmetry measure. |
| 57 | cell long axis tau cap enrichment score | vlm | Scores whether Tau signal is concentrated into one or both cap-like zones at the ends of the cell along its long axis. This differs from general polarization because it specifically evaluates terminal polar cap enrichment rather than arbitrary asymmetric hotspots. Such cap-like localization may reflect compartmentalized transport failure, neurite-origin accumulation, or localized aggregate concentration. |

##### Subcellular architecture · Cytoplasmic, somatic, and diffuse intracellular-region scale (8 features)

**Approximate scale.** Approximately 5-30+ um, depending on soma/cytoplasm area; in the 2D MIP this corresponds to large regions of the cell mask excluding the nucleus.

| # | Feature name | Method | Description |
|---|---|---|---|
| 58 | cell periphery tau enrichment | code | Measures Tau enrichment near the cell boundary by taking the ratio of mean Tau intensity in an outer cytoplasmic shell to mean Tau intensity in the remaining inner cytoplasm. This can capture redistribution of Tau toward neuritic or peripheral cellular regions, which is distinct from nucleus-centered radial summaries. The feature is computed on the 2D MIP using the cell and nucleus masks. |
| 59 | cytoplasm tau interior periphery variance ratio | code | Computes the ratio of robust Tau intensity dispersion in the cell interior to that in the peripheral cytoplasmic shell, using distance-to-cell-boundary bins. The interior region can be defined by higher distance-transform values, while the periphery is defined by the outer cytoplasmic band. This feature may distinguish soma-centered or perinuclear Tau heterogeneity from edge- or neurite-associated Tau heterogeneity. |
| 60 | cytoplasmic tau opening spectrum centroid | code | This feature applies a series of morphological openings with increasing disk radii to the cytoplasmic Tau image and measures the characteristic scale at which bright Tau signal is removed. The scalar is the intensity-loss-weighted centroid of the opening-radius spectrum. It provides a robust estimate of dominant Tau granule or aggregate scale without relying on a single hard object threshold. |
| 61 | soma dominant tau visual score | vlm | Scores how strongly the Tau signal in the 2D MIP appears concentrated in the soma-like cell body region rather than being confined to thin processes. The feature should use the cell and nucleus masks to judge cytoplasmic somatic Tau enrichment while excluding obvious nuclear interior signal when possible. A higher scalar score indicates visually stronger somatic Tau accumulation, which is relevant to Tau redistribution in patient neuronal cells. |
| 62 | soma edge tau leakage score | vlm | Estimates whether Tau signal appears to extend diffusely beyond the main cell boundary or blur across the soma edge in the 2D MIP. High scores indicate poorly confined Tau fluorescence around the cell perimeter, suggesting extracellular leakage, remnant material, or loss of compact cellular organization. This differs from extracellular aggregate scoring by focusing on boundary leakage and edge ambiguity rather than distinct outside-cell aggregates. |
| 63 | somatic tau dominance score | vlm | A continuous visual score for how strongly Tau fluorescence dominates the soma/cell-body cytoplasm in the 2D Tau MIP, relative to neurite-like extensions and the rest of the cell. The nucleus should be visually excluded when possible, using the nucleus mask, so the score reflects cytoplasmic Tau accumulation rather than nuclear area. This captures Tau mislocalization into the soma, a key pathological phenotype. |
| 64 | tau compartment mislocalization visual | vlm | Visual scalar score estimating how strongly Tau fluorescence has shifted from a predominantly neuritic or axon-like distribution into the soma and dendrite-like compartments in the 2D Tau MIP. Higher values indicate broad non-axonal Tau accumulation, especially bright soma and proximal dendrite-like signal relative to long thin processes. This is directly relevant because pathological Tau is often characterized by loss of normal axonal enrichment and increased somatodendritic Tau. |
| 65 | tau structure compartmental segregation score | vlm | Scores whether different Tau structural classes, such as droplets, filaments, and bundles, occupy visually distinct subcellular zones rather than being intermixed. A high value indicates clear compartmental separation, for example droplets concentrated near the soma while filaments or bundles dominate the periphery. This captures mesoscale organization of Tau morphology across the 2D single-cell MIP. |

#### Subcellular architecture · Whole-cell geometry, global polarity, and spatial-distribution scale (22 features)

**Approximate scale.** Approximately 10-100+ um; larger if the full neurite arbor is included in the 2D projection.

| # | Feature name | Method | Description |
|---|---|---|---|
| 66 | cell tau distal geodesic mass fraction | code | Computes the fraction of total cellular Tau mass located in the most geodesically distal region of the cell, defined by the highest 20% of geodesic distances from the nuclear boundary within the cell mask. This captures whether Tau accumulates in remote neurite-like or peripheral regions rather than near the soma. It complements geodesic dispersion by focusing specifically on distal tail burden. |
| 67 | cell tau geodesic distance dispersion | code | This feature measures the intensity-weighted dispersion of Tau signal over geodesic distance from the nucleus through the cell mask. Unlike Euclidean radial measures, geodesic distance respects the actual 2D cell shape and neurite-like extensions in the MIP. A high value indicates Tau mass spread across long cellular paths rather than concentrated near the soma or nucleus. |
| 68 | cell tau medial axis affinity | code | Compute the medial-axis skeleton of the cell mask and measure the Tau-weighted affinity of pixels to that skeleton, for example using an exponential decay with distance from the medial axis. High values indicate Tau concentrated along neurite-like centerlines or filamentous cellular structures, while low values indicate diffuse somatic or peripheral Tau. This captures organization relative to cell geometry rather than only nucleus or boundary distance. |
| 69 | cell tau radial entropy | code | Quantifies how evenly Tau intensity is distributed across radial distance bins from the nucleus centroid to the cell boundary. Tau intensity is summed in normalized radial shells within the cell mask, and Shannon entropy of the radial distribution is returned. Low entropy indicates concentrated perinuclear or peripheral Tau, whereas high entropy indicates broad diffuse redistribution throughout the cell. |

| # | Feature name | Method | Description |
|---|---|---|---|
| 70 | cytoplasm tau isolevel centroid tortuosity | code | Within the cytoplasmic mask, threshold Tau intensity at a sequence of quantiles and compute the intensity-weighted centroid of each superlevel region. The feature is the total path length of these centroids divided by their endpoint displacement, with stable handling when the endpoint displacement is small. It captures whether progressively brighter Tau structures shift smoothly or wander through multiple spatial foci, complementing simple centroid drift measures. |
| 71 | cytoplasm tau radial decile heterogeneity | code | This feature partitions the cytoplasm in the 2D Tau MIP into ten normalized radial bands from the nuclear boundary toward the cell boundary, then computes the dispersion of band-wise Tau intensity means relative to within-band variation. It captures whether Tau is organized into uneven perinuclear, medial, or peripheral zones rather than being diffusely distributed. Such radial heterogeneity may reflect Tau redistribution, perinuclear accumulation, or cytoplasmic exclusion patterns linked to neuronal Tau pathology. |
| 72 | cytoplasm tau radial mass iqr width | code | Measures the interquartile width of Tau signal along the normalized cytoplasmic radial coordinate from the nuclear boundary to the cell boundary in the 2D MIP. Tau intensities within the cytoplasm are used as weights, and the feature is the weighted 75th percentile minus weighted 25th percentile of this normalized distance. A narrow value indicates Tau concentrated in a restricted radial zone, while a broad value indicates diffuse cytoplasmic spreading, complementing but not duplicating simple radial slope features. |
| 73 | cytoplasm tau radial quantile bow | code | Quantifies nonlinearity in the Tau radial mass distribution using weighted radial quantiles in the cytoplasm. Compute normalized nuclear-to-cell-boundary distance for each cytoplasmic pixel, then calculate (q90 + q10 - 2*q50) / (q90 - q10 + epsilon), where q10, q50, and q90 are Tau-intensity-weighted distance quantiles. This captures whether Tau mass is concentrated toward both extremes, centrally shifted, or peripherally shifted in a way that is distinct from a monotonic radial gradient. |
| 74 | cytoplasmic tau radial enrichment bandwidth | code | Quantifies the radial width of the main Tau-enriched band in the cytoplasm of the 2D Tau MIP. After excluding the nucleus, the Tau radial profile from nuclear boundary toward the cell boundary is computed and the width of the dominant enrichment zone above half of its local peak is normalized by cytoplasmic radial extent. This captures whether Tau accumulation is tightly shell-like, broadly diffuse, or spread across much of the cytoplasm. |
| 75 | cytoplasmic tau radial peak prominence | code | Within the 2D Tau MIP, this feature uses the cell mask minus the nucleus mask to build a normalized radial profile from the nuclear boundary toward the cell boundary. It measures how strongly the most Tau-enriched cytoplasmic annulus stands out from the typical annular Tau density, for example as a robust peak-to-background prominence. This captures shell-like or band-like cytoplasmic Tau redistribution that is distinct from total radial mass displacement or simple radial entropy. |
| 76 | cytoplasmic tau radial transition total variation | code | Measures how abruptly Tau intensity changes across concentric cytoplasmic shells between the nucleus boundary and the cell boundary in the 2D Tau MIP. The feature is the normalized total variation of the shell-wise median Tau profile, using the cell mask minus nucleus mask as cytoplasm. It may capture sharp radial redistribution or annular accumulation patterns that differ from smoother diffuse Tau spread. |
| 77 | high tau centroid delaunay length cv | code | Segments high-Tau connected components inside the cell using a robust local or quantile threshold, extracts their centroids, constructs a Delaunay graph, and returns the coefficient of variation of graph edge lengths. High values indicate irregular spacing with mixed clustered and isolated aggregates, while low values indicate more uniform aggregate spacing. This provides a spatial-graph view of aggregate organization distinct from nearest-neighbor crowding or Voronoi inequality. |
| 78 | radial tau distribution entropy | code | Quantifies how evenly Tau signal is distributed across concentric distance zones from the nucleus within the cell mask. The total Tau intensity in each radial bin is converted to a probability distribution, and Shannon entropy is computed. Low entropy suggests Tau is concentrated in a restricted compartment such as perinuclear soma or distal neurites, while high entropy suggests broad redistribution across the cell. |
| 79 | tau geodesic profile inflection count | code | This feature constructs a smoothed Tau intensity profile along geodesic distance from the nucleus through the cell mask and counts significant inflection points in that profile. Multiple inflections indicate alternating Tau-rich and Tau-poor zones, such as perinuclear rings, medial aggregates, or peripheral enrichment, while few inflections indicate a simpler monotonic distribution. It complements radial mass and distance features by focusing on profile shape complexity rather than total displacement or skew. |
| 80 | tau intensity centroid centrality | code | Measures how centrally the Tau intensity-weighted centroid lies relative to the nucleus and cell geometry. It is calculated as one minus the normalized distance between the Tau intensity centroid and the nucleus centroid, using the effective radius of the cell mask for normalization. Higher values indicate Tau concentrated near the soma/nucleus, whereas lower values indicate Tau shifted toward distal processes or peripheral cell regions. |
| 81 | tau mass covariance elongation | code | Computes the anisotropy of the Tau intensity-weighted spatial covariance matrix within the cytoplasmic cell mask on the 2D MIP image. The feature is defined as the normalized difference between the major and minor eigenvalues, so high values indicate Tau mass concentrated along an elongated axis rather than isotropically filling the cell. This may capture loss of normal compartmental organization or emergence of polarized aggregate-rich structures without duplicating simple angular polarity metrics. |
| 82 | tau mass principal axis kurtosis | code | Measures the fourth standardized moment of Tau intensity projected onto the principal mass axis within the cell mask. In the 2D MIP, each cytoplasmic Tau-positive pixel is projected onto the intensity-weighted major axis, and the weighted kurtosis of this one-dimensional mass distribution is reported. High values indicate Tau mass concentrated in central or terminal extremes along a dominant axis, while lower values reflect more evenly distributed Tau. |

| # | Feature name | Method | Description |
|---|---|---|---|
| 83 | tau radial chord continuity median | code | Measures whether high Tau signal forms long continuous runs along radial paths from the nucleus toward the cell edge. For many rays starting at the nuclear boundary and ending at the cell boundary, threshold Tau locally and compute the longest contiguous high-Tau run as a fraction of that ray length, then take the median across rays. This complements transition-count features by emphasizing uninterrupted radial continuity rather than the number of on/off changes. |
| 84 | tau radial mass earthmover distance | code | Measures the 1D Wasserstein distance between the observed radial Tau mass distribution and an area-normalized uniform radial distribution from the nuclear boundary toward the cell boundary. The feature is computed on the 2D MIP using nucleus and cell masks, with higher values indicating strong radial redistribution of Tau toward perinuclear or peripheral zones. It provides a complementary view to radial skewness by quantifying whole-distribution displacement rather than a single moment. |
| 85 | tau radial shell mass skewness | code | This feature quantifies the skewness of Tau mass over normalized radial shells from the nuclear boundary toward the cell boundary in the 2D MIP. Tau intensity is summed in cytoplasmic distance shells and the skewness of the resulting radial mass distribution is reported. It complements half-mass and radial entropy features by capturing whether Tau burden is biased toward perinuclear, mid-cytoplasmic, or peripheral regions. |
| 86 | cytoplasmic tau shear flow alignment score | vlm | Scores the extent to which Tau filaments, streaks, and elongated granular textures appear arranged in a common shear-flow-like direction across the cytoplasm of the 2D MIP. A high value indicates coherent directional streaming or laminar organization of Tau signal, whereas a low value indicates isotropic haze, random puncta, or tangled disorganization. This may reflect cytoskeletal remodeling and pathological reorientation of Tau-bearing structures. |
| 87 | tau mass centroid offcenter score | vlm | A scalar score for how strongly the visually dominant Tau signal mass is displaced toward one side of the cell body or whole cell. It should be low when Tau is evenly distributed around the soma and neurites, and high when a large inclusion, cap, or polarized aggregate pulls the Tau distribution off-center. This complements polarity and nuclear-displacement features by focusing on the apparent center of Tau burden rather than a specific compartment boundary. |

#### Morphology-defined biological process · Abnormal neuronal process

**Description.** This feature class focuses on Tau continuity, obstruction, swelling, branch-point enrichment, terminal retention, and shaft organization within neurites, axons, dendrites, branches, terminals, and shafts. It reflects the integrity of neuronal-process architecture and intracellular transport.

20 features · 2 subcellular architectures

##### Subcellular architecture · Cytoplasmic, somatic, and diffuse intracellular-region scale (2 features)

**Approximate scale.** Approximately 5-30+ um, depending on soma/cytoplasm area; in the 2D MIP this corresponds to large regions of the cell mask excluding the nucleus.

| # | Feature name | Method | Description |
|---|---|---|---|
| 88 | soma neurite texture transition abruptness score | vlm | A scalar visual score measuring how abruptly Tau texture changes between the soma and emerging neurites. Low values correspond to a smooth, continuous Tau appearance across compartments, while high values indicate coarse/mottled somatic Tau with sharply different smooth, dim, fragmented, or thread-like neuritic signal. This may capture compartment-specific Tau remodeling beyond simple soma-versus-neurite intensity differences. |
| 89 | soma process tau blending score | vlm | Scores the degree to which Tau signal appears visually continuous from the soma into neurite-like processes rather than being clearly compartmentalized. A high score indicates blurred or seamless Tau-positive spread across the soma-process junctions, suggesting loss of normal compartment restriction. This is relevant to Tau mislocalization and complements features focused only on soma dominance or distal neurite continuity. |

##### Subcellular architecture · Neurites, dendrite-/axon-like processes, and branching arbors (18 features)

**Approximate scale.** Diameter approximately 0.1 to several um; length can reach tens to hundreds of um or more, often extending beyond the soma in 2D projection.

| # | Feature name | Method | Description |
|---|---|---|---|
| 90 | axonal polarity loss visual | vlm | Visual score for loss of Tau polarity, comparing whether Tau appears concentrated in one or a few long thin axon-like processes versus diffusely distributed across the entire cell and shorter branches. Higher values represent weaker axonal enrichment and more uniform whole-cell Tau signal. This complements total intensity by focusing on the spatial organization of Tau rather than its absolute brightness. |
| 91 | branch like tau invasion score | vlm | A continuous score estimating the extent to which Tau signal visibly invades multiple branch-like neurite or dendrite-like processes emerging from the soma. High values indicate Tau-positive branching arbors, thick proximal processes, or widespread non-somatic extensions rather than Tau confined to a single thin track. This reflects pathological redistribution of Tau into dendrite-like compartments and loss of axon-restricted localization. |
| 92 | cell process tau taper preservation score | vlm | Scores how well Tau-positive cell processes preserve a smooth proximal-to-distal taper in width and brightness. High values indicate continuous processes that gradually narrow and fade, while low values indicate abrupt swellings, breaks, or irregular Tau loading along processes. This complements arborization features by emphasizing local taper integrity rather than branch number or span. |

| # | Feature name | Method | Description |
|---|---|---|---|
| 93 | dendritic branch order tau persistence score | vlm | Visual scalar score for how strongly Tau signal persists into higher-order, finer neurite branches rather than fading after primary proximal branches in the 2D Tau MIP. High values indicate broad dendritic/neuritic spread of Tau into distal arbor tips, consistent with loss of normal axonal restriction and pathological redistribution. |
| 94 | dendritic branchpoint tau nodal score | vlm | A scalar visual score estimating whether Tau signal is preferentially concentrated at dendrite-like branch points or neurite junctions. It should be high when branch nodes show bright punctate or swollen Tau accumulations relative to adjacent shaft segments. This feature may reflect abnormal Tau aggregation or transport stalling at neuritic junctions, which is distinct from general dendritic thread burden. |
| 95 | dendritic tau branch order gradient reversal score | vlm | Scores whether Tau signal or aggregate burden appears unexpectedly stronger in distal or higher-order dendritic branches than in proximal dendritic segments near the soma. The feature uses the 2D MIP to visually compare branch-order-dependent Tau distribution, rather than simply asking whether dendrites remain Tau-positive. A high score may indicate abnormal propagation or distal seeding of pathological Tau in dendritic arbors. |
| 96 | dendritic thread inclusion score | vlm | Visual scalar score for thread-like or filamentous Tau inclusions running along dendrite-like branches rather than being restricted to a single axon-like process. Higher values indicate clear Tau-positive linear threads, thickened dendritic shafts, or elongated inclusions in multiple branch-like processes. This feature targets dendritic Tau pathology and complements somatic aggregate features. |
| 97 | dendritic thread knot entanglement score | vlm | Evaluates whether dendritic thread-like Tau inclusions appear knotted, tangled, or mutually intertwined within dendritic processes. The feature emphasizes local thread entanglement and crossing-like complexity inside dendrite-associated Tau structures, not merely the amount of dendritic Tau. Higher values suggest more advanced thread-like inclusion organization in dendritic compartments. |
| 98 | distal neurite tau continuity score | vlm | Estimates how continuously Tau signal persists along thin distal cellular processes visible in the 2D MIP cell mask. Unlike filament fragmentation or orientation coherence, this feature focuses on long-range maintenance versus fading of Tau signal toward distal neurite regions, which may relate to axonal Tau retention or transport disruption. |
| 99 | distal neurite terminal bulb tau score | vlm | A scalar score for bright bulb-like Tau accumulations at distal ends of neurite-like processes. The feature should capture terminal swelling, rounded end-clubs, or growth-cone-like Tau aggregates rather than puncta distributed along the shaft. It is relevant because Tau pathology can begin or concentrate in distal axonal or terminal regions before becoming strongly somatic. |
| 100 | neurite aggregate comet tail orientation score | vlm | Scores whether bright Tau aggregates along neurites show asymmetric diffuse tails or comet-like trails aligned with the neurite axis. The feature captures directional aggregate morphology in the 2D Tau MIP, potentially reflecting transported, spreading, or recently deposited Tau material. Higher values indicate more frequent and more consistently oriented aggregate-tail structures along neurites. |
| 101 | neurite bead size distance trend score | vlm | Measures whether bead-like Tau accumulations along neurites show a systematic size trend with distance from the soma, such as progressively larger distal beads or proximal bead swelling. This differs from general neurite beading or puncta spacing because it captures directional size progression along the neurite axis. It may reflect transport failure, local aggregation kinetics, or distal Tau accumulation. |
| 102 | neurite beading fragmentation score | vlm | Visual score quantifying how beaded, discontinuous, or fragmented the Tau signal appears along neurite-like processes in the 2D MIP. Higher values indicate punctate beads, broken tracks, alternating bright and dim segments, or irregular granular neuritic Tau rather than smooth continuous labeling. This may capture early pathological disruption of axonal or dendritic Tau organization. |
| 103 | neurite shaft tau swelling score | vlm | A scalar score estimating whether Tau-positive neurite shafts appear abnormally thickened, swollen, or locally expanded along their length. It should emphasize broad shaft enlargement and uneven caliber rather than isolated bead-like puncta or terminal bulbs. Such swelling may indicate neuritic degeneration, Tau accumulation, or disrupted cytoskeletal organization complementary to fragmentation scores. |
| 104 | neurite tau phase slip banding score | vlm | Assesses whether Tau signal along neurite-like filaments shows alternating bright and dim bands that appear locally shifted or out of phase across adjacent parallel segments. A high score reflects discontinuous banding with misaligned periodicity, rather than simple beading or fragmentation. This captures subtle nanoscale-to-mesoscale organization of Tau along neuronal processes in the 2D MIP. |
| 105 | neurite tau taper mismatch score | vlm | Visual score for mismatch between neurite geometry and Tau signal width or intensity along the shaft. High values indicate Tau remaining broad, swollen, or abruptly thick in visually thin/tapering neurites, which may reflect abnormal neuritic Tau accumulation rather than normal smooth axonal labeling. |
| 106 | proximal neurite tau choke point score | vlm | Scores whether Tau intensity is visually concentrated at one or more proximal neurite emergence sites near the soma, forming a bottleneck-like bright zone in the 2D MIP. This complements radial and edge-leakage features by emphasizing Tau accumulation at process origins, a possible marker of altered axonal or dendritic transport. |
| 107 | terminal sheet like tau spread score | vlm | Visual score for Tau signal spreading into flattened, sheet-like, lamellipodial, or fan-plate terminal regions rather than ending as thin fibers or compact bulbs. This captures a distinct distal terminal morphology in the 2D Tau projection that may indicate abnormal terminal Tau redistribution. |

#### Morphology-defined biological process · Liquid-liquid phase separation

**Description.** This feature class captures puncta, droplets, condensate-like signal, granules, microgranules, beading, spot-to-haze transitions, and stress-granule-like Tau puncta. It reflects processes related to liquid–liquid phase separation, RNA/RBP granule interactions, and early nanoscale-to-microscale reorganization of Tau-positive signal, without assuming the presence of bona fide Tau aggregates.

49 features · 5 subcellular architectures

#### Subcellular architecture · Local texture and pixel-neighborhood scale (3 features)

**Approximate scale.** Approximately 0.2-1 um imaging texture scale; the lower bound is constrained by conventional fluorescence microscopy lateral resolution (~200 nm).

| # | Feature name | Method | Description |
|---|---|---|---|
| 108 | cytoplasm tau spot to haze energy ratio | code | This feature separates punctate Tau signal from diffuse cytoplasmic haze by comparing high-frequency spot-enhanced energy, such as Laplacian-of-Gaussian or white-top-hat response, to low-frequency background-smoothed Tau energy. A high value indicates punctate or granular Tau aggregation, while a low value indicates diffuse somatic/cytoplasmic Tau accumulation. This directly targets the biologically important diffuse-versus-aggregate transition in Tau pathology. |
| 109 | cytoplasmic granular haze score | vlm | Scores the extent of fine, diffuse, grainy Tau texture filling the cytoplasm of the cell in the 2D MIP, excluding obvious large droplets, filaments, or bundles. High values indicate a hazy granular cytoplasmic Tau background suggestive of diffuse early aggregation or stress-granule-like punctation. This complements features focused on discrete puncta or filaments by emphasizing unresolved fine-scale cytoplasmic texture. |
| 110 | diffuse to punctate soma conversion score | vlm | Measures the apparent transition of somatic Tau from smooth diffuse cytoplasmic fluorescence to a granular or punctate aggregate-dominated pattern. In the 2D Tau MIP, the VLM should compare the relative visual dominance of haze-like soma filling versus discrete bright granules within the segmented soma. Higher values indicate a stronger punctate conversion phenotype, which may reflect pathological Tau condensation or aggregation. |

#### Subcellular architecture · Small puncta, granules, foci, or condensate-like image objects (11 features)

**Approximate scale.** Approximately 0.1-2 um; comparable in order of magnitude to stress granules / cytoplasmic RNP granules and small vesicles/lysosomes.

| # | Feature name | Method | Description |
|---|---|---|---|
| 111 | cytoplasmic tau blob scale geometric mean | code | Estimates the characteristic size scale of discrete Tau-positive blobs in the cytoplasm. A multiscale Laplacian-of-Gaussian detector is applied within the cytoplasmic mask, and the intensity-weighted geometric mean of detected blob radii is normalized by the square root of cell area. This complements granularity-ratio features by measuring the dominant physical scale of punctate Tau structures rather than only the relative energy at small versus large scales. |
| 112 | cytosolic tau log blob scale cv | code | Measures the coefficient of variation of Tau-positive blob scales detected by multi-scale Laplacian-of-Gaussian filtering in the cytoplasm of the 2D Tau MIP. It captures whether cytoplasmic Tau puncta are uniform, diffraction-limited spots or a heterogeneous mixture of small puncta and larger condensate-like aggregates, which is relevant to Tau stress-granule recruitment and aggregation states. The feature uses the cell mask to restrict analysis and the nucleus mask to exclude nuclear pixels; if the nucleus mask is unavailable, an eroded central cell region can be conservatively excluded. |
| 113 | droplet boundary tau gradient enrichment | code | For samples with a provided droplet mask, this feature measures the Tau gradient magnitude along droplet boundaries and normalizes it to the typical Tau gradient magnitude in nearby cytoplasm. High values indicate sharp Tau recruitment or exclusion interfaces around droplets. This is related to droplet-associated Tau organization but focuses on boundary sharpness rather than centroid offset or mean recruitment contrast. |
| 114 | droplet tau asymmetric cap fraction | code | For each segmented droplet, this feature measures whether nearby Tau signal is distributed uniformly around the droplet or concentrated into a localized cap. Tau intensity is measured in an annulus around each droplet and the fraction contained in the brightest angular quadrant is computed, then averaged across droplets. It complements droplet proximity features by capturing polarized Tau recruitment around droplet surfaces. |
| 115 | droplet tau centroid offset index | code | For each segmented droplet, computes the distance between the droplet geometric centroid and the Tau intensity-weighted centroid inside the droplet, normalized by droplet equivalent radius. The final scalar is the Tau-mass-weighted mean across droplets. This captures asymmetric Tau recruitment within droplets using a centroid-shift formulation distinct from cap-fraction measurements. |
| 116 | droplet tau fraction of cell signal | code | Measures the fraction of total cell Tau intensity that lies inside mask_droplet objects, treating binary masks as connected components or label masks as provided. If the droplet mask is absent or empty, the value can be defined as zero to represent no segmented droplet burden. This captures Tau concentrated in punctate droplet-like structures that may reflect stress-granule-associated or aggregate-prone Tau states. |
| 117 | droplet tau hotspot proximity index | code | Measures whether segmented droplets lie unusually close to Tau hotspots by comparing the mean distance from droplet centroids to nearest high-Tau local maxima against a cell-size-normalized distance scale. Higher values indicate stronger spatial coupling between droplets and bright Tau accumulations. This is related to Tau recruitment but focuses on centroid-to-hotspot geometry rather than intensity contrast inside the droplet mask. |

| # | Feature name | Method | Description |
|---|---|---|---|
| 118 | droplet tau recruitment contrast | code | Uses the provided droplet mask to compare mean Tau intensity inside droplets with mean Tau intensity in the surrounding local cytoplasmic neighborhood. The feature can be summarized as the median droplet-to-local-background contrast across droplets, with zero assigned when no droplet structure is present. It is designed to capture Tau recruitment into condensate-like puncta or stress-granule-like droplets, rather than droplet size or spatial clustering. |
| 119 | tau log punctateness index | code | Quantifies small bright Tau puncta and speckled aggregation by applying a Laplacian-of-Gaussian or difference-of-Gaussian filter to the 2D Tau MIP within the cell mask. The feature can be defined as the positive LoG response energy normalized by total cell Tau intensity or cytoplasmic area. Higher values indicate granular or punctate Tau morphology that may correspond to oligomeric or stress-granule-like Tau-positive structures. |
| 120 | punctate tau condensate burden score | vlm | Estimates the visual burden of discrete bright Tau-positive puncta, droplets, or small condensates within the segmented cell in the 2D MIP. The scalar score should reflect both the apparent number and prominence of punctate objects while avoiding isolated noise speckles. This captures granular Tau organization that may relate to aggregation or stress-granule-like recruitment patterns. |
| 121 | somatic tau microgranule density score | vlm | Scores the abundance of very fine, dot-like Tau granules within the somatic cytoplasm outside the nucleus. A high value indicates a dense peppered texture of small Tau-positive grains, distinct from broad diffuse haze or large compact inclusions. This may reflect early condensate-like or stress-granule-associated Tau redistribution. |

#### Subcellular architecture · Filaments, skeletons, ridges, or short linear structures (9 features)

**Approximate scale.** Biological fiber diameters are often nanometer-scale (actin ~7 nm, intermediate filaments ~8-11 nm, microtubules ~25 nm); in optical images they appear as linear/skeletonized structures >=~0.2 um wide, with lengths of several um or more.

| # | Feature name | Method | Description |
|---|---|---|---|
| 122 | cytoplasmic tau hessian blob ridge balance | code | Uses multiscale Hessian filtering within the cytoplasm to compare blob-like Tau responses with ridge-like Tau responses. The scalar is a normalized balance score between isotropic punctate structure and elongated filamentous structure, weighted by local Tau intensity. This may capture shifts between granular Tau aggregates and fibrillar or thread-like Tau organization in the 2D super-resolution MIP. |
| 123 | filament tau beading coefficient | code | Measures intensity intermittency along segmented Tau filaments by skeletonizing mask_filament and computing the coefficient of variation of Tau intensity sampled along the filament skeleton after mild smoothing. High values indicate beaded or fragmented Tau signal along filaments, while lower values indicate more continuous filamentous Tau. This is relevant to pathological neuritic beading and fragmented Tau organization in 2D MIP images. |
| 124 | bundle surface puncta decoration score | vlm | Scores the degree to which bundle surfaces are decorated by small bright Tau puncta or bead-like deposits that appear attached to the bundle exterior. The VLM should focus on puncta aligned along bundle edges rather than internal bundle texture or unrelated cytoplasmic dots. If bundle masks are absent, the feature should be scored based on visible bundle-like Tau structures if present, otherwise near zero. |
| 125 | droplet bundle wetting flattening score | vlm | Scores whether Tau droplets adjacent to bundles appear flattened, elongated, or spread along the bundle surface rather than remaining round and isolated. This is distinct from droplet coalescence because it focuses on droplet wetting behavior at bundle interfaces. High values may indicate physical interaction between condensate-like Tau droplets and fibrillar Tau assemblies. |
| 126 | droplet filament anchor multiplicity score | vlm | Scores how often Tau droplets act as multi-valent anchor points where several filaments contact or radiate from the same droplet. A high value indicates droplets that visually behave like hubs with multiple filament attachments, rather than isolated round puncta or simple bead-on-filament structures. This may reflect condensate-mediated nucleation or crosslinking of Tau fibrillar assemblies. |
| 127 | droplet filament endpoint tethering score | vlm | Measures whether Tau droplets preferentially appear attached to or positioned at the visible endpoints of Tau filaments rather than along filament midshafts or in free cytoplasm. The feature uses the 2D Tau MIP together with droplet and filament masks when present. Endpoint tethering may indicate nucleation, fragmentation, or growth sites of Tau aggregates. |
| 128 | droplet filament interface seeding score | vlm | Scores the visual association of Tau droplets with nearby filaments, emphasizing droplets that appear attached to, aligned along, or nucleating filament-like extensions in the 2D MIP. This differs from droplet peripheral enrichment by focusing on droplet-filament interfaces and potential seeding relationships rather than droplet position within the cell. |
| 129 | droplet size gradient along filaments score | vlm | Measures whether droplets arranged along Tau filaments show a directional size trend, such as progressively larger or smaller beads along a strand. A high score indicates visually coherent bead-size gradients along filament paths rather than random droplet sizes. This is distinct from radial sorting or neighborhood assortativity because it evaluates ordering specifically along filament trajectories. |
| 130 | punctate to filamentous tau balance | vlm | A scalar continuum score describing whether visible Tau aggregates appear predominantly as compact droplets/puncta versus elongated filaments or strands. Low values correspond to mostly round puncta or droplets, while high values correspond to filamentous, thread-like, or rod-like Tau morphology. This captures aggregate morphotype differences that may reflect different Tau assembly states. |

#### Subcellular architecture · Composite Tau-positive object morphologies and inter-object relationships (18 features)

**Approximate scale.** Approximately 0.1-20+ um; a multi-scale class spanning small puncta, short filaments, thick bundles and larger objects.

| # | Feature name | Method | Description |
|---|---|---|---|
| 131 | cytosolic tau puncta spacing mad ratio | code | Quantifies irregularity in the spatial spacing of cytoplasmic Tau puncta by detecting local high-Tau maxima and computing the median absolute deviation of nearest-neighbor distances divided by the median nearest-neighbor distance. Pathological Tau often forms uneven clustered puncta or bead-like deposits rather than regularly spaced sparse signal, so this captures a complementary spatial organization phenotype. The computation is performed within the cytoplasm using the cell and nucleus masks, with a conservative cell-only fallback if the nucleus mask is missing. |
| 132 | tau puncta ripley l peak | code | Detects LoG-like Tau puncta inside the cytoplasm and computes the maximum normalized Ripley L(r) - r value across biologically reasonable radii. The feature quantifies whether puncta are spatially clustered rather than randomly scattered, independent of simple puncta count. Clustered cytoplasmic Tau puncta may reflect stress-granule-like recruitment, oligomeric seeding, or local aggregation foci. |
| 133 | cytoplasmic tau puncta lattice order score | vlm | Visual scalar score for whether small cytoplasmic Tau puncta show quasi-regular spacing or lattice-like ordering rather than random clustering. This may capture ordered nanoscale or mesoscale Tau organization visible in super-resolution Tau MIPs, while remaining distinct from total puncta burden. |
| 134 | cytoplasmic tau puncta pair spacing dispersion score | vlm | Scores how irregular the visual spacing is among cytoplasmic Tau-positive puncta within the 2D Tau MIP, focusing on the cell cytoplasm outside the nucleus. A high value indicates puncta are neither lattice-like nor uniformly distributed, but instead show mixed close clustering and large empty gaps. This complements prior puncta uniformity/lattice features by emphasizing pairwise spacing dispersion rather than overall order. |
| 135 | droplet attachment fraction visual score | vlm | Estimates the fraction of Tau droplets that appear physically attached to, touching, or embedded along filaments or bundles rather than floating as isolated cytoplasmic puncta. The scalar increases when most droplet-like objects are associated with fibrillar structures. This may reflect coupling between condensate-like Tau droplets and fibrillar Tau scaffolds. |
| 136 | droplet clusteredness score | vlm | A continuous score describing whether Tau droplets or puncta are spatially dispersed throughout the cell or concentrated into local clusters. Low values indicate isolated, evenly distributed puncta, while high values indicate dense neighborhoods of droplets, puncta swarms, or local aggregate hotspots. This captures spatial organization of compact Tau aggregates beyond simple puncta count. |
| 137 | droplet coalescence necking score | vlm | Scores whether Tau droplets appear partially fused through narrow Tau-positive necks or bridges. A high value indicates adjacent rounded condensates connected by constricted links, suggesting coalescence or liquid-like aggregate maturation. If no droplet-like structures are visible, the score should be low rather than missing. |
| 138 | droplet contact side tau polarity score | vlm | Measures whether Tau droplets show asymmetric brightness concentrated on the side facing a nearby filament or bundle. High values indicate visibly polarized droplet interiors or rims, with stronger Tau signal at the contact-facing side than the free side. This captures directional interaction between droplet-like condensates and fibrillar Tau structures. |
| 139 | droplet interstitial tau web trapping score | vlm | Scores the degree to which droplets appear embedded within a web of Tau-positive strands or bridges in the spaces between droplets on the 2D MIP. A high value indicates that inter-droplet gaps are crossed by thin Tau filaments or reticular strands, suggesting droplet-associated trapping within a Tau network. This is distinct from measuring droplet rim brightness or direct filament anchoring because it emphasizes the intervening mesh-like Tau architecture. |
| 140 | droplet peripheral enrichment score | vlm | Measures whether Tau-positive droplets are preferentially located near the cell periphery rather than near the nucleus or central soma in the 2D Tau MIP. High scores indicate a peripheral rim or edge-biased distribution of droplets, while low scores indicate central or uniformly distributed droplets. This captures a spatial organization of punctate Tau aggregates not addressed by simple droplet clusteredness. |
| 141 | droplet population dominance score | vlm | Scores whether the droplet mask population is dominated by one or a few large Tau-associated droplets versus many similarly sized small droplets. If droplet segmentation is absent or no droplets are visible, the score should be near zero rather than missing. This captures condensate size hierarchy, which may reflect coalescence or maturation of Tau-associated droplet-like structures. |
| 142 | droplet satellite puncta collar score | vlm | Scores the presence of small Tau-bright puncta forming a satellite-like collar around segmented droplets in the 2D Tau MIP. The feature emphasizes nearby punctate halos around droplets rather than direct filament-droplet contact. Such satellite puncta may reflect local nucleation, condensate-associated Tau recruitment, or stress-granule-like organization. |
| 143 | droplet shadow overlap field score | vlm | Measures whether Tau-depleted regions surrounding multiple droplets visually merge into larger continuous low-signal fields in the 2D MIP. A high score indicates that individual droplet-associated shadows overlap or coalesce into broad Tau-poor basins, rather than appearing as isolated halos around each droplet. This extends droplet-shadow analysis from single-droplet depletion strength to collective spatial field organization. |
| 144 | droplet size assortative neighborhood score | vlm | Scores whether Tau droplets of similar apparent size tend to be spatially grouped together within the cell, rather than large and small droplets being randomly intermixed. A high value reflects local size assortativity among droplet neighborhoods. This may capture coordinated condensate growth or maturation zones distinct from global droplet clustering or radial size sorting. |

| # | Feature name | Method | Description |
|---|---|---|---|
| 145 | droplet size radial sorting score | vlm | Measures whether Tau droplets show an ordered size pattern as a function of distance from the nucleus, such as larger droplets preferentially near the nucleus or preferentially near the cell periphery. The score should consider the 2D cell and nucleus masks plus droplet morphology in the Tau channel. This feature captures spatial sorting of condensate-like aggregates rather than simply droplet abundance or clustering. |
| 146 | droplet string spacing regularity score | vlm | Measures how regularly Tau droplets are arranged as bead-like sequences along curvilinear paths, filaments, or neurite-like axes in the 2D MIP. A high score indicates repeated, nearly periodic droplet spacing along one or more tracks, while a low score indicates randomly scattered or irregularly spaced droplets. This feature targets spatial periodicity of condensates rather than simply their density or endpoint tethering. |
| 147 | somatic puncta radial bias score | vlm | A scalar score describing whether small somatic Tau puncta are preferentially located near the nuclear/perinuclear region, near the soma edge, or evenly distributed, encoded as the strength of radial bias regardless of direction. Low values indicate uniform puncta distribution, while high values indicate a pronounced radial concentration pattern. This may reveal spatially organized aggregate nucleation that is not captured by total puncta burden alone. |
| 148 | stress granule like tau puncta uniformity score | vlm | Visual score for the presence of many discrete, round, cytoplasmic Tau-positive puncta of relatively uniform size and brightness, resembling stress-granule-like punctate recruitment in the Tau channel. This does not infer TIA-1 colocalization, but captures a Tau-only punctate phenotype that may relate to pathological Tau condensation or stress-associated states. |

##### Subcellular architecture · Cell boundary, cortical, peripheral-shell, and extracellular-neighborhood scale (8 features)

**Approximate scale.** Set against a whole-cell diameter of ~10-100 um; the measured region is typically a thin band or boundary arc inside/outside the cell edge.

| # | Feature name | Method | Description |
|---|---|---|---|
| 149 | tau condensate intensity size allometry | code | Measures the scaling relationship between size and Tau brightness of high-intensity condensate-like components. High-Tau components are detected within the cell, and the slope of log integrated intensity versus log component area is computed using robust regression. Values above or below linear scaling may distinguish dense compact inclusions from diffuse enlarged accumulations. |
| 150 | droplet internal tau mottling score | vlm | Scores the degree to which segmented droplet-like Tau objects show uneven, mottled, or patchy internal fluorescence rather than smooth homogeneous interiors. The feature is assessed in the 2D Tau MIP using droplet masks when available, and should be near zero when droplets are absent. Internal mottling may indicate heterogeneous condensate composition or partial transition toward aggregate-like Tau structure. |
| 151 | droplet rim bead angular periodicity score | vlm | Scores whether bright Tau beads or puncta along droplet rims occur at roughly regular angular intervals around the droplet perimeter. This feature focuses on angular periodicity of rim-associated bead placement rather than simple rim polarization or satellite abundance. Higher values indicate more ordered bead spacing around droplets, potentially reflecting structured condensate-rim organization. |
| 152 | droplet rim sector polarization score | vlm | Measures whether Tau-positive droplets show sectorized or one-sided rim brightening, such as crescent-like caps or polarized rim segments, instead of a uniformly circular rim. This feature is distinct from rim-core inversion because it focuses on angular asymmetry around the droplet boundary. High values may indicate droplet wetting, directional interaction with nearby fibrils, or polarized Tau recruitment. |
| 153 | droplet size rim brightness order score | vlm | Scores whether droplet rim Tau brightness is systematically ordered by droplet size, such as larger droplets having brighter or more complete Tau rims than smaller droplets. The VLM should compare multiple droplets within the same cell using the droplet mask and Tau signal, emphasizing monotonic visual ordering rather than absolute brightness. If droplets are absent, the score should be near zero. |
| 154 | droplet tau rim core inversion score | vlm | Measures whether different Tau droplet-like structures within the same cell show mixed rim-dominant and core-dominant intensity patterns in the 2D MIP. A high value indicates coexistence of droplets with opposite internal intensity organization, suggesting heterogeneous condensate maturation states. This is distinct from a single core-shell contrast measure because it emphasizes within-cell inversion diversity across multiple droplets. |
| 155 | tau condensate core shell contrast score | vlm | Scores whether Tau-positive droplet-like or condensate-like structures show a visually distinct bright rim with dimmer core, or bright core with dimmer surrounding shell, in the single Tau MIP. A high value indicates strong internal radial contrast within condensates rather than uniformly bright puncta. This may reflect phase-separated or maturing aggregate states and is complementary to simple droplet size or clusteredness measures. |
| 156 | tau condensate shell thickness heterogeneity score | vlm | Scores how variable the apparent rim or shell thickness is across Tau condensates in the cell. This differs from core-shell contrast because it emphasizes geometric shell width heterogeneity, including condensates with thick asymmetric shells, thin halos, or mixed shell maturation states. Such variability may reflect heterogeneous condensate aging, compaction, or transition toward insoluble aggregates. |

#### Morphology-defined biological process · Microtubule cytoskeletal remodeling

**Description.** This feature class quantifies the length, orientation, curvature, gaps, intersections, and hierarchical organization of Tau-positive filament-like, bundle-like, ridge-like, fibril-like, thread-like, skeletonized, striated, swirl-like, or vortex-like networks.

67 features · 2 subcellular architectures

#### Subcellular architecture · Filaments, skeletons, ridges, or short linear structures (37 features)

**Approximate scale.** Biological fiber diameters are often nanometer-scale (actin ~7 nm, intermediate filaments ~8-11 nm, microtubules ~25 nm); in optical images they appear as linear/skeletonized structures >=~0.2 um wide, with lengths of several um or more.

| # | Feature name | Method | Description |
|---|---|---|---|
| 157 | bundle tau orientation radial misalignment | code | Quantifies whether Tau-positive bundles are oriented radially with respect to the nucleus or instead lie tangentially/obliquely. For each bundle component in mask_bundle.tif, compute its major-axis orientation and compare it with the vector from the nuclear centroid to the bundle centroid; the feature is the Tau-intensity-weighted mean sine of the orientation mismatch. This captures spatial organization of bundle-like inclusions relative to the soma-nucleus geometry, complementing aggregate burden and compactness metrics. |
| 158 | bundle tau structure tensor anisotropy | code | Computes the average structure-tensor anisotropy of Tau intensity within segmented bundle regions on the 2D MIP. High values indicate directionally organized Tau signal inside bundles, whereas low values indicate isotropic or disordered bundle-associated Tau. This complements bundle shape and orientation features by measuring the internal Tau texture alignment. |
| 159 | cytoplasm tau laplacian zero crossing density | code | Apply a lightly smoothed Laplacian-of-Gaussian filter to the Tau signal inside the cytoplasm and count significant zero-crossing contours per unit cytoplasmic area. High values indicate fine-scale alternating bright and dark structure such as granular, reticular, or mesh-like Tau organization. This complements intensity-tail and wavelet measures by focusing on spatial sign transitions rather than energy magnitude. |
| 160 | cytoplasmic tau structure tensor radial alignment | code | Measures whether local Tau ridges or textured structures in the cytoplasm are aligned radially away from the nucleus or tangentially around it. A structure tensor is computed from the Tau MIP in the cytoplasmic mask, and local dominant orientations are compared with the radial direction from the nucleus centroid; the mean absolute radial alignment score is returned. This may distinguish neurite-like radial Tau structures from annular or tangled somatic organization. |
| 161 | filament curvature tau intensity coupling | code | Measures whether Tau intensity along segmented filaments is associated with filament bending in the 2D MIP. Filament masks are skeletonized, local curvature is estimated along skeleton paths, Tau intensity is sampled at the same skeleton points, and the within-sample rank correlation between curvature and intensity is returned. This can reveal whether bright Tau accumulates preferentially at curved, stressed, or kinked filament regions. |
| 162 | filament radial orientation bias | code | Measures whether Tau filaments preferentially point radially away from the nucleus or tangentially around it. Local filament skeleton tangents are compared with vectors from the nuclear centroid, and the scalar is the mean absolute cosine of this angle across skeleton pixels. This captures spatial organization of filament orientation relative to the cell body, distinct from global orientation coherence or filament length density. |
| 163 | filament skeleton tau continuity | code | Uses the provided filament mask to skeletonize Tau-positive filament structures and returns the longest skeleton component length divided by total filament skeleton length. Higher values indicate a more continuous dominant filament network, whereas lower values indicate fragmented or beaded filament organization. This directly targets Tau filament continuity and fragmentation in the 2D MIP. |
| 164 | filament skeleton tortuosity | code | Measures the average excess tortuosity of Tau filament objects from the provided filament mask by skeletonizing each component and comparing skeleton path length to endpoint chord distance. Higher values indicate curved, winding, or tangled filaments rather than straight linear structures. This complements filament length density by characterizing filament geometry rather than abundance. |
| 165 | filament tau beadiness cv | code | Samples Tau intensity along the skeleton of the provided filament mask and computes the coefficient of variation of intensity along the filament paths. High values indicate alternating bright beads and dim gaps along filamentous Tau structures, a pattern associated with fragmented or beaded Tau pathology. If no filament mask or skeleton is present, the feature can be defined as zero to indicate no measurable filament beadiness. |
| 166 | filament tau loading heterogeneity index | code | Quantifies how unevenly Tau intensity is loaded across segmented filamentous structures by computing Tau integrated intensity per skeleton length for each filament component and returning a robust coefficient of variation across components. Normal axonal Tau may appear as smoother continuous linear signal, whereas pathological filamentous inclusions and fragmented neurites can show highly variable Tau loading. The feature uses mask_filament when available and the cell mask to restrict intracellular filaments; if mask_filament is absent, a ridge-filter-derived filament candidate mask can be used as a fallback. |
| 167 | filament tau longitudinal autocorrelation peak | code | This feature captures periodic beading or repeated intensity modulation along segmented Tau-positive filaments. Tau intensity is sampled along each filament skeleton, an autocorrelation curve is computed, and the strongest nonzero-lag peak is aggregated across filaments. It provides a complementary view to gap-based filament features by measuring regularity of longitudinal Tau spacing. |
| 168 | filament tau skeleton gap fraction | code | Skeletonizes the filament mask and measures the fraction of skeleton pixels whose local Tau intensity falls below a cytoplasm-normalized threshold. Higher values indicate discontinuous or fragmented Tau labeling along filaments, while lower values indicate continuous filament-associated Tau. This captures gaps along filament geometry rather than bead intensity variability. |

| # | Feature name | Method | Description |
|---|---|---|---|
| 169 | filament tau transverse decay slope | code | Measures how rapidly Tau intensity decays away from segmented filament skeletons into surrounding cytoplasm. The feature is computed by comparing mean Tau signal in the filament core with successive distance-transform bands outside the filament and fitting a radial decay slope. Steeper decay suggests sharply confined filament-associated Tau, while flatter decay suggests diffuse halo-like accumulation around filaments. |
| 170 | tau filamentous vesselness coupling | code | Measures the correlation between Tau intensity and multiscale line-enhancement or vesselness response within the cytoplasm. High values indicate that bright Tau preferentially lies on thin curvilinear structures, consistent with thread-like neuritic inclusions or filamentous Tau organization. Low values suggest that Tau is more diffuse, blob-like, or spatially unstructured. |
| 171 | tau radon projection anisotropy | code | Quantifies global directional organization of Tau signal inside the segmented cell. The Tau MIP is masked by mask_cell, Radon projections are computed over multiple angles, and the scalar is the ratio of maximum projection energy to median projection energy across angles. High values indicate strongly aligned filamentous or bundle-like Tau organization, whereas low values indicate isotropic punctate or diffuse signal. |
| 172 | tau ridge coherence in cytoplasm | code | Measures the average local orientation coherence of Tau intensity gradients within the cytoplasm using a structure-tensor or Hessian-based ridge analysis on the 2D MIP. High values suggest organized linear Tau fibers or bundles, while lower values suggest diffuse, isotropic, or punctate signal. This complements skeleton-length features because it can capture sub-mask filamentous organization directly from the Tau image. |
| 173 | tau ridge orientation coherence length | code | Applies ridge-enhancing or structure-tensor filters to cytoplasmic Tau signal and estimates the spatial correlation length of the dominant local ridge orientation field. Longer coherence length indicates extended aligned Tau fibers or threads, while shorter coherence suggests fragmented, tangled, or locally disordered structures. This is designed to capture organization of filament-like Tau signal without relying only on predefined filament masks. |
| 174 | cell margin filament exit angle bias score | vlm | Scores whether Tau filaments reaching the cell boundary tend to exit or approach the margin at strongly radial angles versus running tangentially along the edge. A high value indicates a consistent biased exit geometry rather than mixed or random boundary contacts. This can reflect altered neurite-like Tau organization, boundary-associated retraction, or peripheral fibril alignment. |
| 175 | curved filament inner outer brightness bias score | vlm | Scores whether curved Tau filaments show a consistent brightness imbalance between the inner and outer sides of their bends. The VLM should inspect curved filament segments in the Tau MIP and estimate whether intensity preferentially accumulates along one side of the curve. This complements bend-related features by focusing on cross-sectional brightness polarity across curved filaments rather than brightness at the bend itself. |
| 176 | filament bend brightness coupling score | vlm | Scores whether curved or sharply bent Tau filaments are preferentially brighter than straighter filament segments. High values indicate that bends, arcs, or kinked portions of filament masks coincide with local Tau intensity enhancement. This may capture stress-induced Tau accumulation at mechanically or structurally abnormal filament regions. |
| 177 | filament cell span fraction score | vlm | Scores how much of the segmented cell's spatial extent is traversed by Tau-positive filament masks. High values indicate filaments spanning across much of the cell body or processes, while low values indicate localized or absent filament signal. This feature measures the macroscopic spread of filamentous Tau pathology rather than filament disorder or orientation alone. |
| 178 | filament crossing complexity score | vlm | Scores the degree to which Tau-positive filament structures form crossings, intersections, or tangled overlaps in the 2D MIP. This is distinct from filament orientation coherence because it emphasizes topological complexity and entanglement rather than alignment, potentially reflecting advanced fibrillar network formation. |
| 179 | filament endpoint fraying score | vlm | Scores whether Tau-positive filament structures show frayed, split, or brush-like ends in the 2D MIP using the filament mask when available. High values indicate filaments whose termini branch into multiple fine strands or dissolve into ragged punctate fragments. This may capture a distinct filament remodeling phenotype not summarized by crossing complexity or orientation coherence. |
| 180 | filament intersection brightness node score | vlm | Measures whether filament crossings, branchpoints, or junctions appear visually brighter and more aggregate-like than the surrounding filament shafts in the Tau channel. A high score indicates Tau accumulation at network nodes rather than uniform intensity along linear structures. This may reflect nucleation or stabilization points in pathological Tau assemblies. |
| 181 | filament length scale bimodality score | vlm | Scores whether the Tau filament population visually contains two distinct length classes, such as many short dash-like fragments coexisting with long continuous filaments. The feature uses the 2D MIP and filament mask when available to assess length-distribution structure at the image level. This may reveal mixed aggregation states or transitions between fragmented and mature filamentous Tau. |
| 182 | filament loop area polydispersity score | vlm | Scores how heterogeneous the apparent hole or loop sizes are within Tau filament networks in the 2D MIP. A high value indicates a mixture of tiny mesh openings and large lacuna-like loops rather than a uniformly meshed network. This may capture distinct stages of Tau network remodeling that are not reduced to simple loop abundance. |

| # | Feature name | Method | Description |
|---|---|---|---|
| 183 | filament midshaft intensity pulsing score | vlm | Scores the degree to which Tau-positive filaments show alternating bright and dim segments along their central shafts, excluding obvious terminal fraying. A high score indicates strong periodic or irregular intensity pulsing along filaments while the filament remains structurally continuous. This captures a distinct form of filament maturation or instability in the 2D Tau projection. |
| 184 | filament network cycle density score | vlm | Scores how strongly Tau-positive filament masks form closed loops, cages, or reticulated cycles within the 2D MIP cell area, rather than only open-ended linear strands. A high value reflects many visually apparent loop-like filament circuits around cytoplasmic or nuclear regions. This may capture advanced Tau network organization distinct from simple radial alignment or orientation coherence. |
| 185 | filament orientation coherence score | vlm | Estimates the degree to which Tau-positive filaments in the 2D MIP share a common orientation within the cell. High scores indicate organized, parallel or bundled filament alignment, while low scores indicate randomly oriented, tangled, or disordered Tau filaments. This may reflect differences between structured fibrillar assemblies and chaotic pathological tangles. |
| 186 | filament radial spoke alignment score | vlm | Scores how strongly Tau-positive filaments in the 2D MIP appear as radial spokes extending from the nuclear or perinuclear region toward the cell periphery. This is complementary to tangential nuclear wrapping because it emphasizes outward-directed filament organization rather than circumferential encirclement. A high value may reflect cytoskeletal reorganization or directed aggregate spread within the neuronal cell. |
| 187 | filament short gap reconnection score | vlm | Measures the visual tendency of broken Tau filament segments to appear reconnectable across short dark gaps, as if the network is fragmented but still aligned. A high score indicates many near-collinear segment interruptions rather than fully separated random fragments. This complements general filament fragmentation by focusing on spatially aligned discontinuities that may reflect partial preservation of neuritic or fibrillar architecture. |
| 188 | filament tip brightness taper score | vlm | Scores whether Tau filaments show systematic brightness tapering toward their tips or endpoints in the 2D maximum intensity projection. A high score indicates filaments with bright central shafts that fade smoothly at tips, or conversely bright terminal caps relative to shafts, depending on the dominant visible taper pattern. This captures endpoint intensity organization rather than endpoint fraying or simple filament count. |
| 189 | somatic fibril counterflow domain score | vlm | Scores whether somatic fibrillar Tau contains adjacent domains whose apparent fibril flow, curvature, or swirl direction opposes each other, creating a visual counterflow seam. This differs from measuring a single spiral pitch or global swirl because it emphasizes competing local orientation fields within the same soma. Higher values indicate stronger multi-domain fibrillar organization, which may reflect complex tangle-like remodeling. |
| 190 | somatic fibril spiral pitch variability score | vlm | Measures how variable the spacing and curvature of spiral or swirl-like somatic Tau fibrils appear within the soma. Unlike vortex-center offset, this feature focuses on the internal pitch irregularity of the swirl pattern rather than where the swirl center lies. High values may reflect disorganized tangle compaction and heterogeneous fibril packing. |
| 191 | somatic fibril swirl directionality score | vlm | Visual score for whether somatic Tau is organized into curved, vortex-like, or circumferential fibrillar streaks rather than isotropic dots or diffuse haze. This captures a somatic tangle-like organization pattern in the 2D Tau MIP that may reflect advanced intracellular aggregate remodeling. |
| 192 | somatic fibril vortex center offset score | vlm | Measures whether curving or swirling somatic Tau fibrils appear organized around a vortex-like center that is displaced from the soma or nuclear center in the 2D MIP. A high score indicates an off-center swirl focus, suggesting asymmetric intracellular organization or aggregate-driven remodeling. This is distinct from general swirl directionality because it emphasizes the spatial offset of the apparent rotational center. |
| 193 | somatic tau lamellar onion skin score | vlm | Scores the presence of concentric or quasi-concentric lamellar Tau bands in the soma, resembling onion-skin layering around the nucleus or a central inclusion. This is distinct from a simple perinuclear ring because it captures multiple nested intensity layers or curved band shells in the 2D MIP. Higher values indicate more visually organized lamellar Tau stratification, potentially marking advanced inclusion remodeling. |

#### Subcellular architecture · Thick bundles, fascicles, and intra-bundle structures (30 features)

**Approximate scale.** Approximately 0.2-5+ um in width and 1-20+ um in length; the optical scale of merged linear Tau-positive regions or multi-filament bundles.

| # | Feature name | Method | Description |
|---|---|---|---|
| 194 | bundle compact inclusion burden | code | Aggregates region-level properties of mask_bundle objects into a scalar burden score, for example summing each bundle's Tau integrated intensity multiplied by its solidity and circularity, then normalizing by total cell Tau intensity. This emphasizes bright, compact, inclusion-like Tau bundles while down-weighting diffuse or irregular regions. It is relevant to dense somatic or neuritic Tau inclusions and NFT-like aggregate burden in the 2D Tau MIP. |
| 195 | bundle tau boundary arc discontinuity | code | Quantifies how fragmented Tau enrichment is along the boundary of segmented bundle regions. For each bundle, Tau intensity is sampled around the bundle perimeter, high-intensity perimeter arcs are identified, and the number of separated enriched arcs is normalized by perimeter length before aggregation across bundles. This detects patchy or discontinuous Tau decoration of bundles, distinct from bundle core-to-halo contrast or global anisotropy. |

| # | Feature name | Method | Description |
|---|---|---|---|
| 196 | bundle tau burden fraction | code | Measures the integrated Tau intensity inside the provided bundle mask divided by the integrated Tau intensity inside the whole cell mask. This estimates the fraction of cellular Tau signal contained in thick bundled or aggregate-like structures. It is relevant for advanced Tau pathology where filamentous Tau collapses into dense bundles or NFT-like inclusions. |
| 197 | bundle tau core to halo contrast | code | This feature compares Tau intensity inside segmented bundle cores with Tau intensity in a local surrounding halo within the cell. The bundle mask is dilated to form an annular neighborhood, and a log or ratio contrast between bundle-core and halo intensity is reported. It may capture how strongly Tau is concentrated into bundles relative to nearby cytoplasmic background, distinct from bundle anisotropy. |
| 198 | bundle tau lateral intensity skewness | code | Measures asymmetry of Tau intensity across the width of segmented bundle-like Tau structures. For each bundle component, pixels are projected onto the minor axis of the component, and the intensity-weighted skewness of this lateral coordinate is computed; the final scalar is the mass-weighted mean absolute skewness across bundles. This may capture one-sided rim loading, uneven compaction, or partial bundle maturation that is not captured by bundle area or thickness alone. |
| 199 | filament bundle orientation coherence | code | Measures the directional alignment of elongated Tau structures using region orientations from available mask_filament.tif and mask_bundle.tif. The scalar can be computed as the circular resultant length of doubled component orientations, optionally weighted by component area or skeleton length. High values indicate parallel aligned filaments or bundles, while low values indicate randomly oriented or tangled structures. |
| 200 | filament bundle skeleton length density | code | Measures the total skeletonized length of segmented filament and bundle masks normalized by cell area. The feature is computed on the 2D MIP segmentation by skeletonizing the union of mask_filament and mask_bundle and counting skeleton pixels after optional pruning of tiny fragments. Higher values indicate more extensive fibrillar or thread-like Tau structures within the projected cell. |
| 201 | bundle axis waviness score | vlm | Scores the apparent waviness or tortuosity of Tau-positive bundle centerlines in the 2D MIP using the bundle mask when present. Low values represent straight, bar-like bundles, while high values represent curved, serpentine, or twisted bundle trajectories. This captures large-scale bundle geometry rather than internal strand resolution or contour roughness. |
| 202 | bundle contour raggedness score | vlm | Scores how irregular, lobulated, or ragged the borders of Tau-positive bundle regions appear in the 2D MIP. High values indicate dense aggregates with non-smooth, jagged, or fragmented outlines, while low values indicate compact smooth rounded bundles. This captures aggregate boundary morphology, which is distinct from measuring bundle abundance or intensity. |
| 203 | bundle core hollowing score | vlm | Scores the extent to which Tau bundles show a darker or less fluorescent central core surrounded by brighter Tau signal in the 2D MIP. This differs from bundle strand resolvability because it focuses on hollow or lumen-like intensity organization inside larger bundled structures. Such internal depletion may capture mature, compact, or reorganized Tau assemblies. |
| 204 | bundle cross section striation anisotropy score | vlm | Scores the degree to which thick Tau bundle regions appear internally striated across their width, with visually resolved parallel sub-lines or ridges in the 2D MIP. A high value indicates anisotropic internal cross-sectional texture rather than a smooth homogeneous bundle. This complements bundle fasciculation and strand resolvability by focusing on cross-bundle striation pattern strength. |
| 205 | bundle crossing brightness hierarchy score | vlm | Scores whether Tau bundles or thick filaments crossing other structures show a consistent apparent brightness hierarchy at intersections in the 2D MIP. A high value indicates that one structure visually dominates or remains continuous through crossings, while other strands appear dimmer, interrupted, or subordinate. This may capture layered bundle organization, occlusion-like effects, or dominance of mature Tau cables within tangled networks. |
| 206 | bundle end taper bluntness score | vlm | Measures whether Tau bundle ends appear blunt, swollen, and abruptly terminated versus smoothly tapered into thinner structures. The score is evaluated on the 2D Tau MIP using the bundle mask when available. Blunt or swollen bundle ends may indicate arrested fibrillar growth, aggregate maturation, or structural destabilization. |
| 207 | bundle internal strand resolvability score | vlm | Assesses whether Tau bundle regions contain visually resolvable internal parallel strands, striations, or sub-bundle lanes in the 2D MIP. This complements bundle contour raggedness by focusing on internal organization rather than outer boundary shape, and may reflect different stages of fibril bundling or compaction. |
| 208 | bundle lateral fibril shedding score | vlm | Measures whether thick Tau bundles appear to shed fine fibrillar strands from their lateral sides. A high score indicates wispy, hair-like Tau extensions emerging from bundle edges rather than only from endpoints. This may reflect loosening or remodeling of dense Tau bundle structures. |
| 209 | bundle local kink concentration score | vlm | Scores the concentration of abrupt angular bends or kink points along thick Tau bundles in the 2D MIP. A high value indicates bundles that change direction sharply at localized points rather than showing smooth curvature. Such kinks may reflect mechanical distortion, aggregate compaction, or tangle-like bundle remodeling. |
| 210 | bundle lumen edge asymmetry score | vlm | Assesses whether thick Tau bundles show one-sided brightness or edge dominance, with one margin appearing brighter, sharper, or more fibrillar than the opposite margin. A high score indicates asymmetric bundle internal organization rather than uniformly filled bundles. This may capture polarized fibril packing or partial hollowing from a perspective distinct from bundle width or core appearance alone. |

| # | Feature name | Method | Description |
|---|---|---|---|
| 211 | bundle striation longitudinal drift score | vlm | Scores the degree to which internal bright-dark striations inside Tau bundles drift, warp, or lose phase coherence along the bundle length. This is complementary to cross-sectional striation anisotropy because it evaluates longitudinal stability rather than local stripe directionality. High values may indicate mechanically distorted or partially unraveling Tau bundles. |
| 212 | bundle striation phase locking score | vlm | Measures whether repeating bright and dim striations inside Tau bundles remain phase-aligned across neighboring parallel bundle segments. In the 2D Tau MIP, the score should be high when multiple adjacent bundle lanes show synchronized transverse banding rather than independent or drifting stripe patterns. This may capture coordinated packing or ordered fibrillar bundle architecture. |
| 213 | bundle terminal fanout score | vlm | Measures whether the ends of thick Tau bundles visually split into several thinner filamentous strands or fan-shaped terminal sprays. A high value indicates bundle terminals that disperse into multiple fine Tau structures rather than ending bluntly or remaining compact. This feature captures a bundle-to-filament terminal remodeling pattern not described by overall bundle waviness or internal hollowing. |
| 214 | bundle width varicosity score | vlm | Scores the degree to which Tau bundle masks show repeated local swellings and constrictions along their length in the 2D MIP. A low value corresponds to bundles with relatively uniform width, while a high value corresponds to varicose, sausage-like, or intermittently thickened bundles. Such width modulation may indicate heterogeneous fibril packing or aggregate remodeling. |
| 215 | filament bundle contact angle diversity score | vlm | Measures the visual diversity of angles at which thin Tau filaments contact or merge into thicker Tau bundles in the 2D MIP. Low values indicate mostly parallel merging, while high values indicate many oblique, perpendicular, or fan-like contacts. This captures architectural complexity of filament-bundle interactions without duplicating simple bundle burden or filament crossing counts. |
| 216 | filament bundle disorganization score | vlm | Measures how visually disorganized Tau-positive filaments or bundles appear in the 2D MIP. Low scores correspond to long, aligned, continuous, organized fibers, while high scores correspond to tangled, crossing, irregular, clumped, or poorly oriented bundles. This feature captures structural disruption of Tau-associated filament architecture. |
| 217 | filament bundle entry splay score | vlm | Measures how widely individual Tau filaments fan or splay as they enter visible bundle regions in the 2D MIP. A high score indicates that many filaments approach bundles from diverse angles before merging, while a low score indicates clean, parallel, sharply organized bundle entry. This complements merge-zone fuzziness by focusing on the angular spread and approach geometry of incoming filaments. |
| 218 | filament bundle hierarchical fasciculation score | vlm | Visual score for hierarchical organization where thin Tau filaments appear to merge into intermediate strands and then into thicker bundles within the same cell. High values indicate multi-scale fasciculation rather than a single uniform bundle or isolated filaments, potentially reflecting progressive aggregate assembly. |
| 219 | filament bundle merge zone fuzziness score | vlm | Scores how visually diffuse or ambiguous the transition zones are where thin Tau filaments meet thicker Tau bundles. High values indicate broad, fuzzy, unresolved merge regions rather than crisp contact boundaries, suggesting structural remodeling or aggregation between filament and bundle states. This uses the 2D Tau MIP together with filament and bundle masks when available. |
| 220 | filament bundle partition score | vlm | Scores whether Tau-positive structured signal is visually dominated by thin filament-like masks or by thicker bundle-like masks when the corresponding segmentations are available. High values indicate bundle-dominant organization, while low values indicate mostly fine filamentous organization; if both masks are absent or empty, the score should reflect minimal structured partition burden. This captures maturation from fine fibrillar Tau structures toward coarser bundled aggregates. |
| 221 | filament to bundle orientation mismatch score | vlm | Scores the angular mismatch between thin Tau filaments and thicker Tau bundles within the same cell. Low values indicate filaments run parallel to or merge coherently with bundles, whereas high values indicate crossing, oblique, or disorganized filament orientations relative to bundle axes. This captures structural coordination between fibrillar Tau subtypes rather than measuring either compartment alone. |
| 222 | filament to bundle transition score | vlm | Scores how strongly thin Tau filaments appear to merge into thicker bundles or compact masses within the 2D MIP. High values indicate visible continuity between fine linear structures and dense bundled inclusions, suggesting progressive fibril coalescence. This complements separate filament and bundle burden measures by focusing on the transition morphology between structural classes. |
| 223 | round to fibrillar transition front score | vlm | Scores whether the image shows a spatial transition from round Tau droplets or puncta into elongated fibrillar or bundled structures, forming an apparent maturation front within the cell. This is distinct from simple droplet-bundle contact because it emphasizes a graded morphotype transition across space. High values may indicate Tau phase separation progressing toward fibrillization. |

#### Morphology-defined biological process · Tau-positive object maturation

**Description.** This feature class quantifies segmented Tau-positive high-intensity objects, hotspots, dominant components, object density, size diversity, eccentricity, solidity, and morphotype complexity. It summarizes the burden and morphological maturation of discrete Tau-positive objects in the image, without interpreting them as Tau aggregates in the biological sense.

40 features · 4 subcellular architectures

##### Subcellular architecture · Small puncta, granules, foci, or condensate-like image objects (13 features)

**Approximate scale.** Approximately 0.1-2 um; comparable in order of magnitude to stress granules / cytoplasmic RNP granules and small vesicles/lysosomes.

| # | Feature name | Method | Description |
|---|---|---|---|
| 224 | compact to linear aggregate balance | code | Compares compact droplet-like Tau aggregation to elongated filamentous or bundled Tau aggregation using the provided segmentation masks. It is computed as Tau intensity or area in mask_droplet divided by Tau intensity or area in the union of mask_filament and mask_bundle, with a small pseudocount for stability. The feature may distinguish punctate/oligomer-like aggregation states from fibrillar or tangle-like Tau organization. |
| 225 | high tau component solidity weighted mean | code | Segments high-intensity Tau-positive components and computes their mean solidity weighted by each component's integrated Tau signal. Compact globular inclusions tend to have high solidity, whereas tangled, crescent-like, fragmented, or thread-like aggregates tend to have lower solidity. This feature captures aggregate compactness in a way that is complementary to intensity burden alone. |
| 226 | high tau object shape intensity correlation | code | Segments high-intensity Tau objects inside the cell and computes the robust Spearman correlation between each object's eccentricity or log aspect ratio and its mean Tau intensity. Positive values indicate that brighter aggregates tend to be elongated or rod-like, whereas negative values indicate brighter aggregates tend to be round or globular. This captures aggregate phenotype heterogeneity beyond total aggregate burden or average shape. |
| 227 | intensity weighted aggregate eccentricity | code | Measures the Tau-intensity-weighted mean eccentricity of segmented aggregate components, using available droplet, filament, and bundle masks restricted to the cell. Round puncta contribute low eccentricity, whereas rod-like or elongated inclusions contribute high eccentricity. This captures aggregate shape state and may distinguish compact droplets from filamentous Tau pathology. |
| 228 | tau aggregate mass partition entropy | code | Measures how evenly high-intensity Tau aggregate mass is distributed across detected aggregate components. After segmenting bright Tau components, each component's integrated Tau signal is converted to a probability and summarized by normalized Shannon entropy. Low values indicate dominance by one large NFT-like inclusion, while high values indicate many similarly weighted puncta or fragments. |
| 229 | tau component shape diversity entropy | code | Detects Tau-positive connected components inside the cell and computes the entropy of their shape classes based on circularity, eccentricity, and solidity bins. High entropy indicates coexistence of puncta, rods, irregular aggregates, and filament-like objects, consistent with heterogeneous Tau aggregation states. This measures aggregate morphology diversity rather than aggregate burden alone. |
| 230 | tau positive convexity defect ratio | code | After identifying Tau-positive pixels within the segmented cell using an adaptive within-cell threshold, this feature compares the convex hull area of the Tau-positive region to the actual Tau-positive area. A high value indicates a spatially dispersed, branched, or fragmented Tau pattern with large convexity defects, whereas a compact inclusion has a lower value. This provides a global geometric spread measure distinct from component count or largest-component occupancy. |
| 231 | aggregate size diversity score | vlm | Measures the visual heterogeneity of Tau aggregate sizes within the cell, considering droplets, filaments, bundles, and compact inclusions in the 2D MIP. High scores indicate coexistence of very small puncta, intermediate rods, and large inclusions, while low scores indicate aggregates of relatively uniform scale. Size diversity may reflect mixed stages of Tau aggregation and maturation. |
| 232 | large tau inclusion prominence score | vlm | Scores the visual prominence of one or more large compact Tau-positive inclusions within the cell. The score should increase when a bright aggregate occupies a substantial fraction of the soma or cell area, has a dense clumped appearance, or visually dominates the Tau image. This complements puncta burden by focusing on large NFT-like or conglomerate-like Tau masses rather than small spots. |
| 233 | tau aggregate morphotype complexity | vlm | Visual score for the diversity and complexity of Tau aggregate morphologies in the image, considering mixtures of dots, rods, elongated fibers, branched fibrils, dense bundles, and conglomerate NFT-like masses. Higher values indicate heterogeneous, complex, or hierarchically organized aggregates rather than a single simple punctate pattern. This provides a complementary summary of Tau structural progression in super-resolution MIP images. |
| 234 | tau bright mass compaction index | vlm | Scores whether the brightest Tau signal within the cell is spatially compacted into one or a few concentrated masses versus dispersed across many regions. It is assessed on the 2D Tau MIP within the cell mask and emphasizes the spatial concentration of high-intensity Tau burden. This may capture advanced aggregation states while remaining distinct from simply recognizing a large inclusion. |
| 235 | tau object class local entropy score | vlm | Scores the local intermixing of different Tau object classes—droplets, filaments, and bundles—within the same cytoplasmic neighborhoods. A high value indicates that multiple object types are repeatedly interspersed at short range, while a low value indicates that each object class occupies separate zones. This may reflect heterogeneous aggregation states coexisting within the same cellular microenvironment. |
| 236 | tau object size intensity rank discordance score | vlm | Measures the visual mismatch between Tau object size and brightness across droplets, filaments, bundles, or compact aggregates. A high score indicates that small objects are often among the brightest structures or that large objects are relatively dim, whereas a low score indicates that larger objects are consistently brighter. This may capture heterogeneous aggregate maturation states or differences between compact oligomer-like puncta and diffuse larger deposits. |

#### Subcellular architecture · Composite Tau-positive object morphologies and inter-object relationships (7 features)

**Approximate scale.** Approximately 0.1-20+ um; a multi-scale class spanning small puncta, short filaments, thick bundles and larger objects.

| # | Feature name | Method | Description |
|---|---|---|---|
| 237 | high tau component crowding inverse distance | code | High-Tau cytoplasmic components are detected in the 2D MIP, and the median nearest-neighbor distance between component centroids is normalized by the square root of cytoplasmic area. The feature is reported as an inverse crowding score, with larger values indicating tightly packed Tau puncta or condensate-like clusters. This measures aggregate spatial crowding rather than total aggregate burden. |
| 238 | high tau component radial size sorting | code | Measures whether larger high-Tau components preferentially occur near the nucleus or toward the cell periphery. High-intensity Tau components are detected within the cell, each component area is paired with its normalized distance from the nucleus, and the Spearman correlation between component size and radial position is returned. This captures spatial ordering of aggregate size, distinct from total aggregate burden or nearest-neighbor crowding. |
| 239 | tau high signal object texture dispersion | code | Measures heterogeneity of internal texture among high-Tau connected objects inside the cell. After segmenting high-intensity Tau objects, local intensity entropy or local coefficient of variation is computed within each object, and the robust dispersion of these object-level texture values is returned. This captures whether aggregates are uniformly compact, granular, or internally complex, providing information beyond aggregate size or total area. |
| 240 | tau hotspot mark correlation decay length | code | Estimates the spatial scale over which high-Tau hotspot intensities are correlated. Local maxima or high-Tau connected components are detected within the cell, each assigned a mark such as peak or integrated Tau intensity, and mark similarity is computed as a function of inter-hotspot distance; the decay length of this correlation curve is reported. This captures whether bright aggregates form locally coordinated clusters versus independent puncta. |
| 241 | tau hotspot voronoi territory inequality | code | Measures spatial inequality in the territories surrounding Tau hotspots within the cell. Local Tau intensity maxima or high-Tau component centroids are used as seeds, a clipped Voronoi tessellation is formed inside the cell mask, and the coefficient of variation or Gini coefficient of Voronoi region areas is returned. This captures whether Tau hotspots are evenly dispersed or clustered into crowded pathological zones. |
| 242 | somatic multifocal seed burden score | vlm | A VLM-rated scalar score for how many visually distinct bright Tau seed-like foci are present within the soma, emphasizing separated compact initiation sites rather than one merged inclusion. The score should be low for diffuse somatic Tau without discrete foci and high when multiple independent bright puncta or small clusters appear inside the cell body. This may capture an early-to-intermediate aggregation state complementary to overall somatic inclusion severity. |
| 243 | tau hotspot radial dispersion score | vlm | Scores how widely bright Tau hotspots are dispersed from the nuclear or somatic center toward the cell periphery in the 2D MIP. Low values indicate hotspots clustered near the soma center, whereas high values indicate bright Tau deposits distributed across peripheral cytoplasm and processes. This captures spatial spread of aggregation beyond simple intensity or object count. |

#### Subcellular architecture · Cytoplasmic, somatic, and diffuse intracellular-region scale (12 features)

**Approximate scale.** Approximately 5-30+ um, depending on soma/cytoplasm area; in the 2D MIP this corresponds to large regions of the cell mask excluding the nucleus.

| # | Feature name | Method | Description |
|---|---|---|---|
| 244 | adaptive tau hotspot area fraction | code | Measures the fraction of cytoplasmic pixels whose Tau intensity exceeds an adaptive within-cell threshold, such as median plus two median absolute deviations. This identifies high-Tau hotspot area without requiring a fixed global threshold across patients or imaging sessions. It may capture granular or inclusion-like Tau pathology even when explicit aggregate masks are incomplete. |
| 245 | aggregate component density normalized | code | Measures the number of connected Tau aggregate components per cytoplasmic area using the union of available mask_droplet.tif, mask_filament.tif, and mask_bundle.tif. Binary masks are connected-component labeled, while label masks can be counted directly after restricting to the cell or cytoplasm mask. This captures aggregate burden as object density rather than signal fraction, complementing existing intensity-fraction features. |
| 246 | cytoplasmic tau threshold largest component auc | code | Sweeps a range of within-cytoplasm Tau intensity quantile thresholds and records the fraction of suprathreshold area occupied by the largest connected component at each threshold. The scalar feature is the area under this largest-component persistence curve. It captures whether high Tau signal forms one dominant inclusion-like structure or many fragmented puncta across intensity levels. |
| 247 | dominant aggregate area occupancy | code | Measures the area of the largest segmented Tau aggregate component divided by cytoplasmic area, using the union of available droplet, filament, and bundle masks. This specifically captures whether pathology is dominated by one large inclusion-like Tau structure rather than many small puncta. Large dominant inclusions are relevant to advanced Tau aggregation and soma-filling tangle-like phenotypes. |
| 248 | high tau component contour curvature skewness | code | Quantifies asymmetry in the boundary curvature distribution of high-intensity Tau components in the 2D MIP. For each detected high-Tau component, contour curvature values are estimated along the object boundary and pooled or area-weighted to compute skewness. This distinguishes components with sharp protrusions or invaginations from smoother aggregates, complementing boundary rugosity with a signed curvature-shape statistic. |

| # | Feature name | Method | Description |
|---|---|---|---|
| 249 | segmented aggregate tau fraction | code | Measures the fraction of total cellular Tau fluorescence that falls inside user-provided aggregate-related masks, using the union of mask_droplet, mask_filament, and mask_bundle where available. Missing optional aggregate masks can be treated as empty masks when mask_cell is present, making the feature robust to absent classes. This captures how much Tau is contained in segmented pathological structures rather than diffuse cellular signal. |
| 250 | tau hotspot interior exterior gradient ratio | code | Identifies high-intensity Tau hotspot objects within the cell and compares the average inward-facing intensity gradient at object borders with the outward-facing gradient into surrounding cytoplasm. The scalar is the object-area-weighted mean interior-to-exterior gradient ratio. This captures boundary asymmetry and object compactness in a way that differs from absolute edge sharpness alone. |
| 251 | tau levelset centroid drift index | code | Measures how the center of Tau mass shifts as progressively brighter Tau superlevel sets are examined. The centroids of Tau-positive pixels above several percentile thresholds within the cell are computed, and the total path length of these centroids is normalized by an effective cell radius. Large values indicate that the brightest Tau structures are spatially displaced from the diffuse Tau distribution, consistent with asymmetric inclusions or localized aggregate maturation. |
| 252 | tau superlevel area curve roughness | code | Summarizes the irregularity of the Tau area-survival curve across intensity thresholds in the segmented cell. For a sequence of Tau percentile thresholds, compute the fraction of cell pixels above each threshold and measure the normalized second-difference roughness of this curve. Abrupt changes suggest a mixture of diffuse Tau and discrete bright inclusions, whereas smoother curves indicate a more continuously graded Tau distribution. |
| 253 | tau superlevel component burstiness | code | Thresholds the Tau signal inside the cell across a sequence of robust intensity quantiles and records the number of connected superlevel components at each threshold. The feature is the largest positive jump in component count divided by the median component count across thresholds. It captures abrupt fragmentation of Tau into many puncta over a narrow intensity range, which may reflect aggregation-state transitions distinct from persistence-area summaries. |
| 254 | tau superlevel component persistence auc | code | Summarizes how long connected high-Tau components persist across a range of increasing intensity thresholds within the cell mask. At each threshold quantile, connected Tau-positive components are counted after small-object removal, and the area under the component-count-versus-threshold curve is reported. Persistent component structure may reflect stable puncta or aggregates rather than threshold-specific noise. |
| 255 | somatic inclusion severity visual | vlm | Visual score of dense Tau-positive inclusion burden inside the soma, considering apparent number, size, brightness, and dominance of compact somatic aggregates. Higher values correspond to large or multiple bright inclusions, NFT-like masses, or aggregates occupying much of the cell body. This captures advanced Tau aggregation phenotypes that may not be well represented by average intensity alone. |

#### Subcellular architecture · Cell boundary, cortical, peripheral-shell, and extracellular-neighborhood scale (8 features)

**Approximate scale.** Set against a whole-cell diameter of ~10-100 um; the measured region is typically a thin band or boundary arc inside/outside the cell edge.

| # | Feature name | Method | Description |
|---|---|---|---|
| 256 | high tau component boundary distance skew | code | Computes the skewness of the distribution of high-Tau component centroid distances to the nearest cell boundary. Components are detected inside the cell mask, their centroid-to-boundary distances are normalized by local cell size, and intensity-weighted skewness is reported. This captures whether aggregates preferentially accumulate near the cell edge, deep cytoplasm, or both in an asymmetric distribution. |
| 257 | high tau component boundary rugosity | code | This feature detects high-intensity Tau components inside the cytoplasm using a robust local or within-cell threshold, then computes an intensity-weighted mean boundary rugosity such as perimeter squared divided by area. Smooth round puncta have lower rugosity, while irregular tangle-like or jagged aggregates have higher values. It captures aggregate boundary complexity rather than aggregate size or eccentricity alone. |
| 258 | tau high intensity object edge sharpness | code | This feature measures how sharply high-intensity Tau objects transition into their local background. High-tail Tau objects are detected within the cell mask, their boundaries are extracted, and the mean gradient magnitude along object edges is normalized by local object intensity. Sharp-edged condensates or inclusions should score higher than diffuse fuzzy Tau accumulations. |
| 259 | somatic inclusion edge crispness score | vlm | Assesses how sharply bounded the brightest somatic Tau inclusions appear relative to surrounding cytoplasmic Tau in the 2D MIP. High values indicate compact, well-demarcated aggregates, whereas low values indicate diffuse or gradually fading accumulations; this may separate mature compact inclusions from early diffuse Tau buildup. |
| 260 | tau aggregate depletion halo score | vlm | Visual scalar score for bright Tau aggregates that are surrounded by locally dimmer cytoplasmic halos or depletion zones. High values suggest Tau sequestration into dense inclusions with reduced diffuse Tau immediately around them, a pattern distinct from uniformly bright cytoplasm. |

| # | Feature name | Method | Description |
|---|---|---|---|
| 261 | tau object boundary feathering directionality score | vlm | Scores whether edges of bright Tau domains show directional feathering, with wispy extensions preferentially pointing toward neurites, bundles, droplets, or the cell edge. The VLM should distinguish directional feathered margins from uniformly soft edges or random noise. This may reflect active spreading or remodeling of Tau aggregates in the 2D MIP. |
| 262 | tau object boundary interdigitation score | vlm | Assesses how strongly droplets, filaments, and bundles visually interleave through shared or closely apposed boundaries within the Tau-positive region. A high value indicates a mixed morphology where object classes are spatially intertwined, rather than segregated into separate droplet-rich, filament-rich, or bundle-rich zones. This may capture transitions between condensate-like and fibrillar Tau states. |
| 263 | tau object edge polarity reversal score | vlm | Measures whether Tau-positive objects show opposite edge behavior on different sides, such as one side having a sharp bright boundary while the opposite side fades into diffuse haze. A high score indicates strong directional edge polarity or reversal across aggregates, bundles, or dense Tau domains in the 2D MIP. This differs from general boundary feathering by emphasizing paired contrast between opposing sides of the same Tau object. |

#### Morphology-defined biological process · Tau boundary organization

**Description.** This feature class specifically describes boundaries, gradients, rim/core organization, halos, edge softness, shells, domain edges, interfaces, and cell-periphery rims of Tau-positive structures. It emphasizes material-state-like and phase-boundary-like properties between Tau-enriched regions and the surrounding cytoplasm, without assuming bona fide aggregate formation.

15 features · 3 subcellular architectures

##### Subcellular architecture · Local texture and pixel-neighborhood scale (2 features)

**Approximate scale.** Approximately 0.2-1 um imaging texture scale; the lower bound is constrained by conventional fluorescence microscopy lateral resolution (~200 nm).

| # | Feature name | Method | Description |
|---|---|---|---|
| 264 | diffuse tau contour conformity score | vlm | Visual score for how well the diffuse Tau cloud conforms to the overall cell body and proximal neurite outline. Low conformity suggests patchy underfilling, retracted Tau distribution, or collapsed intracellular Tau domains, while high conformity indicates broad diffuse Tau occupying the expected cytoplasmic shape. |
| 265 | tau domain edge softness heterogeneity score | vlm | Measures the diversity of boundary sharpness among Tau-positive domains, from fuzzy diffuse edges to crisp compact inclusion borders. The scalar increases when a single cell contains a mixture of soft-edged haze, semi-defined patches, and sharply bounded aggregates. This may reflect coexistence of soluble, condensate-like, and more mature aggregated Tau states. |

##### Subcellular architecture · Cell boundary, cortical, peripheral-shell, and extracellular-neighborhood scale (5 features)

**Approximate scale.** Set against a whole-cell diameter of ~10-100 um; the measured region is typically a thin band or boundary arc inside/outside the cell edge.

| # | Feature name | Method | Description |
|---|---|---|---|
| 266 | cell boundary tau penetration depth | code | Measures the Tau-intensity-weighted average inward distance from the cell boundary within the cell mask. Low values indicate Tau concentrated near the membrane or thin peripheral processes, while high values indicate deeper somatic or central accumulation. This provides a continuous complement to periphery enrichment and contact-based features. |
| 267 | cell edge tau halo excess ratio | code | Compares Tau signal immediately outside the cell boundary to Tau signal immediately inside the cell boundary in the 2D MIP. The feature is the mean Tau intensity in a thin extracellular dilation ring around mask_cell divided by the mean Tau intensity in a matched inner cell-edge ring. It may capture pericellular Tau debris, released aggregate material, or edge-associated pathological Tau while remaining distinct from intracellular aggregate burden. |
| 268 | cell edge tau inward gradient skewness | code | Captures asymmetry in Tau intensity gradients moving inward from the cell boundary into the cytoplasm. Tau intensity is sampled along short inward normals or distance-transform shells from the cell edge, the first derivative profile is computed, and the skewness of gradient values across boundary locations is reported. This detects whether Tau entry from the cell periphery is uniform or concentrated in sharp localized boundary-associated patches. |
| 269 | tau cell edge arc run entropy | code | Quantifies how fragmented or continuous high Tau signal is along the inner cell perimeter. Tau intensity is sampled in a narrow band just inside the cell boundary, thresholded relative to cytoplasmic background, and contiguous high-Tau boundary arc lengths are summarized by Shannon entropy. This captures whether Tau accumulates as a few long peripheral arcs or many broken edge-associated patches. |
| 270 | cell periphery tau rim sharpness score | vlm | Scores how sharply Tau fluorescence is enriched along the outer cell boundary in the 2D MIP using the cell mask. A high score indicates a crisp peripheral Tau rim or cortical band, while a low score indicates diffuse cytoplasmic signal with no clear edge enrichment. This captures boundary-localized Tau organization distinct from soma-edge leakage or general cytoplasmic accumulation. |

##### Subcellular architecture · Whole-cell geometry, global polarity, and spatial-distribution scale (8 features)

**Approximate scale.** Approximately 10-100+ um; larger if the full neurite arbor is included in the 2D projection.

| # | Feature name | Method | Description |
|---|---|---|---|
| 271 | boundary band tau fourier peak fraction | code | Samples Tau intensity around the inner cell-boundary band of the 2D MIP and computes the fraction of angular Fourier power contained in the strongest nonzero frequency. High values indicate periodically spaced Tau-rich patches along the cell edge, while low values indicate either diffuse boundary signal or irregular patch placement. This complements boundary arc entropy by focusing on periodic spatial organization rather than run complexity. |
| 272 | cytoplasm tau entropy radial gradient | code | Computes a local Shannon entropy map of the Tau MIP within cytoplasm, then fits the slope of entropy versus normalized radial distance from the nuclear boundary toward the cell boundary. A positive value indicates more heterogeneous or granular Tau texture toward the periphery, while a negative value indicates stronger perinuclear texture. This complements intensity radial features by focusing on spatial disorder rather than mean signal. |
| 273 | cytoplasmic tau gradient magnitude gini | code | Computes the Gini coefficient of Tau gradient magnitudes within the cytoplasm. A high value indicates that sharp Tau transitions are concentrated in a small subset of pixels, as expected for compact inclusions, rimmed aggregates, or sharp puncta, whereas diffuse accumulation produces a more even gradient distribution. This complements intensity Gini by focusing on spatial edges rather than brightness values themselves. |
| 274 | cytoplasmic tau gradient orientation entropy | code | Computes Sobel or Scharr gradients of the Tau MIP within the cytoplasm and forms a gradient-orientation histogram weighted by gradient magnitude. The scalar output is the circular entropy of this orientation distribution. Low entropy indicates strongly aligned fibrillar or bundle-like Tau organization, while high entropy indicates isotropic granular or diffuse cytoplasmic Tau texture. |
| 275 | tau boundary distance information coupling | code | Computes the normalized mutual information between Tau intensity and distance to the cell boundary within the 2D cell mask. Unlike a simple radial gradient, this detects any structured dependence of Tau signal on boundary proximity, including pericellular rims, central accumulations, or non-monotonic bands. It may capture pathological redistribution of Tau toward or away from the neuronal periphery. |
| 276 | tau boundary distance isotonic fit error | code | This feature evaluates how well Tau intensity can be explained as a monotonic function of normalized distance from the cell boundary. It fits an isotonic regression between cytoplasmic Tau intensity and inward boundary distance, then reports the normalized residual error. High error indicates non-monotonic, patchy, or compartmentalized Tau organization, while low error indicates a smooth peripheral-to-central gradient. |
| 277 | tau boundary distance quantile spline curvature | code | Within the cell mask, bin pixels by normalized distance from the cell boundary and compute a robust Tau intensity quantile, such as the 75th percentile, in each bin. Fit or discretely approximate the resulting distance-intensity curve and report the summed absolute second differences. This measures non-monotonic layering of Tau from edge to interior, complementing simpler inward-gradient or boundary-enrichment features. |
| 278 | tau radial shell gradient score | vlm | Scores the apparent radial organization of Tau signal from the nucleus outward toward the cell boundary in the 2D MIP. High positive values correspond to strong peripheral or outer-cytoplasmic enrichment, while low values correspond to central or perinuclear dominance without a clear outward gradient. This captures broad subcellular redistribution of Tau using a radial-shell perspective rather than a single compartment dominance score. |

#### Morphology-defined biological process · Proteostasis stress

**Description.** This feature class includes texture statistics such as lacunarity, Gini index, entropy, wavelet/spectral features, gray-level co-occurrence matrix metrics, variogram measures, local contrast, haze, mosaicity, and tail-index features. It describes the transition of cytoplasmic Tau signal from a relatively homogeneous diffuse pattern to more patchy, granular, or haze-like heterogeneity.

45 features · 3 subcellular architectures

##### Subcellular architecture · Local texture and pixel-neighborhood scale (21 features)

**Approximate scale.** Approximately 0.2-1 um imaging texture scale; the lower bound is constrained by conventional fluorescence microscopy lateral resolution (~200 nm).

| # | Feature name | Method | Description |
|---|---|---|---|
| 279 | cytoplasm tau directional variogram anisotropy | code | This feature computes rank-normalized Tau semivariograms within the cytoplasm along several 2D directions and reports the ratio or contrast between the strongest and weakest directional semivariance. High anisotropy suggests oriented fibrils, bundles, or elongated Tau structures, while low anisotropy suggests isotropic punctate or diffuse Tau texture. It provides a directional spatial-correlation measurement distinct from local gradient orientation entropy or run-length nonuniformity. |
| 280 | cytoplasm tau gray run length nonuniformity | code | Quantize Tau intensities inside the cytoplasm and compute a gray-level run-length matrix over several 2D directions. The scalar is the normalized gray-level nonuniformity averaged across directions. It captures whether cytoplasmic Tau texture is dominated by repeated similar-intensity runs, streaks, and bands versus heterogeneous granular patches. |
| 281 | cytoplasm tau log gabor orientation dispersion | code | This feature applies a bank of oriented log-Gabor or steerable filters to the Tau signal within the cytoplasm and summarizes the circular dispersion of orientation-specific energy. Low dispersion indicates aligned fibrillar or bundle-like Tau organization, while high dispersion indicates isotropic granular or tangled Tau signal. This may capture super-resolution Tau filament organization and mesh-like pathology not fully represented by simple intensity or object counts. |

| # | Feature name | Method | Description |
|---|---|---|---|
| 282 | cytoplasmic tau bandpass granularity ratio | code | Quantifies small-to-intermediate scale Tau granularity in the cytoplasm of the 2D MIP by applying bandpass filters such as Difference-of-Gaussians or Laplacian-of-Gaussian at biologically plausible puncta scales. The feature is the bandpass energy normalized by total cytoplasmic Tau intensity or low-frequency energy. It captures granular or punctate Tau pathology while remaining distinct from simple high-intensity area fractions or global intensity inequality. |
| 283 | cytoplasmic tau coarse to fine wavelet ratio | code | This feature measures the ratio of large-scale to small-scale Tau texture energy within the cytoplasm, defined as the cell mask excluding the nucleus mask. The 2D MIP Tau image is decomposed with multiscale wavelet or difference-of-Gaussian filters, and coarse-scale energy is divided by fine-scale energy. It may distinguish broad diffuse cytoplasmic Tau accumulation from fine granular or punctate Tau pathology. |
| 284 | cytoplasmic tau fine energy coarse intensity correlation | code | Measures whether fine-scale Tau texture is preferentially located in bright coarse-scale Tau regions. Within the cytoplasm, compute a fine-detail energy map using a high-pass or wavelet detail filter and a coarse Tau intensity map using Gaussian smoothing, then report their Spearman correlation. This distinguishes diffuse bright haze from bright regions that contain embedded punctate or fibrillar substructure. |
| 285 | cytoplasmic tau glcm homogeneity anisotropy | code | Measures directional anisotropy of local Tau texture in the cytoplasm using gray-level co-occurrence matrices. After robust intensity quantization within the cytoplasmic mask, GLCM homogeneity is computed at several orientations, and the feature is the coefficient of variation across orientations. This may capture oriented fibrillar or bundled Tau texture that differs from isotropic granular cytoplasmic accumulation. |
| 286 | cytoplasmic tau local contrast skewness | code | Computes the skewness of a local contrast map within the cytoplasm of the 2D Tau MIP. Local contrast can be estimated using a sliding-window robust standard deviation or percentile range normalized by local median intensity. High positive skewness indicates rare sharply contrasted inclusions or puncta embedded in otherwise smoother Tau signal, which may mark pathological aggregation. |
| 287 | cytoplasmic tau local dynamic range | code | Computes the median local intensity dynamic range within cytoplasm, for example the local 90th minus 10th percentile Tau intensity in sliding windows inside the cell excluding nucleus. High values indicate locally heterogeneous granular or punctate Tau signal, while low values indicate smoother diffuse Tau. This captures cytoplasmic texture at a local scale without relying only on global intensity distribution. |
| 288 | cytoplasmic tau moran autocorrelation | code | Measures spatial autocorrelation of Tau intensity within the cytoplasm using a local-neighborhood Moran's I statistic on the 2D MIP. High values indicate clustered or spatially coherent Tau domains, whereas lower values indicate more random or fine-grained intensity variation. This captures Tau spatial organization from a statistical perspective distinct from radial gradients, puncta scores, or aggregate fractions. |
| 289 | cytoplasmic tau multiscale mad slope | code | Applies Gaussian smoothing at multiple spatial scales to the cytoplasmic Tau signal and computes the median absolute deviation of the residual high-pass image at each scale. The feature is the slope of log residual MAD versus log scale. It distinguishes fine granular Tau punctateness from broader diffuse heterogeneity using robust dispersion rather than wavelet energy ratios. |
| 290 | cytoplasmic tau phase congruency density | code | Measures the fraction of cytoplasmic pixels with high multiscale phase congruency or phase-symmetric edge/ridge response in the Tau MIP. Unlike raw gradient magnitude, phase congruency is relatively invariant to absolute brightness and highlights coherent ridges, sharp aggregate borders, and mesh-like Tau structures. This may help separate diffuse Tau accumulation from structurally organized pathological Tau. |
| 291 | cytoplasmic tau rank variogram slope | code | Computes a spatial variogram of rank-transformed Tau intensities within the cytoplasm at multiple 2D pixel lags and fits the log-log slope of semivariance versus distance. A steep slope indicates smooth large-scale organization, whereas a flatter or rapidly saturating profile indicates fine-grained heterogeneity and fragmented Tau texture. Rank transformation reduces sensitivity to staining intensity while preserving spatial ordering. |
| 292 | cytoplasmic tau rof residual fraction | code | Quantifies the fraction of cytoplasmic Tau signal present as fine punctate or granular residual after edge-preserving total-variation denoising of the 2D MIP. The Tau image inside the cytoplasm is decomposed into a smooth component and a residual component, and the feature is the robust residual energy divided by total cytoplasmic Tau energy. Higher values indicate speckled or granular Tau pathology beyond diffuse cytoplasmic accumulation. |
| 293 | tau cell glcm correlation decay length | code | Estimates the spatial correlation length of Tau texture within the segmented cell using gray-level co-occurrence matrix correlation at multiple pixel offsets. A longer decay length indicates broad smooth Tau domains, while a shorter decay length indicates fine-grained speckled or fragmented Tau organization. This provides a texture-autocorrelation view of Tau distribution that is complementary to intensity heterogeneity and puncta-count features. |
| 294 | tau granularity small to large scale ratio | code | Measures the ratio of small-scale Tau granularity to large-scale diffuse Tau structure within the cell mask. It can be computed as the summed positive Laplacian-of-Gaussian or bandpass response at puncta-like scales divided by the low-frequency Gaussian-smoothed Tau signal. This distinguishes granular or stress-granule-like Tau patterns from broad diffuse accumulation in the 2D Tau MIP. |

| # | Feature name | Method | Description |
|---|---|---|---|
| 295 | tau midband spectral flatness | code | Computes the spectral flatness of Tau intensity fluctuations in an intermediate spatial-frequency band from the 2D MIP without requiring object segmentation. After background normalization and optional rank transformation, the image is windowed, Fourier transformed, and the geometric-to-arithmetic mean ratio of annular midband power is returned. Higher flatness suggests broadband granular or noisy Tau organization, whereas lower flatness suggests dominant structured periodic or smooth spatial organization. |
| 296 | tau multiscale granularity ratio | code | Computes the ratio of small-scale Tau granularity energy to larger-scale Tau structure energy within the cell mask, for example using Difference-of-Gaussian or Laplacian-of-Gaussian filter responses at two spatial scales. High values indicate fine punctate or speckled Tau, while lower values indicate smoother diffuse signal or larger aggregates. This captures puncta-like Tau organization in the 2D MIP without relying solely on connected-component segmentation. |
| 297 | tau rank power spectrum slope | code | Measures the slope of the radially averaged Fourier power spectrum of the rank-transformed Tau MIP. Rank transformation reduces sensitivity to absolute brightness and saturation, while the spectral slope summarizes whether Tau texture is dominated by coarse diffuse structures or fine punctate patterns. This image-wide feature can be computed without object segmentation and may capture global pathological granularity complementary to compartment-based metrics. |
| 298 | tau wavelet scale energy entropy | code | Computes the normalized Shannon entropy of Tau-channel wavelet energy across multiple 2D spatial scales within the cell or cytoplasm mask. A low value indicates Tau texture dominated by one characteristic scale, while a high value indicates mixed diffuse, granular, and aggregate-scale structure. This complements coarse-to-fine wavelet ratios by measuring scale diversity rather than a single scale contrast. |
| 299 | somatic tau texture scale mixing score | vlm | Scores the degree to which somatic cytoplasmic Tau contains multiple texture scales simultaneously, such as fine speckles, intermediate droplets, and broad diffuse patches in the same cell body. This is distinct from microgranule density because it emphasizes coexistence of several aggregate scales rather than only small puncta. High scale mixing may reflect heterogeneous stages of Tau condensation within the soma. |

#### Subcellular architecture · Composite Tau-positive object morphologies and inter-object relationships (10 features)

**Approximate scale.** Approximately 0.1-20+ um; a multi-scale class spanning small puncta, short filaments, thick bundles and larger objects.

| # | Feature name | Method | Description |
|---|---|---|---|
| 300 | cell tau boxcount fractal dimension | code | Estimates the box-counting fractal dimension of adaptively thresholded Tau-positive pixels within the cell mask on the 2D MIP. Higher values indicate more space-filling, branched, or mesh-like Tau organization, while lower values indicate sparse puncta or compact inclusions. This complements Euler and lacunarity features by quantifying scale-dependent occupancy complexity. |
| 301 | cytoplasmic tau multiscale lacunarity | code | Measures multiscale spatial gap heterogeneity of Tau fluorescence within the cytoplasmic cell region, excluding the nucleus. Box-counting lacunarity is computed over several 2D window sizes on the Tau MIP restricted to mask_cell minus mask_nucleus, then summarized as an average or area under the lacunarity curve. High values capture patchy, honeycomb-like, granular, or void-containing Tau organization rather than smooth diffuse signal. |
| 302 | cytoplasmic tau percolation threshold span | code | This feature measures how gradually Tau-positive pixels form connected structures as the intensity threshold is swept within the cytoplasm. It is defined as the normalized intensity range over which the largest connected Tau component grows from a low occupancy fraction to a higher occupancy fraction. Diffuse mesh-like Tau should percolate differently from isolated puncta or compact inclusions. |
| 303 | pericellular tau contact fraction | code | Measures the fraction of the cell boundary that is adjacent to locally high Tau signal within a thin inner peripheral band. Unlike a periphery enrichment ratio, this feature asks whether Tau contacts the cell edge broadly around the perimeter or only in sparse focal regions. It may capture peripheral Tau accumulation, neuritic attachment zones, or edge-associated aggregate spread in the 2D MIP. |
| 304 | tau lacunarity cell texture | code | Measures lacunarity of the Tau signal within the cell mask using box-counting on a locally thresholded or intensity-weighted Tau image. High lacunarity reflects heterogeneous gaps, voids, and clustered bright structures, consistent with reticular or honeycomb-like Tau organization. This complements mean intensity and granularity by quantifying spatial gap structure in the 2D MIP. |
| 305 | tau microterritory mass entropy | code | The cytoplasmic region is partitioned into a fixed number of approximately equal-area spatial microterritories in the 2D MIP, such as by distance-ordered grid bins restricted to the cell mask. Tau mass is summed in each territory and converted to a normalized entropy. Low values indicate Tau concentrated into a few local domains, while high values indicate spatially diffuse cytoplasmic distribution. |
| 306 | tau superpixel mosaic transition entropy | code | Partitions the cell area into compact superpixels on the Tau MIP, assigns each superpixel to an intensity state such as low, medium, or high, and computes the entropy of neighboring-state transitions across the superpixel adjacency graph. High transition entropy indicates a mosaic-like mixture of diffuse, punctate, and bright regions, while low entropy indicates smoother compartmental organization. This provides a coarse spatial-organization descriptor complementary to pixel-level texture metrics. |

| # | Feature name | Method | Description |
|---|---|---|---|
| 307 | cytoplasmic tau mosaic irregularity score | vlm | Measures how strongly the cytoplasmic Tau signal appears as an irregular mosaic of bright and dim territories rather than a smooth diffuse field or a few isolated aggregates. The feature focuses on spatially extended patchwork heterogeneity inside the cell mask while excluding the nucleus. It may capture intermediate Tau redistribution states linked to cellular stress and aggregation-prone cytoplasmic organization. |
| 308 | somatic tau patch domain scale score | vlm | Scores whether somatic Tau is organized into large patch-like intensity domains rather than fine granules, diffuse haze, or single compact inclusions. The feature uses the 2D MIP Tau channel within the cell mask and may capture intermediate-scale cytoplasmic Tau redistribution linked to altered protein aggregation state. |
| 309 | tau cytoplasmic lacunarity index | vlm | Scores the prominence of dark voids, gaps, or lacunae within Tau-positive cytoplasm after excluding the nucleus in the 2D MIP Tau image. It captures porous or sponge-like organization as a spatial texture property distinct from overall haze or simple honeycomb appearance, and may indicate dense aggregate remodeling of the soma. |

#### Subcellular architecture · Cytoplasmic, somatic, and diffuse intracellular-region scale (14 features)

**Approximate scale.** Approximately 5-30+ um, depending on soma/cytoplasm area; in the 2D MIP this corresponds to large regions of the cell mask excluding the nucleus.

| # | Feature name | Method | Description |
|---|---|---|---|
| 310 | cell tau gini heterogeneity | code | Computes the Gini coefficient of Tau pixel intensities within the cell mask on the 2D MIP. Higher values indicate unequal Tau distribution with bright puncta, aggregates, bundles, or localized inclusions against lower diffuse background. This provides a robust scalar measure of intracellular Tau heterogeneity independent of explicit object counting. |
| 311 | cell tau intensity gini | code | Calculates the Gini coefficient of Tau intensities over all pixels inside mask_cell. High values indicate that Tau signal is concentrated into a small subset of bright pixels, consistent with puncta, aggregates, or compact inclusions, while lower values indicate diffuse signal. This captures intensity inequality independent of absolute staining level. |
| 312 | cytoplasm tau peak prominence gini | code | Detect local Tau maxima in the cytoplasm after mild background correction and estimate each peak's prominence relative to its local neighborhood. The feature is the Gini coefficient of these peak prominences. It distinguishes cells with one or a few dominant inclusions from cells with many similarly prominent puncta, both of which can have similar total high-intensity Tau burden. |
| 313 | cytoplasm tau tail balance ratio | code | Measures the asymmetry between bright-tail and dim-tail Tau intensity spread within the cytoplasm, defined for example as (Q95 - Q75) / (Q25 - Q05) after background correction and robust normalization. It is computed on the 2D Tau MIP inside the cell mask excluding the nucleus. This may capture whether pathology is dominated by high-intensity puncta or aggregates versus broad diffuse cytoplasmic signal and low-signal voids. |
| 314 | cytoplasmic tau bimodality coefficient | code | Computes the bimodality coefficient of Tau intensities within the cytoplasmic mask using skewness and kurtosis. Higher values suggest separation between diffuse background Tau and bright aggregate-like pixels, while lower values suggest a more unimodal diffuse distribution. This captures diffuse-versus-punctate transition in a distributional way distinct from simple upper-tail contrast. |
| 315 | cytoplasmic tau extreme value tail index | code | Quantifies how heavy the brightest tail of cytoplasmic Tau intensities is within the 2D MIP. It can be computed as a Hill-type tail index or log-slope using the top few percent of cytoplasmic Tau pixels after robust background normalization. This captures rare very bright Tau condensates or inclusions while being distinct from total intensity or mean cytoplasmic brightness. |
| 316 | cytoplasmic tau intensity gini | code | Computes the Gini coefficient of Tau fluorescence intensities across all cytoplasmic pixels in the 2D MIP. Low values indicate diffuse, relatively homogeneous Tau, whereas high values indicate that Tau signal is concentrated into a subset of bright pixels such as puncta, droplets, or inclusions. This captures intensity inequality in a way that is complementary to hotspot area or upper-tail contrast features. |
| 317 | cytoplasmic tau smooth haze fraction | code | Measures the fraction of cytoplasmic Tau signal carried by a smooth low-frequency component after background correction and removal of bright punctate or aggregate-like residuals. It is computed on the 2D Tau MIP within the cell mask excluding the nucleus, using Gaussian smoothing or morphological opening to estimate diffuse haze. This may capture diffuse somatic Tau mislocalization that is biologically distinct from compact aggregates or filamentous inclusions. |
| 318 | cytoplasmic tau upper tail contrast | code | Measures the robust upper-tail intensity contrast of Tau within the cytoplasm, for example as the cytoplasmic 95th percentile divided by the cytoplasmic median plus a small offset. It captures whether a cell contains unusually bright Tau-enriched subregions beyond the diffuse cytoplasmic background. This is complementary to aggregate mask fractions because it is computed from raw Tau intensity distribution inside the cytoplasm. |
| 319 | high tau area fraction in cell | code | Computes the fraction of cell-mask area occupied by unusually bright Tau pixels, using a robust within-cell threshold such as median plus three median absolute deviations. This estimates the burden of high-intensity Tau deposits without depending on a separate aggregate mask. It is relevant to compact Tau inclusions and hyperintense pathological puncta in the 2D MIP. |
| 320 | robust high tau area fraction | code | Measures the fraction of cell-mask pixels whose Tau intensity exceeds a robust within-cell threshold such as median plus three median absolute deviations. This estimates the area occupied by high-intensity Tau accumulations while adapting to sample-specific staining intensity. It is relevant for detecting compact aggregates, bright somatic inclusions, and high-burden Tau regions in the MIP. |

| # | Feature name | Method | Description |
|---|---|---|---|
| 321 | intracellular tau heterogeneity score | vlm | Measures the overall visual heterogeneity of Tau signal within the segmented cell in the 2D MIP. A low score corresponds to smooth diffuse Tau distribution, while a high score corresponds to patchy, mottled, granular, locally saturated, or spatially uneven Tau fluorescence. This feature captures mixed diffuse-and-aggregate phenotypes that may not be represented by a single compartment intensity or object count. |
| 322 | somatic diffuse tau haze score | vlm | Visual scalar estimate of diffuse, non-punctate Tau brightness within the somatic cytoplasm of the 2D MIP. Higher values indicate a broad hazy or cloud-like somatic Tau signal rather than a dark or weakly labeled soma. Diffuse somatic Tau accumulation can represent an early pathological redistribution state before formation of dense inclusions. |
| 323 | tau peak to haze balance score | vlm | Scores the visual balance between sharp punctate Tau peaks and broad diffuse Tau haze within the cell. Higher values indicate dominance of discrete bright peaks over diffuse background, while lower values indicate mostly smooth or carpet-like Tau signal with few distinct peaks. This complements puncta-burden and diffuse-accumulation concepts by explicitly measuring their relative visual mixture. |

#### Morphology-defined biological process · Cytoplasmic compartment disruption

**Description.** This feature class focuses on negative-space and network-like patterns, including mesh-like, reticular or honeycomb structures, voids, pores, holes, channels, corridors, gaps, basins, and barriers. It reflects spatial partitioning, exclusion, and channelization of the cytoplasm by Tau-enriched regions.

22 features · 2 subcellular architectures

##### Subcellular architecture · Filaments, skeletons, ridges, or short linear structures (5 features)

**Approximate scale.** Biological fiber diameters are often nanometer-scale (actin ~7 nm, intermediate filaments ~8-11 nm, microtubules ~25 nm); in optical images they appear as linear/skeletonized structures >=~0.2 um wide, with lengths of several um or more.

| # | Feature name | Method | Description |
|---|---|---|---|
| 324 | cytoplasmic tau channel connectivity score | vlm | Measures whether low-Tau cytoplasmic channels form continuous paths from the perinuclear region toward the cell periphery. A high score indicates visually connected Tau-poor corridors cutting through otherwise Tau-positive cytoplasm, rather than isolated small voids. This captures the topology of cytoplasmic exclusion spaces in the 2D Tau MIP. |
| 325 | cytoplasmic tau isthmus connectivity score | vlm | Scores the extent to which separated Tau-rich cytoplasmic islands are linked by narrow bridge-like isthmuses within the cell mask. In the 2D Tau MIP, high values indicate a connected but constricted network of bright Tau domains, while low values indicate either diffuse signal or clearly isolated deposits. This feature captures an intermediate aggregation architecture distinct from simple puncta burden or whole-cell connectivity. |
| 326 | radial tau corridor alignment score | vlm | Scores whether Tau-rich structures form visually coherent radial corridors extending from the perinuclear cytoplasm toward the cell periphery. High values indicate aligned channels or streaks of Tau connecting the nucleus-adjacent region to outer cell regions, while low values indicate random, circumferential, or patchy organization. This captures spatial organization of Tau transport-like paths in the 2D cell projection. |
| 327 | tau free cytoplasmic corridor continuity score | vlm | Measures the visual continuity of Tau-poor corridors or channels running through the cytoplasm between Tau-rich aggregates, droplets, filaments, or bundles. The feature is computed conceptually from the 2D Tau MIP within the cell mask and excludes the nucleus when available. It captures organized cytoplasmic void pathways rather than simply the size or heterogeneity of isolated Tau-negative holes. |
| 328 | tau gap skip pattern severity score | vlm | A scalar visual score for long irregular dark gaps interrupting otherwise Tau-positive neurite-like structures. The score should be high when processes show skip-like labeling with alternating bright segments and extended signal voids, rather than uniformly weak or smoothly continuous Tau. This may capture loss of axonal or dendritic Tau continuity in a way distinct from simple beading or distal continuity assessment. |

##### Subcellular architecture · Reticular, honeycomb, pore/void, and cytoplasmic-domain patterns (17 features)

**Approximate scale.** Approximately 1-20+ um; composed of multiple pores, low-signal corridors, mesh walls or Tau-positive/negative cytoplasmic domains.

| # | Feature name | Method | Description |
|---|---|---|---|
| 329 | cytoplasmic tau void size entropy | code | Identifies low-Tau void-like regions inside the cytoplasmic mask after excluding the nucleus and summarizes the diversity of their connected-component areas using Shannon entropy. High entropy indicates a heterogeneous mesh with both small and large dark holes, while low entropy indicates uniform holes or absence of void structure. This complements total hole-area measurements by describing the size distribution of internal Tau-free spaces. |
| 330 | tau dark basin depth entropy | code | This feature analyzes low-signal basins inside the cytoplasmic Tau field by applying watershed-like basin detection to the inverted, smoothed Tau image and computing the entropy of basin depth values. It captures whether Tau forms a uniform haze, a few dominant voids, or many heterogeneous dark holes within a bright reticular pattern. Such dark-basin organization may reflect honeycomb-like Tau networks or cytoplasmic exclusion zones. |

| # | Feature name | Method | Description |
|---|---|---|---|
| 331 | tau honeycomb wall hole contrast | code | Detects low-intensity cytoplasmic holes after local smoothing and compares their Tau intensity to the surrounding narrow rim or wall region. The feature is the median log ratio of surrounding-wall Tau intensity to internal-hole Tau intensity across detected holes, with zero returned when no valid holes are detected. It is designed to capture STED-like honeycomb or mesh organization where bright Tau walls surround darker voids. |
| 332 | tau mesh hole area fraction | code | Estimates the fraction of cytoplasmic area occupied by low-Tau holes enclosed within locally high-Tau mesh-like signal. The feature is designed to capture reticular or honeycomb-like Tau organization in super-resolution 2D MIP images, where bright Tau ridges may surround darker voids. It provides a topology-oriented texture measurement distinct from general lacunarity or global entropy. |
| 333 | tau network euler characteristic density | code | Thresholds cytoplasmic Tau signal using an adaptive within-cell threshold and computes the Euler characteristic of the resulting Tau-positive binary network, normalized by cytoplasmic area. Positive values reflect many disconnected Tau components, while negative values reflect connected mesh-like structures with holes. This provides a topological view of Tau organization that is related to but distinct from measuring only total mesh hole area. |
| 334 | tau superlevel hole lifetime mean | code | Measures the average threshold persistence of dark holes enclosed by high-Tau regions inside the cell. Across multiple Tau intensity thresholds, holes within superlevel Tau masks are detected, matched approximately by overlap, and their lifetimes across thresholds are averaged. This feature targets honeycomb-like or reticular Tau organization distinct from simple aggregate area or mesh-hole fraction. |
| 335 | cytosolic tau void size heterogeneity score | vlm | Scores the visual heterogeneity of dark Tau-poor voids or cavities within the cell cytoplasm, excluding the nucleus, in the single Tau-channel 2D MIP. The feature focuses on whether the cytoplasmic Tau texture contains similarly sized gaps or a broad mixture of small and large dark spaces. This complements overall lacunarity by emphasizing perceived variation in void scale rather than only the presence of gaps. |
| 336 | nuclear tau gap peninsula ruggedness score | vlm | Measures how irregular the Tau-negative or Tau-poor nuclear gap appears due to jagged Tau protrusions, peninsulas, or bays extending along its boundary. This feature focuses on rugged boundary interdigitation around the nuclear clearing rather than simple nuclear Tau intrusion or exclusion. Higher values indicate a more serrated nucleus-adjacent Tau interface, which may reflect pressure from dense somatic inclusions or uneven perinuclear aggregation. |
| 337 | nucleus adjacent tau void penetration score | vlm | Measures whether Tau-poor voids or dark channels extend from the nuclear boundary outward into the surrounding cytoplasm in the 2D Tau MIP. A high score indicates clear penetration of low-signal corridors from the nucleus into Tau-rich cytoplasmic regions, potentially reflecting displacement, exclusion, or compartmental disruption. This complements perinuclear enrichment features by focusing on negative-space intrusions rather than bright Tau accumulation. |
| 338 | reticular honeycomb soma score | vlm | Visual scalar score for a reticular or honeycomb-like Tau pattern in the soma, where bright Tau-positive ridges surround darker holes or excluded compartments. Higher values indicate a clear mesh-like, porous, or lattice-like somatic organization rather than uniform diffuse signal or isolated compact puncta. Such patterns have been described in super-resolution Tau imaging and may reflect dense pathological organization within the cell body. |
| 339 | reticular honeycomb tau pattern score | vlm | Scores the presence of a reticular, mesh-like, or honeycomb-like Tau pattern within the cell body or dense Tau regions of the 2D MIP. A high score indicates bright interconnected ridges surrounding darker holes or compartments rather than uniform diffuse fluorescence. This may capture super-resolution-visible dense Tau network organization that is not equivalent to simple intensity or puncta count. |
| 340 | reticular mesh wall thickness cv score | vlm | Measures the apparent variability in thickness of the bright Tau-positive walls forming a somatic reticular or honeycomb-like mesh. Unlike pore-shape alignment, this feature focuses on whether the mesh walls are uniformly thin, locally swollen, or highly uneven across the soma. High values may indicate remodeling from a fine physiological network toward heterogeneous aggregated Tau strands. |
| 341 | reticular pore coalescence front asymmetry score | vlm | Evaluates whether dark pores within a somatic reticular Tau mesh appear to merge into larger voids along one side or front of the soma rather than being evenly distributed. The feature focuses on asymmetric coalescence of pore spaces in the 2D Tau projection, not simply pore size or wall thickness. Higher values indicate a polarized front of pore enlargement, suggesting spatially heterogeneous reticular remodeling. |
| 342 | somatic dark pore rim enhancement score | vlm | A scalar visual score for dark intracellular pore-like voids whose edges are outlined by brighter Tau signal inside the soma. This differs from a general honeycomb score by emphasizing rim contrast around individual dark cavities rather than overall mesh abundance. It may reflect organelle-exclusion or reticular Tau organization associated with dense pathological somatic remodeling. |
| 343 | somatic honeycomb mesh score | vlm | A continuous score measuring whether dense somatic Tau signal has a mesh-like or honeycomb-like texture, with bright walls surrounding darker holes or cavities. High values indicate sponge-like, reticular, or internally patterned Tau aggregates rather than uniform blobs. This may capture super-resolution-visible organization within dense pathological Tau regions. |

| # | Feature name | Method | Description |
|---|---|---|---|
| 344 | somatic reticular pore radial maturation score | vlm | Scores whether the somatic Tau reticular pattern shows a radial organization of dark pores, such as small irregular pores near the nucleus and larger or more ordered pores toward the soma edge, or the reverse. This is measured visually on the 2D Tau MIP, using soma and nucleus context to judge radial pore zonation rather than overall pore alignment. It may capture a distinct stage of somatic tangle maturation and Tau mesh remodeling. |
| 345 | tau void partitioning barrier score | vlm | Scores the extent to which Tau-poor dark voids form barrier-like corridors that partition the cell into separate Tau-rich domains. A high value indicates connected dark channels or clefts that split aggregated Tau regions, not merely isolated small holes. This may capture advanced spatial reorganization of cytoplasmic Tau into separated inclusion territories. |

#### Morphology-defined biological process · Morphomechanics

**Description.** This feature class measures the influence of Tau-enriched signal on the nucleus, cell boundary, cell area, cytoplasmic thickness, and cell–nucleus geometric offset. It captures cytoskeletal load, cytoplasmic crowding, and abnormalities in cellular morphomechanics.

13 features · 3 subcellular architectures

##### Subcellular architecture · Nuclear and nucleocytoplasmic compartmentalization scale (7 features)

**Approximate scale.** Typical mammalian nucleus diameter approximately 5-10 um.

| # | Feature name | Method | Description |
|---|---|---|---|
| 346 | cell nucleus geometric offset ratio | vlm | Scores the normalized displacement of the nucleus centroid from the overall cell-mask centroid in the 2D Tau MIP, using the nucleus and cell masks. This captures cell-body asymmetry or nuclear eccentric positioning that may accompany severe intracellular Tau reorganization, while being distinct from Tau-mass-based displacement features. The scalar should increase when the nucleus is visibly off-center relative to the segmented cell body/cytoplasm. |
| 347 | nuclear cell axis mirror tau asymmetry score | vlm | Scores how unevenly Tau signal is distributed on the two sides of the axis connecting the nucleus centroid to the cell centroid in the 2D MIP. A high value indicates that Tau aggregates, haze, or filaments preferentially occupy one lateral half of the cell relative to this biologically meaningful polarity axis, rather than being symmetrically arranged. This complements simple centroid-offset or projection features by judging bilateral pattern imbalance around the nucleus-cell geometry. |
| 348 | nuclear displacement by tau inclusion score | vlm | Scores how strongly the nucleus appears displaced from the center of the cell in association with nearby dense Tau-positive inclusions or bundles in the 2D Tau MIP. A high value indicates an eccentric nucleus adjacent to or compressed by a dominant Tau aggregate mass, a visual signature of advanced somatic Tau pathology. This complements general NFT burden by focusing on spatial deformation of the nucleus-cell geometry rather than aggregate brightness alone. |
| 349 | nuclear displacement by tau mass score | vlm | Scores whether the nucleus appears visually displaced, compressed, or pushed toward the cell edge in association with a bright Tau-rich inclusion or mass. The feature is based on the spatial relationship between the nucleus, cell body, and dominant Tau accumulation in the 2D MIP. A higher score suggests a large asymmetric intracellular Tau burden that may reflect advanced aggregate formation. |
| 350 | nucleus aggregate axis torque score | vlm | Measures the angular mismatch between the dominant orientation of large somatic Tau aggregates and the apparent nucleus displacement or nuclear clearing axis. Rather than only detecting nuclear displacement, this feature asks whether Tau inclusions appear to exert an off-axis, twisting, or torque-like spatial relationship around the nucleus. High values may mark severe asymmetric aggregate growth and mechanical reorganization of the soma. |
| 351 | nucleus cell axis tau projection score | vlm | Scores how strongly the Tau signal is visually projected along the geometric axis defined by the nucleus position relative to the whole-cell mask, rather than being radially balanced around the nucleus. In the 2D Tau MIP, the VLM should judge whether bright Tau mass lies preferentially on the cell side away from or toward the displaced nucleus, using the provided nucleus and cell masks. This complements simple nucleus-cell offset by measuring whether Tau intensity is directionally organized along that offset axis. |
| 352 | nucleus cytoplasm area balance score | vlm | Scores the apparent balance between nuclear area and available cytoplasmic cell area using the nucleus and cell masks in the 2D MIP. Higher values indicate a relatively large nucleus or reduced cytoplasmic compartment, while lower values indicate abundant cytoplasm around the nucleus. This geometry-based feature may reflect cell state, degeneration, or stress-associated morphology independent of raw Tau intensity. |

##### Subcellular architecture · Cell boundary, cortical, peripheral-shell, and extracellular-neighborhood scale (3 features)

**Approximate scale.** Set against a whole-cell diameter of ~10-100 um; the measured region is typically a thin band or boundary arc inside/outside the cell edge.

| # | Feature name | Method | Description |
|---|---|---|---|
| 353 | cell boundary tau indent pressure score | vlm | Scores whether dense Tau masses, bundles, or aggregate-rich zones appear to press against or locally indent the cell boundary in the 2D MIP. A high value indicates that Tau structures visually crowd the cell cortex and create asymmetric boundary deformation, narrowing, or protrusive pressure. This may reflect severe intracellular Tau accumulation affecting cellular shape rather than merely peripheral Tau enrichment. |

| # | Feature name | Method | Description |
|---|---|---|---|
| 354 | cell boundary tau retraction mantle score | vlm | Scores the presence of a relatively continuous Tau-poor mantle immediately inside the cell boundary, suggesting that Tau structures are retracted toward the interior rather than extending to the periphery. A high value corresponds to a clear peripheral cytoplasmic rim with little Tau signal around much of the cell outline. This complements peripheral enrichment features by measuring the opposite spatial organization: inward withdrawal of Tau structures. |
| 355 | cell edge tau budding protrusion score | vlm | Measures localized Tau-positive buds or protrusions extending from the cell boundary. A high score indicates discrete outward bulges or knobs of Tau signal at the cell periphery, rather than a smooth rim or diffuse leakage. This may indicate peripheral aggregate extrusion, neurite initiation sites, or localized membrane-associated Tau accumulation. |

##### Subcellular architecture · Whole-cell geometry, global polarity, and spatial-distribution scale (3 features)

**Approximate scale.** Approximately 10-100+ um; larger if the full neurite arbor is included in the 2D projection.

| # | Feature name | Method | Description |
|---|---|---|---|
| 356 | cell mask process arborization score | vlm | Scores how extensively the segmented single-cell mask extends into branched Tau-positive processes rather than remaining as a compact soma-dominated shape. It uses the 2D Tau MIP together with the cell mask to judge visible process number, branching, and spatial spread. This may capture neuronal structural preservation or collapse, which can accompany Tau pathology and patient-specific transcriptional states. |
| 357 | cellular tau failure collapse score | vlm | A scalar visual score for an end-stage collapsed-cell appearance in which normal neuronal architecture is hard to recognize and Tau appears as a compact, tangled, or debris-like mass replacing the cell. It should be high when the image suggests loss of intact soma-neurite organization rather than merely strong intracellular staining. This feature targets severe degeneration complementary to extracellular ghost-tangle likelihood and somatic inclusion severity. |
| 358 | tau positive cell area utilization score | vlm | Scores the fraction of the segmented cell area that appears visibly occupied by Tau fluorescence in the single-channel 2D MIP. High values indicate broad Tau coverage across soma and processes, whereas low values indicate Tau restricted to small objects or limited subregions. This captures spatial utilization of the cell by Tau signal rather than only brightness or aggregate prominence. |

#### Morphology-defined biological process · Extracellular release and spreading

**Description.** This feature class quantifies extracellular Tau-positive signal, orphan deposits or residual structures, ghost/remnant tangle-like image patterns, and extracellular skeleton complexity. It reflects Tau release, extracellular persistence, and intercellular spreading-related processes while avoiding a direct claim of Tau aggregate formation in the cell images.

5 features · 1 subcellular architectures

##### Subcellular architecture · Cell boundary, cortical, peripheral-shell, and extracellular-neighborhood scale (5 features)

**Approximate scale.** Set against a whole-cell diameter of ~10-100 um; the measured region is typically a thin band or boundary arc inside/outside the cell edge.

| # | Feature name | Method | Description |
|---|---|---|---|
| 359 | extracellular tau aggregate burden fraction | code | Quantifies the fraction of high-intensity Tau aggregate signal located outside the segmented cell mask but within the image field. It is computed by detecting bright Tau-positive connected components outside the cell and dividing their integrated intensity by total image Tau or total detected aggregate intensity. This may capture extracellular ghost-tangle-like remnants or released Tau pathology, while remaining distinct from intracellular somatic aggregate burden. |
| 360 | extracellular tau skeleton complexity index | code | Measures the fibrillar complexity of high-intensity Tau-positive material located outside the segmented cell by skeletonizing extracellular super-threshold Tau objects and combining skeleton length and branch-point density normalized by object area. It is designed to distinguish diffuse extracellular background from ghost-tangle-like fibrillar remnants, complementing simple extracellular burden measurements. The feature uses the cell mask to define extracellular space and can ignore regions overlapping nuclei if a nucleus mask is available. |
| 361 | extracellular ghost tangle likelihood | vlm | Visual scalar estimate of Tau-positive tangle-like material outside the intact cell boundary, such as extracellular dense fibrillar remnants or ghost-tangle-like structures. Higher values indicate prominent Tau-rich structures not clearly contained within the segmented cell body or neurites. This feature captures advanced or degenerative Tau phenotypes while distinguishing them from intracellular aggregates. |
| 362 | extracellular orphan tau deposit score | vlm | Scores the presence and prominence of Tau-positive deposits outside the segmented cell boundary in the 2D MIP. The score should be high when bright aggregate-like Tau structures appear detached from the main cell and not obviously part of a continuous neurite or cell body. This may capture ghost-tangle-like or extracellular Tau-positive remnants while remaining a scalar visual feature. |
| 363 | extracellular tau remnant score | vlm | A continuous score estimating the presence of Tau-positive aggregates or tangled remnants outside the segmented cell boundary. High values indicate bright extracellular Tau deposits, ghost-tangle-like material, or detached aggregates lacking clear association with the intact cell body or nucleus. This feature captures possible degenerative Tau remnants or extracellular aggregate burden. |

#### Additional MorphAgent catalog features not assigned to the Fig. 5 morphology hierarchy

37 features are present in the full MorphAgent Tau catalog (n = 400) but were not members of the 11 morphology-defined biological processes used for Fig. 5.

| # | Feature name | Method | Description |
|---|---|---|---|
| 364 | bundle thickness variability | code | Measures spatial variability in the local thickness of Tau bundle structures using the distance transform sampled along the bundle skeleton. High values indicate unevenly thickened or nodular bundles, while low values indicate bundles of relatively uniform width. This provides information about bundle maturation or compaction beyond orientation coherence or aggregate eccentricity. |
| 365 | cytoplasmic tau highpass radial slope | code | Computes the radial slope of high-frequency Tau texture energy from the nuclear boundary toward the cell periphery within the cytoplasm. The Tau MIP is smoothed to estimate diffuse signal, positive high-pass residual energy is binned by normalized distance from nucleus to cell boundary, and a robust slope is fitted across radial bins. This captures whether granular Tau texture is concentrated perinuclearly, broadly cytoplasmic, or peripheral, which may reflect mislocalized somatic Tau and punctate stress-granule-like organization. |
| 366 | cytoplasmic tau radial frequency entropy | code | Computes the mean Tau intensity in concentric cytoplasmic shells from the nucleus outward to the cell boundary on the 2D MIP, detrends this radial profile, and summarizes the normalized Fourier power spectrum by spectral entropy. Low values indicate a dominant radial oscillation or sharp banding, while high values indicate complex multi-scale radial redistribution. This complements radial transition and enrichment features by measuring frequency composition rather than amplitude or monotonicity. |
| 367 | droplet size dispersion index | code | Uses the provided droplet mask to compute the coefficient of variation of segmented Tau droplet areas in the 2D MIP. Higher values indicate a mixture of small puncta and larger inclusions, suggesting heterogeneous aggregation states. This complements aggregate density and dominant-aggregate occupancy by focusing on the spread of object sizes rather than their total amount. |
| 368 | droplet spatial clustering index | code | Quantifies whether segmented Tau droplets are spatially clustered within the cell by computing the mean nearest-neighbor distance between droplet centroids normalized by the expected spacing from cell area and droplet count. Lower values indicate tighter clustering of puncta, while higher values indicate dispersed droplets; if fewer than two droplets are present, the feature can be set to zero. This captures spatial organization of punctate Tau beyond simple droplet number or intensity. |
| 369 | droplet spatial clustering score | code | Quantifies whether Tau droplets are spatially clustered within the cell by comparing observed nearest-neighbor distances among droplet centroids to the expected spacing for the same droplet count and cell area. Positive values indicate droplets are closer together than expected from a random spatial distribution, while values near zero indicate weak clustering or too few droplets. This captures aggregate organization independent of simple droplet count or area fraction. |
| 370 | droplet tau core rim log ratio | code | For segmented droplet regions, compares Tau signal in droplet cores versus droplet rims in the 2D Tau MIP. Each droplet mask is split by its internal distance transform into a central core and peripheral rim, and the log ratio of mean Tau intensity in core versus rim is aggregated across droplets. This captures whether Tau is internally concentrated, rim-recruited, or excluded from droplet interiors. |
| 371 | droplet tau load area allometric exponent | code | Measures how Tau loading scales with droplet size using the provided droplet mask. Droplet components are identified from mask_droplet.tif, integrated Tau intensity and area are computed for each component, and the feature is the slope of log integrated Tau versus log droplet area. This captures whether larger droplets disproportionately recruit Tau, a biologically relevant condensate-loading property distinct from total droplet Tau fraction. |
| 372 | filament branchpoint density | code | Computes the number of branch points in the skeletonized mask_filament divided by total filament skeleton length. This captures network-like or tangled filament complexity rather than just filament amount. Increased branching or reticular organization may reflect advanced fibrillar Tau aggregation or tangled neuritic pathology. |
| 373 | filament branchpoint frequency | code | Computes the number of skeleton branch points in the Tau filament mask normalized by total filament skeleton length. A high value indicates network-like, branched, or tangled filament organization, whereas a low value indicates mostly unbranched linear filaments. This is relevant to distinguishing simple thread-like Tau structures from complex fibrillar networks. |
| 374 | high tau peak mst edge skewness | code | Measures spatial heterogeneity in the spacing of high Tau local maxima inside the segmented cell. Local maxima are detected after mild smoothing and robust thresholding, a minimum spanning tree is built over their coordinates, and the skewness of normalized MST edge lengths is reported. High values indicate mixtures of dense hotspot clusters and isolated Tau aggregates, which may reflect heterogeneous aggregation or stress-granule-like puncta organization. |
| 375 | nuclear centroid displacement index | code | Measures the distance between the nucleus centroid and the whole-cell centroid, normalized by the equivalent cell radius derived from cell area. This feature uses only the 2D cell and nucleus masks and is independent of Tau intensity. Nuclear displacement may reflect severe cytoplasmic Tau inclusion burden or asymmetric aggregate pressure within the soma. |

| # | Feature name | Method | Description |
|---|---|---|---|
| 376 | perinuclear tau angular autocorrelation halfwidth | code | Measures the angular coherence scale of Tau intensity in a narrow cytoplasmic annulus surrounding the nucleus on the 2D MIP. The annular Tau profile is sampled by angle around the nuclear centroid, autocorrelated circularly, and summarized as the angular lag where autocorrelation first falls below half its zero-lag value. Broad perinuclear Tau caps and continuous rings produce larger values, whereas fragmented punctate perinuclear Tau produces smaller values. |
| 377 | tau low signal void chord anisotropy | code | Inside the cell, identify low-Tau void regions using a lower-intensity quantile threshold and measure chord lengths through those voids at multiple orientations. The feature is the anisotropy ratio between the largest and smallest robust median chord length across orientations. It measures whether Tau-poor spaces are isotropic holes or elongated channels, providing a complementary view of mesh-like and exclusion patterns. |
| 378 | tau radial gradient slope from nucleus | code | Computes the robust slope of mean Tau intensity as a function of distance from the nucleus within the cell mask. Distances are measured in the 2D MIP from the nuclear boundary or centroid, and intensity is averaged in radial distance bins. A strongly negative slope indicates soma/perinuclear enrichment, while a flatter or positive slope suggests more distal or neuritic Tau distribution. |
| 379 | tau reticular saddlepoint density | code | This feature detects saddle-like points in the smoothed cytoplasmic Tau intensity surface using the Hessian determinant and normalizes their count by cytoplasmic area. A high density of saddle points indicates a reticular or tangled mesh with alternating ridges and holes, whereas a low density indicates smoother diffuse signal or isolated compact aggregates. This may capture honeycomb-like or fibrillar Tau organization in super-resolution MIP images. |
| 380 | tau superlevel boundary complexity auc | code | For Tau superlevel masks inside the cell at several upper-intensity quantiles, compute the normalized boundary complexity perimeter squared divided by area. The scalar is the area under this boundary-complexity curve across thresholds. It measures whether high-Tau material forms smooth compact inclusions or jagged fragmented aggregate fronts, which may reflect different Tau aggregation states. |
| 381 | tau top decile nuclear distance shift | code | Computes the normalized shift in distance from the nuclear boundary for the brightest 10% of cytoplasmic Tau pixels relative to all cytoplasmic pixels. Negative values indicate preferential perinuclear or somatic concentration of the brightest Tau, while positive values indicate brighter Tau toward distal cytoplasm or processes. This complements centroid-based features by focusing specifically on the spatial placement of the high-intensity Tau tail. |
| 382 | beaded filament fragmentation score | vlm | Scores the degree to which Tau-positive filamentous structures appear broken into bead-like segments or string-of-pearls patterns rather than smooth continuous fibers. The score is assessed on the 2D MIP using visible Tau signal and filament masks when present. A higher value indicates stronger fragmentation or beadiness of Tau along neurite-like or filament-like structures. |
| 383 | bundle midspan unraveling score | vlm | Measures how much Tau bundles appear to loosen or split into multiple thinner strands along their middle portions rather than only at their ends. A high score indicates mid-bundle fraying, partial separation, or unraveling within the bundle shaft. This complements terminal fan-out and lateral shedding concepts by focusing specifically on internal bundle disassembly. |
| 384 | cell quadrant tau load imbalance score | vlm | Measures the visual imbalance of Tau burden across four approximate quadrants of the segmented single cell in the 2D MIP, using the nucleus or cell centroid as the reference point. A high score indicates that Tau intensity, aggregates, or fibrillar mass are concentrated in one sector rather than distributed evenly. This captures coarse subcellular polarization without assuming a predefined axon or dendrite identity. |
| 385 | cell shape tau axis decoupling score | vlm | Measures the mismatch between the dominant orientation of the whole cell shape and the dominant orientation of Tau filaments or bundles. A high score indicates that Tau structures run obliquely, transversely, or in multiple directions relative to the cell's long axis rather than aligning with overall cell polarity. This may capture loss of organized neuronal Tau architecture and altered cytoskeletal alignment. |
| 386 | condensate coarsening stage score | vlm | Estimates the apparent Tau condensate maturation state from many fine puncta toward fewer larger rounded droplets or coalesced bodies within the 2D cell projection. Low values represent dispersed small granules, while high values represent visually coarsened droplet-like aggregates. This may reflect phase-separation-like aggregation behavior relevant to pathological Tau and stress-granule-associated phenotypes. |
| 387 | cytoplasmic fibril free halo asymmetry score | vlm | Measures whether fibrillar Tau is unevenly excluded from one side of the cytoplasm, creating an asymmetric fibril-free halo or cleared region inside the cell mask. A high score indicates that Tau filaments and bundles occupy one portion of the cytoplasm while another similarly available cytoplasmic region remains visually depleted. This may capture polarized redistribution or compartmental clearing not fully described by total Tau intensity. |
| 388 | dense bundle nft like burden | vlm | A continuous score estimating the burden of dense, tangled, bundle-like Tau structures within the cell. High values indicate large compact bundles, tangled masses, or neurofibrillary-tangle-like conglomerates occupying a substantial fraction of the cell body or cytoplasm. This feature targets advanced Tau aggregation beyond simple diffuse intensity. |
| 389 | distal process punctate burden visual | vlm | Visual score estimating the burden of bright Tau puncta or small aggregates in distal neurite-like regions relative to the soma and proximal processes. Higher values indicate many puncta concentrated near neurite tips or far from the cell body, even if proximal regions remain relatively smooth or dim. This is relevant because some Tau pathology can emerge first in distal axonal regions before strong somatic aggregation. |

| # | Feature name | Method | Description |
|---|---|---|---|
| 390 | droplet pearling on bundle edges score | vlm | Scores the extent to which small rounded Tau droplets or puncta decorate the edges of thicker Tau bundles like pearls along a boundary. High values indicate edge-localized bead formation rather than droplets randomly scattered in cytoplasm or embedded uniformly inside bundles. This may capture nucleation of condensate-like Tau objects at fibrillar aggregate interfaces. |
| 391 | filament fragmentation beading score | vlm | A continuous score for how discontinuous, beaded, or broken the Tau-positive filamentous structures appear in the 2D MIP. Low values indicate smooth continuous strands, while high values indicate string-of-pearls morphology, interrupted segments, or many bright beads along thin Tau tracks. This captures loss of organized Tau continuity and early aggregate nucleation along processes. |
| 392 | filament mesh chord irregularity score | vlm | Scores variability in the apparent spacing between neighboring Tau mesh walls or filament chords within network-like regions of the MIP. High values indicate an irregular mesh with widely varying gap widths, while low values indicate more uniform spacing or absence of mesh-like organization. This may reflect heterogeneous remodeling of dense somatic or cytoplasmic Tau networks. |
| 393 | filament parallel track lane switch score | vlm | Scores whether Tau filaments within elongated structures run as parallel tracks that repeatedly cross, merge, or exchange relative positions along their length. Low values correspond to straight, consistently separated lanes, while high values indicate woven or lane-switching fibrillar organization. This may reflect altered filament packing or pathological remodeling of Tau assemblies. |
| 394 | nuclear moat breach frequency score | vlm | Measures the visual frequency with which Tau filaments, bundles, droplets, or dense patches cross or interrupt the normally Tau-poor gap around the nucleus. A high score indicates repeated breaches of a perinuclear clearance zone rather than uniform nuclear exclusion. This may capture pathological Tau encroachment on nuclear-adjacent cytoplasm in a way distinct from global nuclear Tau exclusion. |
| 395 | perinuclear microbead collar score | vlm | Scores how strongly small Tau-positive puncta or droplet-like objects form a discontinuous collar immediately outside the nuclear boundary in the 2D Tau MIP. This differs from a smooth perinuclear ring by emphasizing bead-like, granular decoration along the nucleus-facing cytoplasm. It may reflect stress-granule-like or early aggregate recruitment around the perinuclear compartment. |
| 396 | reticular pore shape alignment score | vlm | Measures whether dark pores or holes within reticular somatic Tau patterns have aligned, elongated shapes rather than random round cavities. A high score indicates that pore shapes share a common orientation or flow direction within the Tau mesh. This complements honeycomb or lacunarity concepts by focusing specifically on orientation coherence of the negative spaces inside the Tau reticulum. |
| 397 | tau depletion shadow around droplets score | vlm | Scores whether segmented droplets are bordered by asymmetric local Tau-depleted shadows or clear zones in the surrounding cytoplasm. The VLM should compare Tau intensity immediately around droplet masks with nearby non-droplet cytoplasm and judge whether droplets appear to repel or exclude Tau on one side or around their perimeter. If droplet masks are absent or no droplets are visible, the feature should be scored near zero rather than treated as missing. |
| 398 | tau object nested enclosure score | vlm | Scores how often smaller Tau droplets, puncta, or short fragments appear visually enclosed within larger looped, ring-like, or cage-like Tau structures. High values indicate nested aggregate organization, where one Tau object class is contained inside another Tau-defined boundary. This may reflect complex tangle maturation, compartmental trapping, or higher-order aggregate assembly. |
| 399 | tau percolating island dominance score | vlm | Visual scalar score for whether Tau-positive signal is dominated by one large connected island spanning soma and processes versus many separated islands or speckles. This captures the percolation-like organization of Tau domains in the 2D MIP, complementing simple connectivity by emphasizing dominance of a single contiguous Tau compartment. |
| 400 | tau polarization asymmetry score | vlm | A continuous score measuring whether Tau intensity and aggregates are distributed symmetrically around the cell or concentrated in one pole, one side of the soma, or one major process. High values indicate strong spatial polarization or eccentric aggregation relative to the cell and nucleus. This may reflect asymmetric Tau trafficking, localized pathology, or nuclear displacement by aggregates. |
