## Supplementary feature list 5 for "Biologically grounded cell profiling across microscopy modalities"

### MorphAgent nucleus-associated size and shape features

| # | Feature name |
|---|---|
| 1 | cell nucleus area ratio mean |
| 2 | cyto nucleus area ratio mean |
| 3 | cytoplasm nucleus area ratio |
| 4 | cytoplasm to nucleus area ratio |
| 5 | cytoskeleton nucleus area ratio |
| 6 | dapi area coefficient of variation |
| 7 | dapi boundary intensity gradient |
| 8 | dapi boundary intensity mean |
| 9 | dapi boundary intensity ratio |
| 10 | dapi boundary irregularity |
| 11 | dapi boundary irregularity index |
| 12 | dapi boundary roughness |
| 13 | dapi boundary sharpness mean |
| 14 | dapi high intensity area fraction mean |
| 15 | dapi nuclear area mean |
| 16 | dapi nuclear boundary intensity ratio |
| 17 | dapi nuclear circularity mean |
| 18 | dapi solidity mean |
| 19 | nuclear area cv |
| 20 | nuclear boundary intensity gradient |
| 21 | nuclear boundary solidity mean |
| 22 | nuclear circularity median |
| 23 | nuclear eccentricity std |
| 24 | nuclear eccentricity variance |
| 25 | nuclear form factor mean |
| 26 | nuclear major axis orientation coherence |
| 27 | nuclear roundness mean |
| 28 | nuclear shape circularity mean |
| 29 | nuclear shape eccentricity mean |
| 30 | nuclear shape factor form factor |
| 31 | nuclear shape irregularity index |
| 32 | nuclear shape irregularity mean |
| 33 | nuclear shape roundness mean |
| 34 | nucleus area mean |
| 35 | nucleus area median |
| 36 | nucleus boundary roughness |
| 37 | nucleus boundary roughness mean |
| 38 | nucleus cell area ratio mean |
| 39 | nucleus compactness mean |
| 40 | nucleus cytoplasm area ratio |
| 41 | nucleus cytoplasm area ratio mean |
| 42 | nucleus eccentricity mean |
| 43 | nucleus major axis length |
| 44 | nucleus major axis orientation entropy |
| 45 | nucleus shape form factor mean |
| 46 | nucleus to cell area ratio mean |
| 47 | nucleus to cytoplasm area ratio |
| 48 | vlm nuclear shape irregularity |
