## Supplementary feature list 6 for "Biologically grounded cell profiling across microscopy modalities"

### CellProfiler nucleus-associated size and shape features

| # | Feature name |
|---|---|
| 1 | Mean_Nuclei_AreaShape_Area |
| 2 | Mean_Nuclei_AreaShape_BoundingBoxArea |
| 3 | Mean_Nuclei_AreaShape_BoundingBoxMaximum_X |
| 4 | Mean_Nuclei_AreaShape_BoundingBoxMaximum_Y |
| 5 | Mean_Nuclei_AreaShape_BoundingBoxMinimum_X |
| 6 | Mean_Nuclei_AreaShape_BoundingBoxMinimum_Y |
| 7 | Mean_Nuclei_AreaShape_Center_X |
| 8 | Mean_Nuclei_AreaShape_Center_Y |
| 9 | Mean_Nuclei_AreaShape_Compactness |
| 10 | Mean_Nuclei_AreaShape_Eccentricity |
| 11 | Mean_Nuclei_AreaShape_EquivalentDiameter |
| 12 | Mean_Nuclei_AreaShape_Extent |
| 13 | Mean_Nuclei_AreaShape_FormFactor |
| 14 | Mean_Nuclei_AreaShape_MajorAxisLength |
| 15 | Mean_Nuclei_AreaShape_MaxFeretDiameter |
| 16 | Mean_Nuclei_AreaShape_MaximumRadius |
| 17 | Mean_Nuclei_AreaShape_MeanRadius |
| 18 | Mean_Nuclei_AreaShape_MedianRadius |
| 19 | Mean_Nuclei_AreaShape_MinFeretDiameter |
| 20 | Mean_Nuclei_AreaShape_MinorAxisLength |
| 21 | Mean_Nuclei_AreaShape_Orientation |
| 22 | Mean_Nuclei_AreaShape_Perimeter |
| 23 | Mean_Nuclei_AreaShape_Solidity |
| 24 | Mean_Nuclei_AreaShape_Zernike_0_0 |
| 25 | Mean_Nuclei_AreaShape_Zernike_1_1 |
| 26 | Mean_Nuclei_AreaShape_Zernike_2_0 |
| 27 | Mean_Nuclei_AreaShape_Zernike_2_2 |
| 28 | Mean_Nuclei_AreaShape_Zernike_3_1 |
| 29 | Mean_Nuclei_AreaShape_Zernike_3_3 |
| 30 | Mean_Nuclei_AreaShape_Zernike_4_0 |
| 31 | Mean_Nuclei_AreaShape_Zernike_4_2 |
| 32 | Mean_Nuclei_AreaShape_Zernike_4_4 |
| 33 | Mean_Nuclei_AreaShape_Zernike_5_1 |
| 34 | Mean_Nuclei_AreaShape_Zernike_5_3 |
| 35 | Mean_Nuclei_AreaShape_Zernike_5_5 |
| 36 | Mean_Nuclei_AreaShape_Zernike_6_0 |
| 37 | Mean_Nuclei_AreaShape_Zernike_6_2 |
| 38 | Mean_Nuclei_AreaShape_Zernike_6_4 |
| 39 | Mean_Nuclei_AreaShape_Zernike_6_6 |
| 40 | Mean_Nuclei_AreaShape_Zernike_7_1 |
| 41 | Mean_Nuclei_AreaShape_Zernike_7_3 |
| 42 | Mean_Nuclei_AreaShape_Zernike_7_5 |
| 43 | Mean_Nuclei_AreaShape_Zernike_7_7 |
| 44 | Mean_Nuclei_AreaShape_Zernike_8_0 |
| 45 | Mean_Nuclei_AreaShape_Zernike_8_2 |
| 46 | Mean_Nuclei_AreaShape_Zernike_8_4 |
| 47 | Mean_Nuclei_AreaShape_Zernike_8_6 |
| 48 | Mean_Nuclei_AreaShape_Zernike_8_8 |
| 49 | Mean_Nuclei_AreaShape_Zernike_9_1 |
| 50 | Mean_Nuclei_AreaShape_Zernike_9_3 |
| 51 | Mean_Nuclei_AreaShape_Zernike_9_5 |
| 52 | Mean_Nuclei_AreaShape_Zernike_9_7 |
| 53 | Mean_Nuclei_AreaShape_Zernike_9_9 |
| 54 | Median_Nuclei_AreaShape_Area |
| 55 | Median_Nuclei_AreaShape_BoundingBoxArea |
| 56 | Median_Nuclei_AreaShape_BoundingBoxMaximum_X |
| 57 | Median_Nuclei_AreaShape_BoundingBoxMaximum_Y |
| 58 | Median_Nuclei_AreaShape_BoundingBoxMinimum_X |
| 59 | Median_Nuclei_AreaShape_BoundingBoxMinimum_Y |
| 60 | Median_Nuclei_AreaShape_Center_X |
| 61 | Median_Nuclei_AreaShape_Center_Y |
| 62 | Median_Nuclei_AreaShape_Compactness |
| 63 | Median_Nuclei_AreaShape_Eccentricity |
| 64 | Median_Nuclei_AreaShape_EquivalentDiameter |
| 65 | Median_Nuclei_AreaShape_Extent |
| 66 | Median_Nuclei_AreaShape_FormFactor |
| 67 | Median_Nuclei_AreaShape_MajorAxisLength |
| 68 | Median_Nuclei_AreaShape_MaxFeretDiameter |
| 69 | Median_Nuclei_AreaShape_MaximumRadius |
| 70 | Median_Nuclei_AreaShape_MeanRadius |
| 71 | Median_Nuclei_AreaShape_MedianRadius |
| 72 | Median_Nuclei_AreaShape_MinFeretDiameter |
| 73 | Median_Nuclei_AreaShape_MinorAxisLength |
| 74 | Median_Nuclei_AreaShape_Orientation |
| 75 | Median_Nuclei_AreaShape_Perimeter |
| 76 | Median_Nuclei_AreaShape_Solidity |
| 77 | Median_Nuclei_AreaShape_Zernike_0_0 |
| 78 | Median_Nuclei_AreaShape_Zernike_1_1 |
| 79 | Median_Nuclei_AreaShape_Zernike_2_0 |
| 80 | Median_Nuclei_AreaShape_Zernike_2_2 |
| 81 | Median_Nuclei_AreaShape_Zernike_3_1 |
| 82 | Median_Nuclei_AreaShape_Zernike_3_3 |
| 83 | Median_Nuclei_AreaShape_Zernike_4_0 |
| 84 | Median_Nuclei_AreaShape_Zernike_4_2 |
| 85 | Median_Nuclei_AreaShape_Zernike_4_4 |
| 86 | Median_Nuclei_AreaShape_Zernike_5_1 |
| 87 | Median_Nuclei_AreaShape_Zernike_5_3 |
| 88 | Median_Nuclei_AreaShape_Zernike_5_5 |
| 89 | Median_Nuclei_AreaShape_Zernike_6_0 |
| 90 | Median_Nuclei_AreaShape_Zernike_6_2 |
| 91 | Median_Nuclei_AreaShape_Zernike_6_4 |
| 92 | Median_Nuclei_AreaShape_Zernike_6_6 |
| 93 | Median_Nuclei_AreaShape_Zernike_7_1 |
| 94 | Median_Nuclei_AreaShape_Zernike_7_3 |
| 95 | Median_Nuclei_AreaShape_Zernike_7_5 |
| 96 | Median_Nuclei_AreaShape_Zernike_7_7 |
| 97 | Median_Nuclei_AreaShape_Zernike_8_0 |
| 98 | Median_Nuclei_AreaShape_Zernike_8_2 |
| 99 | Median_Nuclei_AreaShape_Zernike_8_4 |
| 100 | Median_Nuclei_AreaShape_Zernike_8_6 |
| 101 | Median_Nuclei_AreaShape_Zernike_8_8 |
| 102 | Median_Nuclei_AreaShape_Zernike_9_1 |
| 103 | Median_Nuclei_AreaShape_Zernike_9_3 |
| 104 | Median_Nuclei_AreaShape_Zernike_9_5 |
| 105 | Median_Nuclei_AreaShape_Zernike_9_7 |
| 106 | Median_Nuclei_AreaShape_Zernike_9_9 |
| 107 | StDev_Nuclei_AreaShape_Area |
| 108 | StDev_Nuclei_AreaShape_BoundingBoxArea |
| 109 | StDev_Nuclei_AreaShape_BoundingBoxMaximum_X |
| 110 | StDev_Nuclei_AreaShape_BoundingBoxMaximum_Y |
| 111 | StDev_Nuclei_AreaShape_BoundingBoxMinimum_X |
| 112 | StDev_Nuclei_AreaShape_BoundingBoxMinimum_Y |
| 113 | StDev_Nuclei_AreaShape_Center_X |
| 114 | StDev_Nuclei_AreaShape_Center_Y |
| 115 | StDev_Nuclei_AreaShape_Compactness |
| 116 | StDev_Nuclei_AreaShape_Eccentricity |
| 117 | StDev_Nuclei_AreaShape_EquivalentDiameter |
| 118 | StDev_Nuclei_AreaShape_Extent |
| 119 | StDev_Nuclei_AreaShape_FormFactor |
| 120 | StDev_Nuclei_AreaShape_MajorAxisLength |
| 121 | StDev_Nuclei_AreaShape_MaxFeretDiameter |
| 122 | StDev_Nuclei_AreaShape_MaximumRadius |
| 123 | StDev_Nuclei_AreaShape_MeanRadius |
| 124 | StDev_Nuclei_AreaShape_MedianRadius |
| 125 | StDev_Nuclei_AreaShape_MinFeretDiameter |
| 126 | StDev_Nuclei_AreaShape_MinorAxisLength |
| 127 | StDev_Nuclei_AreaShape_Orientation |
| 128 | StDev_Nuclei_AreaShape_Perimeter |
| 129 | StDev_Nuclei_AreaShape_Solidity |
| 130 | StDev_Nuclei_AreaShape_Zernike_0_0 |
| 131 | StDev_Nuclei_AreaShape_Zernike_1_1 |
| 132 | StDev_Nuclei_AreaShape_Zernike_2_0 |
| 133 | StDev_Nuclei_AreaShape_Zernike_2_2 |
| 134 | StDev_Nuclei_AreaShape_Zernike_3_1 |
| 135 | StDev_Nuclei_AreaShape_Zernike_3_3 |
| 136 | StDev_Nuclei_AreaShape_Zernike_4_0 |
| 137 | StDev_Nuclei_AreaShape_Zernike_4_2 |
| 138 | StDev_Nuclei_AreaShape_Zernike_4_4 |
| 139 | StDev_Nuclei_AreaShape_Zernike_5_1 |
| 140 | StDev_Nuclei_AreaShape_Zernike_5_3 |
| 141 | StDev_Nuclei_AreaShape_Zernike_5_5 |
| 142 | StDev_Nuclei_AreaShape_Zernike_6_0 |
| 143 | StDev_Nuclei_AreaShape_Zernike_6_2 |
| 144 | StDev_Nuclei_AreaShape_Zernike_6_4 |
| 145 | StDev_Nuclei_AreaShape_Zernike_6_6 |
| 146 | StDev_Nuclei_AreaShape_Zernike_7_1 |
| 147 | StDev_Nuclei_AreaShape_Zernike_7_3 |
| 148 | StDev_Nuclei_AreaShape_Zernike_7_5 |
| 149 | StDev_Nuclei_AreaShape_Zernike_7_7 |
| 150 | StDev_Nuclei_AreaShape_Zernike_8_0 |
| 151 | StDev_Nuclei_AreaShape_Zernike_8_2 |
| 152 | StDev_Nuclei_AreaShape_Zernike_8_4 |
| 153 | StDev_Nuclei_AreaShape_Zernike_8_6 |
| 154 | StDev_Nuclei_AreaShape_Zernike_8_8 |
| 155 | StDev_Nuclei_AreaShape_Zernike_9_1 |
| 156 | StDev_Nuclei_AreaShape_Zernike_9_3 |
| 157 | StDev_Nuclei_AreaShape_Zernike_9_5 |
| 158 | StDev_Nuclei_AreaShape_Zernike_9_7 |
| 159 | StDev_Nuclei_AreaShape_Zernike_9_9 |
