## Supplementary Table 1 for "Biologically grounded cell profiling across microscopy modalities"

**Table 1: Summary of the imaging datasets analysed in this study.** For each dataset we list the sample size and biological labels, the imaging modality and acquisition parameters, the imaging channels and profiling input, and any paired molecular or orthogonal measurements.

| Dataset | Sample size and biological labels | Imaging modality and acquisition | Channels and profiling input | Paired molecular / orthogonal data |
|---|---|---|---|---|
| **(a) BBBC021 Cell Painting benchmark** (wide-field) | 3,552 image-level profiles from 37 compounds spanning 26 mechanism-of-action (MoA) categories in MCF7 cells; 7 MoA groups contained $\geq$2 compounds and were used for same-MoA matching. | Conventional wide-field fluorescence microscopy at 20× on the original BBBC021 platform; 49 plates (weeks 1–10). | Three channels: DNA, $\beta$-tubulin and F-actin. The analysis unit is one image-level feature vector. | Matched L1000 Connectivity Map profiles for the 37 compounds (compound × concentration); MoA and compound labels. |
| **(b) HSC omics-paired discovery dataset** (spinning-disk confocal) | 110 mouse hematopoietic stem cells (48 young, 62 aged) with paired single-cell Smart-seq2 transcriptomes. | Spinning-disk confocal; lateral pixel size 38.2 nm, axial step 200 nm; 200 nm lateral resolution. | Mitochondria (MitoTracker Green, 488 nm) and nucleus (Hoechst, 405 nm); profiling on the mitochondrial-channel maximum-intensity projection (MIP). | Smart-seq2 transcriptomes and young/aged labels used as validation signals (correlation with the top-500 highly variable genes; aging discrimination). |
| **(c) HSC independent validation dataset** (spinning-disk confocal + synthetic low-resolution) | 162 mouse hematopoietic stem cells (99 young, 63 aged); imaging only, for cross-dataset transfer. | Spinning-disk confocal (CSU-X1 Yokogawa on an inverted Olympus IX81, ×100 oil objective); 38.2 nm lateral, 200 nm axial, ≈120 nm lateral resolution; higher image quality than the discovery set. Matched synthetic low-resolution MIPs generated by convolution with a 2D Gaussian kernel (FWHM 200 nm; $\sigma$ ≈84.9 nm / ≈2.22 pixels), yielding a nominal effective lateral resolution of ≈233 nm. | Mitochondria (488 nm); the mitochondrial-channel MIP was analysed for both the confocal (≈120 nm) and the synthetic low-resolution (≈233 nm) views. | No paired transcriptome; young/aged labels used to evaluate transfer of the fixed 25-feature panel. |
| **(d) Tau paired imaging–transcriptomic discovery dataset** (SIM) | 58 SH-SY5Y cells overexpressing wild-type human 0N4R tau-EGFP, each with a paired single-cell Smart-seq2 transcriptome (27,476 genes). | Three-dimensional structured illumination microscopy (HIS-SIM); lateral pixel size 32.5 nm, axial step 300 nm; ≈100 nm reconstructed lateral resolution. | Acquired channels: nucleus, tau-EGFP, mitochondria and microtubule. Tau profiling used the tau-EGFP channel as a two-dimensional MIP. | Smart-seq2 transcriptomes used for morphology–transcriptome association, morphology-guided target-gene selection, bidirectional prediction, and the morphology–gene hierarchy and correspondence. |
| **(e) Tau mutant validation dataset** (SIM + matched wide-field) | 495 COS-7 cells expressing wild-type or mutant human 0N4R tau-EGFP (S320F, S305I, P301L, P301L+S320F, P301S+S320F), with paired super-resolution and wide-field views of the same cells. | HIS-SIM super-resolution (Wiener reconstruction) with matched wide-field images generated by averaging the corresponding raw SIM frames. | tau-EGFP profiling channel; two-dimensional Tau-channel projections analysed for both modalities. | No paired transcriptome; genotype labels used as orthogonal annotations for binary and six-class mutation-discrimination analyses. |
| **(f) BBBC022 Cell Painting benchmark** (wide-field) | Single-plate subset of the public BBBC022 release (plate 20585): 1,026 wide-field Cell Painting fields from U2OS cells, comprising 450 compound-treated images (50 bioactive compounds, nine sites per well) and 576 matched mock controls (64 wells, nine sites each); used for perturbation-detection benchmarks. | Wide-field fluorescence microscopy (Cell Painting assay). | Five fluorescence channels: ER, DNA, mitochondria, AGP/actin and RNA. | Metadata (plate, well, site, compound name, BROAD ID and condition [compound or mock]) used as perturbation labels; the detailed BBBC022 retrieval protocol is provided in the Supplementary Methods. |
