## Supplementary Table 2 for "Biologically grounded cell profiling across microscopy modalities"

| Curated feature | Operational definition | Biological interpretation |
|---|---|---|
| **Whole-cell mean intensity** | Mean Tau fluorescence intensity measured across the entire cell. | Cellular Tau expression; the feature value provides a proxy for total cellular Tau abundance. |
| **Nuclear Tau intensity at the cell body (z-stack)** | Mean Tau fluorescence intensity measured in the nucleus from the z-stack at the cell body. | Nuclear accumulation of Tau; the feature value reports the amount of Tau detected in the nucleus. |
| **Cytoplasmic Tau intensity at the cell body (z-stack)** | Mean Tau fluorescence intensity measured in the cytoplasm from the z-stack at the cell body. | Cytoplasmic Tau pool; the feature value reports the amount of Tau retained in the cytoplasm. |
| **Tau intensity on microtubule bundles** | Mean Tau fluorescence intensity measured on microtubule bundles. | Bundle-associated Tau, reflecting Tau bound to microtubules and associated with Tau-mediated microtubule bundling. |
| **Tau intensity on individual filaments** | Mean Tau fluorescence intensity measured on individual microtubule filaments. | Microtubule-bound Tau; the feature value provides a proxy for Tau occupancy on individual microtubules. |
| **Nuclear/background intensity ratio** | Ratio of nuclear fluorescence intensity to background fluorescence intensity. | Nuclear enrichment of Tau relative to background; the feature value reports nuclear Tau accumulation. |
| **Nuclear/cytoplasmic intensity ratio** | Ratio of nuclear fluorescence intensity to cytoplasmic fluorescence intensity. | Balance between nuclear Tau accumulation and the cytoplasmic Tau pool. |
| **Individual-filament/cytoplasmic intensity ratio** | Ratio of fluorescence intensity on individual microtubule filaments to cytoplasmic fluorescence intensity. | Relative enrichment of Tau on individual microtubules compared with the cytoplasmic Tau pool. |
| **Individual-filament/bundle intensity ratio** | Ratio of fluorescence intensity on individual microtubule filaments to fluorescence intensity on microtubule bundles. | Relative distribution of Tau between individual microtubules and bundled microtubules. |
| **Tau continuity** | Annotator-assisted assessment of the continuity of Tau localization along microtubules. | Morphology of microtubule-bound Tau; the feature value captures the continuity of Tau signal along microtubules. |
| **Bundle width** | Maximum width of the thickest microtubule bundle. | Morphology of Tau-associated microtubule bundles; the feature value quantifies bundle thickness. |
| **Individual-filament abundance** | Annotator-assisted score for the abundance of resolvable individual microtubule filaments. | Microtubule-network morphology; the feature value summarizes the abundance of individual filaments. |
| **Microtubule-organizing center visibility** | Annotator-assisted assessment of whether a microtubule-organizing center (MTOC) is visible. | MTOC organization; the feature value indicates whether an MTOC is detectable in the image. |
| **Droplet number** | Number of Tau liquid-liquid phase separation (LLPS) droplets. | Tau LLPS; the feature value quantifies condensate abundance. |
| **Total droplet area** | Total area of all Tau LLPS droplets. | Tau LLPS; the feature value quantifies the cumulative area occupied by condensates. |
| **Droplet intensity** | Mean fluorescence intensity measured across all Tau LLPS droplets. | Tau LLPS; the feature value quantifies Tau enrichment within condensates. |

*Abbreviations: LLPS, liquid-liquid phase separation; MTOC, microtubule-organizing center.*
